## Supplementary material for "Genomic and chemical decryption of the Bacteroidetes phylum for its potential to biosynthesize natural products": General Supporting Material

**This PDF file includes:**

R-script for chemotype-barcoding matrix
Figs. S1 to S39
Tables S1 to S7

**Other Supplementary Materials for this manuscript include the following:**

Synthesis of hydroxy amino acids
Tab. S8

**R-script for chemotype-barcoding matrix:**

```
library(readr)
library(reshape)
library(ggplot2)

setwd("E:/Stephan/") # Working directory
experiment<-"2020-12-16_chitinophaga_v1" #Experiment name
project<-(paste(c(experiment,""), collapse=""))

presencetable.gesamt<-as.matrix(read_tsv(paste(c(project,".txt"), collapse=""),col_names=TRUE, col_types =
NULL, na = c("", "NA"), trim_ws = FALSE, skip = 0, n_max = Inf, progress = show_progress(), skip_empty_rows
= TRUE)) # load project file
#presencetable.transformed<-t(presencetable.gesamt)
n<-ncol(presencetable.gesamt)
presencetable.ges<-presencetable.gesamt[,2:n]
names<-presencetable.gesamt[,1]
rownames(presencetable.ges)<-names
melt.data<-melt(presencetable.ges)
colnames(melt.data)<-c("Bucket","Condition","Value")
png(filename=paste(c(project,"_test.png"), collapse=""),
    width = 50*400,
    height = 50*400,
    res = 1000,
    pointsize = 5)
qplot(data=melt.data,
          x=Bucket,
          y=Condition,
          fill=factor(Value),
        #   geom="tile")+scale_fill_manual(values=c("more than one"="black", "R2A"="green", "5065"="blue",
"3018"="orange","3021"="pink","5294"="brown","empty"="white", "isolated"="red" ))
          geom="tile")+scale_fill_manual(values=c("A - more than one"="black", "F - R2A"="green", "D -
5065"="blue", "B - 3018"="orange","E - 3021"="pink","C - 5294"="brown","G - empty"="white", "H -
isolated"="red" ))
dev.off()
```

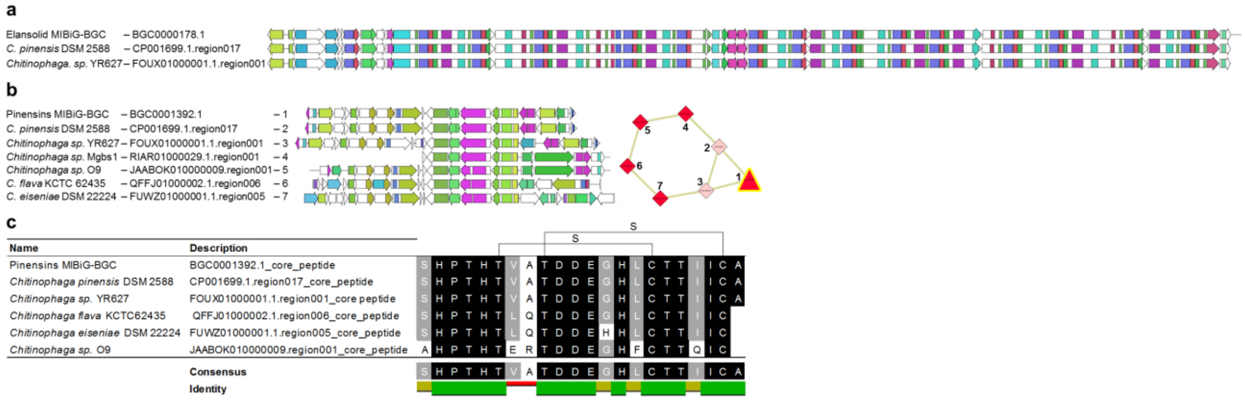


**Fig. S1. Gene cluster families (GCF) of the elansolids and pinensins color coded by PFAM domains. a**, Reference biosynthetic gene cluster of the elansolids deposited at MIBiG and two further identical BGCs identified on genomes of different strains. **b**, Reference biosynthetic gene cluster of the pinensins deposited at MIBiG and six modified derivatives found on genomes of different strains are depicted with the PFAM domains color coded and the gene cluster family shown to the right. BiG-SCAPE results with a cutoff of 0.6. **c**, Clustal W alignment[101] of the pinensin core peptide with those found in its GCF. No core peptide was found on the contig of *Chitinophaga sp.* Mgbs1 (RIAR01000029.1.region001).

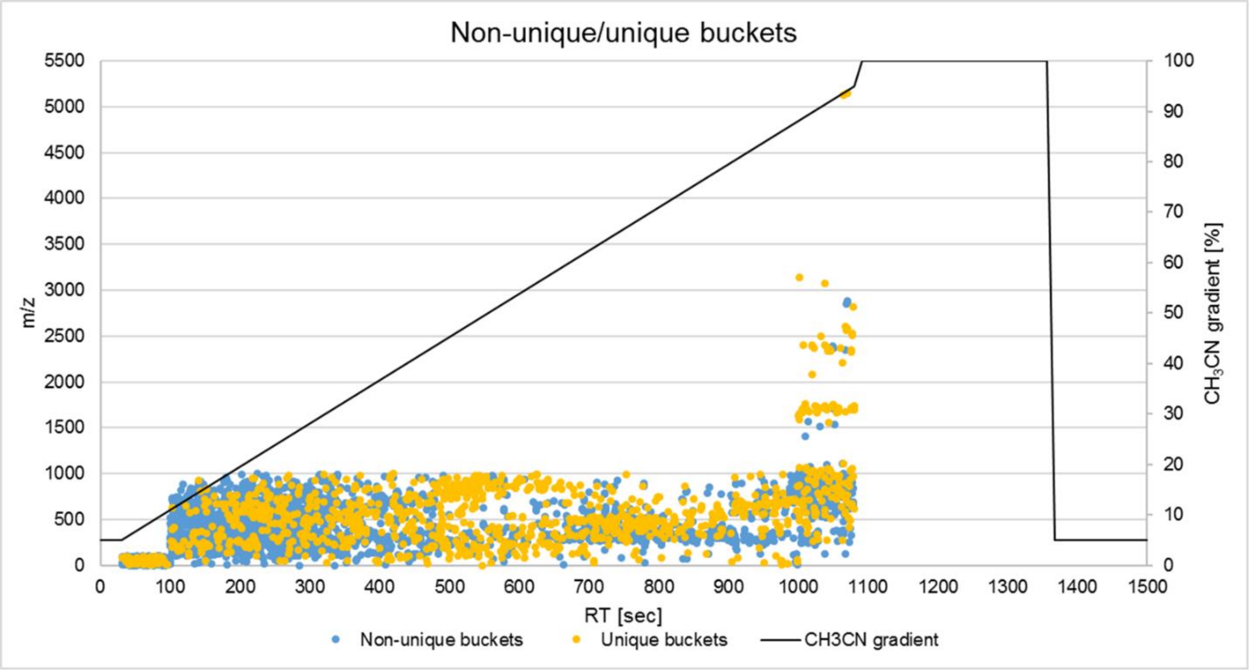


**Fig. S2 Distribution of non-unique/unique buckets revealed by metablomics analysis shown as 2D-scatterplot.** RT, retention time in seconds; *m/z*, mass to charge ratio; $CH_3CN$, acetonitrile.

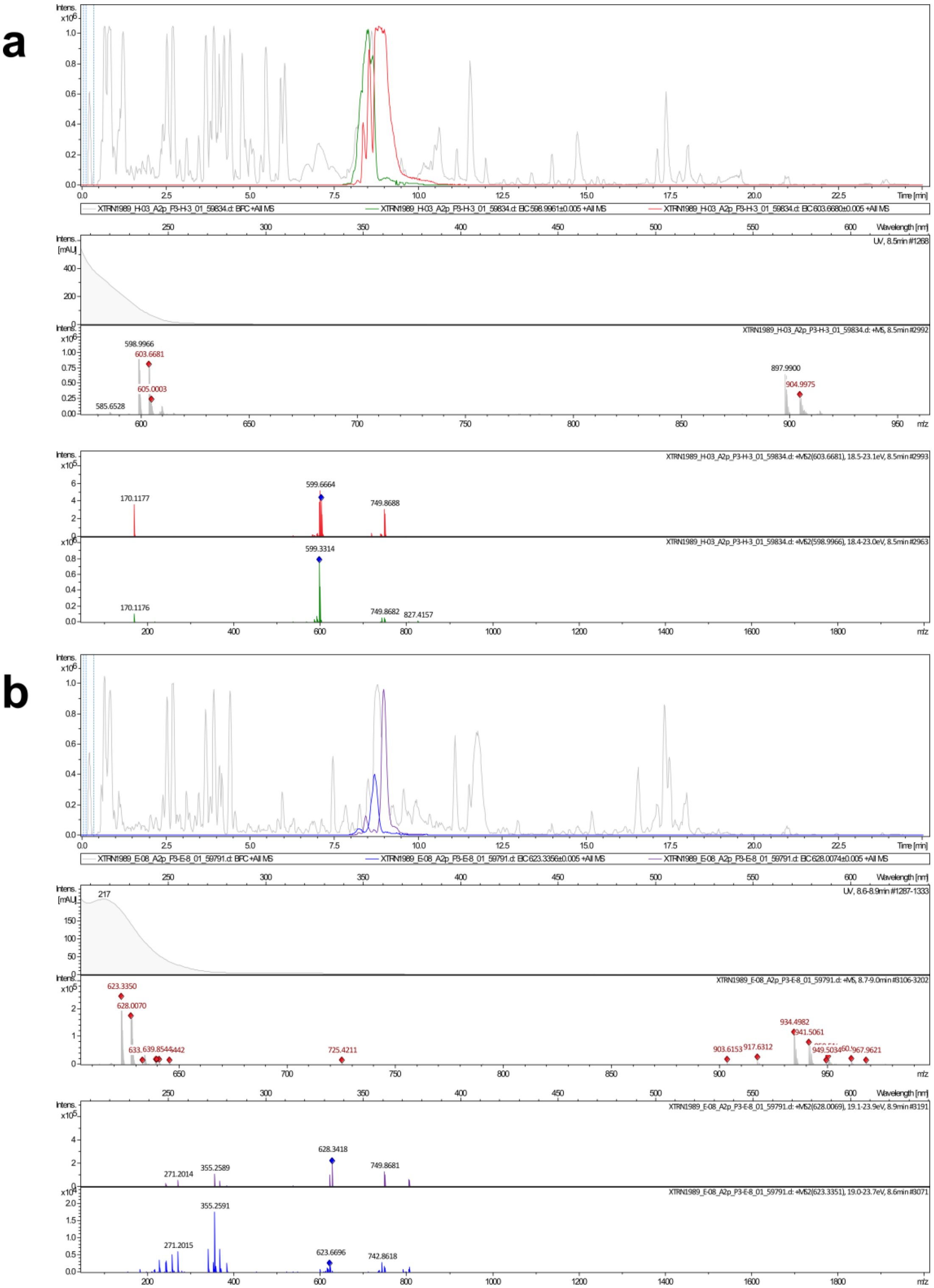


**Fig. S3** (**A**) Analytics (extracted ion chromatograms and MS/MS fragments) of chitinopeptin A with *m/z* 603.6680 $[M+3H]^{3+}$ (red) and chitinopeptin B with m/z 598.6661 $[M+3H]^{3+}$ (green), (**B**) and chitinopeptin C1+C2 with *m/z* 628.0074 $[M+3H]^{3+}$ (purple) and chitinopeptin D1+D2 with *m/z* 623.3356 $[M+3H]^{3+}$ (blue).

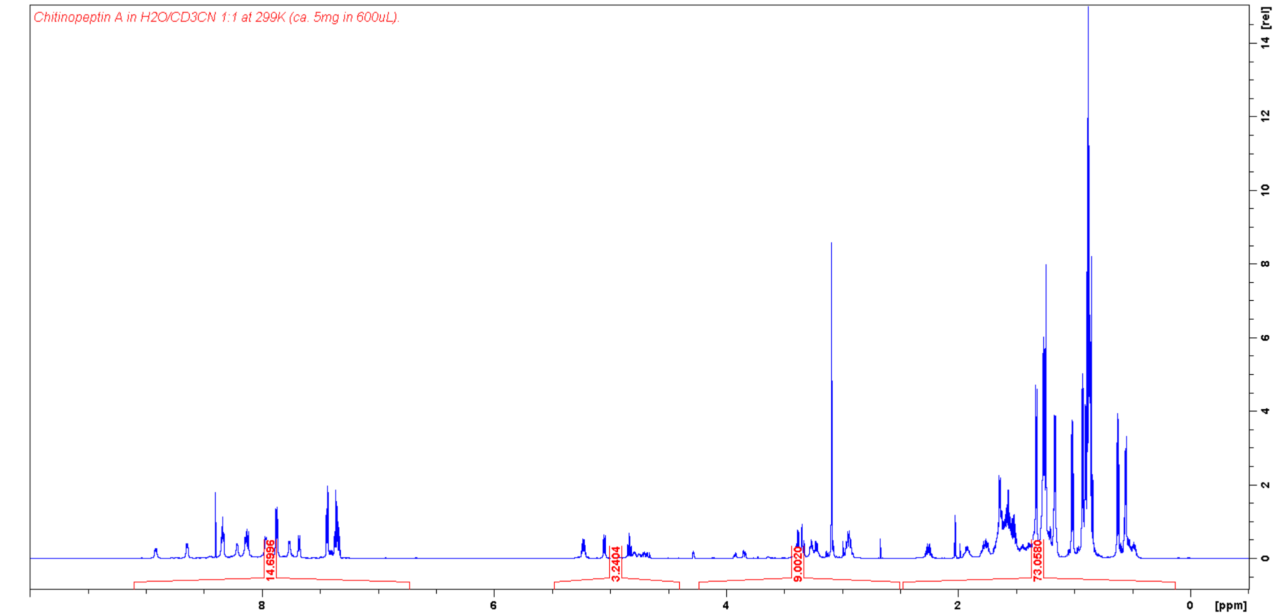


**Fig. S4** $^{1}H$ NMR spectrum (700 MHz, $H_2O/CD_3CN$ 1:1) of chitinopeptin A.

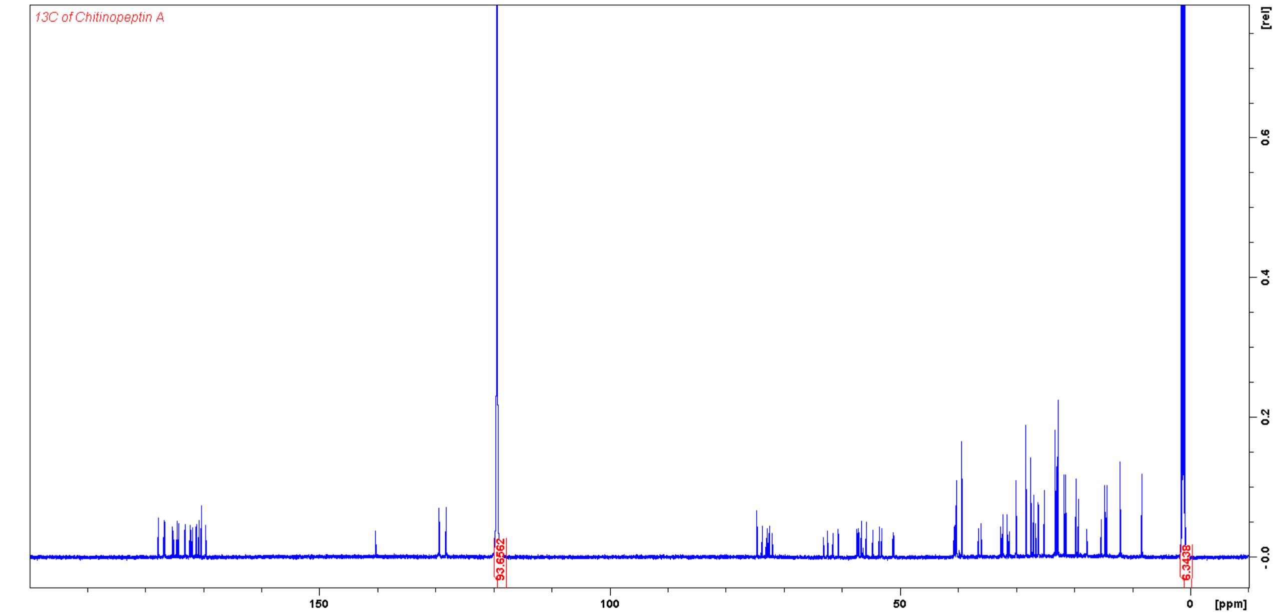


**Fig. S5** $^{13}C$ NMR spectrum (175 MHz, $H_2O/CD_3CN$ 1:1) of chitinopeptin A.

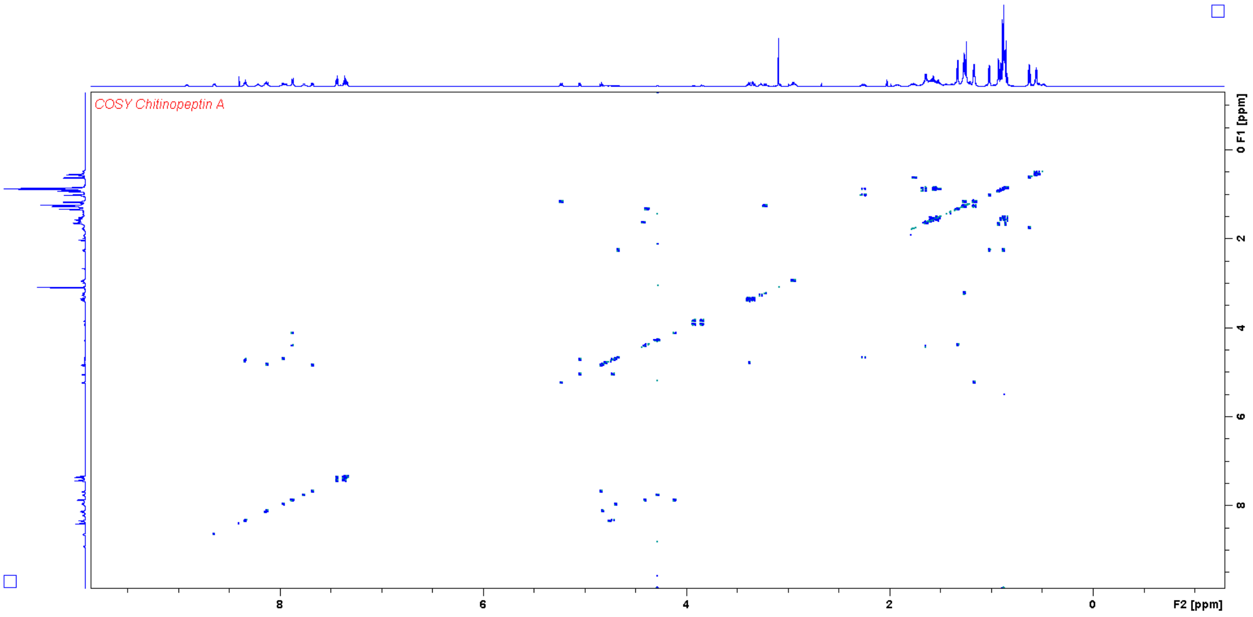


**Fig. S6** COSY NMR spectrum (700 MHz, $H_2O/CD_3CN$ 1:1) of chitinopeptin A.

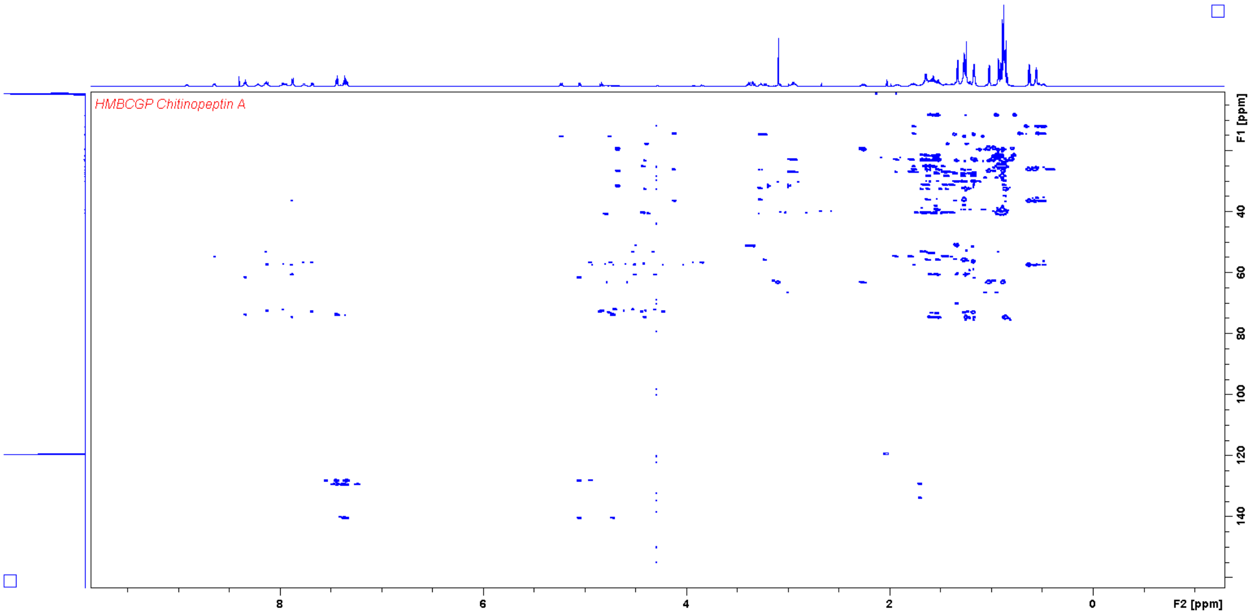


**Fig. S7** HMBC NMR spectrum (700 MHz, $H_2O/CD_3CN$ 1:1) of chitinopeptin A.

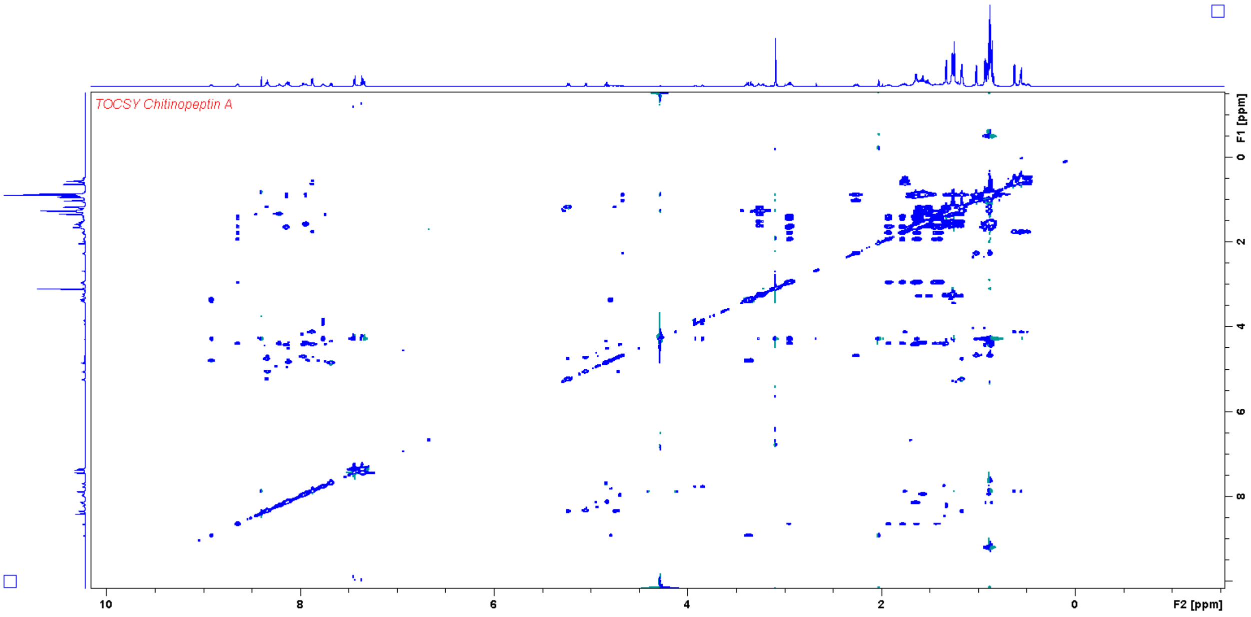


**Fig. S8** TOCSY NMR spectrum (700 MHz, $H_2O/CD_3CN$ 1:1) of chitinopeptin A.

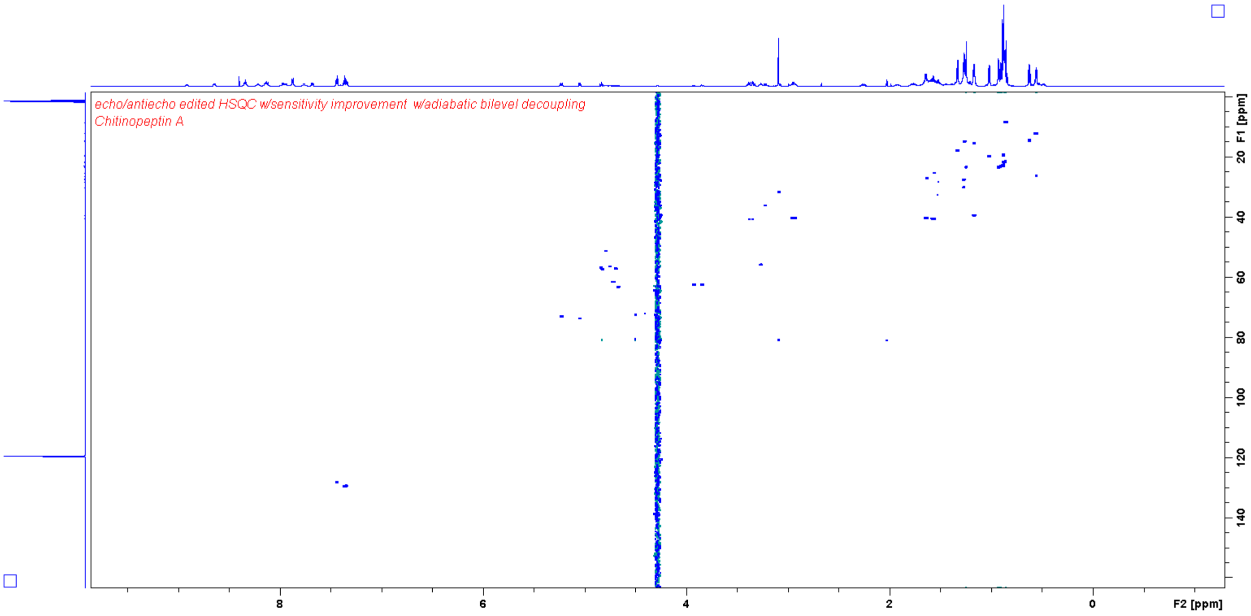


**Fig. S9** HSQC NMR spectrum (700 MHz, $H_2O/CD_3CN$ 1:1) of chitinopeptin A.

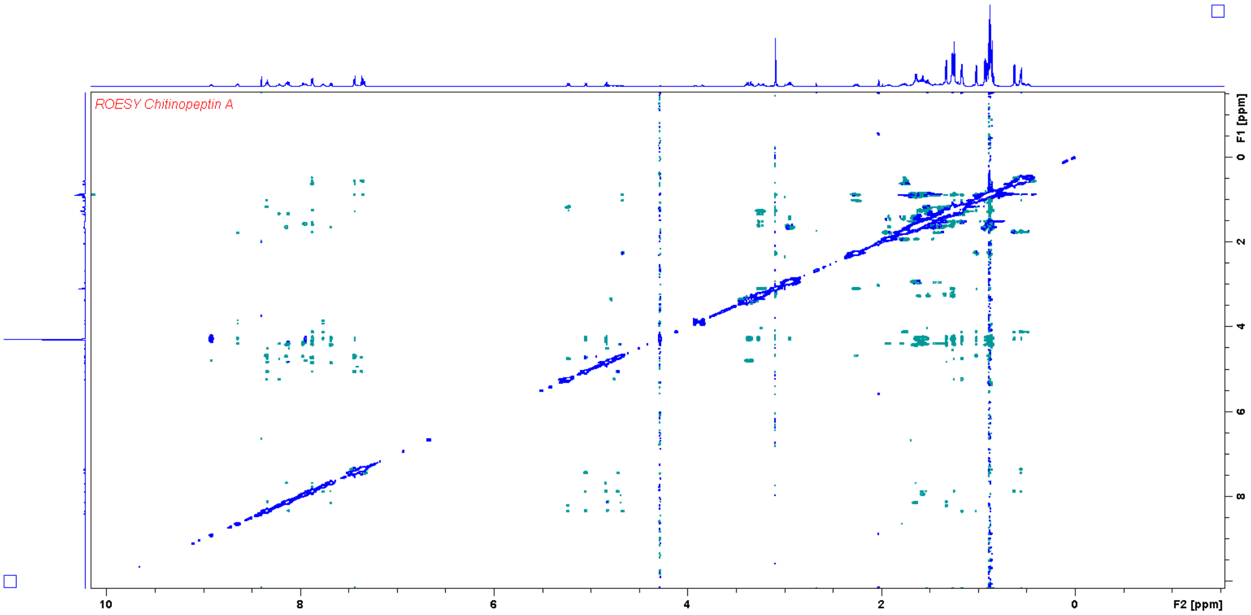


**Fig. S10** ROESY NMR spectrum (700 MHz, $H_2O/CD_3CN$ 1:1) of chitinopeptin A.

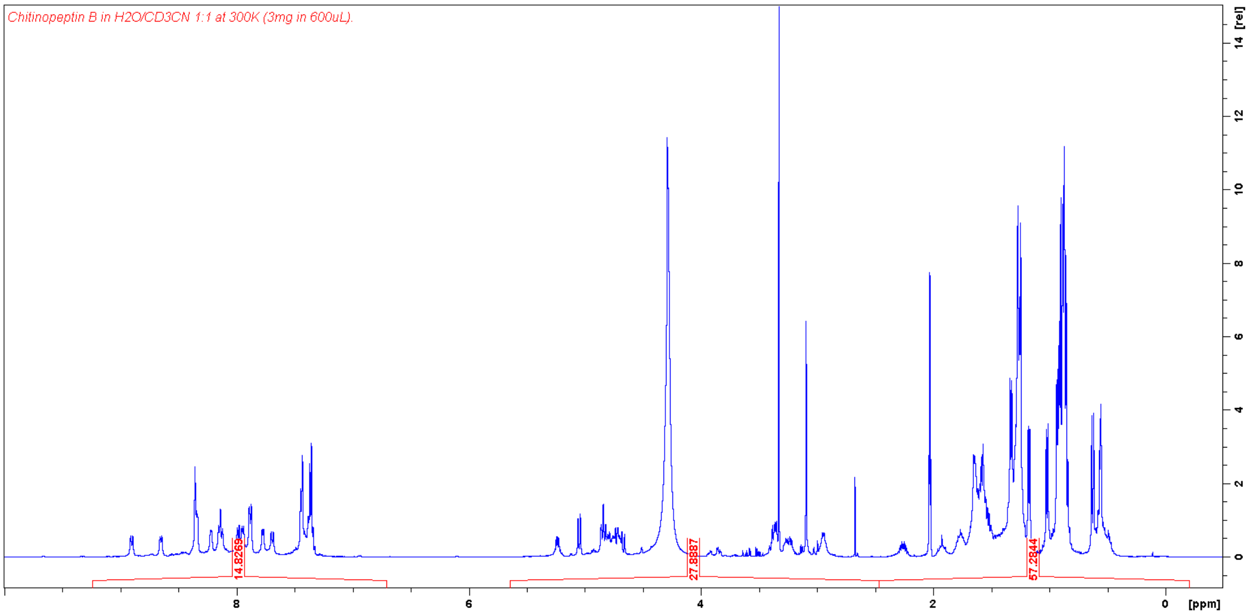


**Fig. S11** $^1$H NMR spectrum (500 MHz, $H_2O/CD_3CN$ 1:1) of chitinopeptin B.

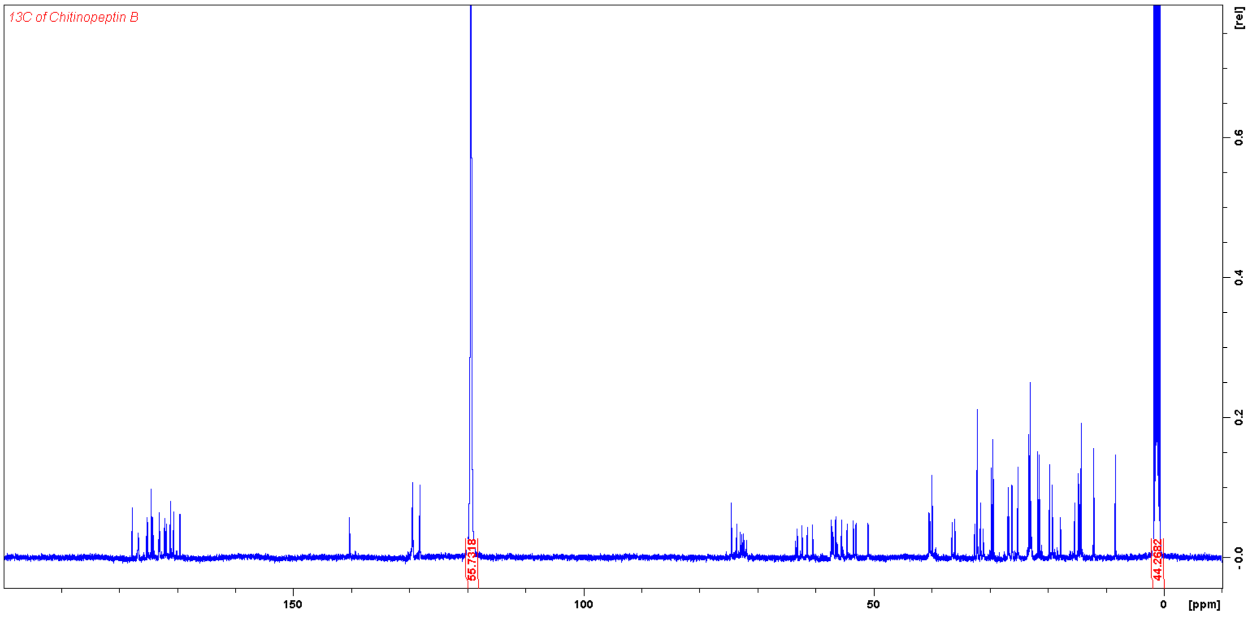


**Fig. S12** $^{13}$C NMR spectrum (125 MHz, $D_2O/CD_3CN$ 1:1) of chitinopeptin B.

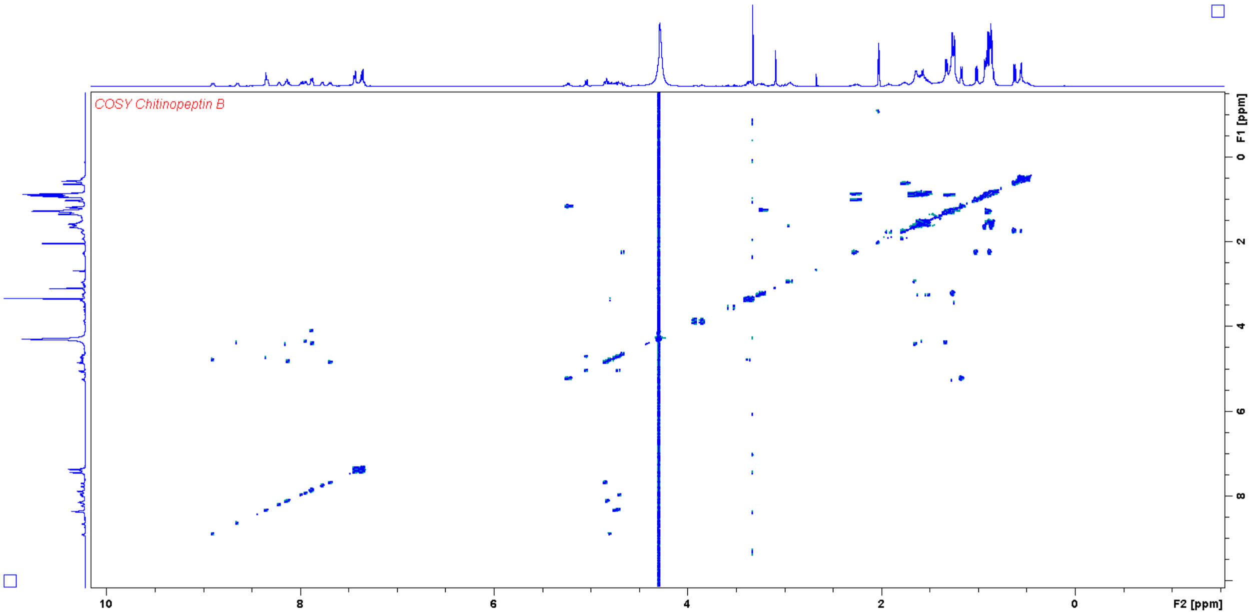


**Fig. S13** COSY NMR spectrum (500 MHz, $H_2O/CD_3CN$ 1:1) of chitinopeptin B.

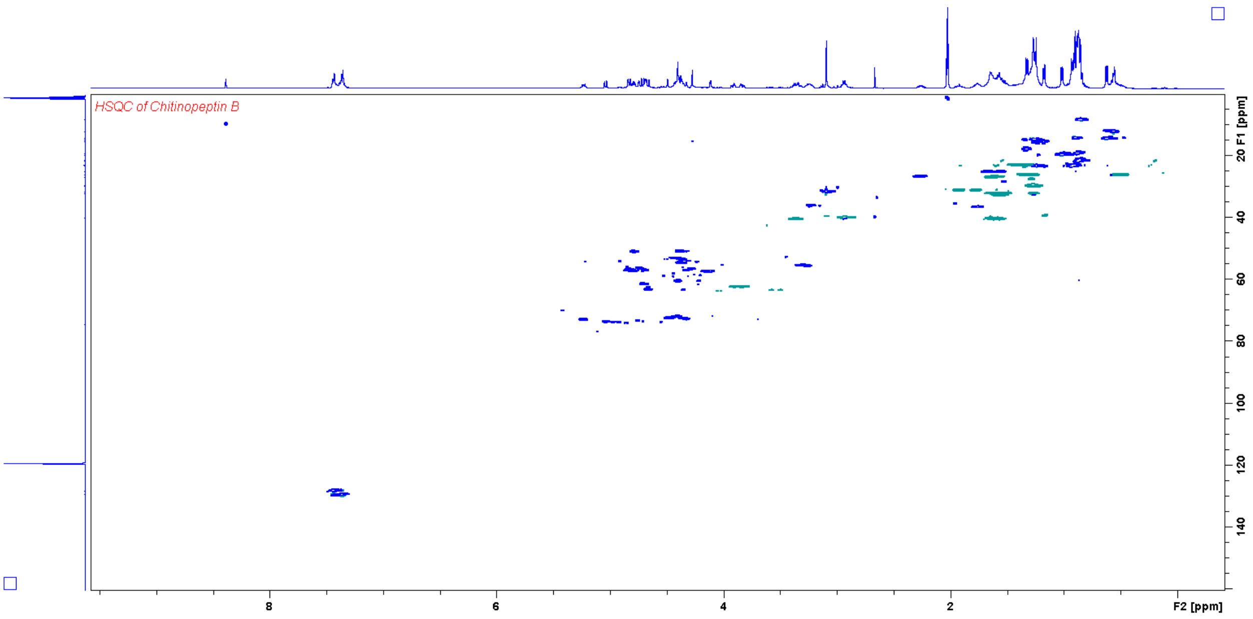


**Fig. S14** HSQC NMR spectrum (500 MHz, $D_2O/CD_3CN$ 1:1) of chitinopeptin B.

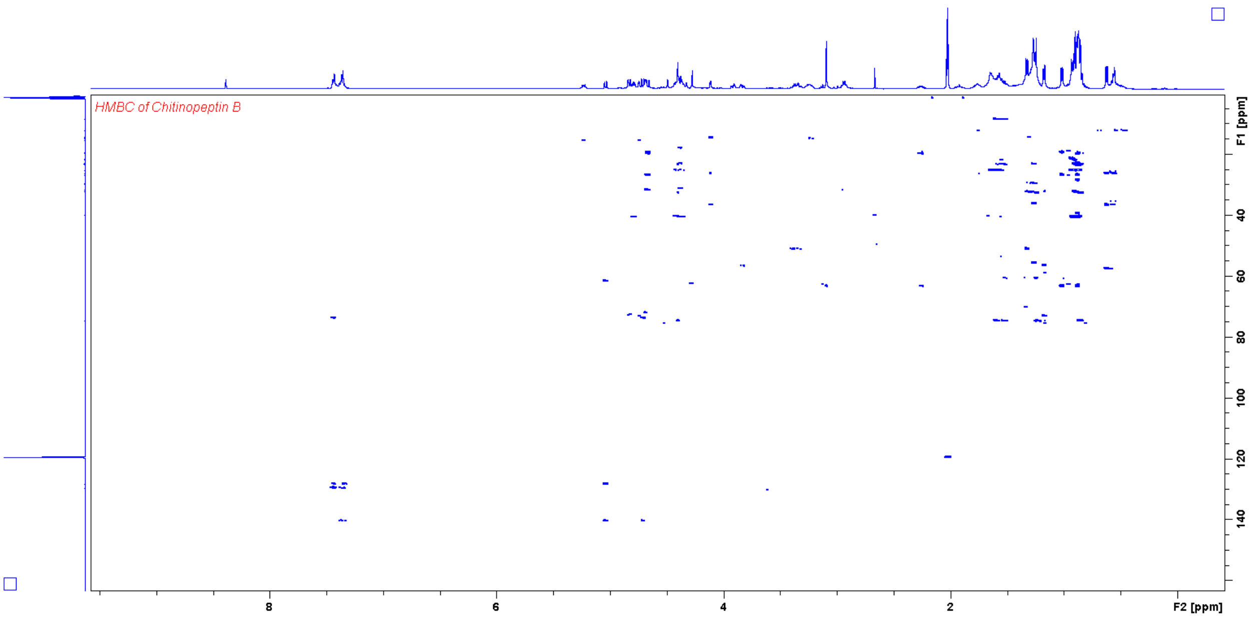


**Fig. S15** HMBC NMR spectrum (500 MHz, $D_2O/CD_3CN$ 1:1) of chitinopeptin B.

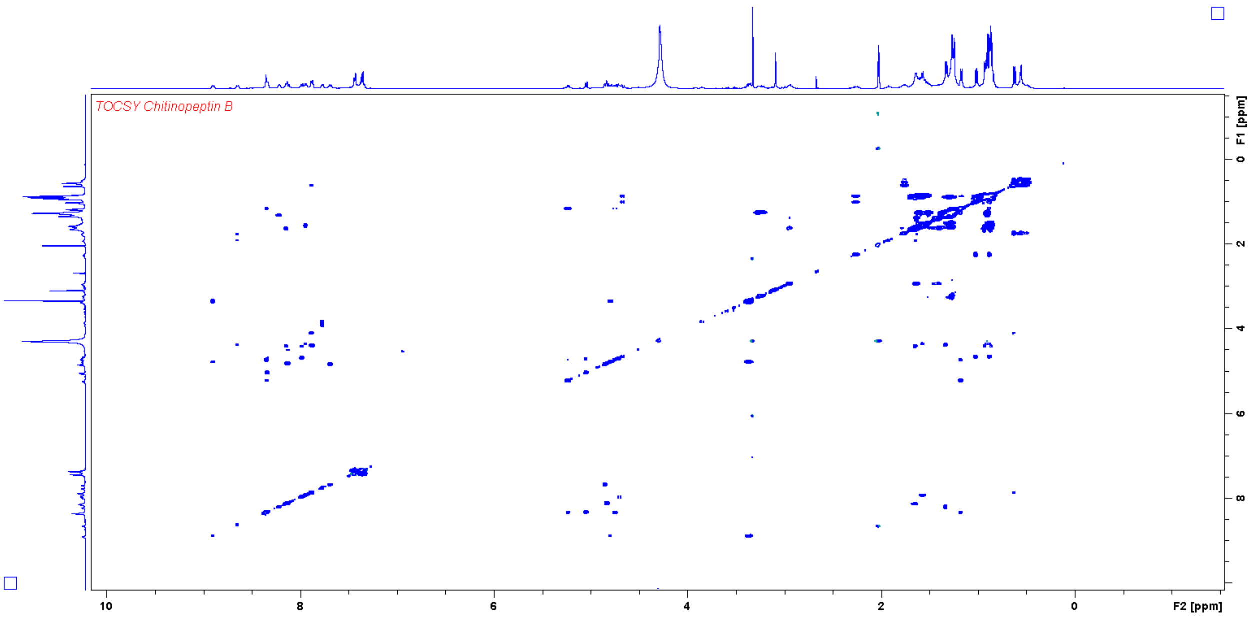


**Fig. S16** TOCSY NMR spectrum (500 MHz, $H_2O/CD_3CN$ 1:1) of chitinopeptin B.

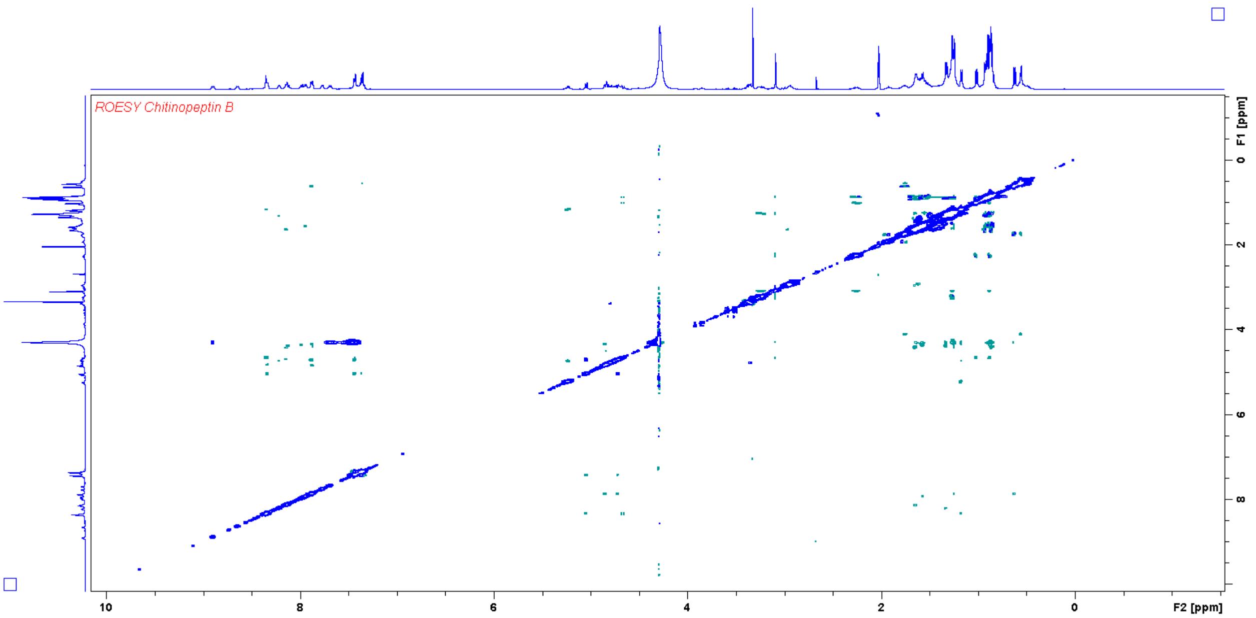


**Fig. S17** ROESY NMR spectrum (500 MHz, $H_2O/CD_3CN$ 1:1) of chitinopeptin B.

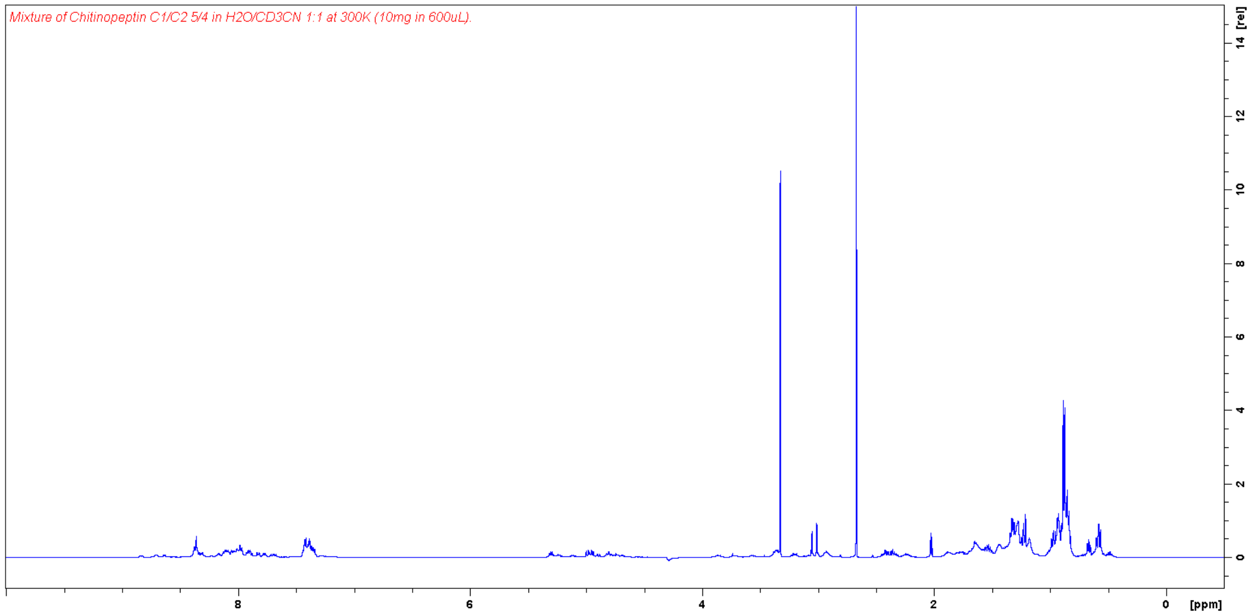


**Fig. S18** $^{1}H$ NMR spectrum (500 MHz, $H_2O/CD_3CN$ 1:1) of chitinopeptin C1 and C2 5/4 mixture.

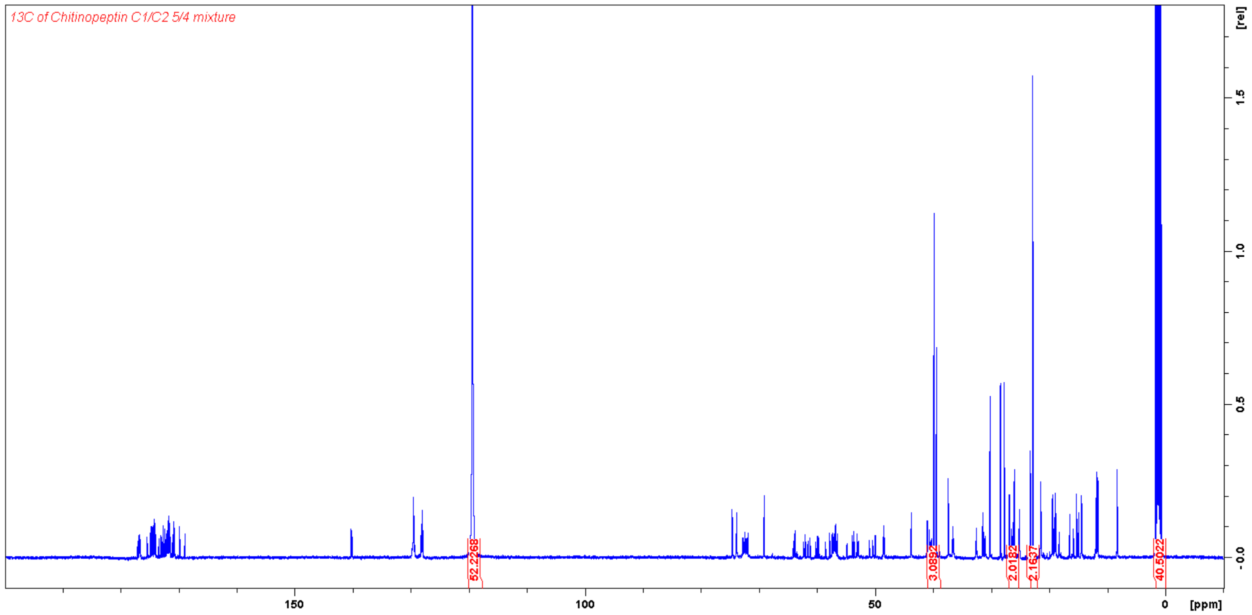


**Fig. S19** $^{13}C$ NMR spectrum (125 MHz, $D_2O/CD_3CN$ 1:1) of chitinopeptin C1 and C2 5/4 mixture.

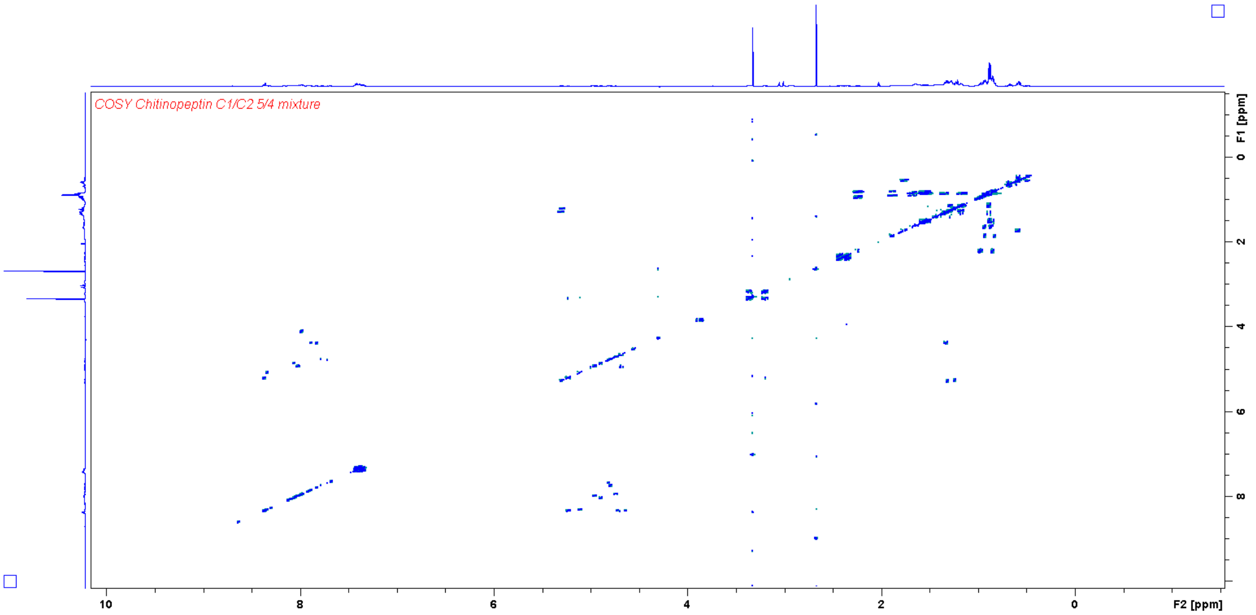


**Fig. S20** COSY NMR spectrum (500 MHz, $H_2O/CD_3CN$ 1:1) of chitinopeptin C1 and C2 5/4 mixture.

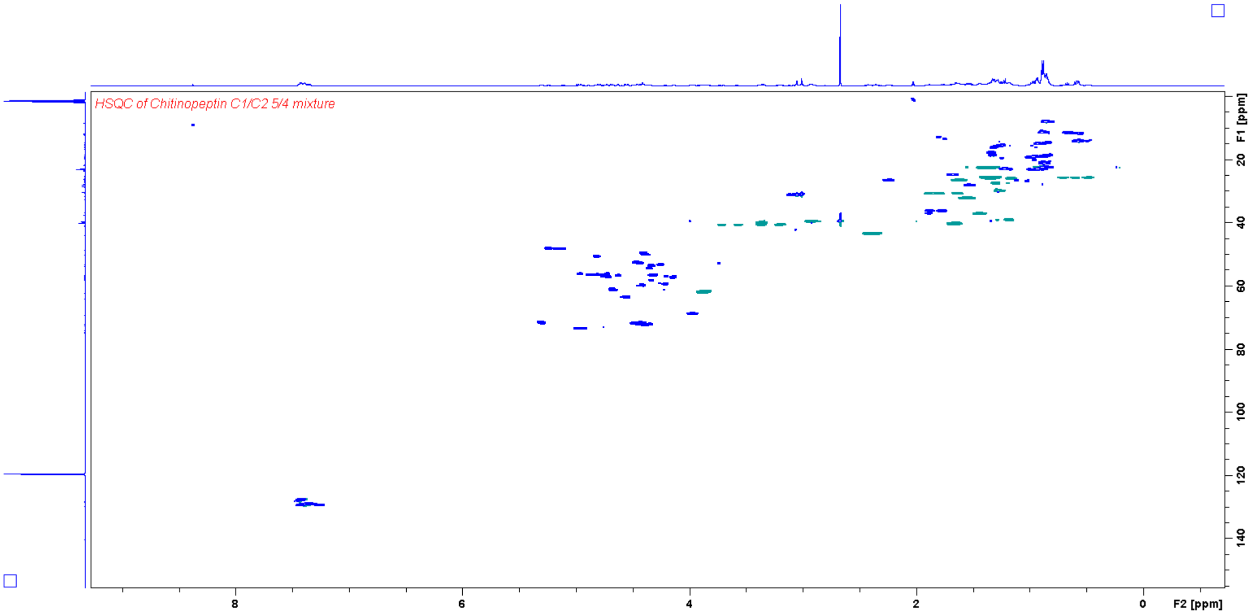


**Fig. S21** HSQC NMR spectrum (500 MHz, $D_2O/CD_3CN$ 1:1) of chitinopeptin C1 and C2 5/4.

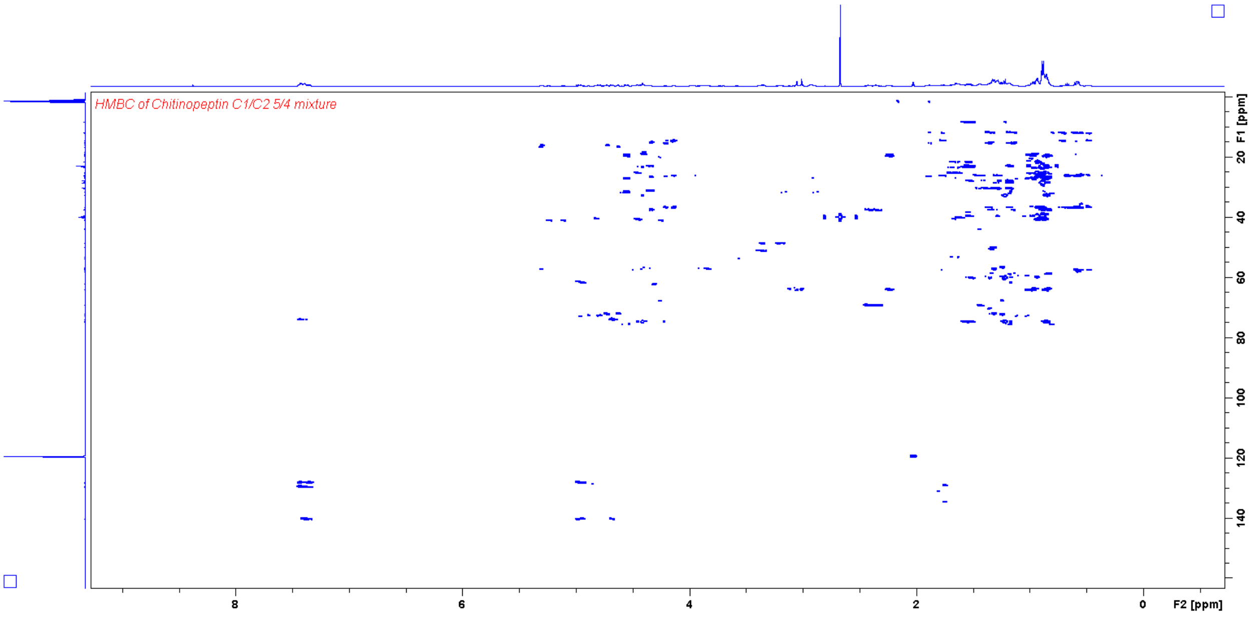


**Fig. S22** HMBC NMR spectrum (500 MHz, $D_2O/CD_3CN$ 1:1) of chitinopeptin C1 and C2 5/4.

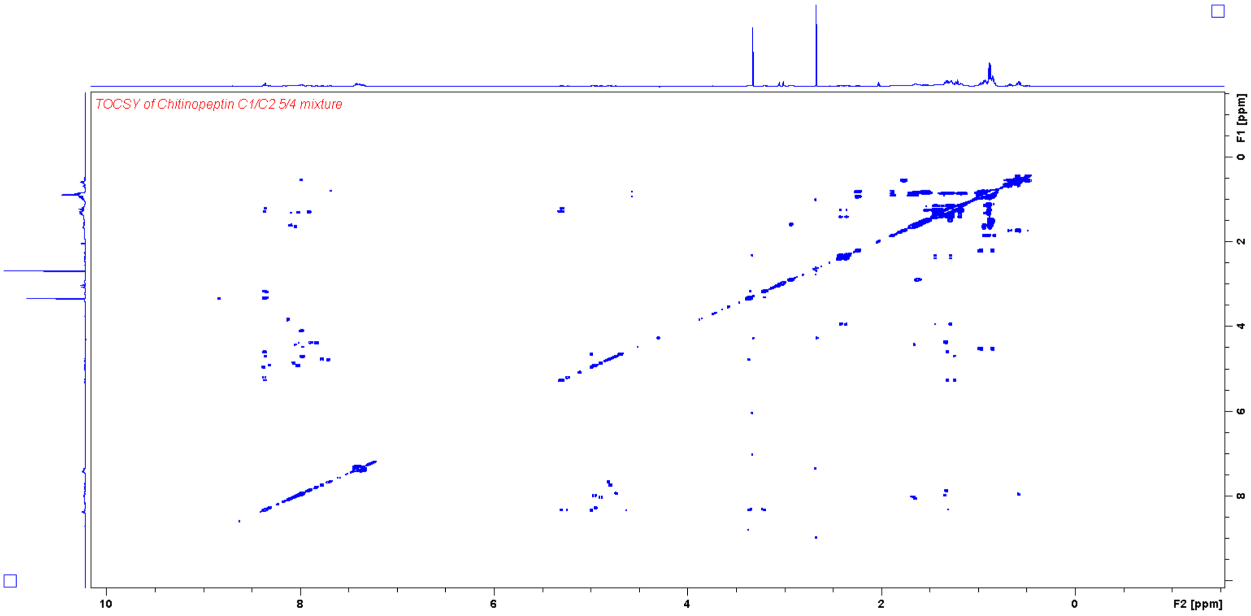


**Fig. S23** TOCSY NMR spectrum (500 MHz, $H_2O/CD_3CN$ 1:1) of chitinopeptin C1 and C2 5/4 mixture.

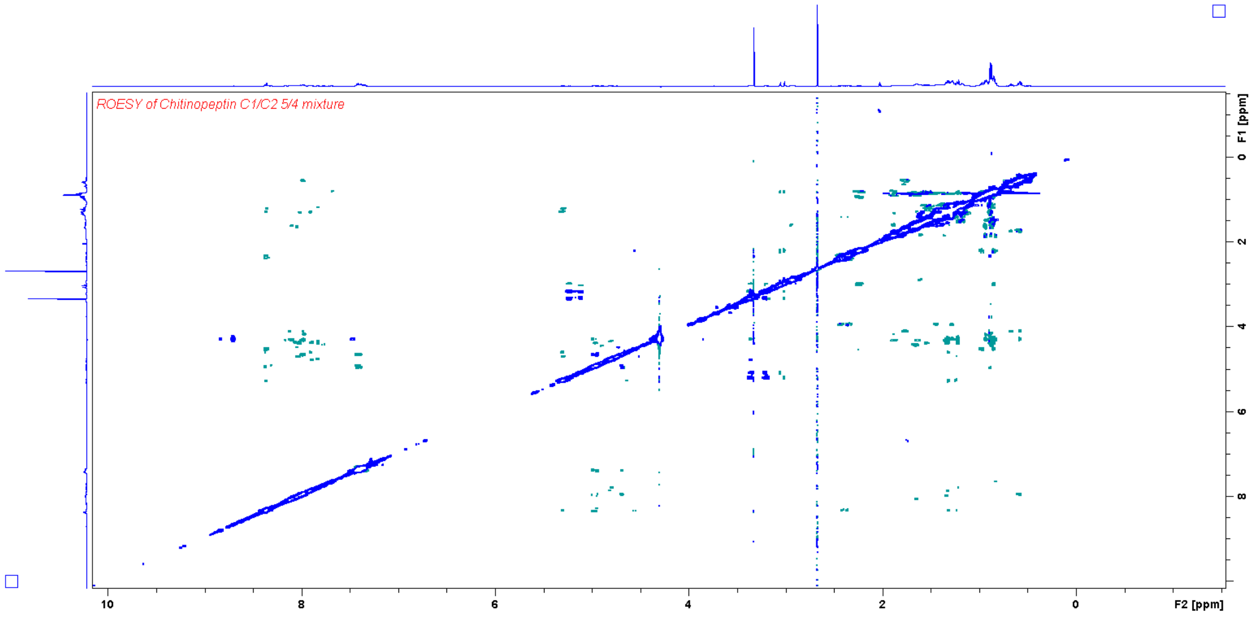


**Fig. S24** ROESY NMR spectrum (500 MHz, $H_2O/CD_3CN$ 1:1) of chitinopeptin C1 and C2 5/4 mixture.

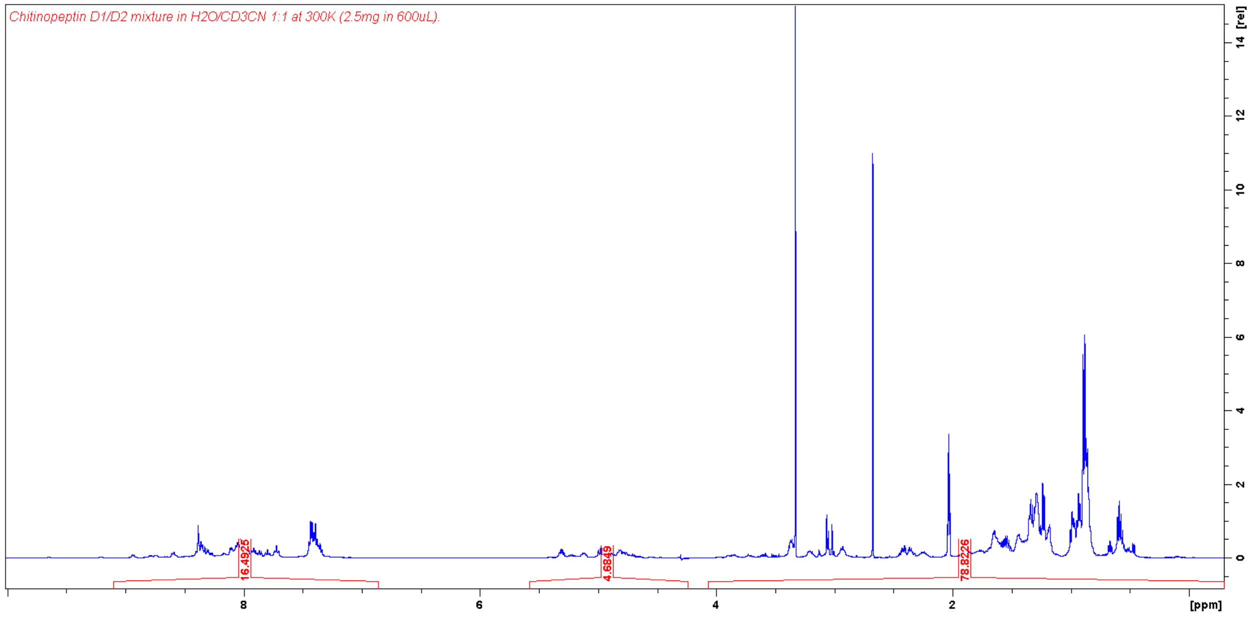


**Fig. S25** $^1H$ NMR spectrum (500 MHz, $H_2O/CD_3CN$ 1:1) of chitinopeptin D1 and D2 mixture.

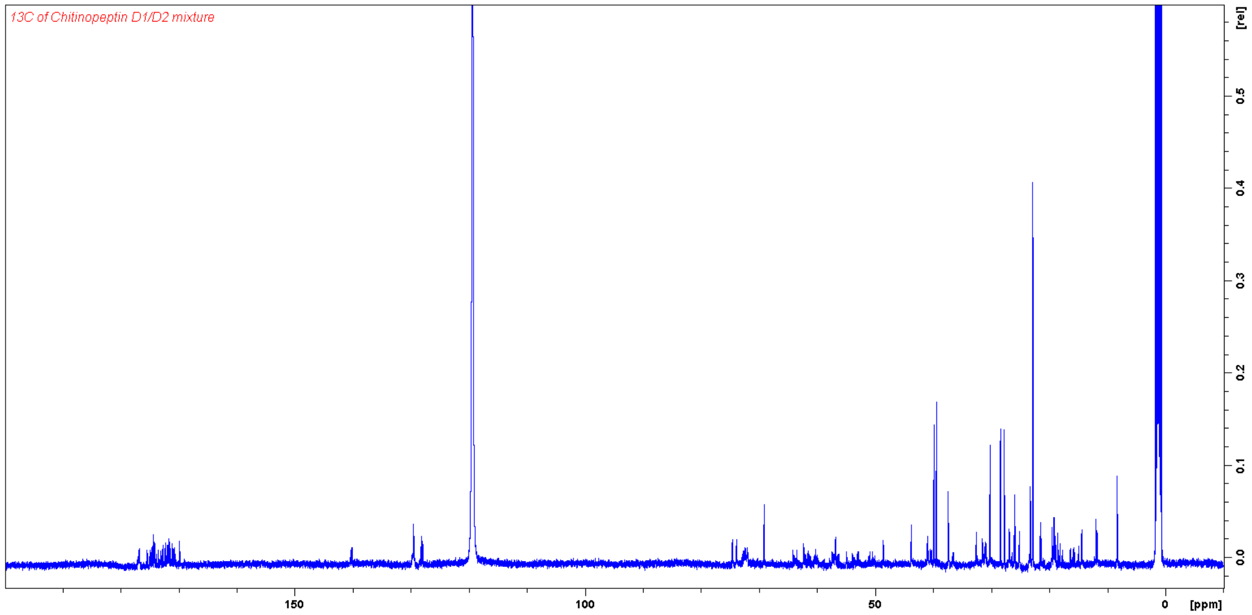


**Fig. S26** $^{13}$C NMR spectrum (125 MHz, $D_2O/CD_3CN$ 1:1) of chitinopeptin D1 and D2.

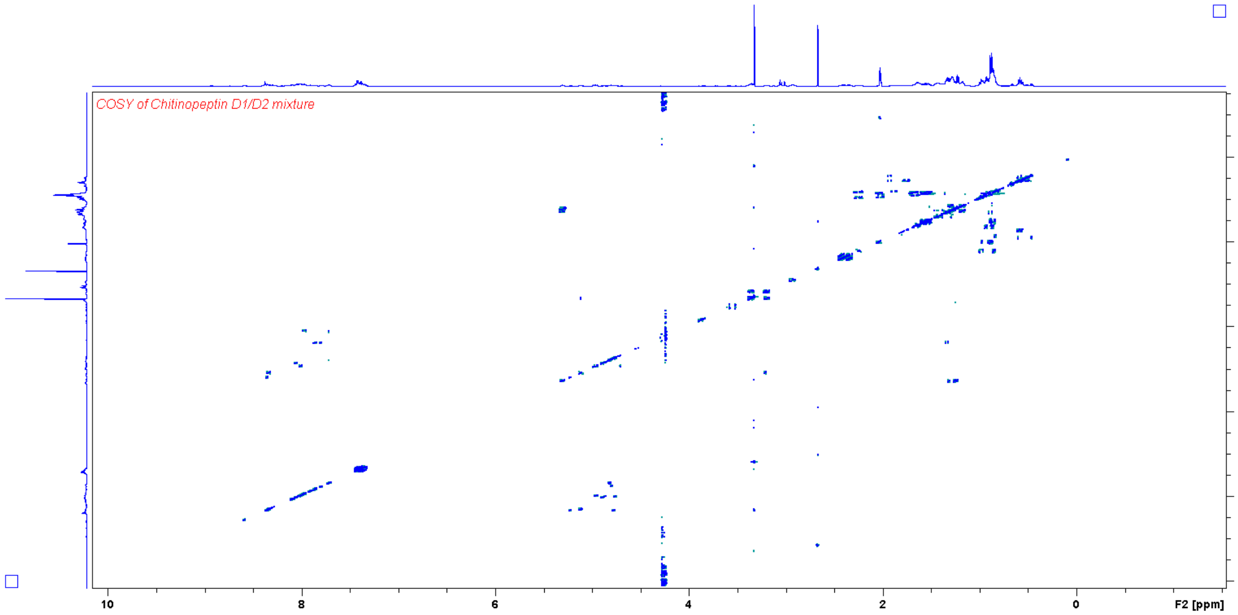


**Fig. S27** COSY NMR spectrum (500 MHz, $H_2O/CD_3CN$ 1:1) of chitinopeptin D1 and D2.

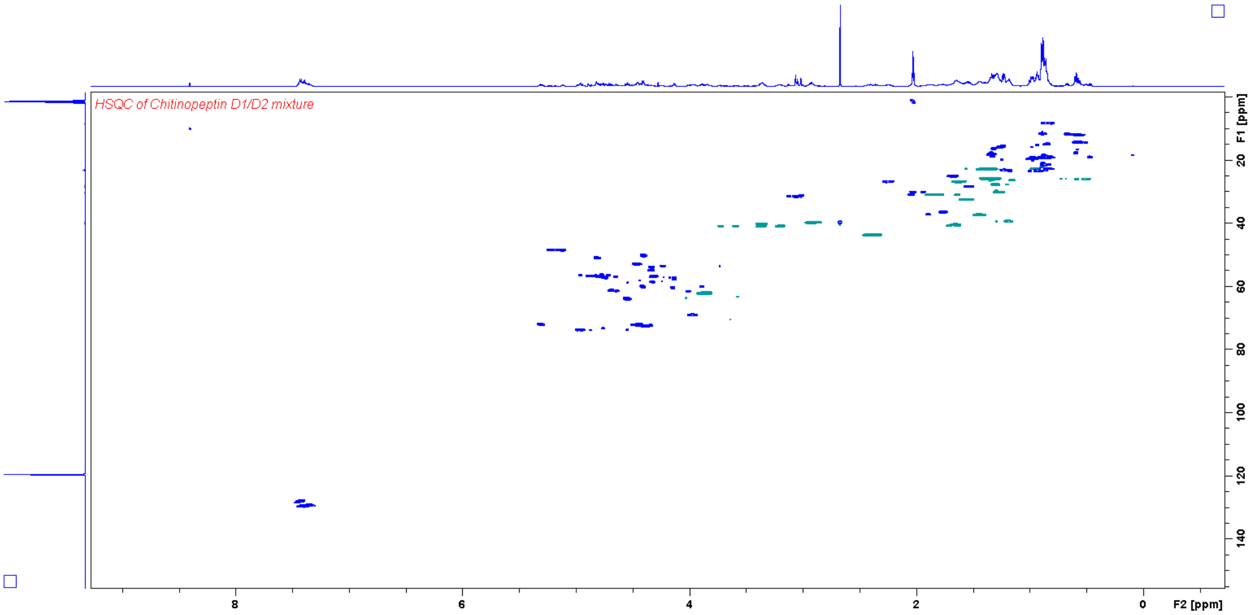


**Fig. S28** HSQC NMR spectrum (500 MHz, $D_2O/CD_3CN$ 1:1) of chitinopeptin D1 and D2.

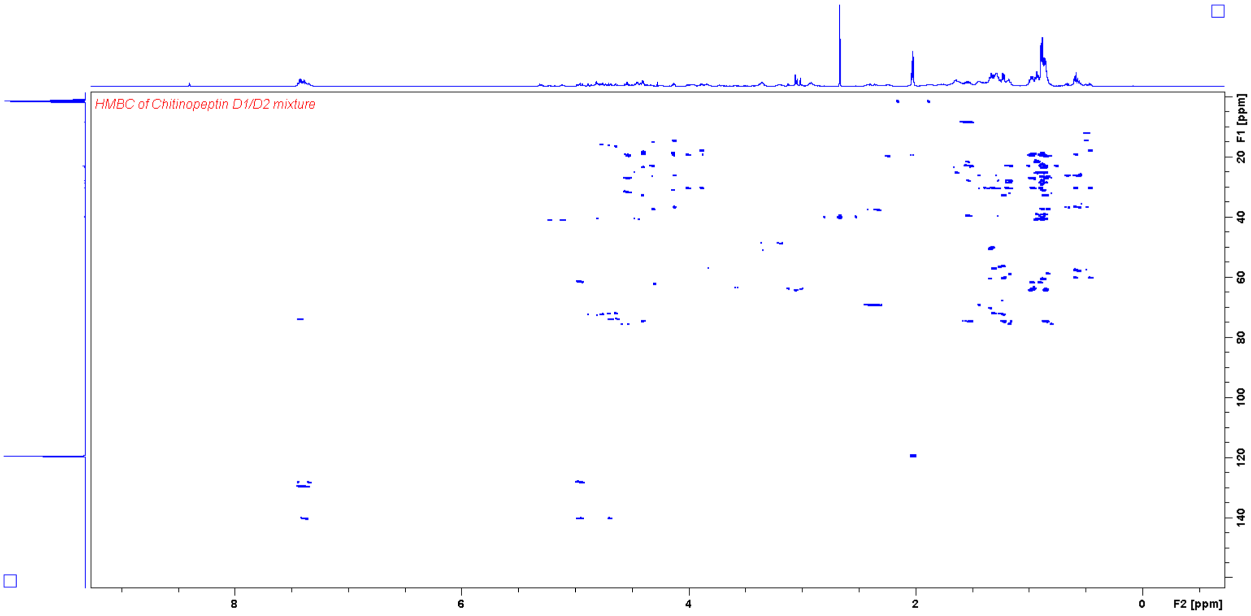


**Fig. S29** HMBC NMR spectrum (500 MHz, $D_2O/CD_3CN$ 1:1) of chitinopeptin D1 and D2.

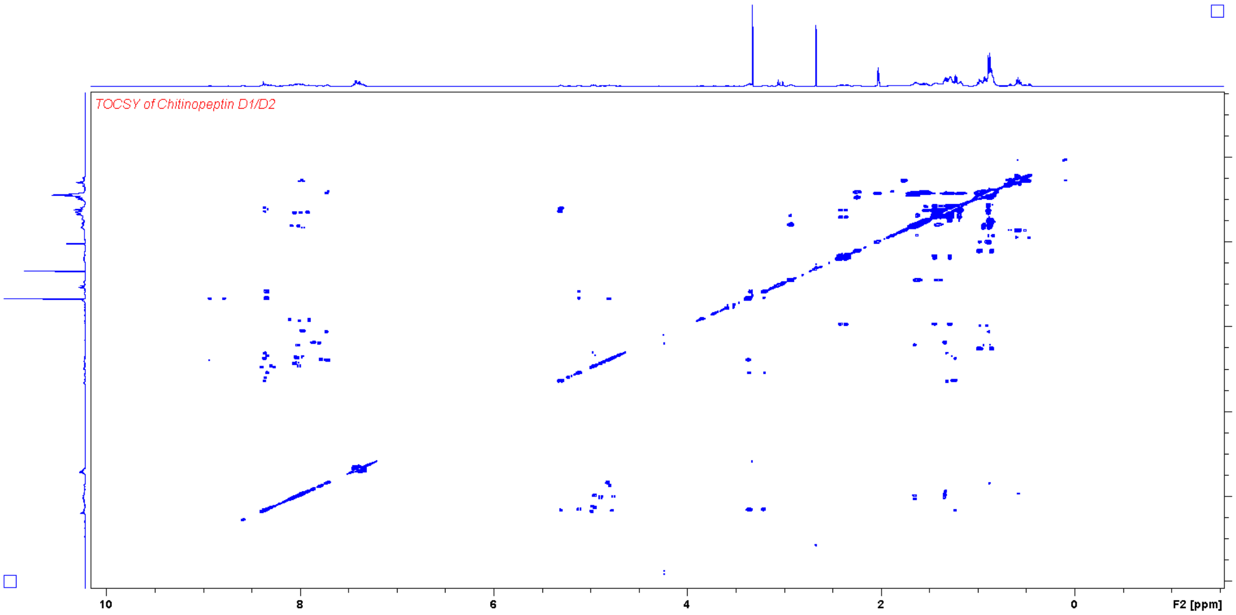


**Fig. S30** COSY NMR spectrum (500 MHz, $H_2O/CD_3CN$ 1:1) of chitinopeptin D1 and D2.

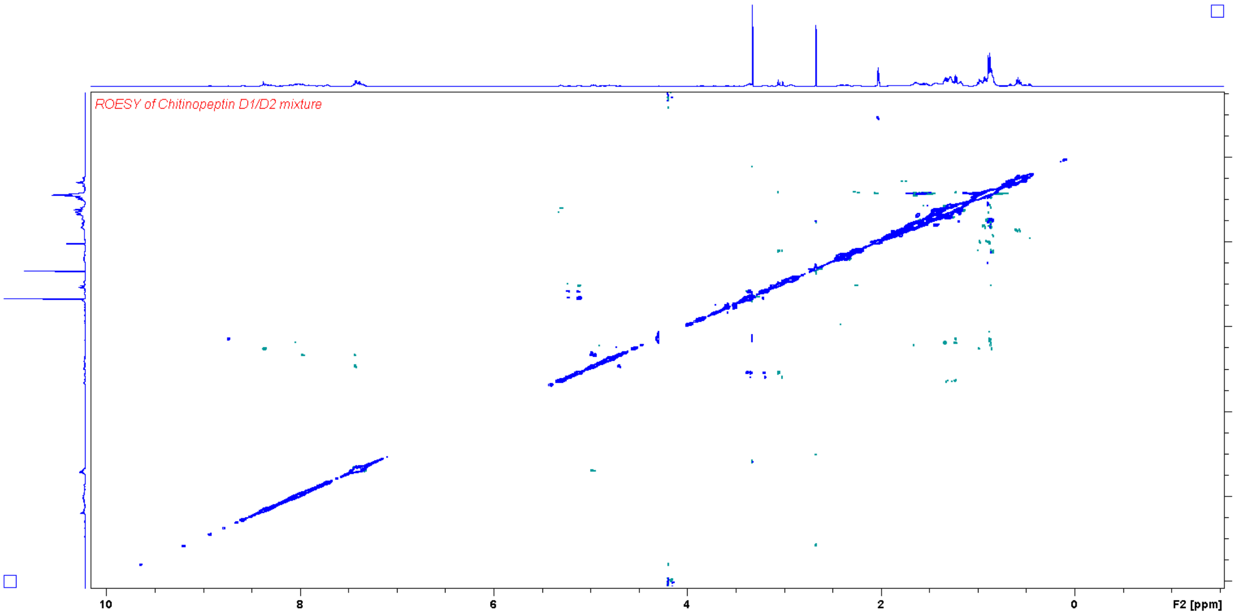


**Fig. S31** ROESY NMR spectrum (500 MHz, $H_2O/CD_3CN$ 1:1) of chitinopeptin D1 and D2.

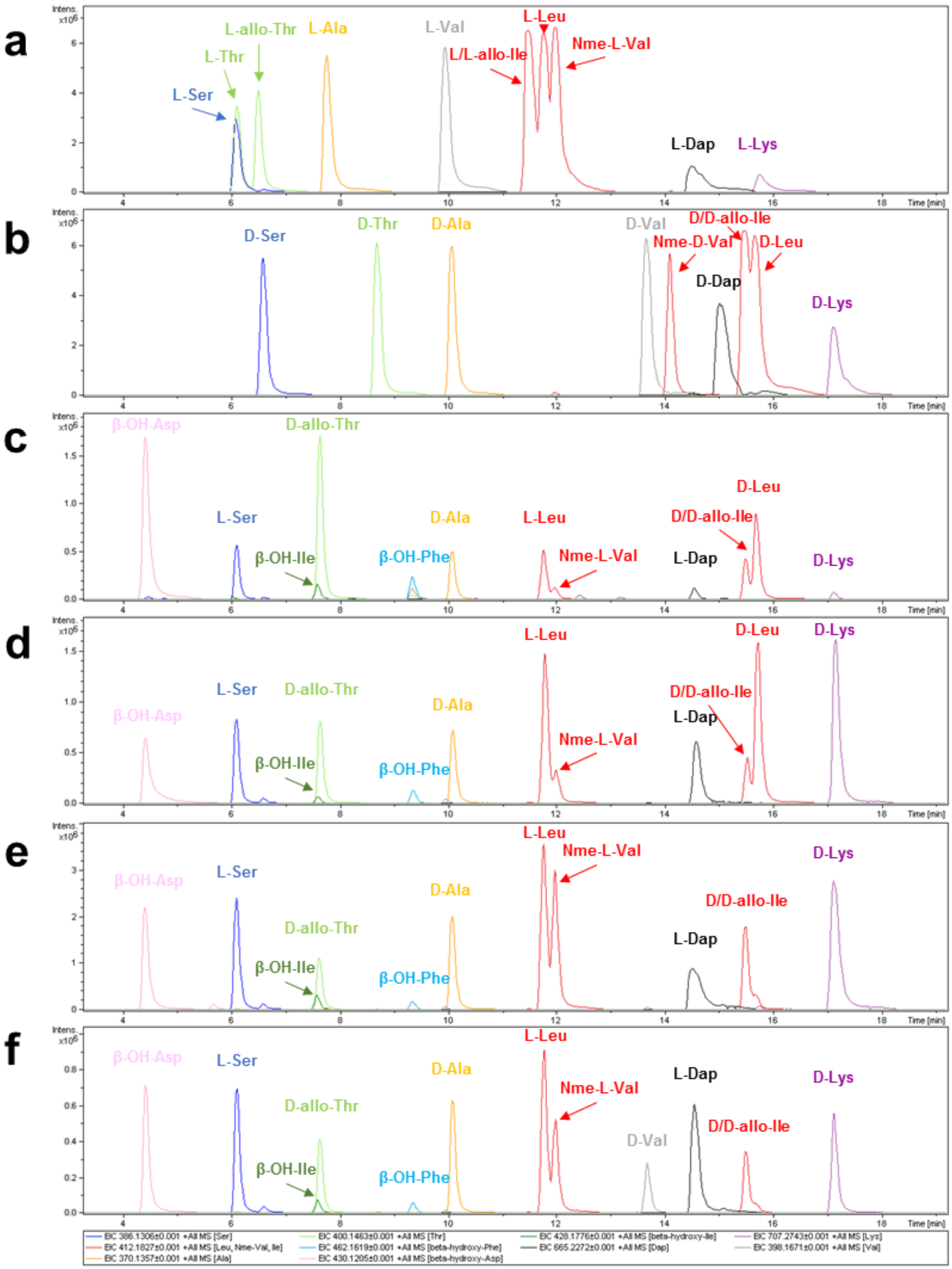


**Fig. S32 Comparison of the Marfey derivatization products of the chitinopeptin A-D DCl hydrolysates and commercially available amino acid standards derivatized with L-FDVA.** (**A**) Commercially available L-amino acid standards. (**B**) Commercially available D-amino acid standards. (**C**) DCl hydrolysate of chitinopeptin A. (**D**) DCl hydrolysate of chitinopeptin B. (**E**) DCl hydrolysate of chitinopeptin C1+C2. (**F**) DCl hydrolysate of chitinopeptin D1+D2.

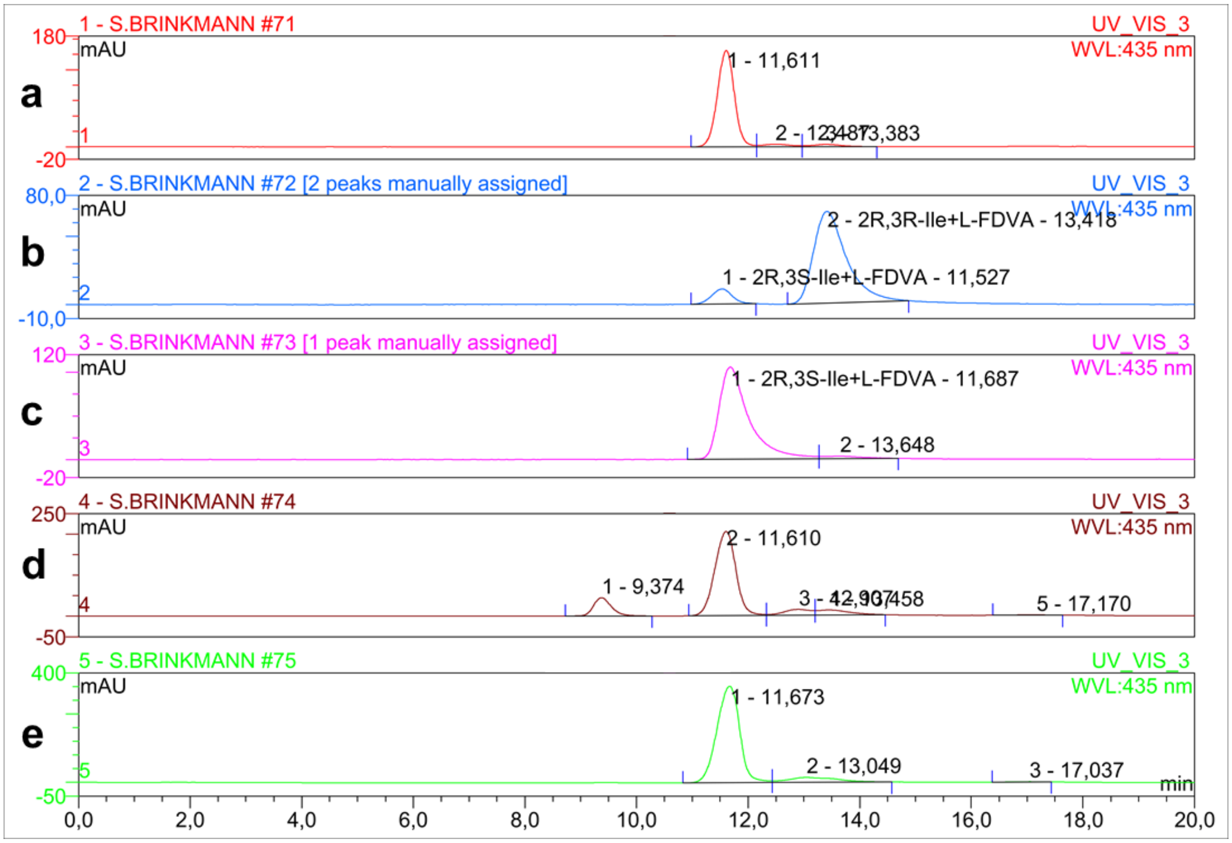


**Fig. 33 Chiral HPLC of 2*R*,3*S*-Ile and 2*R*,3*R*-Ile standards and chitinopeptin samples derivatized with L-FDVA.** Chiral HPLC conditions: Chiralpak IC; hexane:isopropyl alcohol:formic acid 75:25:0.2. (**A**) chitinopeptin D1+D2, (**B**) 2*R*,3*R*-Ile, (**C**) 2*R*,3*S*-Ile, (**D**) mixture of chitinopeptin A and B, (**E**) chitinopeptin C1+C2

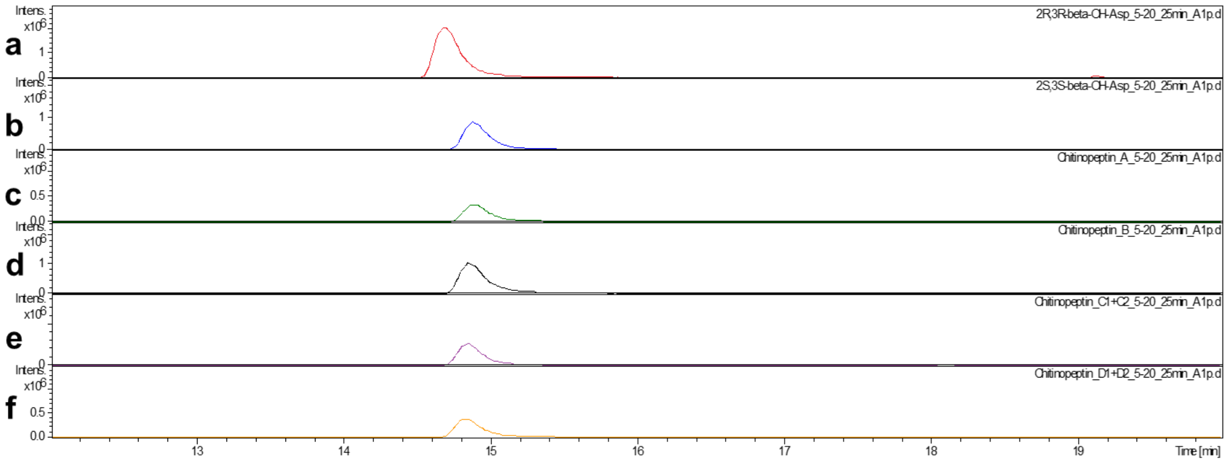


**Fig. S34 C18-RP UHPLC-MS extracted ion chromatograms of β-hydroxyaspartic acids derivatized with L-FDVA at *m/z* 430.1205 $[M+H]^+$ within synthesized β-hydroxyaspartic acid standards and chitinopeptin A-D samples.** (**A**) (2*R*,3*R*)-3-hydroxyaspartic acid, (**B**) (2*S*,3*S*)-3-hydroxyaspartic acid, (**C**) chitinopeptin A, (**D**) chitinopeptin B, (**E**) chitinopeptin C1+C2, (**F**) chitinopeptin D1+D2

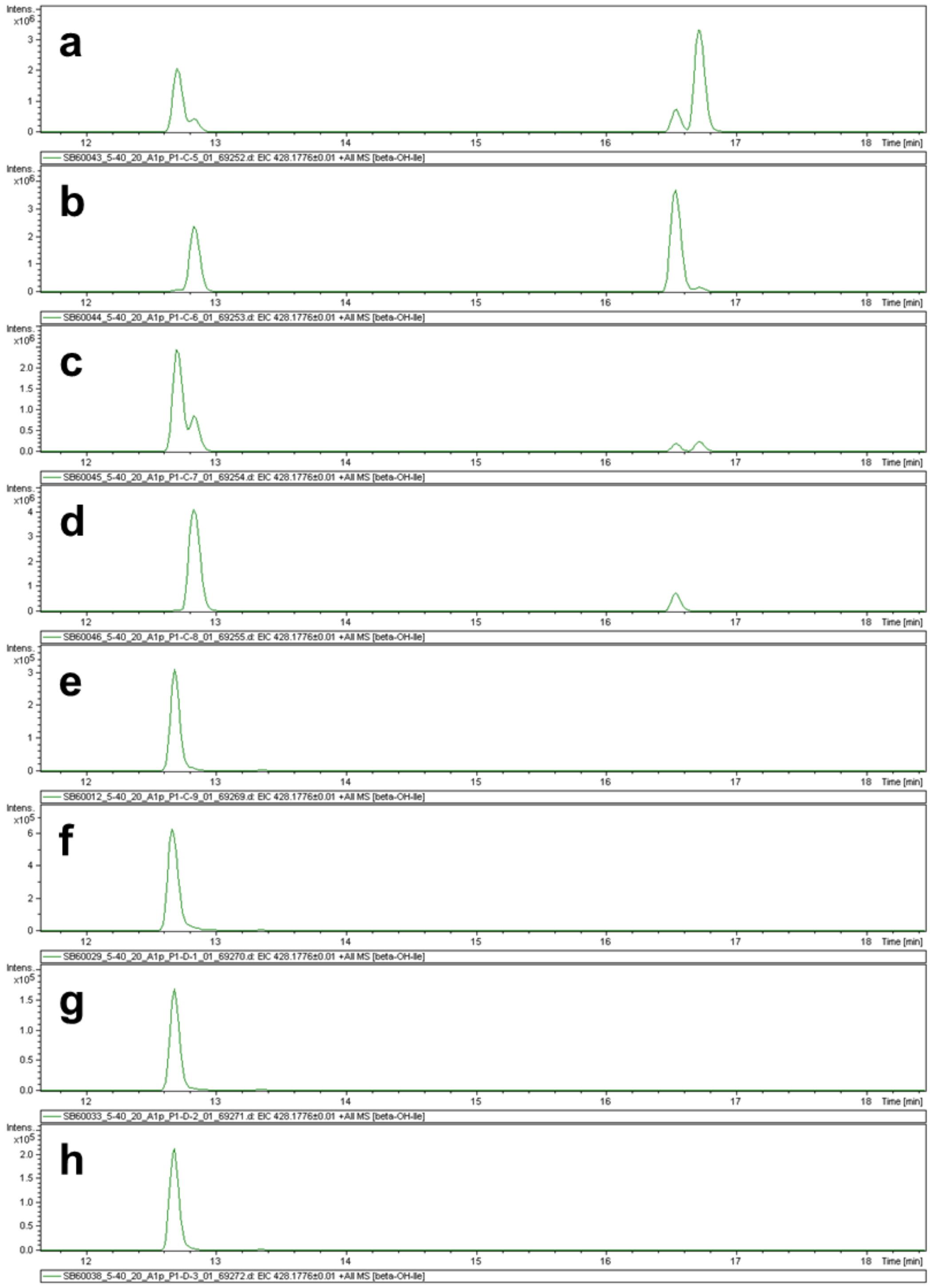


**Fig. S35 C18-RP UHPLC-MS extracted ion chromatograms of β-hydroxyisoleucines derivatized with L-FDVA at *m/z* 428.1776 [M+H]$^+$ within synthesized β- hydroxyisoleucine standards and chitinopeptin A-D samples.** (**A**) (2*S*,3*R*)- and (2*R*,3*S*)-3-hydroxyisoleucine, (**B**) (2*S*,3*S*)- and (2*R*,3*R*)-3-hydroxyisoleucine, (**C**) (2*S*,3*R*)-3-hydroxyisoleucine, (**D**) (2*S*,3*S*)-3-hydroxyisoleucine, (**E**) chitinopeptin A, (**F**) chitinopeptin B, (**G**) chitinopeptin C1+C2, (**H**) chitinopeptin D1+D2

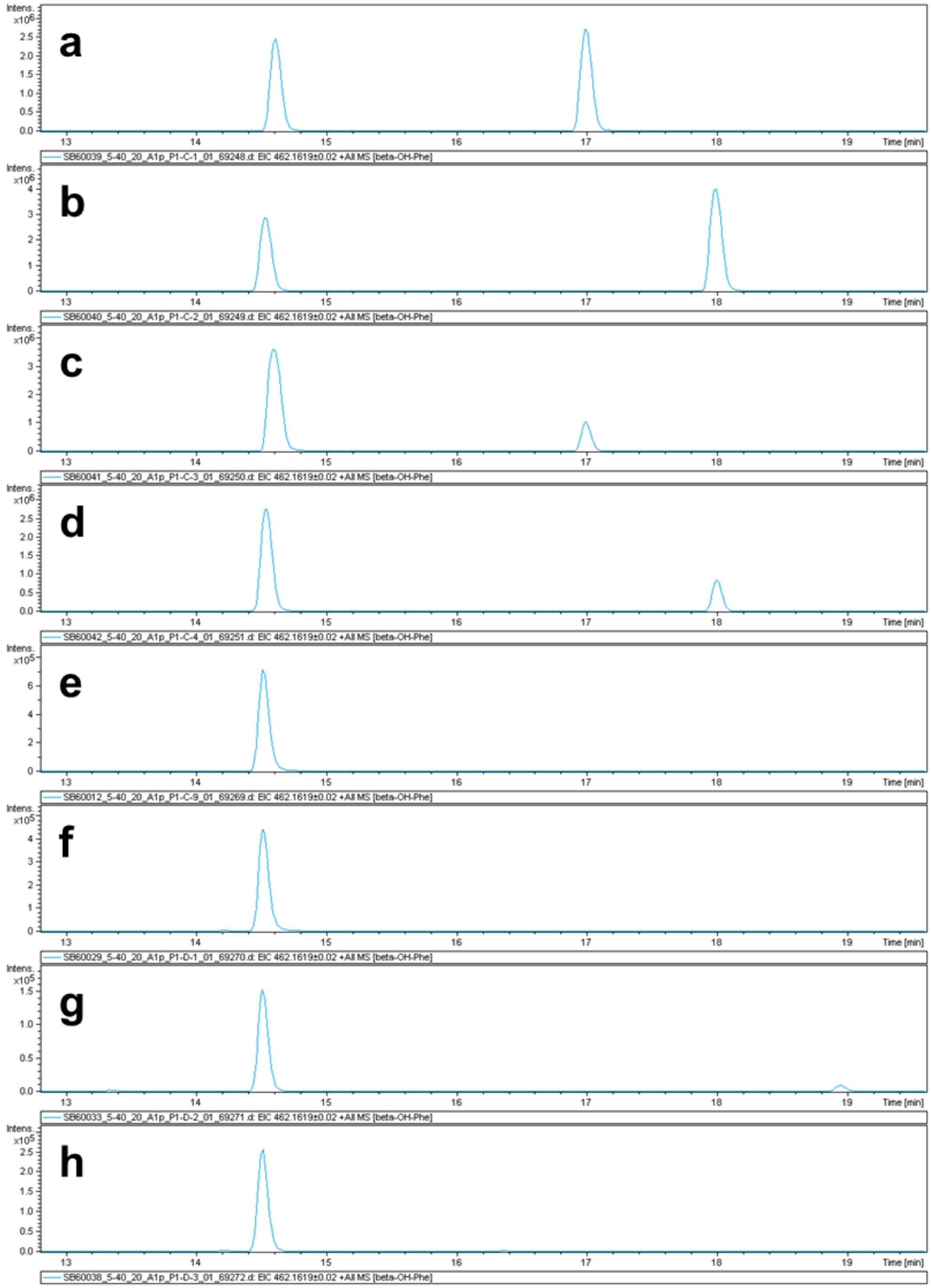


**Fig. S36 C18-RP UHPLC-MS extracted ion chromatograms of β hydroxyphenylalanines derivatized with L-FDVA at *m/z* 462.1619 $[M+H]^+$ within synthesized β-hydroxyphenylalanine standards and chitinopeptin A-D samples.** (**A**) (2*S*,3*S*)- and (2*R*,3*R*)-3-hydroxyphenylalanine, (**B**) (2*S*,3*R*)- and (2*R*,3*S*)-3-hydroxyphenylalanine, (**C**) (2*S*,3*S*)-3-hydroxyphenylalanine, (**D**) (2*S*,3*R*)-3-hydroxyphenylalanine, (**E**) chitinopeptin A, (**F**) chitinopeptin B, (**G**) chitinopeptin C1+C2, (**H**) chitinopeptin D1+D2

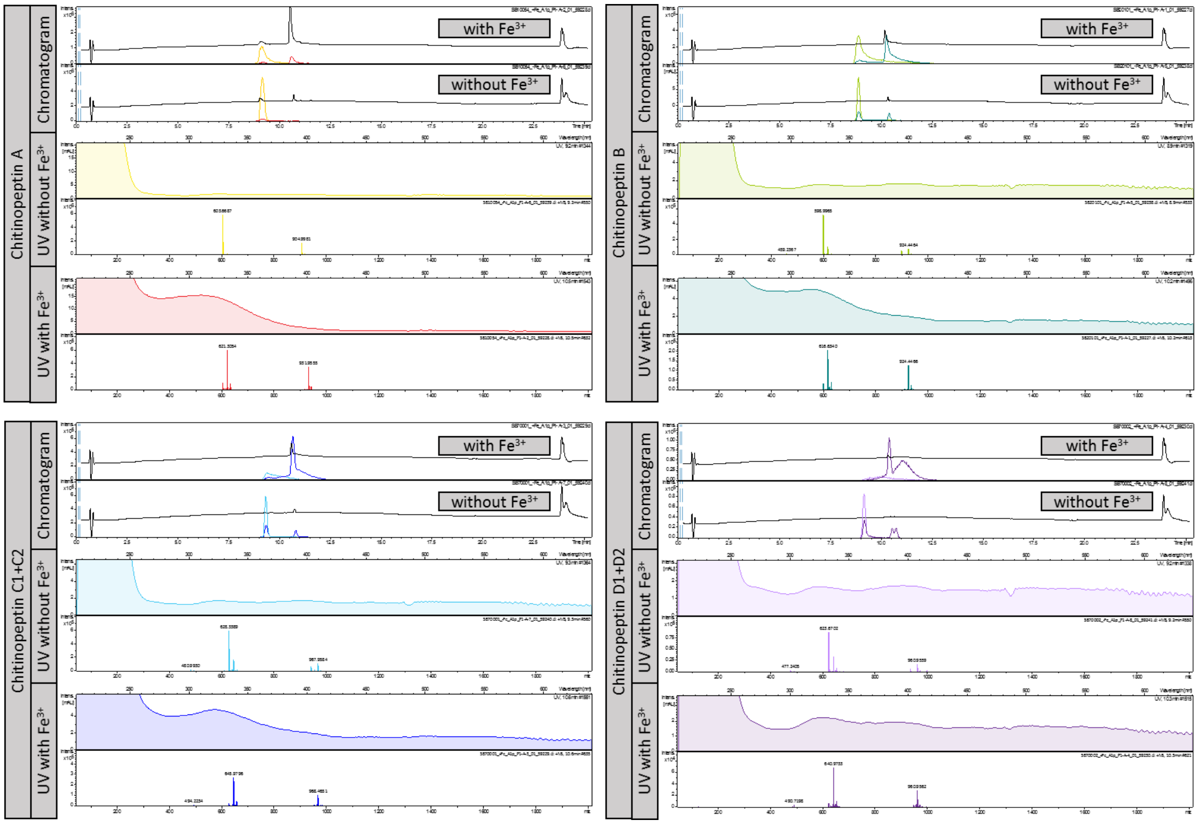


**Fig. S37 Iron chelating properties of chitinopeptin A-D.** Fe(III)Citrate was added in excess to pure compounds.

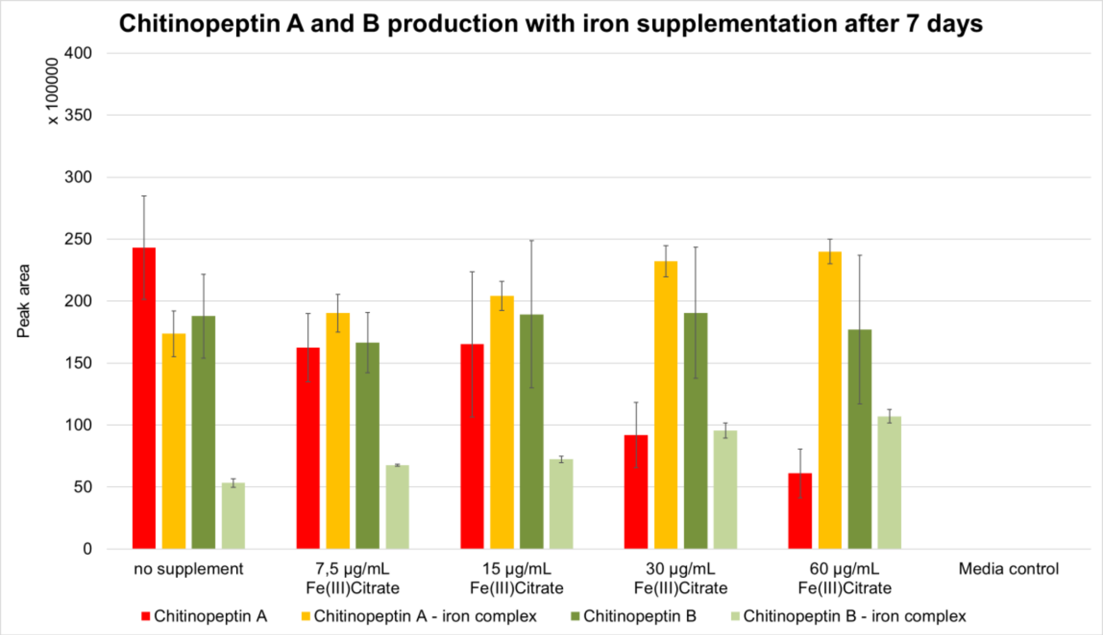


**Fig. S38** Production of chitinopeptin A and B in medium 3018 after 7 days of cultivation in 24 well plate cultivation.

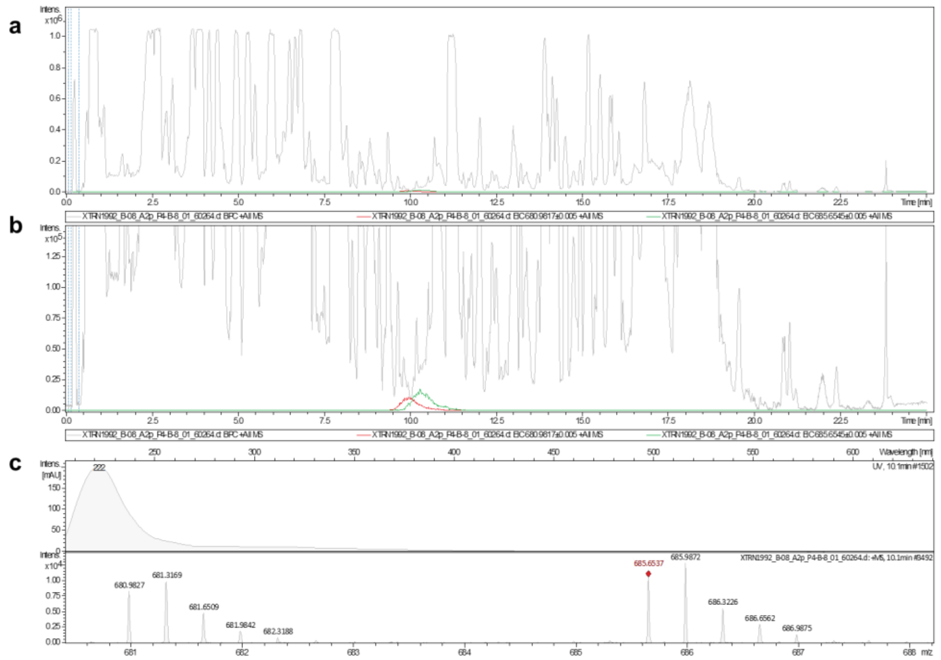


**Fig. S 39 Analytics of putatively cyclic lipodepsipeptides produced by *C. niastensis* DSM 24859 with m/z of 680.9817 $[M+3H]^{3+}$ and 685.6545 $[M+3H]^{3+}$.** (**A**) overlaid Base Peak Chromatogram (BPC, grey) and Extracted Ion Chromatogram (EIC) of *m/z* 680.9817 $[M+3H]^{3+}$ (red) and 685.6545 $[M+3H]^{3+}$ (green). (**B**) Zoom into the chromatogram. (**C**) Isotope pattern of both derivatives.

**Tab. S1** Strain list of all *Chitinophaga* strains used for the chemical barcoding matrix and bioactivity-guided NP discovery process.

| Phylum | Class | Order | Family | Genus | Species | Strain |
|---|---|---|---|---|---|---|
| Bacteroidetes | Chitinophagia | Chitinophagales | Chitinophagaceae | *Chitinophaga* | *alhagiae* | KCTC62518 |
| Bacteroidetes | Chitinophagia | Chitinophagales | Chitinophagaceae | *Chitinophaga* | *arvensicola* | DSM3695 |
| Bacteroidetes | Chitinophagia | Chitinophagales | Chitinophagaceae | *Chitinophaga* | *barathri* | KCTC42472 |
| Bacteroidetes | Chitinophagia | Chitinophagales | Chitinophagaceae | *Chitinophaga* | *caeni* | KCTC62265 |
| Bacteroidetes | Chitinophagia | Chitinophagales | Chitinophagaceae | *Chitinophaga* | *cymbidii* | KCTC23738 |
| Bacteroidetes | Chitinophagia | Chitinophagales | Chitinophagaceae | *Chitinophaga* | *dinghuensis* | DSM29821 |
| Bacteroidetes | Chitinophagia | Chitinophagales | Chitinophagaceae | *Chitinophaga* | *eiseniae* | DSM22224 |
| Bacteroidetes | Chitinophagia | Chitinophagales | Chitinophagaceae | *Chitinophaga* | *filiformis* | DSM527 |
| Bacteroidetes | Chitinophagia | Chitinophagales | Chitinophagaceae | *Chitinophaga* | *flava* | KCTC62435 |
| Bacteroidetes | Chitinophagia | Chitinophagales | Chitinophagaceae | *Chitinophaga* | *ginsengisegetis* | DSM18108 |
| Bacteroidetes | Chitinophagia | Chitinophagales | Chitinophagaceae | *Chitinophaga* | *ginsengisoli* | DSM18107 |
| Bacteroidetes | Chitinophagia | Chitinophagales | Chitinophagaceae | *Chitinophaga* | *japonensis* | DSM13484 |
| Bacteroidetes | Chitinophagia | Chitinophagales | Chitinophagaceae | *Chitinophaga* | *jiangningensis* | DSM27406 |
| Bacteroidetes | Chitinophagia | Chitinophagales | Chitinophagaceae | *Chitinophaga* | *niabensis* | DSM24787 |
| Bacteroidetes | Chitinophagia | Chitinophagales | Chitinophagaceae | *Chitinophaga* | *niastensis* | DSM24859 |
| Bacteroidetes | Chitinophagia | Chitinophagales | Chitinophagaceae | *Chitinophaga* | *pinensis* | DSM2589 |
| Bacteroidetes | Chitinophagia | Chitinophagales | Chitinophagaceae | *Chitinophaga* | *pinensis* | DSM2588 |
| Bacteroidetes | Chitinophagia | Chitinophagales | Chitinophagaceae | *Chitinophaga* | *rupis* | DSM21039 |
| Bacteroidetes | Chitinophagia | Chitinophagales | Chitinophagaceae | *Chitinophaga* | *sancti* | DSM784 |
| Bacteroidetes | Chitinophagia | Chitinophagales | Chitinophagaceae | *Chitinophaga* | *sedimenti* | KCTC52590 |
| Bacteroidetes | Chitinophagia | Chitinophagales | Chitinophagaceae | *Chitinophaga* | *silvisoli* | KCTC62860 |
| Bacteroidetes | Chitinophagia | Chitinophagales | Chitinophagaceae | *Chitinophaga* | *skermanii* | DSM23857 |
| Bacteroidetes | Chitinophagia | Chitinophagales | Chitinophagaceae | *Chitinophaga* | *sp.* | DSM18078 |
| Bacteroidetes | Chitinophagia | Chitinophagales | Chitinophagaceae | *Chitinophaga* | *terrae* | DSM23920 |
| Bacteroidetes | Chitinophagia | Chitinophagales | Chitinophagaceae | *Chitinophaga* | *varians* | KCTC52926 |

**Tab. S2 |** $^{1}$H-NMR spectroscopic data of chitinopeptins in a mixture of $H_2O$ and $CD_3CN$ in a ratio of 1:1. Chitinopeptin A: 700.13 MHz, 299 K; chitinopeptins B-D: 500.30 MHz, 300 K; $^{1}$H-chemical shifts are referenced to sodium-3-(trimethylsilyl)propionate-2,2,3,3-$d_4$. The chitinopeptin D sample was a mixture of at least four derivatives with chitinopeptin D1 being the major component. Only the relevant part (Val$_{10}$-Dab$_9$) of chitinopeptin D2 was assigned due to the signal overlap.[a]

| Pos. | Chitinopeptin A $\delta_H$, mult. [J (Hz)] | Chitinopeptin B $\delta_H$, mult. [J (Hz)] | Chitinopeptin C1 $\delta_H$, mult. [J (Hz)] | Chitinopeptin C2 $\delta_H$, mult. [J (Hz)] | Chitinopeptin D1 $\delta_H$, mult. [J (Hz)] | Chitinopeptin D2 $\delta_H$, mult. [J (Hz)] |
|---|---|---|---|---|---|---|
| **FA** | | | | | | |
| 1 | - | - | - | - | - | |
| 2 | 3.22, m | 3.23, m | 2.44/2.35, m | 2.42/2.34, m | 2.41/2.35, m | |
| 2-Me | 1.26, m | 1.26, m | - | | - | |
| 3 | 3.27, m | 3.26,m | 3.98, b | 3.96, b | 3.96, m | |
| 4 | 1.61/1.51, m | 1.61/1.52, m | 1.44, m | 1.44, m | 1.44, m | |
| 5 | 1.34/1.27, m | 1.33/1.27, m | 1.38/1.29, m/b | 1.38/1.29, m/b | 1.38/1.28, m | |
| 6 | 1.27/1.21, m | 1.29/1.24, m | 1.28, b | 1.28, b | 1.28, m | |
| 7 | 1.27, m | 1.27, m | 1.29, b | 1.29, b | 1.29, m | |
| 8 | 1.17, m | 1.27, m | 1.18, b | 1.18, b | 1.18, b | |
| 9 | 1.52, m | 1.30, m | 1.53, m | 1.53, b | 1.53, b | |
| 10 | 0.88, m | 0.90, m | 0.88, d (6.7) | 0.88, d (6.7) | 0.89, m | |
| 11 | 0.88, m | - | 0.88, (6.7) | 0.88, d (6.7) | 0.89, m | |
| **Dab0** | | | | | | |
| NH | | | 8.37, d (8.1) | 8.345, b | 8.34, b | |
| α | | | 5.24, dd (7.2/15.1) | 5.12, dd (7.5/13.8) | 5.11, b | |
| β | | | 3.37/3.20, m | 3.36/3.22, m | 3.36/3.20, m | |
| C‘ | | | - | - | - | |
| **NMe-Val1** | | | | | | |
| NMe | 3.09, s | 3.09, s | 3.01, s | 3.05, s | 3.06, s | |
| α | 4.67, d (11.1) | 4.67, d (11.3) | 4.56, d (10.9) | 4.55, d (10.9) | 4.55, d (6.1) | |
| β | 2.26, m | 2.26, m | 2.23, m | 2.24, m | 2.24, m | |
| γ | 1.02, d (6.7) | 1.02, d (6.7) | 0.96, b | 0.98, d (3.1) | 0.97, m | |
| γ' | 0.88, m | 0.88, m | 0.85, m | 0.85, m | 0.85, m | |
| C‘ | - | - | - | - | - | |
| **Thr2** | | | | | | |
| NH | 8.35, d (7.5) | 8.35, d (5.4) | 8.37, b | 8.36, b | 8.37, b | |
| α | 4.75, m | 4.75, m | 4.63, m | 4.73, m | 4.78, m | |
| β | 5.23, m | 5.23, m | 5.31, m | 5.30, m | 5.30, m | |
| γ | 1.17, d (6.5) | 1.17, d (6.6) | 1.31, d (5.5) | 1.24, d (7.5) | 1.23, m | |
| C‘ | - | - | - | - | - | |
| **Ala3** | | | | | | |
| NH | 8.22 (b) | 8.22, d (4.6) | 7.91, d (7.5) | 8.02 | 8.05, d (5.9) | |
| α | 4.39, m | 4.39, m | 4.41, b | 4.415 | 4.40, b | |
| β | 1.33, d (7.0) | 1.33, d (7.0) | 1.33, d (5.1) | 1.34 | 1.34, d (9.4) | |
| C‘ | - | - | - | - | - | |
| **β-OH-Asp4** | | | | | | |
| NH | 8.13, d (8.9) | 8.13, d (9.3) | 8.02, b | 8.07, b | 8.06, b | |
| α | 4.83,m | 4.83, m | 4.96, m | 4.895, m | 4.89, b | |
| β | 4.50, m | 4.51, b | 4.42, d (7.0) | 4.48, b | 4.46, b | |
| γ | - | - | - | - | - | |
| C‘ | - | - | - | - | - | |
| **β-OH-Phe5** | | | | | | |
| NH | 8.34, 3 (6.8) | 8.34, 2 (3.4) | 8.38, b | 8.315, d (5.8) | 8.39, b | |
| α | 4.72, m | 4.72, m | 4.68, m | 4.69, m | 4.70, m | |
| β | 5.05, d (9.5) | 5.05, d (9.4) | 4.99, d (8.3) | 4.95, d (8.8) | 4.98, d (9.0) | |
| γ | - | - | - | - | - | |
| δ | 7.44, m | 7.44, m | 7.42, m | 7.44, m | 7.44, m | |
| ε | 7.37, m | 7.37, m | 7.39, m | 7.39, m | 7.39, m | |
| ζ | 7.35,m | 7.35, m | 7.35, m | 7.35, m | 7.35, m | |
| C‘ | - | - | - | - | - | |
| **Ile6** | | | | | | |
| NH | 7.88, d (8.1) | 7.88, d (7.8) | 7.99, d (7.7) | 7.98, d (7.9) | 7.97, b | |
| α | 4.12, d (4.5) | 4.11, m | 4.14, m | 4.13, m | 4.13, m | |
| β | 1.75, m | 1.75, m | 1.77, m | 1.77, m | 1.76, m | |
| β-Me | 0.63, d (7.2) | 0.62, d (7.1) | 0.57, d (7.5) | 0.59, d (7.8) | 0.59, b | |
| γ | 0.54/0.49, m/m | 0.54/0.49, m/m | 0.69/0.48, m/m | 0.60/0.48, m/m | 0.59/0.50, m/m | |
| δ | 0.56, m | 0.56, m | 0.67, t (6.4) | 0.58, m | 0.59, t (7.5) | |
| C‘ | - | - | - | - | - | |

**Tab. S2** (continued)

| Pos. | Chitinopeptin A $\delta_H$, mult. [J (Hz)] | Chitinopeptin B $\delta_H$, mult. [J (Hz)] | Chitinopeptin C1 $\delta_H$, mult. [J (Hz)] | Chitinopeptin C2 $\delta_H$, mult. [J (Hz)] | Chitinopeptin D1 $\delta_H$, mult. [J (Hz)] | Chitinopeptin D2 $\delta_H$, mult. [J (Hz)] |
|---|---|---|---|---|---|---|
| **Ser7** | | | | | | |
| NH | 7.77, d (6.1) | 7.77, d (6.1) | 8.12, d (6.6) | 7.92, d (8.0) | 7.90, b | |
| α | 4.28, m | 4.29, m | 4.32, b | 4.32, b | 4.32, b | |
| β | 3.93/3.84, d/d (4.4/4.4) | 3.92/3.85 (4.2/4.4) | 3.87, dd (4.6/12.1) | 3.92/3.86, b/b | 3.90/3.85, d/b (5.8/-) | |
| C' | - | - | - | - | - | |
| **Lys8** | | | | | | |
| NH | 8.65, d (8.1) | 8.65, d (7.9) | 8.095, b | 8.64, d (7.2) | 8.59, d (7.6) | |
| α | 4.39, m | 4.38, m | 4.34, m | 4.36, m | 4.33, m | |
| β | 1.93/1.78, m/m | 1.93/1.77, m/m | 1.85/1.64, m/m | 1.90/1.79, m/m | 1.88/1.78, m/m | |
| γ | 1.44/1.38, m/m | 1.41, m | 1.35, m | 1.42, m | 1.41, m | |
| δ | 1.64, m | 1.64, m | 1.64, m | 1.64, m | 1.63, m | |
| ε | 2.95, m | 2.95, m | 2.93, m | 2.94, m | 2.93, m | |
| C' | - | - | - | - | - | |
| **β-$NH_2$-Ala9/ β-Dab9/Dab9** | | | | | | |
| NH | 8.92, d (8.3) | 8.90, d (8.5) | 8.17, d (5.5) | 8.84, d (8.4) | 8.93, d (8.3) | 3.72/3.59 |
| α | 4.79, m | 4.79 m | 3.72/3.56, m | 4.82, m | 4.82, m | 4.23 |
| β | 3.38/3.34, d/d (7.3/5.0) | 3.36, m | 4.24, m | 3.37, m | 3.37, m | - |
| C' | - | - | - | - | - | - |
| **Leu/Ile/Val10** | | | | | | |
| NH | 7.94, d (7.3) | 7.94, d (6.7) | 8.71, d (6.9) | 7.68, d (7.6) | 7.73, d (7.9) | 8.74, d (5.8) |
| α | 4.37, m | 4.37, m | 4.21, m | 4.34, m | 4.15, m | 4.01, m |
| β | 1.57, m | 1.57, m | 1.88, m | 1.89, m | 2.04, (a) | 2.03, (a) |
| γ | 1.56, m | 1.56, m | 1.36/1.17, m/m | 1.34/1.15, m/m | 0.89, d (1.6) | 0.97, b |
| γ' | - | - | 0.93, m | 0.83, m | 0.88, d (1.6) | 0.90, b |
| δ | 0.91, d (6.3) | 0.90, d (6.5) | 0.89, m | 0.89, m | | |
| δ' | 0.87, m | 0.88, m | - | - | | |
| C' | - | - | - | - | | |
| **β-OH-Asp11** | | | | | | |
| NH | 7.97, d (8.8) | 7.98, d (8.5) | 7.97, d (8.0) | 7.98, d (8.0) | 7.97, b | |
| α | 4.70, m | 4.70, m | 4.74, b | 4.75, b | 4.75, d (2.8) | |
| β | 4.41, b | 4.43, b | 4.52, b | 4.45, b | 4.46, b | |
| γ | - | - | - | - | - | |
| C' | - | - | - | - | - | |
| **Leu12** | | | | | | |
| NH | 8.14, b | 8.15, d (7.9) | 8.05, d (8.0) | 8.10, d (7.6) | 8.02, b | |
| α | 4.42, b | 4.42, b | 4.45, b | 4.466, b | 4.47, b | |
| β | 1.64, m | 1.65, m | 1.67, m | 1.65, m | 1.65, m | |
| γ | 1.64, m | 1.65, m | 1.67, m | 1.65, m | 1.67, m | |
| δ | 0.93, d (6.3) | 0.93, d (6.3) | 0.94, b | 0.927, d (2.1) | 0.93, b | |
| δ' | 0.86, m | 0.86, m | 0.88, b | 0.86, b | 0.87, b | |
| C' | - | - | - | - | - | |
| **β-OH-Asp13** | | | | | | |
| NH | 7.68, d (5.9) | 7.69, d (8.9) | 7.78, d (8.6) | 7.71, b | 7.71, d (9.3) | |
| α | 4.84, m | 4.85, m | 4.79, m | 4.82, m | 4.82, m | |
| β | 4.33, b | 4.35, m | 4.36, m | 4.38, m | 4.36, m | |
| γ | - | - | - | - | - | |
| C' | - | - | - | - | - | |
| **β-OH-Ile14** | | | | | | |
| NH | 7.88, d (8.3) | 7.88, d (7.9) | 7.83, d (8.5) | 7.89, d (9.3) | 7.86, d (8.6) | |
| α | 4.40, b | 4.41, b | 4.42, b | 4.41, b | 4.40, b | |
| β | - | - | - | - | - | |
| β-Me | 1.25, s | 1.25, s | 1.21, s | 1.23, s | 1.23, s | |
| γ | 1.58/1.53, m | 1.58/1.54, m | 1.55, m | 1.57, m | 1.56, m | |
| δ | 0.85, m | 0.86, m | 0.85, m | 0.86, m | 0.86, m | |
| C' | - | - | - | - | - | |

[a]Abbreviations: b = broad signal; a = $^1$H-signal below solvent signal; n.a. = not assigned

**Tab. S3** $^{13}$C-NMR spectroscopic data of chitinopeptins A-D in a mixture of $D_2O$ and $CD_3CN$ in a ratio of 1:1. Chitinopeptin A: 176.05 MHz, 299 K; chitinopeptins B-D: 125.82 MHz, 300 K. $^{13}$C-chemical shifts were referenced to the solvent signal ($CD_3CN$, $^{13}$C: 1.30 ppm). The chitinopeptin D sample was a mixture of at least four derivatives with chitinopeptin D1 being the major component. Only the relevant part ($Val_{10}$-$Dab_9$) of chitinopeptin D2 was assigned due to the signal overlap.[a]

| Pos. | Chitinopeptin A $\delta_C$, mult. | Chitinopeptin B $\delta_C$, mult. | Chitinopeptin C1 $\delta_C$, mult. | Chitinopeptin C2 $\delta_C$, mult. | Chitinopeptin D1 $\delta_C$, mult. | Chitinopeptin D2 $\delta_C$, mult. |
|---|---|---|---|---|---|---|
| **FA** | | | | | | |
| 1 | 177.81, C | 177.79, C | 174.27,C | 174.35, C | n.a. | |
| 2 | 36.08, CH | 36.05, CH | 43.85, $CH_2$ | 43.80, $CH_2$ | 43.80, $CH_2$ | |
| 2-Me | 14.77, $CH_3$ | 14.76, $CH_3$ | - | - | - | |
| 3 | 55.85, CH | 55.60, CH | 69.19, CH | 69.18, CH | 69.19 | |
| 4 | 32.37, $CH_2$ | ~32.2 (b), $CH_2$ | 37.49, $CH_2$ | 37.49, $CH_2$ | 37.48, CH | |
| 5 | 26.29, $CH_2$ | 26.25, $CH_2$ | 26.02, $CH_2$ | 25.99, $CH_2$ | 25.99, $CH_2$ | |
| 6 | 30.11, $CH_2$ | 29.79, $CH_2$ | 30.28, $CH_2$ | 30.28, $CH_2$ | 30.28, $CH_2$ | |
| 7 | 27.56, $CH_2$ | 29.46, $CH_2$ | 27.83, $CH_2$ | 27.83, $CH_2$ | 27.83, $CH_2$ | |
| 8 | 39.40, $CH_2$ | 32.24, $CH_2$ | 39.52, $CH_2$ | 39.52, $CH_2$ | 39.52, $CH_2$ | |
| 9 | 28.37, CH | 23.09, $CH_2$ | 28.47, CH | 28.47, CH | 28.47, $CH_2$ | |
| 10 | 22.84, $CH_3$ | 14.31, $CH_3$ | 22.88, $CH_3$ | 22.88, $CH_3$ | 22.88, $CH_3$ | |
| 11 | 22.81, $CH_3$ | - | 22.88, $CH_3$ | 22.88, $CH_3$ | 22.88, $CH_3$ | |
| **Dab0** | | | | | | |
| NH | - | | - | - | - | |
| α | - | | 48.52, CH | 48.67, CH | 48.68, CH | |
| β | - | | 41.02, $CH_2$ | 41.07, $CH_2$ | 41.04, $CH_2$ | |
| C‘ | - | | 171.75, C | 171.76, C | 171.76, C | |
| **N-Me-Val1** | | | | | | |
| NMe | 31.64, $CH_3$ | 31.65, $CH_3$ | 31.44, $CH_3$ | 31.60, $CH_3$ | 31.65, $CH_3$ | |
| α | 63.22, CH | 63.21, CH | 63.88, CH | 64.14, CH | 64.17, CH | |
| β | 26.70, CH | 26.71, CH | 26.98, CH | 26.90, CH | 26.93, CH | |
| γ | 19.74, $CH_3$ | 19.73, $CH_3$ | 19.51, $CH_3$ | 19.60, $CH_3$ | 19.53, $CH_3$ | |
| γ' | 19.30, $CH_3$ | 19.28, $CH_3$ | 19.06, $CH_3$ | 19.23, $CH_3$ | 19.09, $CH_3$ | |
| C‘ | 172.16, C | 172.11, C | 171.90, C | 172.17, C | 171.76, C | |
| **Thr2** | | | | | | |
| NH | - | - | - | - | - | |
| α | ~56.4 (b), CH | ~56.3 (b), CH | 57.14, CH | 56.56, CH | 56.34, CH | |
| β | 73.06, CH | 73.03, CH | 71.96, CH | 72.21, CH | 72.10, CH | |
| γ | 15.40, $CH_3$ | 15.44, $CH_3$ | 16.54, $CH_3$ | 15.90, $CH_3$ | 15.75, $CH_3$ | |
| C‘ | 169.65, C | 169.57, C | 169.91, C | 169.88, C | 169.90, C | |
| **Ala3** | | | | | | |
| NH | - | - | - | - | - | |
| α | 51.12, CH | 50.98, CH | 50.03, CH | 50.48, CH | 50.55, CH | |
| β | 17.90, $CH_3$ | 17.86, $CH_3$ | 18.93, $CH_3$ | 18.34, $CH_3$ | 18.20, $CH_3$ | |
| C‘ | 175.19, C | 175.10, C | 174.50, C | 174.97, C | 175.00, C | |
| **β-OH-Asp4** | | | | | | |
| NH | - | - | - | - | - | |
| α | 57.36, CH | 57.24, CH | 56.59, CH | 56.86, CH | 56.84, CH | |
| β | 72.56, CH | 72.45, CH | 72.85, CH | 72.00, CH | 72.44, CH | |
| γ | 176.69, C | 176.7 (b), C | 177.11, C | 176.76, C | n.a. | |
| C‘ | 173.27, C | 173.20, C | 172.47, C | 172.95, C | n.a. | |
| **β-OH-Phe5** | | | | | | |
| NH | - | - | - | - | - | |
| α | 61.62, CH | 61.50, CH | 61.26, CH | 61.64, CH | 61.53, CH | |
| β | 73.78, CH | 73.66, CH | 73.91, CH | 73.91, CH | 73.95, CH | |
| γ | 140.39, C | 140.34, CH | 140.32, C | 140.17, C | 140.17, C | |
| δ | 128.23,CH | 128.25, CH | 128.05, CH | 128.26, CH | 128.25, C | |
| ε | 129.49, CH | 129.52, CH | 129.58, CH | 129.64, CH | ~129.6 (a), CH | |
| ζ | 129.38, CH | 129.42, CH | 129.42, CH | 129.53, CH | ~129.5 (a), CH | |
| C‘ | 171.29, C | 171.24, CH | 171.58, C | 171.65, C | 171.57, C | |
| **Ile6** | | | | | | |
| NH | - | - | - | - | - | |
| α | 57.48, CH | 57.39, CH | 57.89, CH | 57.47, CH | 57.49, CH | |
| β | 36.57, CH | 36.55, CH | 36.70, CH | 36.50, CH | 36.55, CH | |
| β-Me | 14.46, $CH_3$ | 14.45, $CH_3$ | 14.58, $CH_3$ | 14.42, $CH_3$ | 14.45, $CH_3$ | |
| γ | 26.21, $CH_2$ | 26.22, $CH_2$ | 26.12, $CH_2$ | 26.12, $CH_2$ | 26.13, $CH_2$ | |
| δ | 12.16, $CH_3$ | 12.15, $CH_3$ | 11.86, $CH_3$ | 12.05, $CH_3$ | 12.04, $CH_3$ | |
| C‘ | 174.31, C | 174.25, C | 174.17, C | 174.12, C | 174.11, C | |

**Tab. S3** (continued)

| Position | Chitinopeptin A $\delta_C$, mult. | Chitinopeptin B $\delta_C$, mult. | Chitinopeptin C1 $\delta_C$, mult. | Chitinopeptin C2 $\delta_C$, mult. | Chitinopeptin D1 $\delta_C$, mult. | Chitinopeptin D2 $\delta_C$, mult. |
|---|---|---|---|---|---|---|
| **Ser7** | | | | | | |
| NH | - | - | - | - | - | |
| α | 56.71, CH | 56.60, CH | 57.02, CH | 56.98, CH | 56.98, CH | |
| β | 62.55, $CH_2$ | 62.40, $CH_2$ | 62.09, $CH_2$ | 62.40, $CH_2$ | 62.36, $CH_2$ | |
| C‘ | 173.16, C | 173.10, C | 172.77, C | 173.09, C | 173.09, C | |
| **Lys8** | | | | | | |
| NH | - | - | - | - | - | |
| α | 54.75, CH | 54.64, CH | 53.98, CH | 54.93, CH | 55.03 | |
| β | 31.26, $CH_2$ | 31.20, $CH_2$ | 31.07, $CH_2$ | 31.18, $CH_2$ | 31.16, $CH_2$ | |
| γ | 22.97, $CH_2$ | 22.97, $CH_2$ | 22.92, $CH_2$ | 22.95, $CH_2$ | 22.97, $CH_2$ | |
| δ | 27.06, $CH_2$ | 26.91, $CH_2$ | 26.88, $CH_2$ | 26.98, $CH_2$ | 26.98, $CH_2$ | |
| ε | 40.31, $CH_2$ | 40.02, $CH_2$ | 39.91, $CH_2$ | 40.04, $CH_2$ | 39.93, $CH_2$ | |
| C‘ | 175.32, C | 175.25, C | 174.85, C | 175.46, C | 175.48, C | |
| **β-$NH_2$-Ala9/ β-Dab9/Dab9** | | | | | | |
| NH | - | - | - | - | - | - |
| α | 51.28, CH | 51.03, CH | 41.07, CH | 51.05, CH | 50.99, CH | 41.04, |
| β | 40.76, $CH_2$ | 40.48, $CH_2$ | 53.70, $CH_2$ | 40.34, $CH_2$ | 40.31, $CH_2$ | 53.67 |
| C‘ | 170.76, C | 170.67, C | 168.96, C | 170.78, C | 170.71, C | ~169.0 (a) |
| **Leu/Ile/Val10** | | | | | | |
| NH | - | - | - | - | - | - |
| α | 53.64, CH | 53.54, CH | 59.84, CH | 58.62, CH | 60.52, CH | ~61.7 (a) |
| β | 40.57, $CH_2$ | 40.52, $CH_2$ | 36.61, CH | 37.43, CH | 31.00, CH | ~ 30.3(a) |
| γ | 25.18, CH | 25.20, CH | 15.33, $CH_2$ | 14.97, $CH_2$ | 19.30, $CH_2$ | ~19.3 (a) |
| γ' | - | - | 26.34, $CH_3$ | 26.53, $CH_3$ | 18.60, $CH_3$ | ~19.3 (a) |
| δ | 23.12, $CH_3$ | 23.12, $CH_3$ | 11.68, $CH_3$ | 11.79, $CH_3$ | - | - |
| δ' | 21.82, $CH_3$ | 21.80, $CH_3$ | - | - | - | - |
| C‘ | 174.57, C | 174.49, C | 174.02, C | 173.47, C | 173.55, C | 174.17 |
| **β-OH-Asp11** | | | | | | |
| NH | - | - | - | - | - | |
| α | 57.18, CH | 57.05, CH | 57.42, CH | 57.20, CH | 57.37, CH | |
| β | 72.06, CH | 71.94, CH | 72.29, CH | 72.48, CH | 72.10, CH | |
| γ | 176.80, C | ~176.8 (b), C | 176.68, C | 176.76, C | n.a. | |
| C‘ | 172.33, C | 172.25, C | 172.05, C | 172.29, C | n.a. | |
| **Leu12** | | | | | | |
| NH | - | - | - | - | - | |
| α | 53.23, CH | 53.11, CH | 53.14, CH | 52.91, CH | ~53.0 (a), CH | |
| β | 40.37, $CH_2$ | 40.33, $CH_2$ | 40.71, $CH_2$ | 40.46, $CH_2$ | 40.57, $CH_2$ | |
| γ | 25.18, CH | 25.18, CH | 25.23, CH | 25.14, CH | 25.15, CH | |
| δ | 23.34, $CH_3$ | 23.34, $CH_3$ | 23.37, $CH_3$ | 23.37, $CH_3$ | 23.36, $CH_3$ | |
| δ' | 21.52, $CH_3$ | 21.51, $CH_3$ | 21.58, $CH_3$ | 21.49, $CH_3$ | 21.53, $CH_3$ | |
| C‘ | 174.55, C | 174.49, C | 174.66, C | 174.58, C | n.a. | |
| **β-OH-Asp13** | | | | | | |
| NH | - | - | - | - | - | |
| α | 56.71, CH | 56.60, CH | 56.86, CH | 56.81, CH | ~56.8 (a), CH | |
| β | 72.88, CH | 72.77, CH | 72.51, CH | 72.68, CH | ~72.7 (a), CH | |
| γ | 176.82, C | ~176.8 (b), C | 176.88, C | 176.88, C | n.a. | |
| C‘ | 171.94, C | 171.87, C | 171.84, C | 171.98, C | n.a. | |
| **β-OH-Ile14** | | | | | | |
| NH | - | - | - | - | - | |
| α | 60.68, CH | 60.59, CH | 60.04, CH | 60.29, CH | 60.35, CH | |
| β | 74.68, C | 74.61, C | 74.72, C | 74.63, C | 74.58, C | |
| β-Me | 23.34, $CH_3$ | 23.28, $CH_3$ | 23.25, $CH_3$ | 23.28, $CH_3$ | 23.26, $CH_3$ | |
| γ | 32.69, $CH_2$ | 32.65, $CH_2$ | 32.61, $CH_2$ | 32.64, $CH_2$ | 32.64, $CH_2$ | |
| δ | 8.41, $CH_3$ | 8.39, $CH_3$ | 8.35, $CH_3$ | 8.35, $CH_3$ | 8.35, $CH_3$ | |

[a]Abbreviations: b = broad signal; a = $^{13}$C-signal below solvent signal; n.a. = not assigned

**Tab. S4** Bioactivity screening data of chitinopeptins A-D.*

| MIC (µg/mL) | | Compounds | Chitinopeptin | | | | | | | | Reference data | | |
|---|---|---|---|---|---|---|---|---|---|---|---|---|---|
| | | Derivatives | A | | B | | C1+C2 | | D1+D2 | | Rifamicin | Tetracycline | Gentamycin/ Isoniazid[a] |
| | | Iron | - | + | - | + | - | + | - | + | Tebuconazole[b] | Amphotericin B[c] | Nystatin[d] |
| Organism and genotype | Gram - | *E. coli* ATCC 35218 | >64 | n.d. | >64 | n.d. | >64 | n.d. | >64 | n.d. | 4 | 4 | 0.125 |
| | | *E. coli* ATCC 25922 ΔTolC | >64 | n.d. | >64 | n.d. | >64 | n.d. | >64 | n.d. | 2 | 0.5 | 0.25 |
| | | *E. coli* MG1655 | >256 | n.d. | n.d. | n.d. | n.d. | n.d. | n.d. | n.d. | 16 | 0.5 | 0.5 |
| | | *P. aeruginosa* ATCC 27853 | >64 | n.d. | >64 | n.d. | >64 | n.d. | >64 | n.d. | 32 | 64 | 1 |
| | | *P. aeruginosa* PAO750 | >64 | n.d. | n.d. | n.d. | n.d. | n.d. | n.d. | n.d. | 16 | 0.25 | 0.125 |
| | | *K. pneumoniae* ATCC 13883 | >64 | n.d. | n.d. | n.d. | n.d. | n.d. | n.d. | n.d. | 8 | 2 | ≤0.031 |
| | | *M. catarrhalis* ATCC 25238 | 2 | 4 | 2 | n.d. | 2 | n.d. | 2 | n.d. | ≤0.031 | 1 | 0.125 |
| | | *A. baumannii* ATCC 19606 | 16 | n.d. | 64 | n.d. | 16 | n.d. | 16 | n.d. | 2 | 16 | 16 |
| | Gram + | *B. subtilis* DSM 10 | 4 | 8 | 4 | n.d. | 4 | n.d. | 4 | n.d. | ≤0.031 | 4 | ≤0.031 |
| | | *S. aureus* ATCC 25923 | >64 | n.d. | >64 | n.d. | >64 | n.d. | >64 | n.d. | ≤0.031 | 1 | 0.063 |
| | | *M. luteus* DSM 20030 | 32 | n.d. | 32 | n.d. | 32 | n.d. | 32 | n.d. | ≤0.031 | 0.5 | ≤0.031 |
| | | *L. monocytogenes* DSM 20600 | >64 | n.d. | n.d. | n.d. | n.d. | n.d. | n.d. | n.d. | ≤0.031 | 0.5 | ≤0.031 |
| | | *M. smegmatis* ATCC 607 | >64 | n.d. | >64 | n.d. | >64 | n.d. | >64 | n.d. | 8 | 0.5 | 1[a] |
| | Yeast/Fungi | *C. albicans* FH2173 | 4 | 32 | 8 | n.d. | 16 | n.d. | 16 | n.d. | 2[b] | 4[c] | 16[d] |
| | | *A. flavus* ATCC 9170 | >64 | n.d. | >64 | n.d. | 64 | n.d. | 64 | n.d. | 2[b] | 16-8[c] | 16d |
| | | *Z. tritici* MUCL45407 | 16 | 32-16 | 16 | n.d. | 16 | n.d. | 16-8 | n.d. | 0.5[b] | 0.031[c] | 0.5[d] |
| | | *F. oxysporum* ATCC 7601 | >64 | n.d. | >64 | n.d. | >64 | n.d. | >64 | n.d. | >64[b] | 4[c] | 4[d] |

*Abbreviation: n.d. = not determined

**Tab. S5** Partial Clustal W alignment(*106*) of the proposed dinuclear binding motif (His-X-His-X-Asp-His) by Makris *et al.* (2010)(*76*) in all three diiron-monooxygenases with other homologs in NRPS antibiotic biosynthetic pathways.

| Species | Antibiotic | Protein-ID | Consensus sequence | | | | | | |
|---|---|---|---|---|---|---|---|---|---|
| *Chitinophaga eiseniae* | Chitinopeptin A-B | SKA37886.1 | $H^{303}$-N-H-Q-D-H | $X_{66}$ | E | $X_{25}$ | D | $X_{26}$ | E |
| *Chitinophaga flava* | Chitinopeptin C-D | RBL90130.1 | $H^{303}$-N-H-Q-D-H | $X_{66}$ | E | $X_{25}$ | D | $X_{26}$ | E |
| *Chitinophaga oryziterrae* | / | MVT41610.1 | $H^{301}$-N-H-Q-D-H | $X_{66}$ | E | $X_{25}$ | D | $X_{26}$ | E |
| *Chitinophaga nistansis* | / | PSL45608.1 | $H^{303}$-N-H-Q-D-H | $X_{66}$ | E | $X_{25}$ | D | $X_{26}$ | E |
| *Streptomyces venezuelae* | Chloramphenicol | CCA54208.1 | $H^{305}$-N-H-Q-D-H | $X_{66}$ | E | $X_{25}$ | D | $X_{26}$ | E |
| *Actinoplanes teichomycetius* | Teicoplanin | CAE53366.1 | $H^{301}$-G-H-S-D-H | $X_{65}$ | E | $X_{25}$ | D | $X_{26}$ | E |
| *Nonomuraea sp.* | A40926 | CAD91223.1 | $H^{301}$-G-H-S-D-H | $X_{65}$ | E | $X_{25}$ | D | $X_{26}$ | E |
| *Streptomyces toyacaensis* | A47934 | AAM80528.1 | $H^{301}$-G-H-S-D-H | $X_{65}$ | E | $X_{25}$ | D | $X_{26}$ | E |
| *Burkholderia pyrrocinia* | Occidiofungin | KFL51884.1 | $H^{308}$-S-H-H-D-H | $X_{66}$ | E | $X_{25}$ | D | $X_{26}$ | E |
| *Lysobacter sp.* | Lysobactin | AEH59101.1 | $H^{303}$-P-H-Q-D-H | $X_{66}$ | E | $X_{25}$ | D | $X_{26}$ | E |
| *Burkholderia thailandensis* | Bactobolin | ABC35075.1 | $H^{300}$-S-H-H-D-H | $X_{66}$ | E | $X_{25}$ | D | $X_{26}$ | E |

**Tab. S6** Manual BLASTp search results between chitinopeptin enzymes no. 5+6 and reported enzymes to L-Dap biosynthesis.

| Organism | Related products | Homologous enzymes | Protein-ID | Identities (enzyme no. 5/ no. 6) | | | | Positives (enzyme no. 5/ no. 6) | | | | Ref. |
|---|---|---|---|---|---|---|---|---|---|---|---|---|
| | | | | *C. eiseniae* | *C. flava* | *C. oryziterrae* | *C. niastensis* | *C. eiseniae* | *C. flava* | *C. oryziterrae* | *C. niastensis* | |
| *Bacillus thuringiensis* | Zwittermycin A | Zwa5A/Zwa5B | ACM79805/ACM79806 | 48/ | 48/52 | 49/52 | 49/52 | 66/ | 66/73 | 68/71 | 66/71 | [82] |
| *Staphylococcus aureus* | Staphyloferrin B | SbnA/SbnB | TLW72643/TLW72644 | 37/ | 38/34 | 40/34 | 38/35 | 57/ | 58/54 | 60/53 | 58/54 | [77,78] |
| *Saccharothrix mutabilis* | Capreomycin | CmnB/CmnK | ABR67745/ABR67754 | 33/ | 32/24 | 33/24 | 31/23 | 50/ | 49/41 | 50/41 | 49/42 | [80,81] |
| *Streptomyces* sp. | Viomycin | VioB/VioK | AAP92492/AAP92501 | 30/ | 26/24 | 27/24 | 30/23 | 47/ | 47/41 | 49/40 | 49/42 | [79] |
| *Amycolatopsis japonicum* | [S,S]-EDDS | AesC/AesA | AIG74590/AIG74588 | 25/ | 26/17 | 28/18 | 26/- | 45/ | 44/37 | 47/37 | 46/- | [83] |
| *Streptomyces* sp. | EDHA | MA_5143a_00503/00505 | - | 23/ | 24/23 | 25/24 | 24/23 | 46/ | 45/52 | 47/48 | 45/48 | [84] |

**Tab. S7** Comparison of the amino acid prediction based on Stachelhaus code(*85*) and the assembled amino acids verified by NMR and Marfey's analysis. Amino acids marked in grey are most likely all L-Dap based on the same Stachelhaus code.

| *C. eiseniae* | | | | *C. flava* | | | | *C. oryziterrae* | | *C. niastensis* | |
|---|---|---|---|---|---|---|---|---|---|---|---|
| Chitinopeptin A & B | | predicted | | Chitinopeptin C1+C2 & D1+D2 | | predicted | | predicted | | predicted | |
| NMR | | AA | Stachelhaus | NMR | | AA | Stachelhaus | AA | Stachelhaus | AA | Stachelhaus |
| - | - | - | - | 2*S* | Dap | hydrophilic | DiWeLtadDK | hydrophilic | DiWeLtadDK | hydrophilic | DiWeLtadDK |
| 2*S* | *N*-Me-Val | Val (nMT) | DAiWmGGTFK | 2*S* | *N*-Me-Val | Val (nMT) | DAiWmGGTFK | Val (nMT) | DAiWmGGTFK | hydrophilic | DiWeLtadDK |
| 2*R*,3*R* | Thr | Thr (E) | DFWNIGMVHK | 2*R*,3*R* | Thr | Thr (E) | DFWNIGMVHK | Thr (E) | DFWNIGMVHK | Thr (E) | DFWNIGMVHK |
| 2*R* | Ala | Ser (E) | DVWHLSLIeK | 2*R* | Ala | Ser (E) | DVWHISLIEK | Ser (E) | DVWHISLIeK | X (Lys?) (E) | DAEhIGeVtK |
| 2*S*,3*S* | β-OH-Asp (Hya) | Asp | DLTKiGHIGK | 2*S*,3*S* | β-OH-Asp | Asp | DLTKiGHIGK | Asp | DLTKiGHIGK | Asp | DLTKiGHIGK |
| - | - | - | - | - | - | - | - | - | - | Tyr | DASTVAAVCK |
| - | - | - | - | - | - | - | - | - | - | Leu (E) | DAWFLGNVVK |
| - | - | - | - | - | - | - | - | - | - | Asp | DLTKiGHIGK |
| 2*S*,3*R* | β-OH-Phe | Phe | DAyTVAAVCK | 2*S*,3*R* | β-OH-Phe | Phe | DAyTVAAVCK | Phe | DAyTVAAVCK | Phe | DAyTVAAVCK |
| 2*R*,3*S* | Ile | Ile (E) | DAFFIGITFK | 2*R*,3*S* | Ile | Ile (E) | DAFFIGITFK | Val (E) | DAiWmGGTFK | - | - |
| 2*S* | Ser | Ser | DVWHLSLIDK | 2*S* | Ser | Ser | DVWHLSLIDK | Ser | DVWHLSLIDK | - | - |
| 2*R* | Lys | Tyr (E) | DAedVGEVVK | 2*R* | Lys | Tyr (E) | DAedVGEVVK | hydrophilic (E) | DAediGEiVK | - | - |
| 2*S* | Dap | hydrophilic | DiWeLtadDK | 2*S* | Dap | Ala | DiWeMtadDK | hydrophilic | DiWeLtadDK | hydrophilic | DiWeLtadDK |
| 2*R* | Leu | Leu (E) | DAWFLGNVVK | 2*R*,3*S* | Ile | Ile (E) | DAFFIGITFK | Leu (E) | DAWFLGNVVK | Leu (E) | DAWFLGNVVK |
| 2*S*,3*S* | β-OH-Asp (Hya) | Asp | DLTKiGHIGK | 2*S*,3*S* | β-OH-Asp | Asp | DLTKVGHIGK | Asp | DLTKiGHIGK | Asp | DLTKVGHIGK |
| 2*S* | Leu | Leu | DAWFLGNVVK | 2*S* | Leu | Leu | DAWFLGNVVK | Leu | DAWFLGNVVK | Ala | DiWeLtadDK |
| 2*S*,3*S* | β-OH-Asp (Hya) | Asp | DLTKiGHIGK | 2*S*,3*S* | β-OH-Asp | Asn | DLTKIGhVGK | Asp | DLTKiGHIGK | Asp | DLTKVGHIGK |
| - | - | - | - | - | - | - | - | - | - | Gly | DILQLGLIWK |
| 2*S*,3*R* | β-OH-Ile | Ile | DAFFIGITFK | 2*S*,3*R* | β-OH-Ile | Ile | DAFFIGITFK | Ile | DAFFIGITFK | Ile | DAFw LGVTFK |
