## Supplementary material for "Genomic and chemical decryption of the Bacteroidetes phylum for its potential to biosynthesize natural products": Synthesis of Hydroxy Amino Acid Supporting Material

**This PDF file includes:**

Synthesis of beta-hydroxy amino acids (Phe, Ile, Asp)

**Other Supplementary Materials for this manuscript include the following:**

R-script for chemotype-barcoding matrix
Figs. S1 to S39
Tables S1 to S7
Tab. S8

### 1. Synthetic procedures and compound characterization

#### 1.1. General Methods

All chemicals and solvents/anhydrous solvents were commercially supplied and used without further purification. Reactions were monitored using thin layer chromatography (TLC) or using Agilent 1100 series LCMS with UV detection at 254 nm and a low resonance electrospray mode (ESI). TLC was performed on pre-coated silica gel glass plates (Merck TLC Silica gel 60 F254) and compounds were detected under UV light (254 nm) and/or by staining with an aqueous solution of Phosphomolybdic acid, Cerium(IV) sulfate and $H_2SO_4$ followed by heating with a heat gun. Products were purified by using an automated flash column chromatography system (puriFlash® XS520Plus from Intechim) equipped with PF-15SIHC flash columns of different sizes from Interchim (eluants are given in parentheses). Alternatively, purification was performed by HPLC with a Waters AutoPurification HPLC/MS system using RP-18 columns with $H_2O$/TFA 0.1% as mobile phase A and acetonitrile as mobile phase B. The product containing fractions were collected and freeze-dried. NMR spectra were recorded on a Bruker AVANCE II WB spectrometer (400 MHz), a AVANCE III HD spectrometer (400 MHz) or a AVANCE III spectrometer (500 MHz) equipped with a TCI CryoProbe with $CDCl_3$, $D_2O$ or DMSO-$d_6$ as solvent with chemical shifts (δ) quoted in parts per million (ppm) and referenced to the solvent signal ($\delta^1H/^{13}C$: $CDCl_3$ 7.26/77.16, $\delta^1H$: $D_2O$ 4.79, $\delta^1H/^{13}C$: DMSO-$d_6$ 2.50 / 39.52 ppm) or TSPA ($\delta^1H/^{13}C$: 0.00/0.00) as external standard. Assignment was confirmed based on COSY, ROESY, HSQC and HMBC correlations. High resolution mass spectrometry was performed on a maXis II (Bruker) ESI TOF MS equipped with 1290 UPLC (Agilent) with DAD and ELSD. Specific rotation was measured by a polarimeter (P 3000 series) from Krüss.

#### 1.2. Preparation of the β-Hydroxyaspartic acids

All four stereoisomers of β-hydroxyaspartic acid were obtained according to a modified procedure described by Breuning *et al.*[1] Commercial available (−)-dibenzyl D-tartrate or (+)-dibenzyl L-tartrate were chosen as starting material since the benzyl esters allow for accurate *ee*-determination by chiral HPLC (due to their UV absorption at a detection wavelength of 207 nm) and traceless deprotection accompanied by azide reduction *via* catalytic hydrogenation as a last step. While the *anti*-isomers are directly accessible, the *syn*-isomers were obtained by selective base-induced epimerization of the azido-intermediates and separation of the two isomers by HPLC (Scheme S1). The high measured *ee* values of the products **SI2**, **SI4**, ***ent*-SI2** and ***ent*-SI4** proved a selective epimerization at the azido-bearing 2-position.

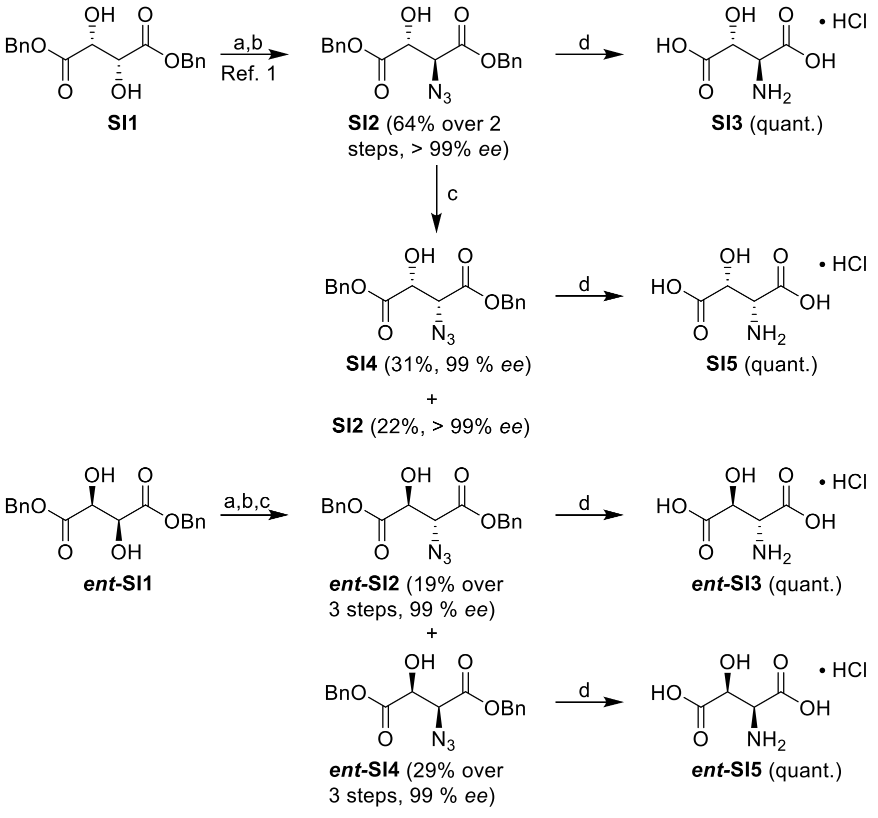

***Scheme S1*: Preparation of the β-Hydroxyaspartic acids, exemplarily shown for one diastereomeric series.**[a]

[a] Conditions: a) $SOCl_2$, DCM; b) $NaN_3$, DMF; c) DBU, THF, 40 °C; d) $H_2$, Pd/C; 1 M $HCl_{aq}$, 1,4-dioxane.

[1] Breuning, A., Vicik, R. & Schirmeister, T. An improved synthesis of aziridine-2,3-dicarboxylates via azido alcohols—epimerization studies. *Tetrahedron: Asymmetry* **14,** 3301–3312; 10.1016/j.tetasy.2003.09.015 (2003).

**Dibenzyl (2*S*,3*R*)-2-azido-3-hydroxysuccinate (SI2):**

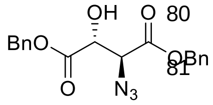

**SI2**

$SOCl_2$ (0.389 mL, 5.45 mmol, 1.20 eq.) was added to a solution of (+)-dibenzyl L –tartrate (**SI1**, 1.50 g, 4.54 mmol, 1.00 eq.) in anhydrous DCM (20 mL) at room temperature. A catalytic amount of anhydrous DMF (1 drop) was added and the reaction mixture was stirred for 7 h, until LC-MS indicated complete conversion. The reaction mixture was concentrated *in vacuo* and crude cyclic sulfite was used for the next step without further purification.

The crude product was dissolved in anhydrous DMF (4 mL), $NaN_3$ (0.590 g, 9.08 mmol, 2.00 eq.) was added and the reaction mixture was stirred overnight at room temperature, until LC-MS indicated complete conversion. DCM (10 mL) and $H_2O$ (3 mL) were added and the reaction mixture was allowed to stir for 2 h. Ethyl acetate (50 mL) was added and the mixture was washed with saturated aqueous NaCl (2 x 20 mL), dried over $MgSO_4$, filtered and concentrated under reduced pressure. The crude product was purified by flash column chromatography (0–50% ethyl acetate in *n*-heptane) to give **SI2** (1.04 g, 2.92 mmol, 64% over two steps) as colorless solid.

**$^1$H-NMR** ($CDCl_3$, 400 MHz): 7.38–7.31 (m, 6H, aryl-*H*), 7.31–7.35 (m, 4H, aryl-*H*), 5.13 (dd, 1H, *J* = 11.9 Hz, C$H_2$), 5.08 (s, 2H, C$H_2$), 5.02 (dd, 1H, *J* = 11.9 Hz, C$H_2$), 4.68 (d, 1H, *J* = 2.7 Hz, C*H*), 4.37 (d, 1H, *J* = 2.7 Hz, C*H*), 3.15 (s, br, 1H, O*H*); **$^{13}$C-NMR** ($CDCl_3$, 100 MHz): 170.7 (*C*O), 166.9 (*C*O), 134.6 (aryl-$C_q$), 134.4 (aryl-$C_q$), 129.0 (aryl-*C*), 128.9 (aryl-*C*), 128.9 (aryl-*C*), 128.8 (aryl-*C*), 128.8 (aryl-*C*), 72.2 (*C*H), 68.6 (*C*$H_2$), 68.3 (*C*$H_2$), 64.5 (*C*H); **HRMS (ESI)** m/z calcd. for $C_{18}H_{17}N_3O_5Na$: $(M+Na)^+$, 378.1060; found: 378.1067 $(M+Na)^+$; **$R_f$** (*n*-heptane/ethyl acetate 2:1): 0.37; **Chiral HPLC** (Chiralpak AS-H/122, 250x4.6 mm; EtOH:MeOH 1:1): **SI2** (Rt = 3.9 min) : ***ent*-SI2** (Rt = 4.3 min) 100 : 0 (> 99% *ee*); **Specific rotation** $[\alpha]_D^{20.6}$ = + 19.8 (c = 1.11; $CHCl_3$).

**Dibenzyl (2*S*,3*R*)-2-azido-3-hydroxysuccinate (SI2) and dibenzyl (2*R*,3*R*)-2-azido-3-hydroxysuccinate (SI4):**

**SI2** (0.300 g, 0.844 mmol, 1.00 eq.) was dissolved in anhydrous THF (10 mL), DBU (0.382 mL, 2.53 mmol, 3.00 eq.) was added and the reaction mixture was stirred at 40 °C. After 1 h the epimerization was stopped by addition of ethyl acetate (50 mL) and 1 M aqueous HCl (10 mL). The layers were separated, the organic layer was washed with saturated aqueous NaCl (10 mL), dried over $MgSO_4$, filtered and concentrated under reduced pressure. The crude product was purified *via* HPLC (RP-18, 15 min, 50-80% MeCN in $H_2O$/TFA 0.1%) to yield **SI2** (0.065 g, 0.18 mmol, 22%) as colorless solid and **SI4** (0.092 g, 0.26 mmol, 31%) as colorless oil.

**Dibenzyl (2*R*,3*R*)-2-azido-3-hydroxysuccinate (SI4)**

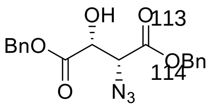

**SI4**

**$^1$H-NMR** ($CDCl_3$, 400 MHz): 7.42–7.32 (m, 10H, aryl-*H*), 5.28 (s, 2H, C$H_2$), 5.27 (d, 2H, $J$ = 2.1 Hz, C$H_2$), 4.81 (d, 1H, $J$ = 2.3 Hz, C*H*), 4.25 (d, 1H, $J$ = 2.3 Hz, C*H*), 3.20 (s, br, 1H, O*H*); **$^{13}$C-NMR** ($CDCl_3$, 100 MHz): 171.1 (*C*O), 167.4 (*C*O), 134.8 (aryl-$C_q$), 134.6 (aryl-$C_q$), 129.1 (aryl-*C*), 128.9 (aryl-*C*), 128.8 (aryl-*C*), 128.7 (aryl-*C*), 128.6 (aryl-*C*), 72.2 (*C*H), 68.6 (*C*$H_2$), 68.3 (*C*$H_2$), 63.5 (*C*H); **HRMS (ESI)** m/z calcd. for $C_{18}H_{17}N_3O_5Na$: $(M+Na)^+$, 378.1060; found: 378.1065 $(M+Na)^+$; **$R_f$** (*n*-heptane/ethyl acetate 2:1): 0.38; **Chiral HPLC** (Chiralpak AS-H/148, 250x4.6 mm; EtOH:MeOH 1:1): **SI4** (Rt = 6.8 min) : ***ent*-SI4** (Rt = 5.5 min) 99.3 : 0.7 (99% *ee*); **Specific rotation** $[\alpha]_D^{20.6}$ = + 59.9 (c = 0.87; $CHCl_3$).

**Dibenzyl (2*R*,3*S*)-2-azido-3-hydroxysuccinate (*ent*-SI2) and dibenzyl (2*S*,3*S*)-2-azido-3-hydroxysuccinate (*ent*-SI4):**

$SOCl_2$ (0.389 mL, 5.45 mmol, 1.20 eq.) was added to a solution of (−)-dibenzyl D-tartrate (***ent*-SI1**, 1.50 g, 4.54 mmol, 1.00 eq.) in anhydrous DCM (20 mL) at room temperature. A catalytic amount of anhydrous DMF (1 drop) was added and the reaction mixture was stirred for 7 h, until LC-MS indicated complete conversion. The reaction mixture was concentrated *in vacuo* and crude cyclic sulfite was used for the next step without further purification.

The crude product was dissolved in anhydrous DMF (4 mL), $NaN_3$ (0.590 g, 9.08 mmol, 2.00 eq.) was added and the reaction mixture was stirred overnight at room temperature, until LC-MS indicated complete conversion. DCM (10 mL) and $H_2O$ (3 mL) were added and the reaction mixture was allowed to stir for 2 h. Ethyl acetate (50 mL) was added and the mixture was washed with saturated aqueous NaCl (2 x 20 mL), dried over $MgSO_4$, filtered and concentrated under reduced pressure. The crude product (1.62 g, 4.54 mmol), quant.) was divided in two parts: 400 mg (1.13 mmol) were used for the epimerization (see below), the rest (1.22 g, 3.41 mmol) was purified *via* HPLC (RP-18, 15 min, 55-75% MeCN in $H_2O$/TFA 0.1%) to yield ***ent*-SI2** (0.470 g, 1.32 mmol, 39% over two steps) as colorless solid.

Crude ***ent*-SI2** (0.400 g, 1.13 mmol, 1.00 eq.) was dissolved in anhydrous THF (10 mL), DBU (0.509 mL, 3.38 mmol, 3.00 eq.) was added and the reaction mixture was stirred at 40 °C. After 1 h the epimerization was stopped by addition of ethyl acetate (50 mL) and 1 M aqueous HCl (10 mL). The layers were separated, the organic layer was washed with saturated aqueous NaCl

(10 mL), dried over $MgSO_4$, filtered and concentrated under reduced pressure. The crude product was purified *via* HPLC (RP-18, 15 min, 60-70% MeCN in $H_2O$/TFA 0.1%) to yield ***ent*-SI2** (0.077 g, 0.22 mmol, 19% over three steps) as colorless solid and ***ent*-SI4** (0.115 g, 0.324 mmol, 29% over three steps) as colorless oil.

**Dibenzyl (2*R*,3*S*)-2-azido-3-hydroxysuccinate (*ent*-SI2)**

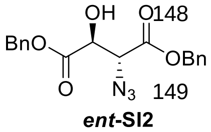

***ent*-SI2**

The NMR data are identical to the ones reported for **SI2**.

**HRMS (ESI)** m/z calcd. for $C_{18}H_{17}N_3O_5Na$: $(M+Na)^+$, 378.1060; found: 378.1066 $(M+Na)^+$; **$R_f$** (*n*-heptane/ethyl acetate 2:1): 0.37; **Chiral HPLC** (Chiralpak AS-H/122, 250x4.6 mm; EtOH:MeOH 1:1): ***ent*-SI2** (Rt = 4.3 min) : **SI2** (Rt = 3.9 min): 99.5 : 0.50 (99% *ee*); **Specific rotation** $[\alpha]_D^{20.6}$ = – 19.3 (c = 1.34; $CHCl_3$).

**Dibenzyl (2*S*,3*S*)-2-azido-3-hydroxysuccinate (*ent*-SI4)**

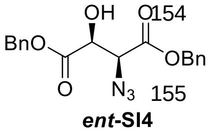

***ent*-SI4**

The NMR data are identical to the ones reported for **SI4**.

**HRMS (ESI)** m/z calcd. for $C_{18}H_{17}N_3O_5Na$: $(M+Na)^+$, 378.1060; found: 378.1065 $(M+Na)^+$; **$R_f$** (*n*-heptane/ethyl acetate 2:1): 0.38; **Chiral HPLC** (Chiralpak AS-H/148, 250x4.6 mm; EtOH:MeOH 1:1): ***ent*-SI4** (Rt = 5.5 min) : **SI4** (Rt = 6.8 min) 99.7 : 0.3 (99% *ee*); **Specific rotation** $[\alpha]_D^{20.6}$ = – 67.9 (c = 0.68; $CHCl_3$).

**General procedure for hydrogenation:**

The acid **SI2**, ***ent*-SI2**, **SI4** or ***ent*-SI4** (0.1–0.2 mmol) was dissolved in 1,4-dioxane (5 mL) and 1 M aqueous HCl (1 mL), Pd/C (10 %, 0.05 eq.) was added and the reaction mixture was hydrogenated at 4 bar pressure in an autoclave at room temperature. After reaction overnight, LC-MS indicated complete hydrogenation to **SI3**, ***ent*-SI3**, **SI5** or ***ent*-SI5**, respectively. The reaction mixture was filtered, diluted with $H_2O$ (20 mL) and freeze-dried to yield the desired product (HCl salt, quant.) as colourless solid.

**(2*S*,3*R*)-2-Amino-3-hydroxysuccinic acid ((2*S*,3*R*)-3-hydroxyaspartic acid) (SI3):**

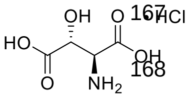

**SI3**

**$^1$H-NMR** ($D_2O$, 400 MHz): 4.73 (d, 1H, $J$ = 2.7 Hz, C*H*), 4.59 (d, 1H, $J$ = 2.7 Hz, C*H*); **$^{13}$C-NMR** ($D_2O$, 100 MHz): 176.1 (*C*OOH), 71.7 (*C*H), 58.5 (*C*H); **HRMS (ESI)** m/z calcd. for $C_4H_8NO_5$: $(M+H)^+$, 150.0397; found: 150.0397 $(M+H)^+$; **Specific rotation** $[\alpha]_D^{20.8}$ = + 46.6 (c = 1.03; $H_2O$).

**(2*R*,3*S*)-2-Amino-3-hydroxysuccinic acid ((2*R*,3*S*)-3-hydroxyaspartic acid) (*ent*-SI3):**

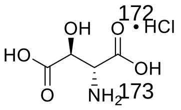

***ent*-SI3**

The NMR data are identical to the ones reported for **SI3**.

**HRMS (ESI)** m/z calcd. for $C_4H_8NO_5$: $(M+H)^+$, 150.0397; found: 150.0397 $(M+H)^+$; **Specific rotation** $[\alpha]_D^{20.7}$ = – 44.3 (c = 0.65; $H_2O$).

**(2*R*,3*R*)-2-Amino-3-hydroxysuccinic acid ((2*R*,3*R*)-3-hydroxyaspartic acid) (SI5):**

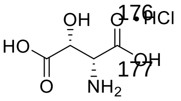

SI5

**$^1$H-NMR** ($D_2O$, 400 MHz): 4.86 (d, 1H, $J$ = 2.4 Hz, C*H*), 4.40 (d, 1H, $J$ = 2.4 Hz, C*H*); **$^{13}$C-NMR** ($D_2O$, 100 MHz): 71.5 (*C*H), 58.4 (*C*H); **HRMS (ESI)** m/z calcd. for $C_4H_8NO_5$: (M+H)$^+$, 150.0397; found: 150.0398 (M+H)$^+$; **Specific rotation** $[\alpha]_D^{20.8}$ = + 9.7 (c = 0.51; $H_2O$).

**(2*S*,3*S*)-2-Amino-3-hydroxysuccinic acid ((2*S*,3*S*)-3-hydroxyaspartic acid) (*ent*-SI5):**

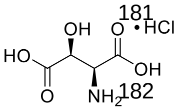

*ent*-SI5

The NMR data are identical to the ones reported for **SI5**.

**HRMS (ESI)** m/z calcd. for $C_4H_8NO_5$: (M+H)$^+$, 150.0397; found: 150.0397 (M+H)$^+$; **Specific rotation** $[\alpha]_D^{20.6}$ = − 8.0 (c = 0.63; $H_2O$).

#### 1.3. Preparation of the β-Hydroxyphenylalanines

The L-isomers of the β-hydroxyphenylalanines and β-hydroxyisoleucines were synthesized by diastereoselective addition of Grignard reagents to orthoester protected L-serine aldehyde **SI7**, initially described by Blaskovich and Lajoie[2]. This synthetic pathway offers several advantages for our envisaged access to reference samples of these β-hydroxyamino acids:

- High *threo*-selectivities are reported for the addition of Grignard reagents to aldehyde **SI7** and ketones **SI18** and **SI25** (Scheme S2 and Scheme S3).
- The addition products are bearing the oxidation stage of the desired amino acids and only protecting group manipulations are required to obtain the reference samples. In contrast to the commonly used synthetic strategy for β-hydroxyamino acids starting from Garner's Aldehyde[3,4], no protection of the secondary alcohol and no further oxidation step are required.
- The Cbz-group allows for accurate *ee*-determination by chiral HPLC (due to their UV absorption at a detection wavelength of 207 nm) and convenient deprotection *via* catalytic hydrogenation or under acidic conditions at the last step.
- The epimerization of aldehyde **SI7** *via* simple flash chromatography on silica exhibits a practical access to racemic reference samples which are required for the *ee*-determination by chiral HPLC and to cover the corresponding *R*-series of the β-hydroxyamino acids.

[2] Blaskovich, M. A. *et al.* Stereoselective Synthesis of Threo and Erythro β-Hydroxy and β-Disubstituted-β-Hydroxy α-Amino Acids. *J. Org. Chem.* **63,** 3631–3646 (1998).
[3] Garner, P. Stereocontrolled addition to a penaldic acid equivalent: an asymmetric of threo-β-hydroxy-L-glutamic acid. *Tetrahedron Lett.* **25,** 5855–5858 (1984).
[4] Passiniemi, M. & Koskinen A. M. P. Garner's aldehyde as a versatile intermediate in the synthesis of enantiopure natural products. *Beilstein J. Org. Chem.* **9,** 2641–2659 (2013).

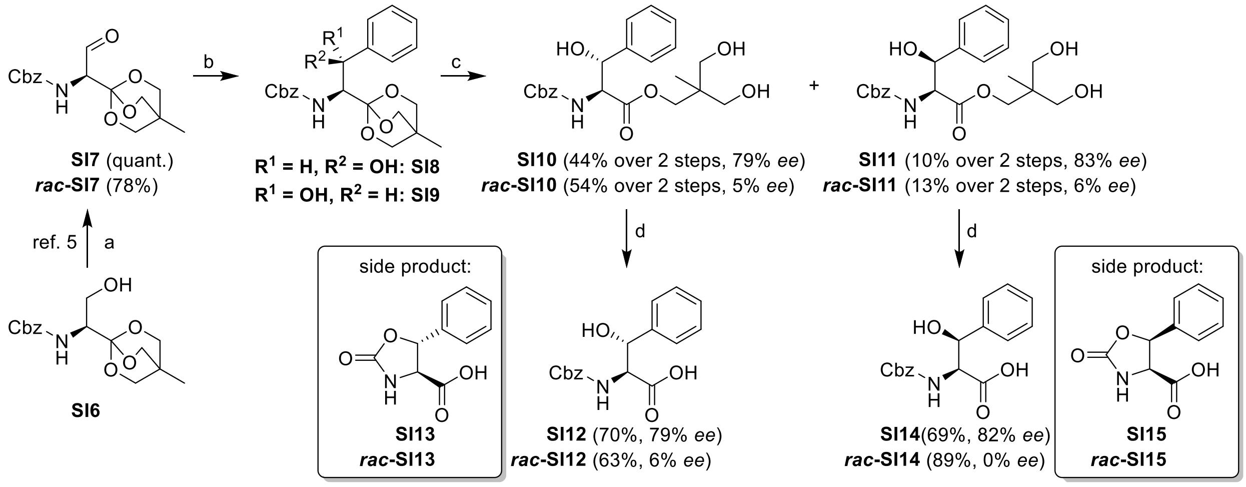

***Scheme S2.*** **Preparation of β-Hydroxyphenylalanines.**[a]

[a] Conditions: a) $(COCl)_2$, DMSO, DCM, −78 °C, 15 min; **SI6**, −78 °C, 90 min; DIPEA, −78 °C to 0 °C, 30 min; **SI7** used as a crude; ***rac*-SI7** purified by flash column chromatography; b) PhMgBr, THF, rt; c) 1 M $HCl_{aq}$, 1,4-dioxane; d) LiOH, MeCN/$H_2O$ 4:1.

The synthesis of **SI7** and ***rac*-SI7** was performed in close analogy to a literature known procedure from **SI6**[5]. To obtain both diastereomers in one reaction and in sufficient quantities, the addition of PhMgBr was performed at room temperature (Scheme S2). Under these conditions, a selectivity of 84:16 for **SI8**:**SI9** is described[2]. In our hands, complete separation of **SI8** and **SI9** *via* flash chromatography was difficult on 1 g-scale. Therefore, the mixture of **SI8** and **SI9** was directly treated by aqueous HCl to achieve orthoester hydrolysis and resulting products **SI10** and **SI11** were separated by preparative HPLC. Previous orthoester hydrolysis is required since the OBO-orthoesters are unstable to acidic HPLC conditions. Contrary to Ref. 2, we performed a saponification of **SI10** and **SI11** to Cbz-protected amino acids **SI12** and **SI14** to obtain UV-active compounds for *ee*-determination. Careful monitoring of the base equivalents and the reaction progress is required since an excess of LiOH resulted in formation of cyclic carbamates **SI13** and **SI15**, respectively.[6] However, we were able to suppress this side reaction to an acceptable level. For the Marfey-protocol, both the ester **SI10** and **SI11** and their corresponding acids **SI12** and **SI14** were suitable because the acidic conditions cleave the Cbz-proteting group and the ester. In our synthesis, we achieved acceptable *ee*s for the *S*-series (79-

[5] Rose, N. G. W. *et al.* Preparation of 1-[N-Benzyloxycarbonyl-(1S)-1-amino-2-oxoethyl]-4-methyl-2,6,7-trioxabicyclo-[2.2.2]octane. *Organic Syntheses, Coll. Vol. 10,* 73 (2004); *Vol. 79,* 216 (2002).

[6] The carbamates were only isolated for the racemic series for characterization.

83%) and the epimerization of **SI7** *via* flash chromatography was successful (*ee*s from 0-6% for the *rac*-series).

**Benzyl (*S*)-(1-(4-methyl-2,6,7-trioxabicyclo[2.2.2]octan-1-yl)-2-oxoethyl)carbamate (SI7):**

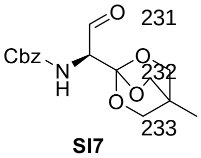

**SI7**

**SI7** was synthesized according to a slightly modified literature-known procedure[5]: The reaction was carried out in moisture-free glassware under inert atmosphere. To a solution of $(COCl)_2$ (0.441 mL, 5.11 mmol, 1.70 eq.) in anhydrous DCM (20 mL) anhydrous DMSO (0.683 mL, 9.62 mmol, 3.20 eq.) was added carefully at −78 °C. After stirring for 15 min, alcohol **SI6** (0.972 g, 3.01 mmol, 1.00 eq.) dissolved in anhydrous DCM (20 mL) was added. The reaction mixture was stirred for 1.5 h at −78 °C. Then DIPEA (3.30 mL, 18.7 mmol, 6.23 eq.) was added and the solution was stirred for 30 min at −78 °C and for further 30 min without cooling bath. A mixture of cooled toluene/saturated aqueous $NH_4Cl$ (4:1, 250 mL, pre-cooled to 0 °C) was added to the reaction mixture. The layers were separated and the organic layer was washed with saturated aqueous $NH_4Cl$ (3 x 50 mL, pre-cooled to 0 °C), with saturated aqueous $NaHCO_3$ (50 mL, pre-cooled to 0 °C), and with saturated aqueous NaCl (50 mL, pre-cooled to 0 °C). The organic layer was dried over $MgSO_4$, filtered, and concentrated under reduced pressure to yield crude aldehyde **SI7** (1.00 g, max. 3.01 mmol, quant.) as slightly yellow oil which was used in the next stage without further purification.

***Rac*-SI7** was synthesized starting from **SI6** in a similar manner with an additional purification step via chromatography, which led to full epimerization of the stereocenter[7]: The reaction was carried out in moisture-free glassware under inert atmosphere. To a solution of $(COCl)_2$ (1.22 mL, 14.0 mmol, 1.70 eq.) in anhydrous DCM (60 mL) anhydrous DMSO (2.09 mL, 30.0 mmol, 3.60 eq.) was added carefully at −78 °C. After stirring for 15 min, alcohol **SI6** (2.65 g, 8.21 mmol, 1.00 eq.) dissolved in anhydrous DCM (60 mL) was added. The reaction mixture was stirred for 1.5 h at −78 °C. Then DIPEA (9.00 mL, 51.2 mmol, 6.23 eq.) was added and the solution was stirred for 30 min at −78 °C and for further 30 min without cooling bath. Saturated aqueous $NH_4Cl$ (100 mL) and DCM (100 mL) were added to the reaction mixture, the layers were separated and the organic layer was washed with saturated aqueous $NH_4Cl$ (50 mL), with saturated aqueous $NaHCO_3$ (50 mL), and with saturated aqueous NaCl (50 mL). The organic

[7] The epimerization of the stereocenter of OBO-protected serinals such as **SI7** *via* chromatography is a known phenomenon and was described by Blascovich and Lajoie (Blaskovich, M. A. & Lajoie, G. A. Synthesis of a chiral serine aldehyde equivalent and its conversion to chiral .alpha.-amino acid derivatives. *J. Am. Chem. Soc.* **115,** 5021–5030 (1993).) In our case, the purification *via* column chromatography resulted in measured *ee*-values of 0–6% for the *rac*-series.

layer was dried over $MgSO_4$, filtered, and concentrated under reduced pressure. The crude product was purified by flash column chromatography[8] (100% DCM, then DCM : EtOAc 4:1) to give ***rac*-SI7** (2.06 g, 6.40 mmol, 78 %) as a colorless solid and was directly used for the next step.

$\mathbf{R_f}$ (*n*-heptane/ethyl acetate 1:3): 0.65

[8] Conditioning of the silica column was performed with a mixture of DCM and $NEt_3$ (98:2) to neutralize the slightly acidic silica and to prevent partial orthoester hydrolysis during chromatography.

**3-Hydroxy-2-(hydroxymethyl)-2-methylpropyl (2*S*,3*R*)-2-(((benzyloxy)carbonyl)amino)-3-hydroxy-3-phenylpro-panoate (SI10) and 3-Hydroxy-2-(hydroxymethyl)-2-methylpropyl (2*S*,3*S*)-2-(((benzyloxy)carbonyl)amino)-3-hydroxy-3-phenylpro-panoate (SI11):**

The reaction was carried out in moisture-free glassware under inert atmosphere. To a solution of aldehyde **SI7** (0.965 g, 3.00 mmol, 1.00 eq.) in anhydrous THF (30 mL) was added PhMgBr (1 M in THF, 12.0 mL, 12.0 mmol, 4.00 eq.) at room temperature. After 90 min, TLC indicated full conversion of the starting material ($R_f$ of **SI8**/**SI9** (*n*-heptane/ethyl acetate 1:3): 0.60). Saturated aqueous $NH_4Cl$ (20 mL) and EtOAc (100 mL) were added to the reaction mixture, the layers were separated and the organic layer was washed with saturated aqueous NaCl (20 mL). The organic layer was dried over $MgSO_4$, filtered, and concentrated under reduced pressure. The crude mixture of **SI8**/**SI9** was dissolved in 1,4-dioxane (20 mL) and aqueous HCl (1 N, 1 mL) and the reaction mixture was stirred for 30 min until full hydrolysis of the orthoester was observed by TLC. All volatiles were removed under reduced pressure and the residue was purified *via* HPLC (RP-18, 15 min, 35-45% MeCN in $H_2O$/TFA 0.1%) to yield **SI10** (0.549 g, 1.32 mmol, 44% over two steps) and **SI11** (0.124 g, 0.297 mmol, 10% over two steps) as colorless oils.

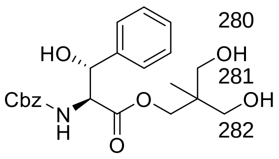

SI10

For **SI10**: **$^1$H-NMR** (DMSO-d$_6$, 400 MHz): 7.42–7.19 (m, 11H, N*H*, aryl-*H*), 5.69 (d, 1H, *J* = 6.2 Hz, CH-O*H*), 5.11 (dd, 1H, *J* = 6.1, 3.9 Hz, C*H*-OH), 4.95 (s, 2H, Ph-C$H_2$), 4.47 (m, 2H, $CH_2$-O*H*), 4.40 (dd, 1H, *J* = 9.3, 3.7 Hz, C*H*-NH), 3.96 (d, 1H, *J* = 10.5 Hz, C$H_2$), 3.90 (d, 1H, *J* = 10.5 Hz, C$H_2$), 3.30–3.21 (m, 4H, C$H_2$-OH), 0.77 (s, 3H, C$H_3$); **$^{13}$C-NMR** (DMSO-d$_6$, 100 MHz): 170.4 (*C*O), 156.2 (*C*O-NH), 141.6 (aryl-$C_q$), 136.9 (aryl-$C_q$), 128.3 (aryl-*C*), 127.9 (aryl-*C*), 127.7 (aryl-*C*), 127.3 (aryl-*C*), 127.2 (aryl-*C*), 126.1 (aryl-*C*), 72.3 (*C*H-OH), 67.1 (*C*$H_2$), 65.4 (Ph-*C*$H_2$), 63.5 (*C*$H_2$-OH), 60.5 (*C*H-NH), 40.7 ($C_q$-$CH_3$), 16.3 (*C*$H_3$); **HRMS (ESI)** m/z calcd. for $C_{22}H_{27}NO_7Na$: $(M+Na)^+$, 440.1680; found: 440.1685 $(M+Na)^+$; **$R_f$** (*n*-heptane/ethyl acetate 1:3): 0.25; **Chiral HPLC** (Chiralpak ID/174, 250x4.6 mm; *n*-heptane:EtOH:MeOH 5:1:1 + 0.1% TFA): **SI10** (Rt = 8.6 min) : ***ent*-SI10** (Rt = 5.6 min) 89.3 : 10.7 (79% *ee*); **Specific rotation** $[\alpha]_D^{20.0} = -35.1$ (c = 0.29; MeCN).

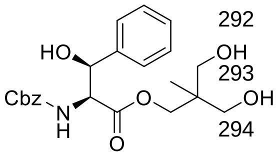

SI11

For **SI11**: **$^1$H-NMR** (DMSO-d$_6$, 400 MHz): 7.64 (d, 1H, *J* = 9.2 Hz, N*H*), 7.42–7.37 (m, 2H, aryl-*H*), 7.35–7.23 (m, 6H, aryl-*H*), 7.21–7.10 (m, 2H, aryl-*H*), 5.76 (d, 1H, *J* = 4.7 Hz, CH-O*H*), 4.93 (d, 1H, *J* = 13.4 Hz, Ph-C$H_2$), 4.90 (d, 1H, *J* = 13.4 Hz, Ph-C$H_2$), 4.76 (dd, 1H, *J*

= 8.3, 4.2 Hz, C*H*-OH), 4.42 (s, br, 2H, $CH_2$-O*H*), 4.24 (t, 1H, *J* = 8.9 Hz, C*H*-NH), 3.96 (d, 1H, *J* = 10.6 Hz, C$H_2$), 3.91 (d, 1H, *J* = 10.6 Hz, C$H_2$), 3.26 (s, 4H, C$H_2$-OH), 0.75 (s, 3H, C$H_3$); **$^{13}$C-NMR** (DMSO-$d_6$, 100 MHz): 170.9 (*C*O), 155.5 (*C*O-NH), 141.9 (aryl-$C_q$), 136.8 (aryl-$C_q$), 128.3 (aryl-*C*), 127.8 (aryl-*C*), 127.7 (aryl-*C*), 127.4 (aryl-*C*), 127.4 (aryl-*C*), 126.8 (aryl-*C*), 72.3 (*C*H-OH), 66.7 (*C*$H_2$), 65.3 (Ph-*C*$H_2$), 63.5 (*C*$H_2$-OH), 60.5 (*C*H-NH), 40.7 ($C_q$-$CH_3$), 16.3 (*C*$H_3$); **HRMS (ESI)** m/z calcd. for $C_{22}H_{27}NO_7Na$: $(M+Na)^+$, 440.1680; found: 440.1685 $(M+Na)^+$; **$R_f$** (*n*-heptane/ethyl acetate 1:3): 0.25; **Chiral HPLC** (Chiralpak IF/181, 250x4.6 mm; *n*-heptane:EtOH:MeOH 5:1:1 + 0.1% TFA): **SI11** (Rt = 13.7 min) : ***ent*-S11** (Rt = 8.5 min) 91.6 : 8.4 (83% *ee*); **Specific rotation** $[\alpha]_D^{19.9}$ = + 6.2 (c = 0.32; MeCN).

***Rac*-SI10** (0.482 g, 1.16 mmol, 54% over two steps) and ***rac*-SI11** (0.118 g, 0.283 mmol, 13% over two steps) were synthesized from ***rac*-SI7** (0.685 g, 2.13 mmol, 1.00 eq.) in analogous manner. The NMR data are identical to the ones reported for **SI10** and **SI11**.

For ***rac*-SI10**: **$R_f$** (*n*-heptane/ethyl acetate 1:3): 0.25; **Chiral HPLC** (Chiralpak ID/174, 250x4.6 mm; *n*-heptane:EtOH:MeOH 5:1:1 + 0.1% TFA): **SI10** (Rt = 8.6 min) : ***ent*-SI10** (Rt = 5.6 min) 52.3 : 47.7 (5% *ee*).

For ***rac*-SI11**: **$R_f$** (*n*-heptane/ethyl acetate 1:3): 0.25; **Chiral HPLC** (Chiralpak IF/181, 250x4.6 mm; *n*-heptane:EtOH:MeOH 5:1:1 + 0.1% TFA): **SI11** (Rt = 14.0 min) : ***ent*-S11** (Rt = 8.5 min) 53.2 : 46.8 (6% *ee*).

**(2*S*,3*R*)-2-(((Benzyloxy)carbonyl)amino)-3-hydroxy-3-phenylpropanoic acid (SI12):**

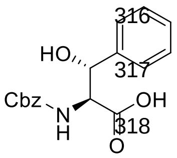

SI12

To a solution of **SI10** (0.509 g, 1.22 mmol, 1.00 eq.) in MeCN/$H_2O$ (4:1, 20.4 mL/5.1 mL) was added LiOH (1 N in $H_2O$, 1.22 mL, 1.22 mmol, 1.00 eq.) at room temperature. After 20 min, another portion of LiOH (1 N in $H_2O$, 0.61 mL, 0.61 mmol, 0.50 eq.) was added to reach full conversion of starting material after 30 min monitored by LC-MS. The reaction was stopped by addition of acetic acid (0.1 mL). All volatiles were removed under reduced pressure and the residue was purified *via* HPLC (RP-18, 15 min, 10-55% MeCN in $H_2O$/TFA 0.1%) to yield **SI12** (0.270 g, 0.856 mmol, 70%) as a colorless solid.[9]

**$^1$H-NMR** (DMSO-d$_6$, 400 MHz): 12.81 (s, br, 1H, COO*H*), 7.45–7.19 (m, 10H, aryl-*H*), 7.08 (d, 1H, *J* = 9.5 Hz, N*H*), 5.68 (s, br, 1H, O*H*), 5.16 (s, br., 1H, C*H*-OH), 4.96 (d, 1H, *J* = 13.5 Hz, Ph-C*H*$_2$), 4.93 (d, 1H, *J* = 13.5 Hz, Ph-C*H*$_2$), 4.33 (dd, 1H, *J* = 9.5, 3.4 Hz, C*H*-NH); **$^{13}$C-NMR** (DMSO-d$_6$, 100 MHz): 171.9 (*C*OOH), 156.2 (*C*O-NH), 142.1 (aryl-*C*$_q$), 137.0 (aryl-*C*$_q$), 128.3 (aryl-*C*), 127.8 (aryl-*C*), 127.7 (aryl-*C*), 127.3 (aryl-*C*), 127.1 (aryl-*C*), 126.2 (aryl-*C*), 72.3 (*C*H-OH), 65.3 (Ph-*C*H$_2$), 60.4 (*C*H-NH); **HRMS (ESI)** m/z calcd. for $C_{17}H_{17}NO_5Na$: $(M+Na)^+$, 338.0999; found: 338.1000 $(M+Na)^+$; **Chiral HPLC** (Chiralpak IF/181, 250x4.6 mm; *n*-heptane:EtOH:MeOH 5:1:1 + 0.1% TFA): **SI12** (Rt = 6.8 min) : ***ent*-SI12** (Rt = 5.5 min) 89.7 : 10.3 (79% *ee*); **Specific rotation** $[\alpha]_D^{19.9}$ = – 23.0 (c = 0.65; MeCN).

***Rac*-SI12** (0.210 g, 0.666 mmol, 63%) and ***rac*-SI13** (0.055 g, 0.27 mmol, 25%) were synthesized from ***rac*-SI10** (0.442 g, 1.06 mmol, 1.00 eq.) in analogous manner. The NMR data of ***rac*-SI12** are identical to the ones reported for **SI12**. ***Rac*-SI12** and ***rac*-SI13** were obtained as colorless solids.

For ***rac*-SI12**: **Chiral HPLC** (Chiralpak IF/181, 250x4.6 mm; *n*-heptane:EtOH:MeOH 5:1:1 + 0.1% TFA): **SI12** (Rt = 6.8 min) : ***ent*-SI12** (Rt = 5.5 min) 52.8 : 47.2 (6% *ee*).

**(4*S*,5*R*)-2-Oxo-5-phenyloxazolidine-4-carboxylic acid (*rac*-SI13):**

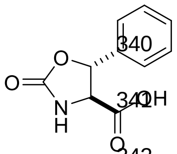

*rac*-SI13

**$^1$H-NMR** (DMSO-d$_6$, 500 MHz): 13.45 (s, br., 1H, COO*H*), 8.36 (s, br., 1H, N*H*), 7.53–7.26 (m, 5H, aryl-*H*), 5.57 (d, 1H, *J* = 4.9 Hz, O-C*H*), 4.23 (dd, 1H, *J* = 4.9, 0.7 Hz, NH-C*H*); **$^{13}$C-NMR** (DMSO-d$_6$, 125 MHz): 171.8 (*C*OOH), 157.7 (*C*O), 139.0 (aryl-*C*$_q$), 128.8 (aryl-*C*), 128.8 (aryl-*C*), 125.8 (aryl-*C*), 78.4 (O-*C*H), 60.6

[9] The formation of **SI13** was observed *via* LC-MS, but was not isolated for the *S*-series.

(NH-*C*H); **HRMS (ESI)** m/z calcd. for $C_{10}H_9NO_4Na$: $(M+Na)^+$, 230.0424; found: 230.0425 $(M+Na)^+$.

**(2*S*,3*S*)-2-(((Benzyloxy)carbonyl)amino)-3-hydroxy-3-phenylpropanoic acid (SI14):**

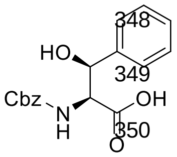

**SI14**

To a solution of **SI11** (0.083 g, 0.20 mmol, 1.00 eq.) in MeCN/$H_2O$ (4:1, 3.2 mL/0.8 mL) was added LiOH (1 N in $H_2O$, 0.20 mL, 0.20 mmol, 1.00 eq.) at room temperature. After 20 min, another portion of LiOH (1 N in $H_2O$, 0.10 mL, 0.10 mmol, 0.50 eq.) was added to reach full conversion of starting material after 30 min monitored by LC-MS. The reaction was stopped by addition of acetic acid (0.1 mL). All volatiles were removed under reduced pressure and the residue was purified *via* HPLC (RP-18, 15 min, 10-55% MeCN in $H_2O$/TFA 0.1%) to yield **SI14** (0.043 g, 0.14 mmol, 69%) as a colorless solid.[10]

**$^{1}$H-NMR** (DMSO-d$_6$, 400 MHz): ~12.51 (s, v. br, 1H, COO*H*), 7.48 (d, 1H, *J* = 9.3 Hz, N*H*), 7.42–7.36 (m, 2H, aryl-*H*), 7.35–7.23 (m, 6H, aryl-*H*), 7.21–7.14 (m, 2H, aryl-*H*), ~5.74 (s, v. br, 1H, O*H*), 4.93 (d, 1H, *J* = 13.3 Hz, Ph-C*H*$_2$), 4.90 (d, 1H, *J* = 13.4 Hz, Ph-C*H*$_2$), 4.73 (d. 1H, *J* = 8.5 Hz, C*H*-OH), 4.15 (t, 1H, *J* = 8.8 Hz, C*H*-NH); **$^{13}$C-NMR** (DMSO-d$_6$, 100 MHz): 172.3 (*C*OOH), 155.5 (*C*O-NH), 142.2 (aryl-*C*$_q$), 136.9 (aryl-*C*$_q$), 128.3 (aryl-*C*), 127.7 (aryl-*C*), 127.7 (aryl-*C*), 127.4 (aryl-*C*), 127.3 (aryl-*C*), 126.9 (aryl-*C*), 72.4 (*C*H-OH), 65.2 (Ph-*C*H$_2$), 60.4 (*C*H-NH); **HRMS (ESI)** m/z calcd. for $C_{17}H_{17}NO_5Na$: $(M+Na)^+$, 338.0999; found: 338.0998 $(M+Na)^+$; **Chiral HPLC** (Chiralpak IF/181, 250x4.6 mm; *n*-heptane:EtOH:MeOH 5:1:1 + 0.1% TFA): **SI14** (Rt = 5.2 min) : ***ent*-SI14** (Rt = 6.2 min) 90.9 : 9.1 (82% *ee*); **Specific rotation** $[\alpha]_D^{20.0}$ = + 32.3 (c = 0.31; MeCN).

***Rac*-SI14** (0.053 g, 0.17 mmol, 89%) and ***rac*-SI15** (0.002 g, 0.01 mmol, 5%) were synthesized from ***rac*-SI11** (0.079 g, 0.19 mmol, 1.00 eq.) in analogous manner. The NMR data of ***rac*-SI14** are identical to the ones reported for **SI14**. ***Rac*-SI14** and ***rac*-SI15** were obtained as colorless solids.

For ***rac*-SI14**: **Chiral HPLC** (Chiralpak IF/181, 250x4.6 mm; *n*-heptane:EtOH:MeOH 5:1:1 + 0.1% TFA): **SI14** (Rt = 5.2 min) : ***ent*-SI14** (Rt = 6.2 min) 50.2 : 49.8 (0% *ee*).

**(4*S*,5*S*)-2-Oxo-5-phenyloxazolidine-4-carboxylic acid (*rac*-SI15):**

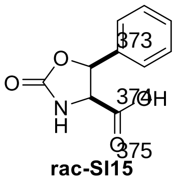

**rac-SI15**

**$^{1}$H-NMR** (DMSO-d$_6$, 500 MHz): 12.71 (s, br., 1H, COO*H*), 8.15 (s, 1H, N*H*), 7.43–7.33 (m, 5H, aryl-*H*), 5.85 (d, 1H, *J* = 8.9 Hz, O-C*H*), 4.59 (d, 1H, *J* = 8.9 Hz, NH-C*H*); **$^{13}$C-NMR** (DMSO-d$_6$, 125 MHz): 170.5 (*C*OOH), 158.4 (*C*O), 135.2 (aryl-*C*$_q$), 128.6 (aryl-*C*), 128.1 (aryl-*C*), 126.4 (aryl-*C*), 77.9 (O-

[10] The formation of **SI15** was observed *via* LC-MS, but was not isolated for the *S*-series.

*C*H), 59.3 (NH-*C*H); **HRMS (ESI)** m/z calcd. for $C_{10}H_{10}NO_4$: $(M+H)^+$, 208.0604; found: 208.0604 $(M+H)^+$.

#### 1.4. Preparation of the β-Hydroxyisoleucines

The synthesis of the β-hydroxyisoleucines was performed in close analogy to the synthesis of the β-hydroxyphenylalanines (see chapter 1.3). Addition of MeMgBr and EtMgBr, respectively, to aldehyde **SI7** was performed at room temperature and the crude mixture of the diastereomers **SI16**/**SI17** and **SI23**/**SI24** was subjected to Swern oxidation to yield crude ketones **SI19** and **SI26** (Scheme S3). High diastereoselectivities > 10:1 were achieved at the addition of MeMgBr and EtMgBr, respectively, to ketones **SI19** and **SI26**. The reactions, however, turned out to be slow and therefore were performed at room temperature. For the racemic series, an additional equivalent of Grignard reagent was added after 90 min, which led to a significant improvement of the isolated yields compared to the previously performed *S*-series. Saponifications of **SI20** and **SI27** were accompanied with faster formation of the cyclic carbamates **SI22** and **SI29** compared to the β-hydroxyphenylalanine series (see chapter 1.3) but sufficient amounts of target products **SI21** and **SI28** were obtained. Noteworthy, the formation of cyclic carbamate **SI22** is significantly quicker than **SI29**, which resulted in different diastereomeric ratios of the isolated products **SI21** and **SI28**. Neither at the stage of the ester **SI20** and **SI27** nor at the stage of the free carboxylic acids **SI21** and **SI28**, separation of the diastereomers was possible. However, the samples were sufficiently diastereomer-enriched to be used for Marfey's protocol. The ratio of the diastereomers was distinguished by 1H-NMR and chiral HPLC.

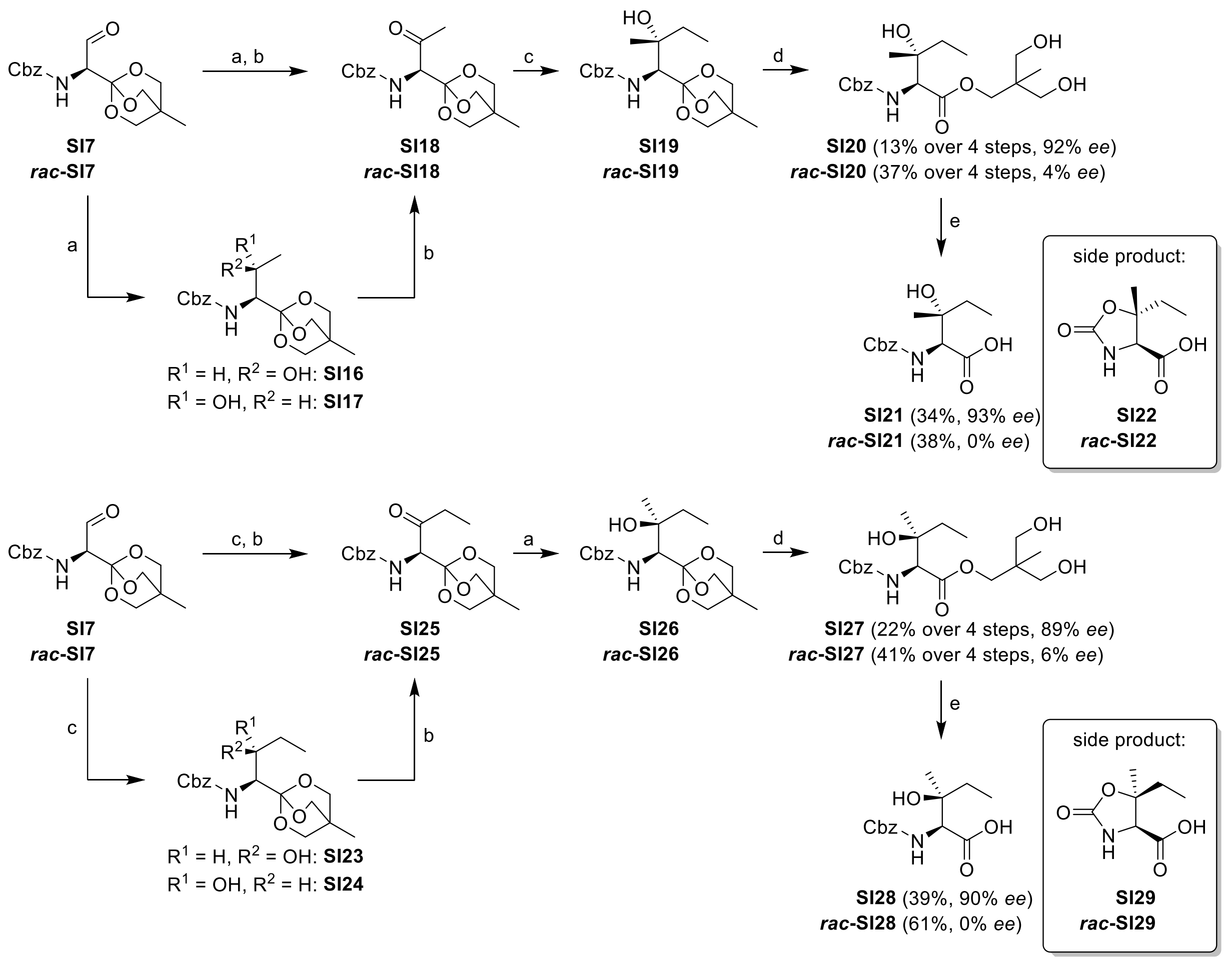

***Scheme S3.* Preparation of β-Hydroxyisoleucines.**[a]

[a] Conditions: a) MeMgBr, THF, rt; b) $(COCl)_2$, DMSO, DCM, −78 °C, 15 min; **SI16/SI17** or **SI23/SI24** or the corresponding racemic mixtures, −78 °C, 90 min; DIPEA, −78 °C to 0 °C, 30 min; **SI18**, ***rac*-SI18**, **SI25** or ***rac*-SI25** used as a crude; c) EtMgBr, THF, rt; d) 1 M $HCl_{aq}$, 1,4-dioxane; e) LiOH, $MeCN/H_2O$ 4:1.

**3-Hydroxy-2-(hydroxymethyl)-2-methylpropyl (2*S*,3*R*)-2-(((benzyloxy)carbonyl)amino)-3-hydroxy-3-methylpenta-oate (SI20):**

The reaction was carried out in moisture-free glassware under inert atmosphere. To a solution of aldehyde **SI7** (0.830 g, 2.58 mmol, 1.00 eq.) in anhydrous THF (30 mL) was added MeMgBr (3 M in $Et_2O$, 3.44 mL, 10.3 mmol, 4.00 eq.) at room temperature. After 90 min, TLC indicated full conversion of the starting material ($R_f$ of **SI16**/**SI17** (*n*-heptane/ethyl acetate 1:3): 0.48). Saturated aqueous $NH_4Cl$ (20 mL) and EtOAc (100 mL) were added to the reaction mixture, the layers were separated and the organic layer was washed with saturated aqueous NaCl (20 mL). The organic layer was dried over $MgSO_4$, filtered, and concentrated under reduced pressure. The crude mixture of **SI16**/**SI17** (0.801 g, 2.37 mmol, 92%) was used for the oxidation without further purification.

The reaction was carried out in moisture-free glassware under inert atmosphere. To a solution of $(COCl)_2$ (0.307 mL, 3.56 mmol, 1.50 eq.) in anhydrous DCM (20 mL) anhydrous DMSO (0.533 mL, 7.50 mmol, 3.16 eq.) was added carefully at −78 °C. After stirring for 15 min, the crude mixture of **SI16**/**SI17** (0.801 g, 2.37 mmol, 1.00 eq.) dissolved in anhydrous DCM (20 mL) was added. The reaction mixture was stirred for 1.5 h at −78 °C. Then DIPEA (2.30 mL, 13.1 mmol, 5.50 eq.) was added and the solution was stirred for 30 min at −78 °C and for further 30 min without cooling bath. A mixture of cooled toluene/saturated aqueous $NH_4Cl$ (4:1, 250 mL, pre-cooled to 0 °C) was added to the reaction mixture. The layers were separated and the organic layer was washed with saturated aqueous $NH_4Cl$ (3 x 50 mL, pre-cooled to 0 °C), with saturated aqueous $NaHCO_3$ (50 mL, pre-cooled to 0 °C), and with saturated aqueous NaCl (50 mL, pre-cooled to 0 °C). The organic layer was dried over $MgSO_4$, filtered, and concentrated under reduced pressure to yield crude ketone **SI18** (0.854 g, max. 2.37 mmol, quant., $R_f$ (*n*-heptane/ethyl acetate 1:3): 0.64) as slightly yellow oil which was used in the next stage without further purification.

The reaction was carried out in moisture-free glassware under inert atmosphere. To a solution of ketone **SI18** (0.854 g, max. 2.37 mmol, 1.00 eq.) in anhydrous THF (30 mL) was added EtMgBr (1 M in THF, 9.49 mL, 9.49 mmol, 4.00 eq.) at room temperature. After 90 min, TLC indicated full conversion of the starting material ($R_f$ of **SI19** (*n*-heptane/ethyl acetate 1:3): 0.61). Saturated aqueous $NH_4Cl$ (20 mL) and EtOAc (100 mL) were added to the reaction mixture, the layers were separated and the organic layer was washed with saturated aqueous NaCl (20 mL). The organic layer was dried over $MgSO_4$, filtered, and concentrated under reduced pressure. The crude product of **SI19** was dissolved in 1,4-dioxane (20 mL) and aqueous HCl (1 N, 1 mL) and the reaction mixture was stirred for 30 min until full hydrolysis of the orthoester was observed by TLC. All volatiles were removed under reduced pressure and the residue was purified *via* HPLC (RP-18, 15 min, 30-40% MeCN in $H_2O$/TFA 0.1%) to yield **SI20** (0.126 g, 0.329 mmol, 13% over four steps)[11] as a colorless oil.

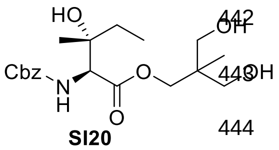

**SI20**

**$^1$H-NMR** (DMSO-$d_6$, 400 MHz): 7.42–7.26 (m, 5H, aryl-*H*), 7.21 (d, 1H, $J$ = 8.8 Hz, N*H*), 5.04 (s, 2H, Ph-C$H_2$), 4.07 (d, 1H, $J$ = 8.8 Hz, NH-C*H*), ~4.06 (s, v. br, 3H, O*H*), 3.90 (s, 2H, C$H_2$), 3.33–3.21 (m, 4H, C$H_2$-OH), 1.48 (m, 2H, C$H_2$-$CH_3$), 1.11 (s, 3H, C$H_3$), 0.85–0.76 (m, 6H, $CH_2$-C$H_3$, C$H_3$);

[11] 14:1 *dr* and >10:1 *dr* were determined for **SI20**:**SI27** by chirale HPLC and $^1$H-NMR analysis.

**$^{13}$C-NMR** (DMSO-d$_6$, 100 MHz): 170.8 (*C*O), 156.1 (*C*O-NH), 136.8 (aryl-*C*$_q$), 128.3 (aryl-*C*), 127.8 (aryl-*C*), 127.7 (aryl-*C*), 72.4 (*C*$_q$-OH), 67.0 (*C*H$_2$), 65.7 (Ph-*C*H$_2$), 63.5/63.5 (*C*H$_2$-OH), 61.7 (*C*H-NH), 40.5 (*C*$_q$-CH$_3$), 31.9 (*C*H$_2$-CH$_3$), 23.1 (*C*H$_3$), 16.5 (*C*H$_3$), 8.0 (CH$_2$-*C*H$_3$); **HRMS (ESI)** m/z calcd. for $C_{19}H_{29}NO_7Na$: (M+Na)$^+$, 406.1836; found: 406.1837 (M+Na)$^+$; **Chiral HPLC** (Chiralpak IF/181, 250x4.6 mm; *n*-heptane:EtOH:MeOH 5:1:1 + 0.1% TFA): **SI20** (Rt = 13.2 min) : ***ent*-SI20** (Rt = 8.2 min) 89.5 : 3.9 (92% *ee*); **Specific rotation** $[\alpha]_D^{20.0}$ = − 21.1 (c = 0.07; MeCN).

***Rac*-SI20** (0.300 g, 0.783 mmol, 37% over four steps)[12] was synthesized from ***rac*-SI7** (0.685 g, 2.13 mmol, 1.00 eq.) in analogous manner. In the conversion of ***rac*-SI18** to ***rac*-Si19**, an additional portion of EtMgBr (1 M in THF, 1.00 eq.) was added after 90 min and the reaction mixture was stirred for 60 min after addition. This modification resulted in a significantly improved overall yield. The NMR data are identical to the ones reported for **SI20**.

For ***rac*-SI20**: **Chiral HPLC** (Chiralpak IF/181, 250x4.6 mm; *n*-heptane:EtOH:MeOH 5:1:1 + 0.1% TFA): **SI20** (Rt = 13.5 min) : ***ent*-SI20** (Rt = 8.2 min) 47.9 : 44.6 (4% *ee*).

**3-Hydroxy-2-(hydroxymethyl)-2-methylpropyl (2*S*,3*S*)-2-(((benzyloxy)carbonyl)amino)-3-hydroxy-3-methylpenta-noate (SI27):**

The reaction was carried out in moisture-free glassware under inert atmosphere. To a solution of aldehyde **SI7** (0.830 g, 2.58 mmol, 1.00 eq.) in anhydrous THF (30 mL) was added EtMgBr (1 M in THF, 10.3 mL, 10.3 mmol, 4.00 eq.) at room temperature. After 90 min, TLC indicated full conversion of the starting material ($R_f$ of **SI23**/**SI24** (*n*-heptane/ethyl acetate 1:3): 0.58). Saturated aqueous $NH_4Cl$ (20 mL) and EtOAc (100 mL) were added to the reaction mixture, the layers were separated and the organic layer was washed with saturated aqueous NaCl (20 mL). The organic layer was dried over $MgSO_4$, filtered, and concentrated under reduced pressure. The crude mixture of **SI23**/**SI24** (0.813 g, 2.31 mmol, 90%) was used for the oxidation without further purification.

The reaction was carried out in moisture-free glassware under inert atmosphere. To a solution of $(COCl)_2$ (0.300 mL, 3.47 mmol, 1.50 eq.) in anhydrous DCM (20 mL) anhydrous DMSO (0.509 mL, 7.17 mmol, 3.10 eq.) was added carefully at −78 °C. After stirring for 15 min, the crude mixture of **SI23**/**SI24** (0.813 g, 2.31 mmol, 1.00 eq.) dissolved in anhydrous DCM (20 mL) was added. The reaction mixture was stirred for 1.5 h at −78 °C. Then DIPEA (2.24 mL, 12.7 mmol, 5.50 eq.) was added and the solution was stirred for 30 min at −78 °C and for further

[12] 12:1 *dr* and >10:1 *dr* were determined for **SI20**:**SI27** by chirale HPLC and $^1$H-NMR analysis.

30 min without cooling bath. A mixture of cooled toluene/saturated aqueous $NH_4Cl$ (4:1, 250 mL, pre-cooled to 0 °C) was added to the reaction mixture. The layers were separated and the organic layer was washed with saturated aqueous $NH_4Cl$ (3 x 50 mL, pre-cooled to 0 °C), with saturated aqueous $NaHCO_3$ (50 mL, pre-cooled to 0 °C), and with saturated aqueous NaCl (50 mL, pre-cooled to 0 °C). The organic layer was dried over $MgSO_4$, filtered, and concentrated under reduced pressure to yield crude ketone **SI25** (0.808 g, 2.31 mmol, quant., $R_f$ (*n*-heptane/ethyl acetate 1:3): 0.68) as slightly yellow oil which was used in the next stage without further purification.

The reaction was carried out in moisture-free glassware under inert atmosphere. To a solution of ketone **SI25** (0.808 g, max. 2.31 mmol, 1.00 eq.) in anhydrous THF (30 mL) was added MeMgBr (3 M in $Et_2O$, 3.08 mL, 9.25 mmol, 4.00 eq.) at room temperature. After 90 min, TLC indicated full conversion of the starting material ($R_f$ of **SI26** (*n*-heptane/ethyl acetate 1:3): 0.61). Saturated aqueous $NH_4Cl$ (20 mL) and EtOAc (100 mL) were added to the reaction mixture, the layers were separated and the organic layer was washed with saturated aqueous NaCl (20 mL). The organic layer was dried over $MgSO_4$, filtered, and concentrated under reduced pressure. The crude product of **SI26** was dissolved in 1,4-dioxane (20 mL) and aqueous HCl (1 N, 1 mL) and the reaction mixture was stirred for 30 min until full hydrolysis of the orthoester was observed by TLC. All volatiles were removed under reduced pressure and the residue was purified *via* HPLC (RP-18, 15 min, 30-40% MeCN in $H_2O$/TFA 0.1%) to yield **SI27** (0.218 g, 0.569 mmol, 22% over four steps).[13] as a colorless oil.

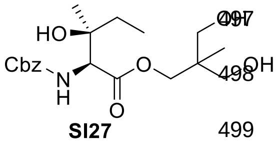

**SI27**

**$^1$H-NMR** (DMSO-$d_6$, 400 MHz): 7.42–7.26 (m, 6H, aryl-*H*, N*H*), 5.04 (s, 2H, Ph-C$H_2$), 4.54 (s, 1H, $C_q$-O*H*), 4.45 (t, 2H, $J$ = 5.2 Hz, $CH_2$-O*H*), 4.10 (d, 1H, $J$ = 9.2 Hz, NH-C*H*), 3.93 (d, 1H, $J$ = 10.7 Hz, C$H_2$), 3.87 (d, 1H, $J$ = 10.7 Hz, C$H_2$), 3.31–3.21 (m, 4H, C$H_2$-OH), 1.56–1.39 (m, 2H, C$H_2$-$CH_3$), 0.86 (s, 3H, C$H_3$), 0.84 (t, 3H, $J$ = 7.4 Hz, $CH_2$-C$H_3$), 0.80 (s, 3H, C$H_3$); **$^{13}$C-NMR** (DMSO-$d_6$, 100 MHz): 170.8 (*C*O), 156.1 (*C*O-NH), 136.9 (aryl-$C_q$), 128.4 (aryl-*C*), 127.9 (aryl-*C*), 127.7 (aryl-*C*), 72.7 ($C_q$-OH), 67.0 (*C*$H_2$), 65.6 (Ph-*C*$H_2$), 63.6 (*C*$H_2$-OH), 61.6 (*C*H-NH), 40.5 ($C_q$-$CH_3$), 31.6 (*C*$H_2$-$CH_3$), 23.1 (*C*$H_3$), 16.5 (*C*$H_3$), 7.9 ($CH_2$-*C*$H_3$); **HRMS (ESI)** m/z calcd. for $C_{19}H_{29}NO_7Na$: $(M+Na)^+$, 406.1836; found: 406.1839 $(M+Na)^+$; **Chiral HPLC** (Chiralpak IF/181, 250x4.6 mm; *n*-heptane:EtOH:MeOH 5:1:1 + 0.1% TFA): **SI27** (Rt = 22.9 min) : ***ent*-SI27** (Rt = 6.9 min) 91.7 :5.2 (89% *ee*); **Specific rotation** $[\alpha]_D^{20.0}$ = − 11.6 (c = 0.35; MeCN).

[13] 32:1 *dr* and >10:1 *dr* were determined for **SI27**:**SI20** by chirale HPLC and $^1$H-NMR analysis.

***Rac*-SI27** (0.337 g, 0.879 mmol, 41% over four steps)[14] was synthesized from ***rac*-SI7** (0.685 g, 2.13 mmol, 1.00 eq.) in analogous manner. In the conversion of ***rac*-SI25** to ***rac*-Si26**, an additional portion of MeMgBr (3 M in $Et_2O$, 1.00 eq.) was added after 90 min and the reaction mixture was stirred for 60 min after addition. This modification resulted in a significantly improved overall yield. The NMR data are identical to the ones reported for **SI27**.

For ***rac*-SI27**: **Chiral HPLC** (Chiralpak IF/181, 250x4.6 mm; *n*-heptane:EtOH:MeOH 5:1:1 + 0.1% TFA): **SI27** (Rt = 23.0 min) : ***ent*-SI27** (Rt = 6.9 min) 51.4 :45.7 (6% *ee*).

**(2*S*,3*R*)-2-(((Benzyloxy)carbonyl)amino)-3-hydroxy-3-methylpentanoic acid (SI21):**

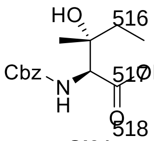

**SI21**

To a solution of **SI20** (0.084 g, 0.22 mmol, 1.00 eq.) in MeCN/$H_2O$ (4:1, 3.2 mL/0.8 mL) was added LiOH (1 N in $H_2O$, 0.22 mL, 0.22 mmol, 1.00 eq.) at room temperature. After 20 min, another portion of LiOH (1 N in $H_2O$, 0.11 mL, 0.11 mmol, 0.50 eq.) was added to reach full conversion of starting material after 30 min monitored by LC-MS. The reaction was stopped by addition of acetic acid (0.1 mL). All volatiles were removed under reduced pressure and the residue was purified *via* HPLC (RP-18, 15 min, 5-55% MeCN in $H_2O$/TFA 0.1%) to yield **SI21** (0.021 g, 0.075 mmol, 34%)[15] as a colorless oil.[16]

**$^1$H-NMR** (DMSO-$d_6$, 400 MHz): 7.42–7.26 (m, 5H, aryl-*H*), 7.03 (d, 1H, $J$ = 8.8 Hz, N*H*), 5.04 (s, 2H, Ph-C$H_2$), 3.96 (d, 1H, $J$ = 9.0 Hz, NH-C*H*), 1.56–1.34 (m, 2H, C$H_2$-$CH_3$), 1.11 (s, 3H, C$H_3$), 0.80 (t, 3H, $J$ = 7.5 Hz, $CH_2$-C$H_3$); **$^{13}$C-NMR** (DMSO-$d_6$, 100 MHz): 172.3 (*C*OOH), 156.2 (*C*O-NH), 137.0 (aryl-$C_q$), 128.4 (aryl-*C*), 127.8 (aryl-*C*), 127.7 (aryl-*C*), 72.4 ($C_q$-OH), 65.6 (Ph-$CH_2$), 61.4 (*C*H-NH), 32.0 ($CH_2$-$CH_3$), 23.1 (*C*$H_3$), 8.0 ($CH_2$-$CH_3$); **HRMS (ESI)** m/z calcd. for $C_{14}H_{19}NO_5Na$: $(M+Na)^+$, 304.1155; found: 304.1154 $(M+Na)^+$; **Chiral HPLC** (Chiralpak IF/181, 250x4.6 mm; *n*-heptane:EtOH 10:1 + 0.1% TFA): **SI21** (Rt = 16.7 min) : ***ent*-SI21** (Rt = 29.3 min) 75.2 : 2.6 (93% *ee*); **Specific rotation** $[\alpha]_D^{19.1}$ = 0 (c = 0.19; MeCN).[17]

***Rac*-SI21** (0.072 g, 0.26 mmol, 38%)[18] and ***rac*-SI22** (0.038 g, 0.22 mmol, 32%) were synthesized from ***rac*-SI20** (0.260 g, 0.678 mmol, 1.00 eq.) in analogous manner. The NMR

[14] 33:1 *dr* and >10:1 *dr* were determined for **SI27**:**SI20** by chirale HPLC and $^1$H-NMR analysis.
[15] 3.5:1 *dr* and 3:1 *dr* were determined for **SI21**:**SI28** by chirale HPLC and $^1$H-NMR analysis.
[16] The formation of **SI22** was observed *via* LC-MS, but was not isolated for the *S*-series.
[17] Since the sample is a mixture of diastereomers and the measured rotation was close to 0, no specific rotation was determined.
[18] 6:1 *dr* and >5:1 *dr* were determined for **SI21**:**SI28** by chirale HPLC and $^1$H-NMR analysis.

data of ***rac*-SI21** are identical to the ones reported for **SI21**. ***Rac*-SI21** was obtained as colorless oil and ***rac*-SI22** as colorless solid.

For ***rac*-SI21**: **Chiral HPLC** (Chiralpak IF/181, 250x4.6 mm; *n*-heptane:EtOH 10:1 + 0.1% TFA): **SI21** (Rt = 16.7 min) : ***ent*-SI21** (Rt = 28.9 min) 43.0 : 43.0 (0% *ee*).

**(4*S*,5*R*)-5-Ethyl-5-methyl-2-oxooxazolidine-4-carboxylic acid (ent-SI22):**

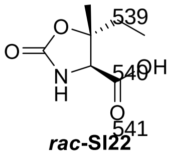

***rac*-SI22**

**$^1$H-NMR** (DMSO-$d_6$, 400 MHz): 13.23 (s, br, 1H, COO*H*), 7.89 (s, 1H, N*H*), 4.03 (d, 1H, *J* = 0.6 Hz, C*H*), 1.72 (q, 2H, *J* = 7.5 Hz, C*H*$_2$), 1.24 (s, 3H, C*H*$_3$), 0.91 (t, 3H, *J* = 7.5 Hz, CH$_2$-C*H*$_3$); **$^{13}$C-NMR** (DMSO-$d_6$, 100 MHz): 171.3 (*C*OOH), 157.4 (*C*O-NH), 82.7 (O-*C*$_q$), 61.2 (*C*H-NH), 33.2 (*C*H$_2$-CH$_3$), 20.7 (*C*H$_3$), 7.5 (CH$_2$-*C*H$_3$); **HRMS (ESI)** m/z calcd. for $C_7H_{12}NO_4$: (M+H)$^+$, 174.0761; found: 174.0761 (M+H)$^+$.

**(2*S*,3*S*)-2-(((Benzyloxy)carbonyl)amino)-3-hydroxy-3-methylpentanoic acid (SI28):**

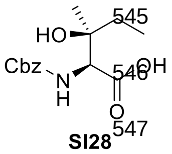

**SI28**

To a solution of **SI27** (0.178 g, 0.464 mmol, 1.00 eq.) in MeCN/$H_2O$ (4:1, 7.2 mL/1.8 mL) was added LiOH (1 N in $H_2O$, 0.46 mL, 0.46 mmol, 1.00 eq.) at room temperature. After 20 min, another portion of LiOH (1 N in $H_2O$, 0.23 mL, 0.23 mmol, 0.50 eq.) was added to reach full conversion of starting material after 30 min monitored by LC-MS. The reaction was stopped by addition of acetic acid (0.1 mL). All volatiles were removed under reduced pressure and the residue was purified *via* HPLC (RP-18, 15 min,10-50% MeCN in $H_2O$/TFA 0.1%) to yield **SI28** (0.051 g, 0.18 mmol, 39%)[19] as a colorless oil.[20]

**$^1$H-NMR** (DMSO-$d_6$, 400 MHz): 7.42–7.27 (m, 5H, aryl-*H*), 7.18 (d, 1H, *J* = 9.2 Hz, N*H*), 5.05 (s, 2H, Ph-C*H*$_2$), 4.01 (d, 1H, *J* = 9.2 Hz, NH-C*H*), 1.58–1.40 (m, 2H, C*H*$_2$-CH$_3$), 1.09 (s, 3H, C*H*$_3$), 0.84 (t, 3H, *J* = 7.4 Hz, CH$_2$-C*H*$_3$); **$^{13}$C-NMR** (DMSO-$d_6$, 100 MHz): 172.3 (*C*OOH), 156.1 (*C*O-NH), 137.0 (aryl-*C*$_q$), 128.3 (aryl-*C*), 127.8 (aryl-*C*), 127.7 (aryl-*C*), 72.6 (*C*$_q$-OH), 65.5 (Ph-*C*H$_2$), 61.2 (*C*H-NH), 31.5 (*C*H$_2$-CH$_3$), 23.1 (*C*H$_3$), 7.8 (CH$_2$-*C*H$_3$); **HRMS (ESI)** m/z calcd. for $C_{14}H_{19}NO_5Na$: (M+Na)$^+$, 304.1155; found: 304.1157 (M+Na)$^+$; **Chiral HPLC** (Chiralpak IF/181, 250x4.6 mm; *n*-heptane:EtOH 10:1 + 0.1% TFA): **SI28** (Rt = 20.6 min) : ***ent*-SI28** (Rt = 27.2 min) 94.2 : 5.1 (90% *ee*); **Specific rotation** $[\alpha]_D^{19.9}$ = + 18.5 (c = 0.05; MeCN).

---

[19] 138:1 *dr* and >100:1 *dr* were determined for **SI28**:**SI21** by chirale HPLC and $^1$H-NMR analysis.

[20] The formation of **SI29** was observed *via* LC-MS, but was not isolated for the *S*-series.

**(4*S*,5*S*)-5-Ethyl-5-methyl-2-oxooxazolidine-4-carboxylic acid (SI29):**

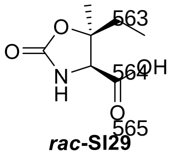

***rac*-SI29**

**$^1$H-NMR** (DMSO-$d_6$, 400 MHz): 13.24 (s, br, 1H, COO*H*), 7.88 (s, 1H, N*H*), 4.05 (d, 1H, *J* = 0.6 Hz, C*H*), 1.73 (q, 2H, *J* = 7.5 Hz, C$H_2$), 1.24 (s, 3H, C$H_3$), 0.91 (t, 3H, *J* = 7.5 Hz, $CH_2$-C$H_3$); **$^{13}$C-NMR** (DMSO-$d_6$, 100 MHz): 171.0 (*C*OOH), 157.4 (*C*O-NH), 82.7 (O-$C_q$), 63.4 (*C*H-NH), 28.3 (*C*$H_2$-$CH_3$), 24.3 (*C*$H_3$), 7.7 ($CH_2$-*C*$H_3$); **HRMS (ESI)** m/z calcd. for $C_7H_{12}NO_4$: $(M+H)^+$, 174.0761; found: 174.0762 $(M+H)^+$.

***Rac*-SI28** (0.132 g, 0.469 mmol, 61%)[21] and ***rac*-SI29** (0.021 g, 0.12 mmol, 16%) were synthesized from ***rac*-SI27** (0.297 g, 0.775 mmol, 1.00 eq.) in analogous manner. The NMR data of ***rac*-SI28** are identical to the ones reported for **SI28**. ***Rac*-SI28** was obtained as colorless oil and ***rac*-SI29** as colorless solid.

For ***rac*-SI28**: **Chiral HPLC** (Chiralpak IF/181, 250x4.6 mm; *n*-heptane:EtOH 10:1 + 0.1% TFA): **SI28** (Rt = 20.8 min) : ***ent*-SI28** (Rt = 26.8 min) 48.4 : 48.7 (0% *ee*).

[21] 33:1 *dr* and >50:1 *dr* were determined for **SI28**:**SI21** by chirale HPLC and $^1$H-NMR analysis.

#### 2. NMR Spectra

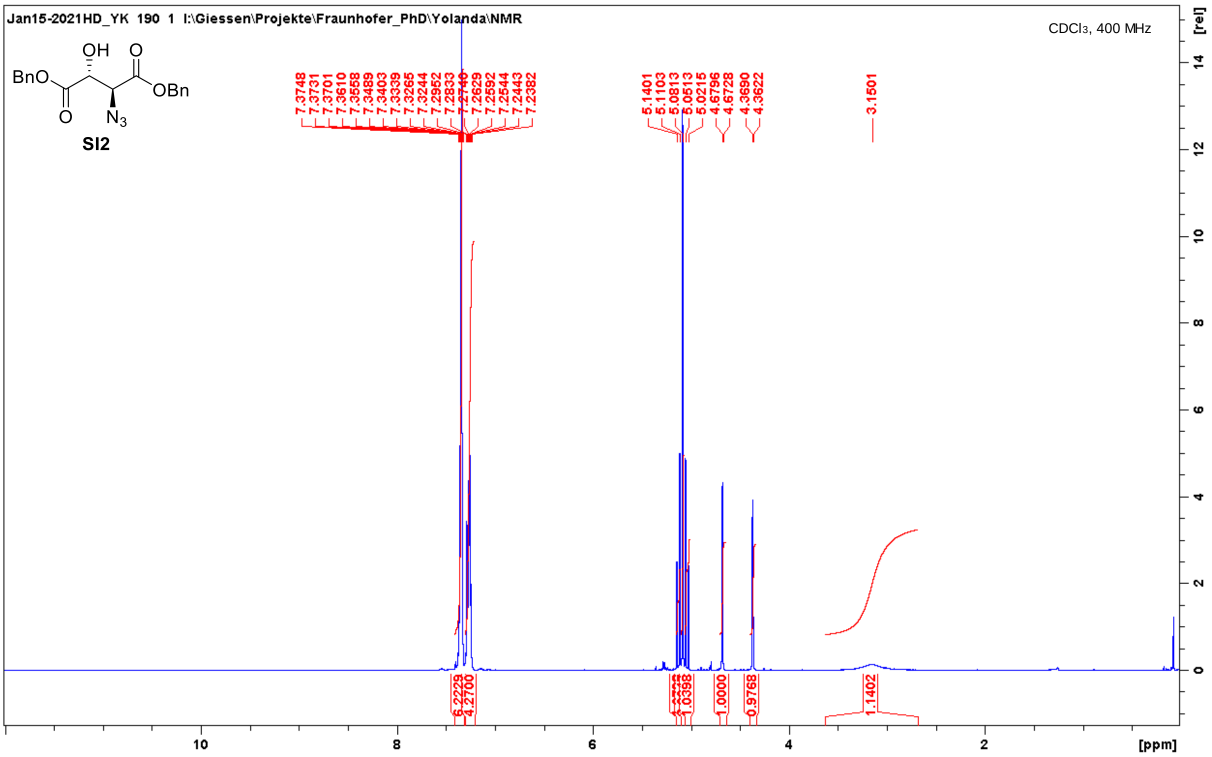

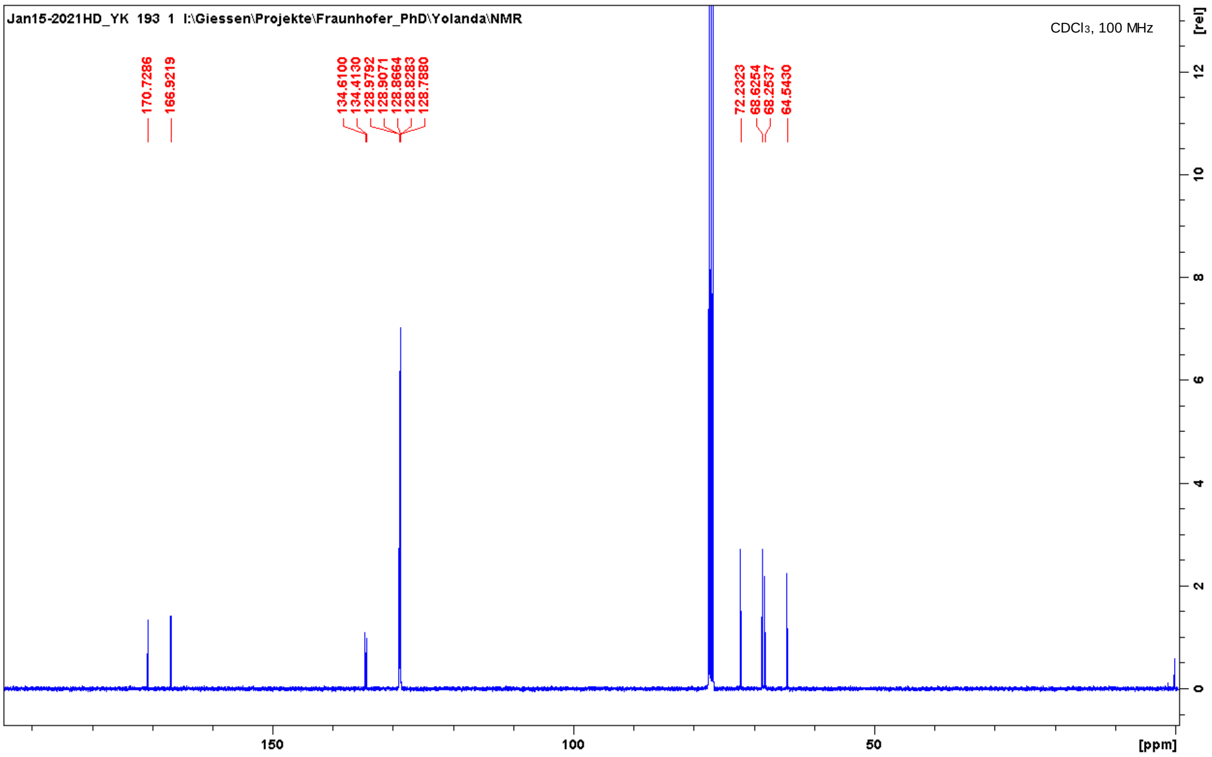

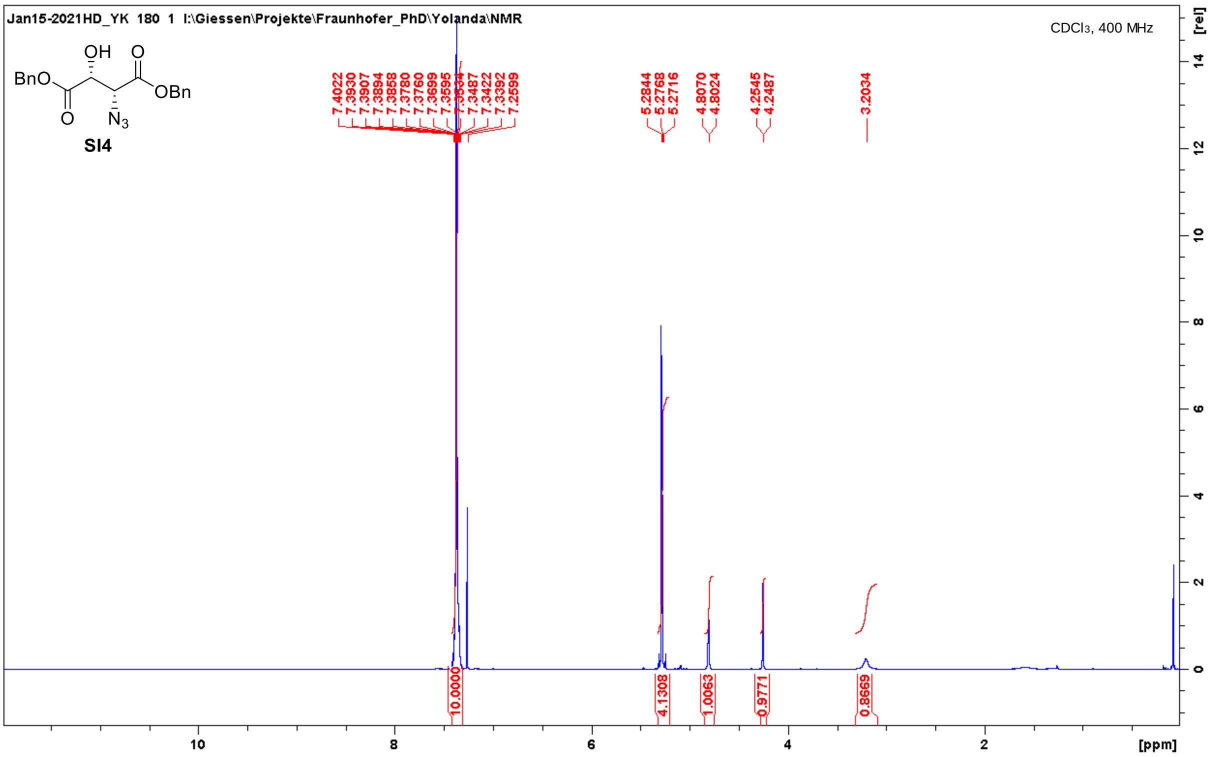
Jan15-2021HD_YK 180 1 I:\Giessen\Projekte\Fraunhofer_PhD\Yolanda\NMR
CDCl3, 400 MHz
SI4
7.4022
7.3930
7.3907
7.3894
7.3858
7.3780
7.3760
7.3699
7.3595
7.3534
7.3487
7.3422
7.3392
7.2599
5.2844
5.2768
5.2716
4.8070
4.8024
4.2545
4.2487
3.2034
10.0000
4.1308
1.0063
0.9771
0.8669
[ppm]
[rel]

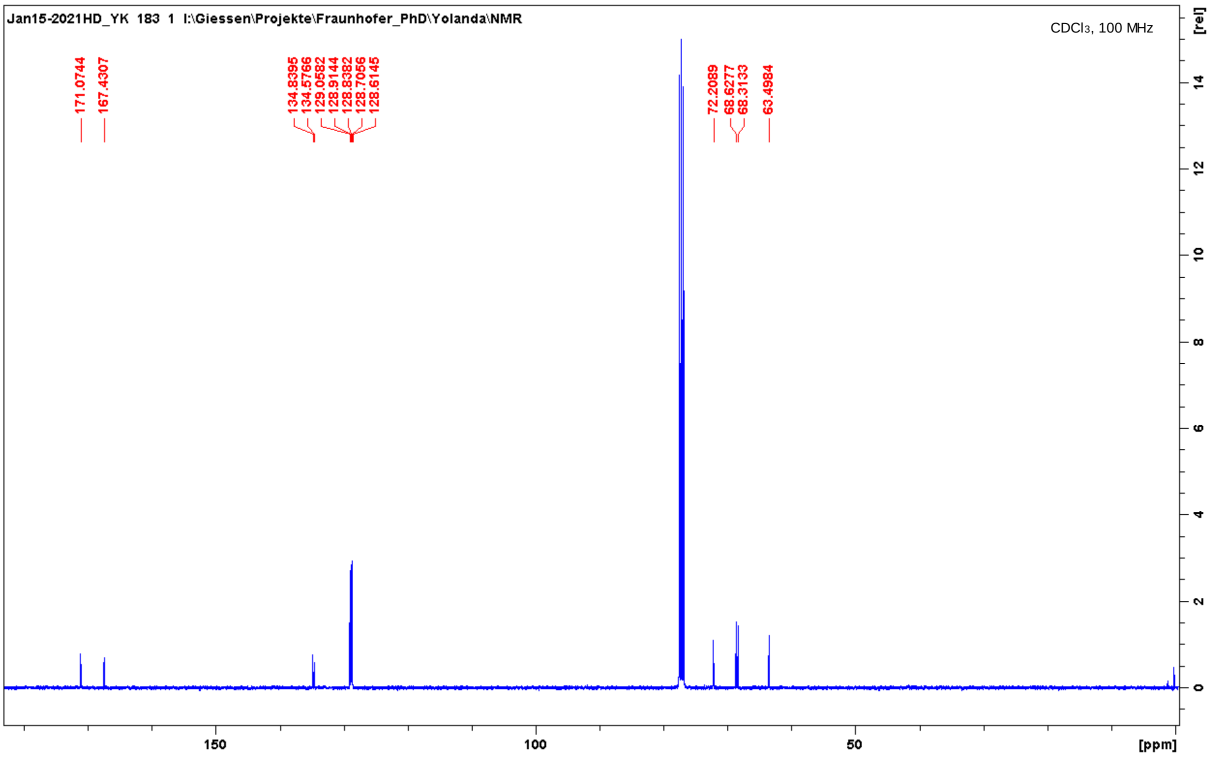
Jan15-2021HD_YK 183 1 I:\Giessen\Projekte\Fraunhofer_PhD\Yolanda\NMR
CDCl3, 100 MHz
171.0744
167.4307
134.8395
134.5766
129.0582
128.9144
128.8382
128.7056
128.6145
72.2089
68.6277
68.3133
63.4984
[ppm]
[rel]

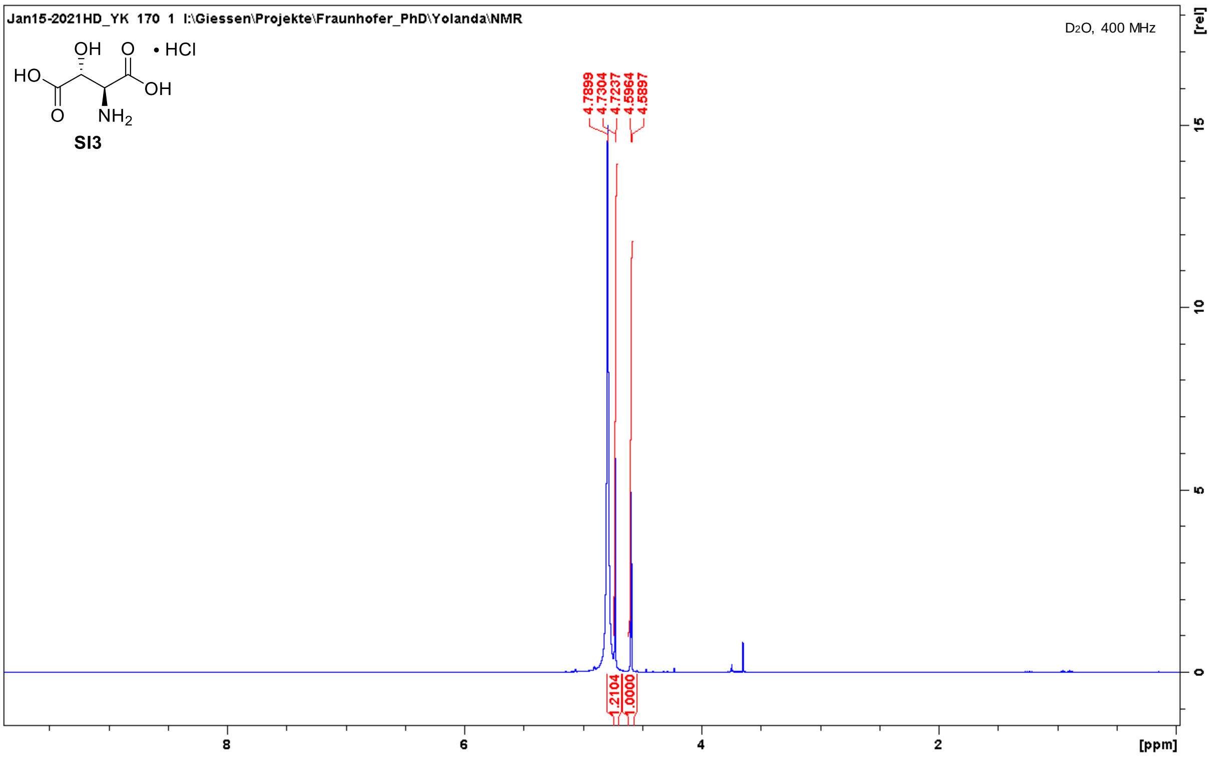
Jan15-2021HD_YK 170 1 I:\Giessen\Projekte\Fraunhofer_PhD\Yolanda\NMR
D2O, 400 MHz
OH
O
• HCl
HO
OH
O
NH2
SI3
4.7899
4.7304
4.7237
4.5964
4.5897
1.2104
1.0000
[rel]
15
10
5
0
8
6
4
2
[ppm]

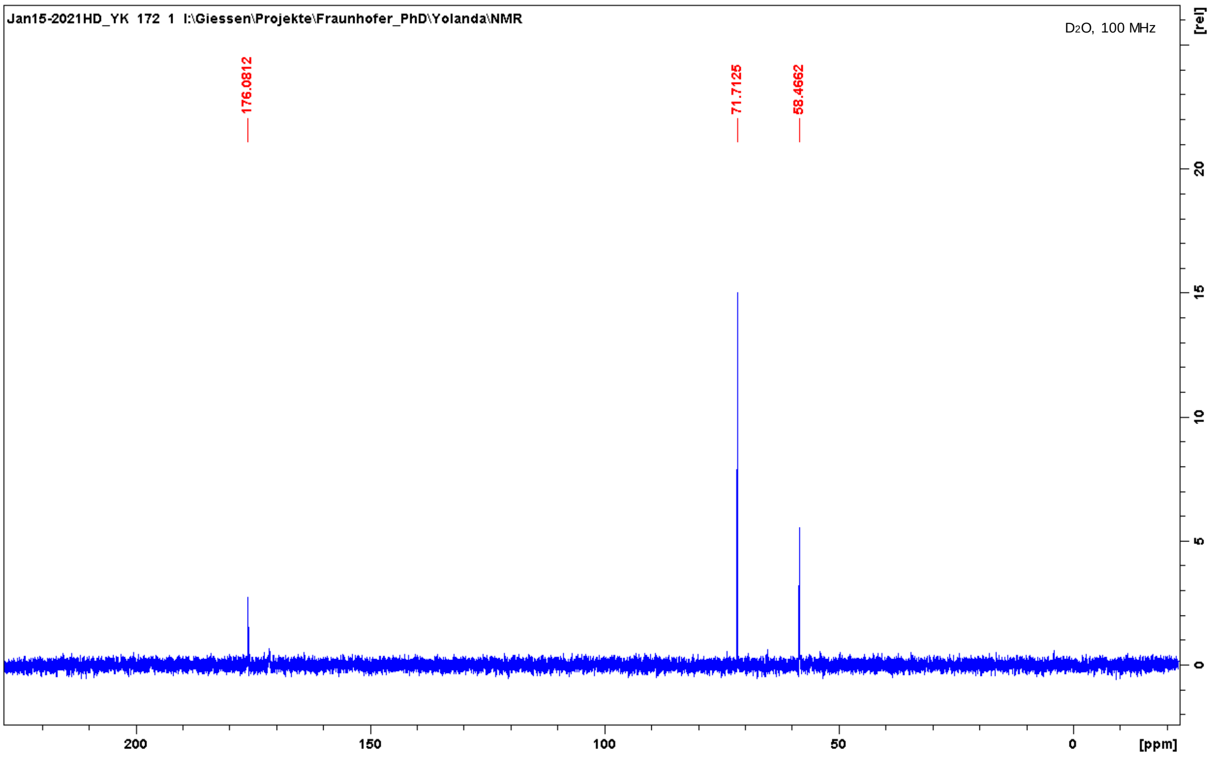
Jan15-2021HD_YK 172 1 I:\Giessen\Projekte\Fraunhofer_PhD\Yolanda\NMR
D2O, 100 MHz
176.0812
71.7125
58.4662
[rel]
20
15
10
5
0
200
150
100
50
0
[ppm]

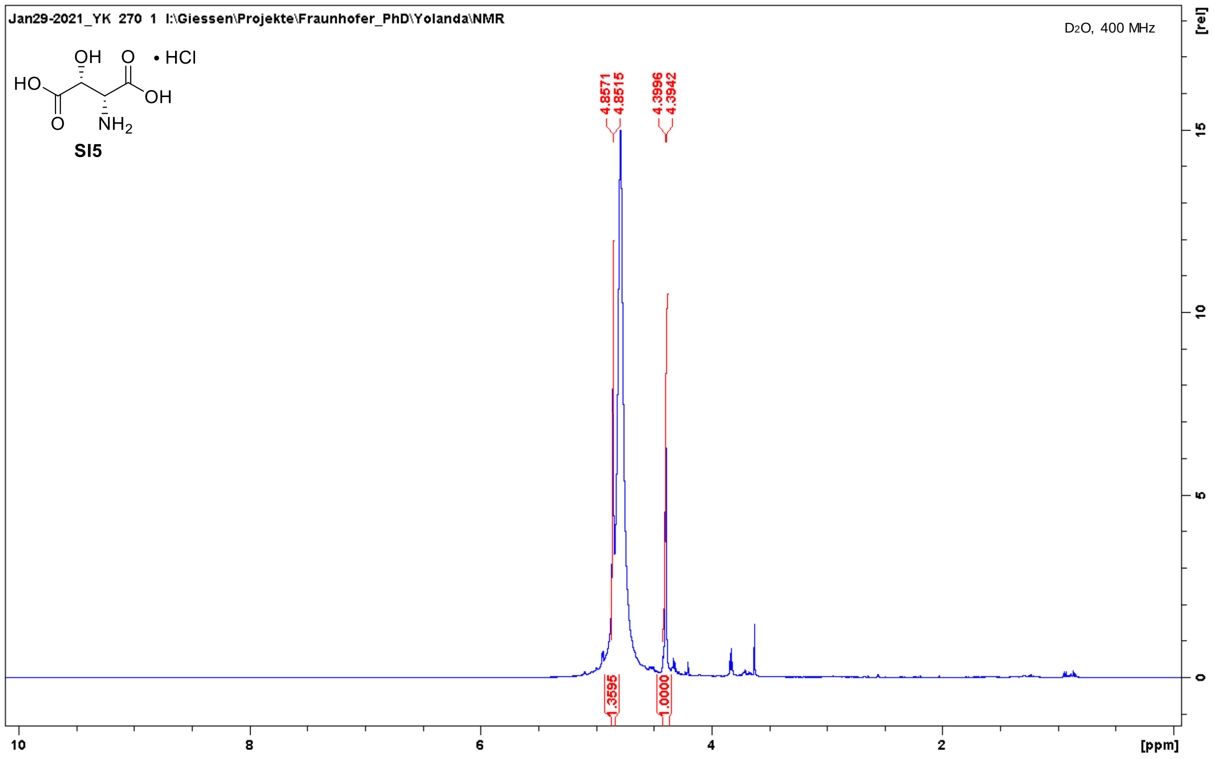
Jan29-2021_YK 270 1 I:\Giessen\Projekte\Fraunhofer_PhD\Yolanda\NMR
D2O, 400 MHz
• HCl
SI5
4.8571
4.8515
4.3996
4.3942
1.3595
1.0000
[rel]
[ppm]

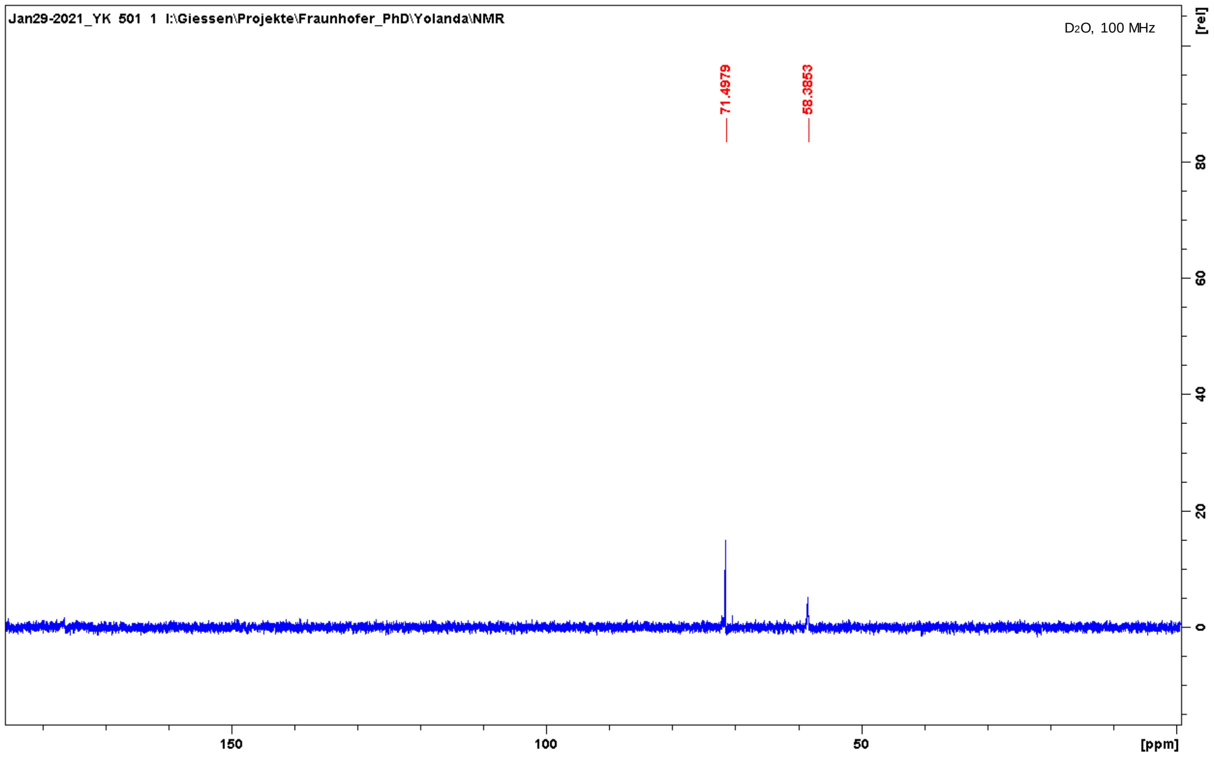
Jan29-2021_YK 501 1 I:\Giessen\Projekte\Fraunhofer_PhD\Yolanda\NMR
D2O, 100 MHz
71.4979
58.3853
[rel]
[ppm]

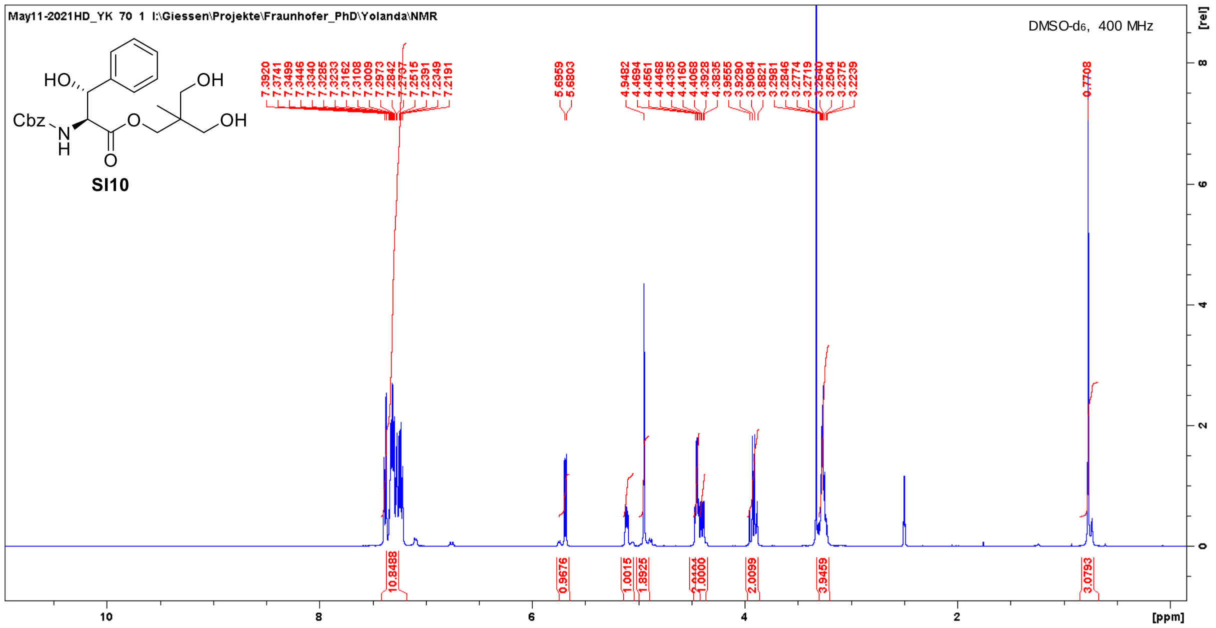
May11-2021HD_YK 70 1 I:\Giessen\Projekte\Fraunhofer_PhD\Yolanda\NMR
DMSO-d6, 400 MHz
HO
OH
Cbz
N
H
O
O
OH
SI10
7.3920
7.3741
7.3499
7.3446
7.3340
7.3285
7.3233
7.3162
7.3108
7.3009
7.2973
7.2842
7.2515
7.2391
7.2349
7.2191
5.6959
5.6803
4.9482
4.4561
4.4468
4.4335
4.4160
4.4068
4.3928
4.3835
3.9555
3.9290
3.9084
3.8821
3.2981
3.2846
3.2774
3.2719
3.2640
3.2504
3.2375
3.2239
0.7708
10.8488
0.9676
1.0015
1.8925
1.0000
2.0099
3.9459
3.0793
[ppm]
[rel]

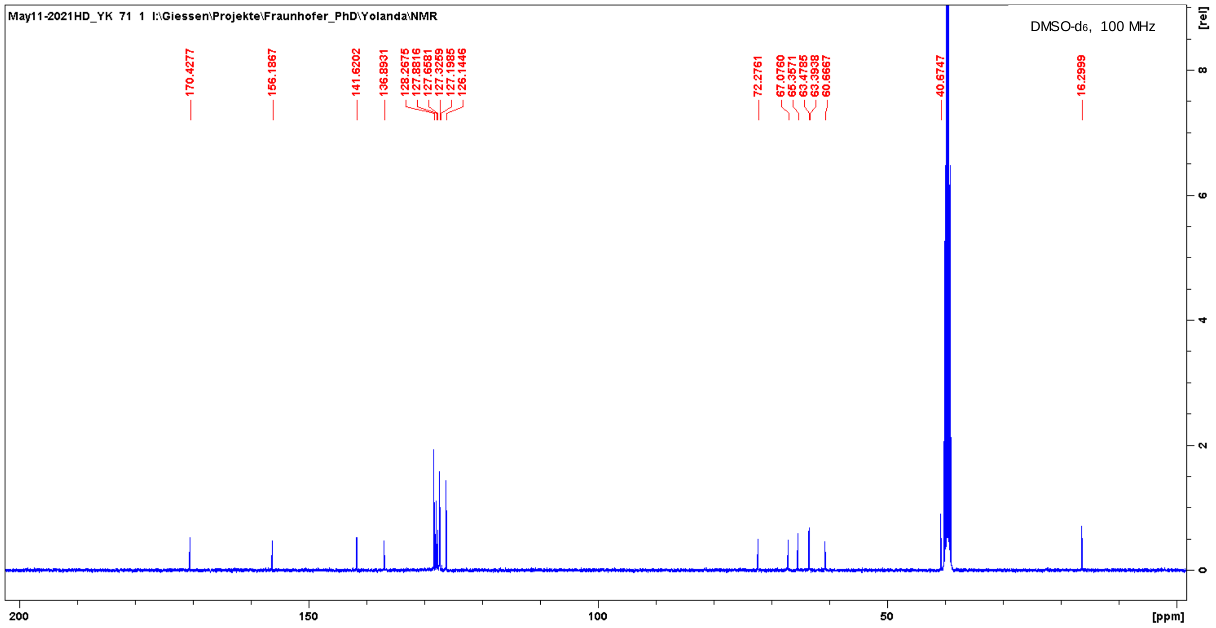
May11-2021HD_YK 71 1 I:\Giessen\Projekte\Fraunhofer_PhD\Yolanda\NMR
DMSO-d6, 100 MHz
170.4277
156.1867
141.6202
136.8931
128.2675
127.8816
127.6581
127.3259
127.1985
126.1446
72.2761
67.0760
65.3571
63.4785
63.3938
60.6667
40.6747
16.2999
[ppm]
[rel]

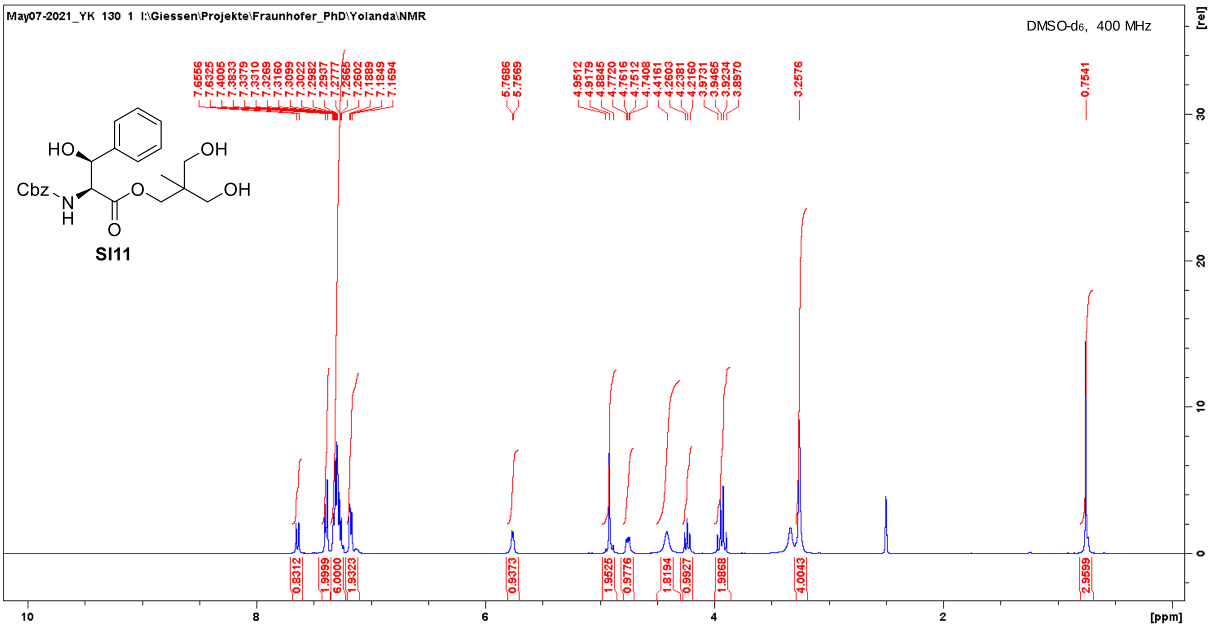
May07-2021_YK 130 1 I:\Giessen\Projekte\Fraunhofer_PhD\Yolanda\NMR
DMSO-d6, 400 MHz
HO
OH
Cbz
N
H
O
O
OH
SI11
[rel]
[ppm]

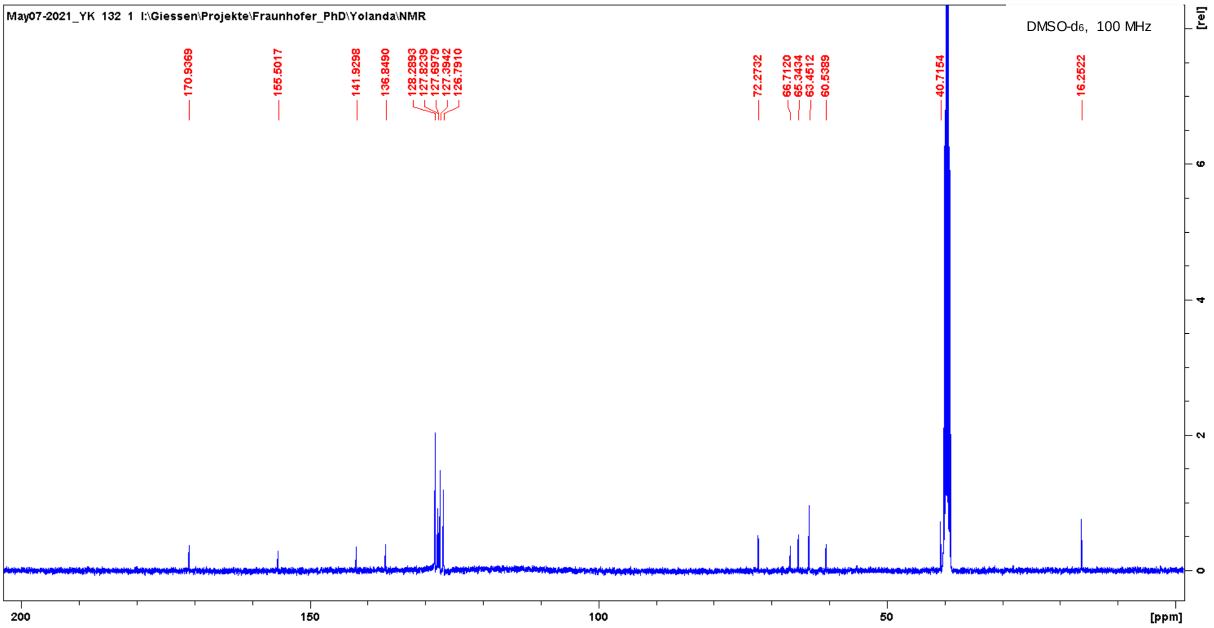
May07-2021_YK 132 1 I:\Giessen\Projekte\Fraunhofer_PhD\Yolanda\NMR
DMSO-d6, 100 MHz
170.9369
155.5017
141.9298
136.8490
72.2732
40.7154
16.2522
[rel]
[ppm]

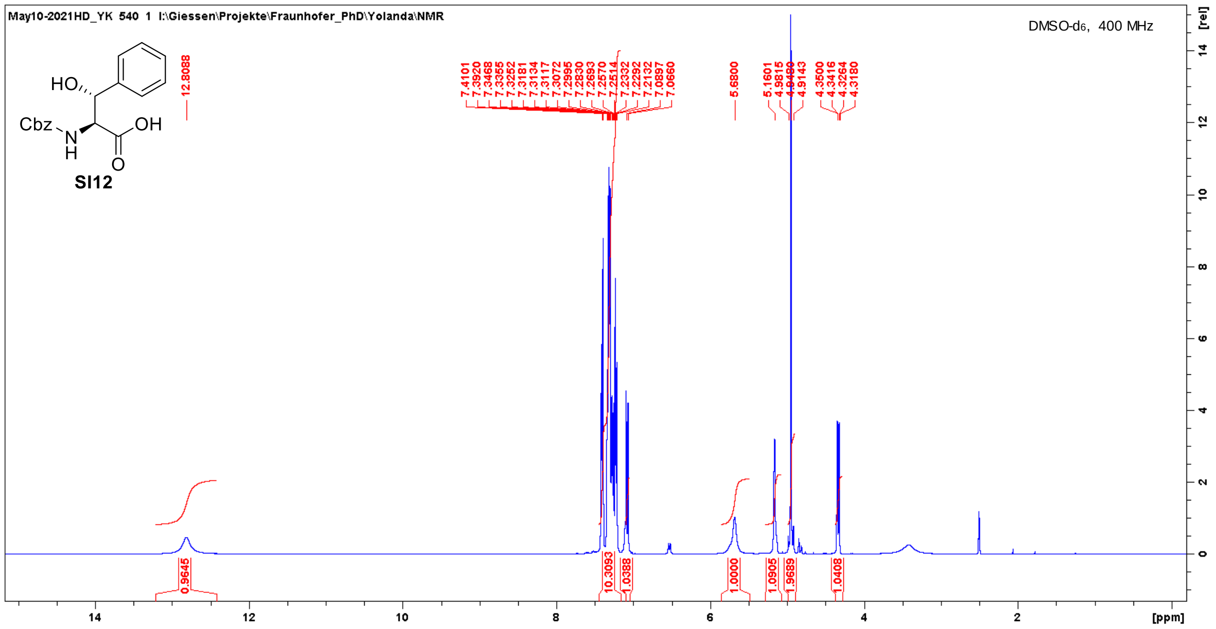
May10-2021HD_YK 540 1 I:\Giessen\Projekte\Fraunhofer_PhD\Yolanda\NMR
DMSO-d6, 400 MHz
HO
Cbz
N
H
OH
O
SI12
12.8088
7.4101 7.3920 7.3468 7.3365 7.3252 7.3181 7.3134 7.3117 7.3072 7.2995 7.2893 7.2830 7.2570 7.2514 7.2332 7.2292 7.2132 7.0897 7.0660
5.6800
5.1601 4.9815
4.9143
4.3500 4.3416 4.3264 4.3180
0.9645
10.3093
1.0388
1.0000
1.0905
1.9689
1.0408
[rel]
[ppm]

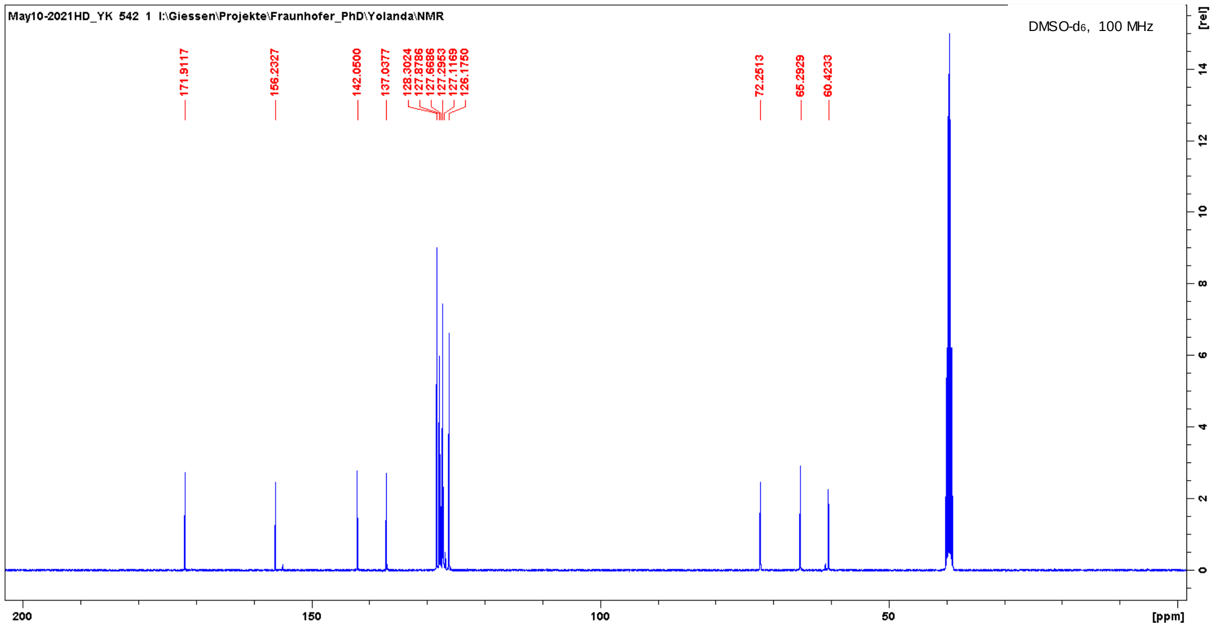
May10-2021HD_YK 542 1 I:\Giessen\Projekte\Fraunhofer_PhD\Yolanda\NMR
DMSO-d6, 100 MHz
171.9117
156.2327
142.0500
137.0377
128.3024 127.8786 127.6686 127.2953 127.1169 126.1750
72.2513
65.2929
60.4233
[rel]
[ppm]

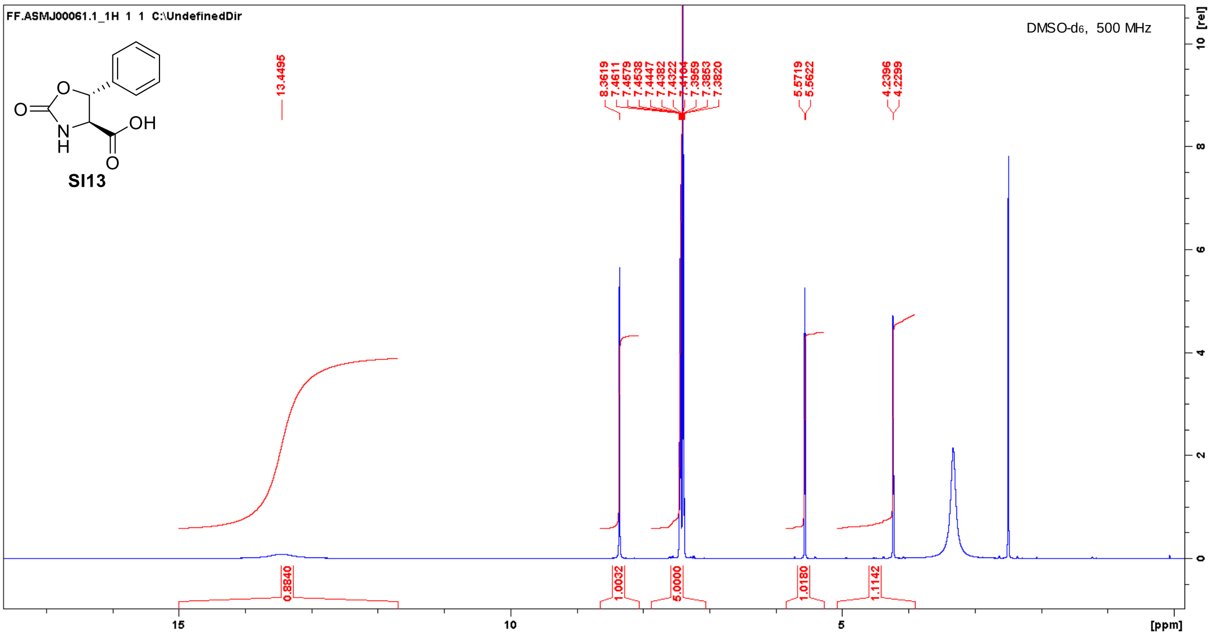
FF.ASMJ00061.1_1H 1 1 C:\UndefinedDir
DMSO-d6, 500 MHz
SI13
O
O
N
H
OH
O
13.4495
8.3619
7.4611
7.4579
7.4538
7.4447
7.4382
7.4322
7.4104
7.3959
7.3853
7.3820
5.5719
5.5622
4.2396
4.2299
0.8840
1.0032
5.0000
1.0180
1.1142
15
10
5
[ppm]
[rel]
10
8
6
4
2
0

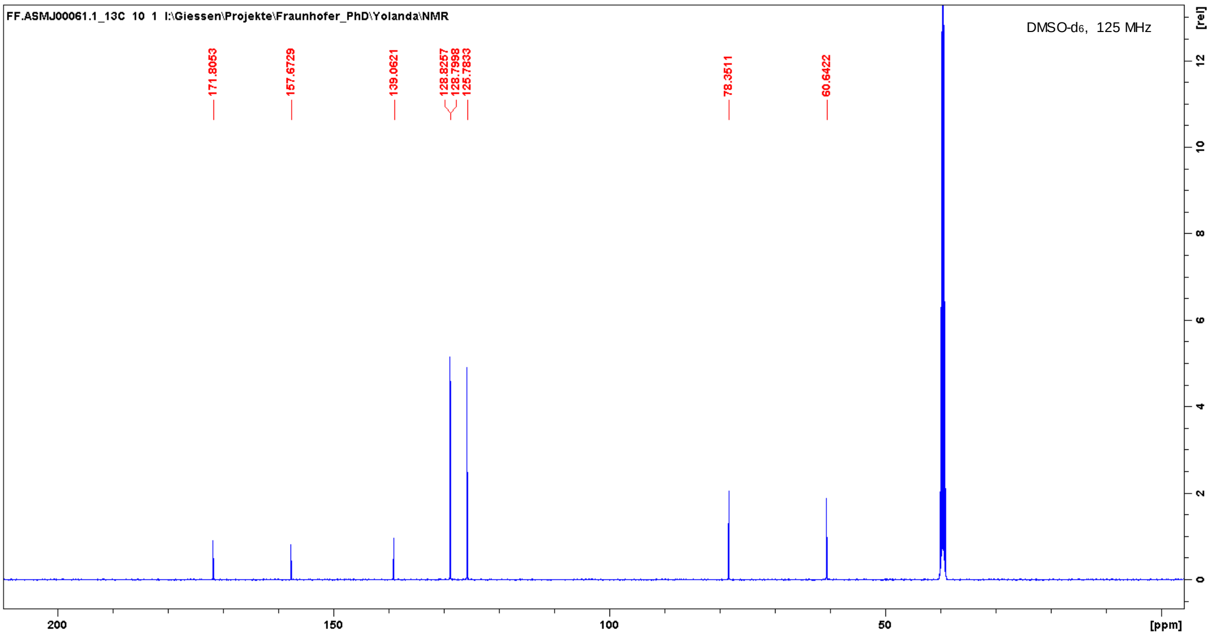
FF.ASMJ00061.1_13C 10 1 I:\Giessen\Projekte\Fraunhofer_PhD\Yolanda\NMR
DMSO-d6, 125 MHz
171.8053
157.6729
139.0621
128.8257
128.7998
125.7833
78.3511
60.6422
200
150
100
50
[ppm]
[rel]
12
10
8
6
4
2
0

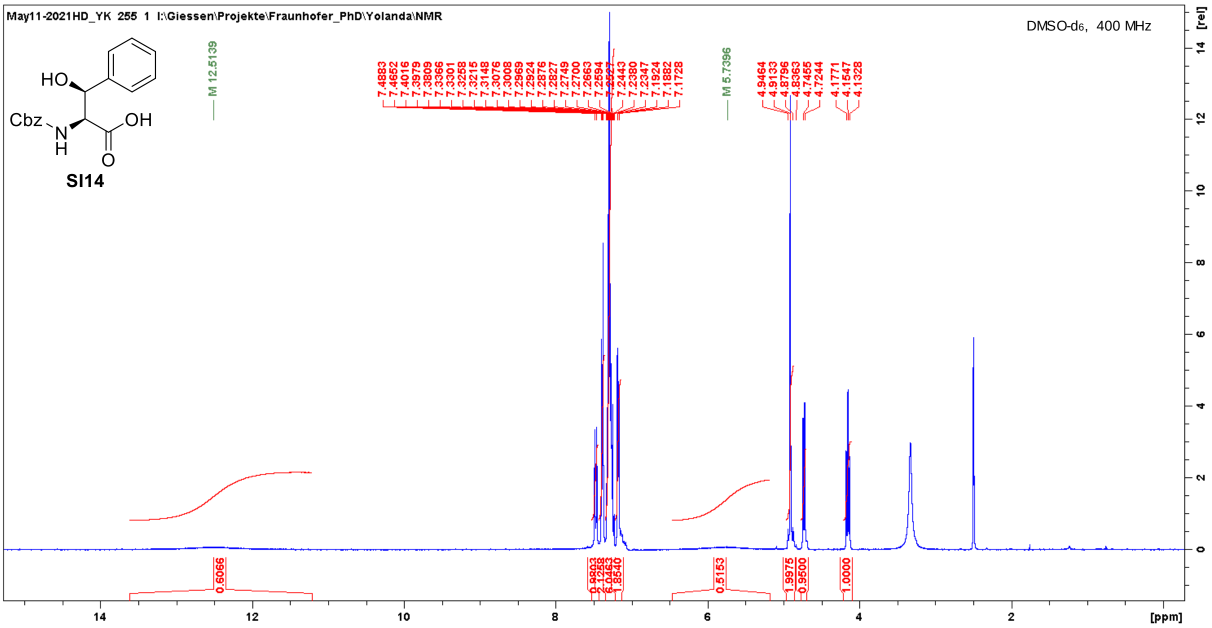
May11-2021HD_YK 255 1 I:\Giessen\Projekte\Fraunhofer_PhD\Yolanda\NMR
DMSO-d6, 400 MHz
HO
Cbz
N
H
OH
O
SI14
M 12.5139
7.4883 7.4652 7.4016 7.3979 7.3809 7.3366 7.3301 7.3258 7.3215 7.3148 7.3076 7.3008 7.2969 7.2924 7.2876 7.2827 7.2749 7.2700 7.2663 7.2594 7.2521 7.2443 7.2380 7.2347 7.1924 7.1882 7.1728
M 5.7396
4.9464 4.9133 4.8796 4.8363 4.7455 4.7244 4.1771 4.1547 4.1328
0.6066
0.5153
1.9975
0.9500
1.0000
[ppm]
[rel]

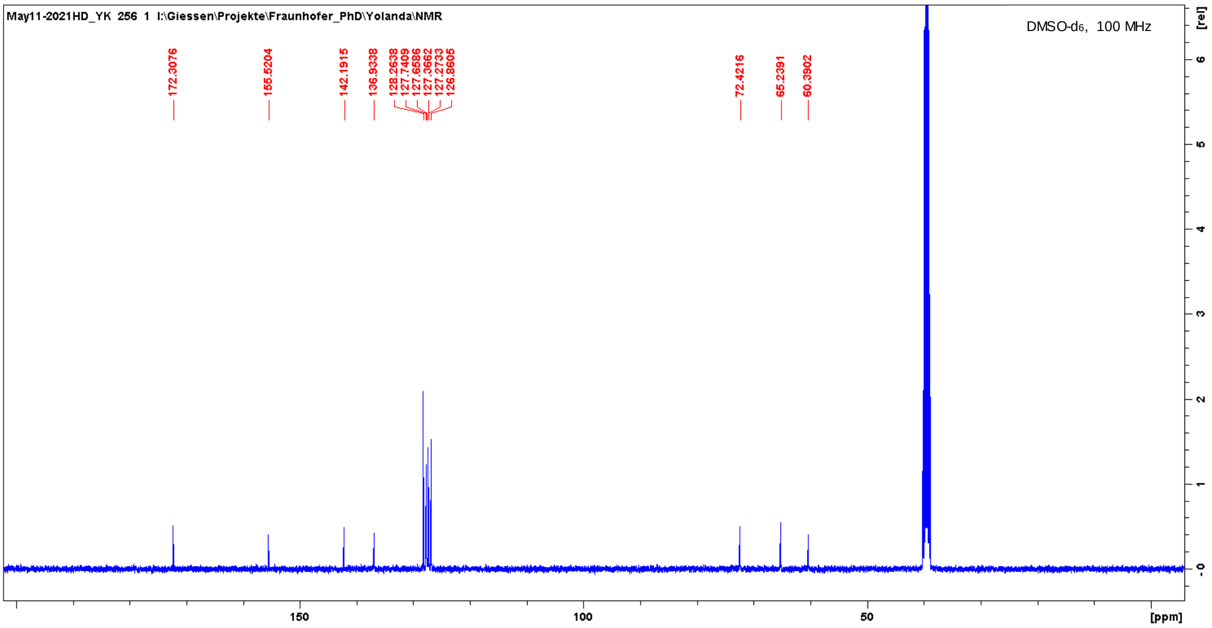
May11-2021HD_YK 256 1 I:\Giessen\Projekte\Fraunhofer_PhD\Yolanda\NMR
DMSO-d6, 100 MHz
172.3076
155.5204
142.1915
136.9338
128.2638 127.7409 127.6586 127.3662 127.2733 126.8605
72.4216
65.2391
60.3902
[ppm]
[rel]

604

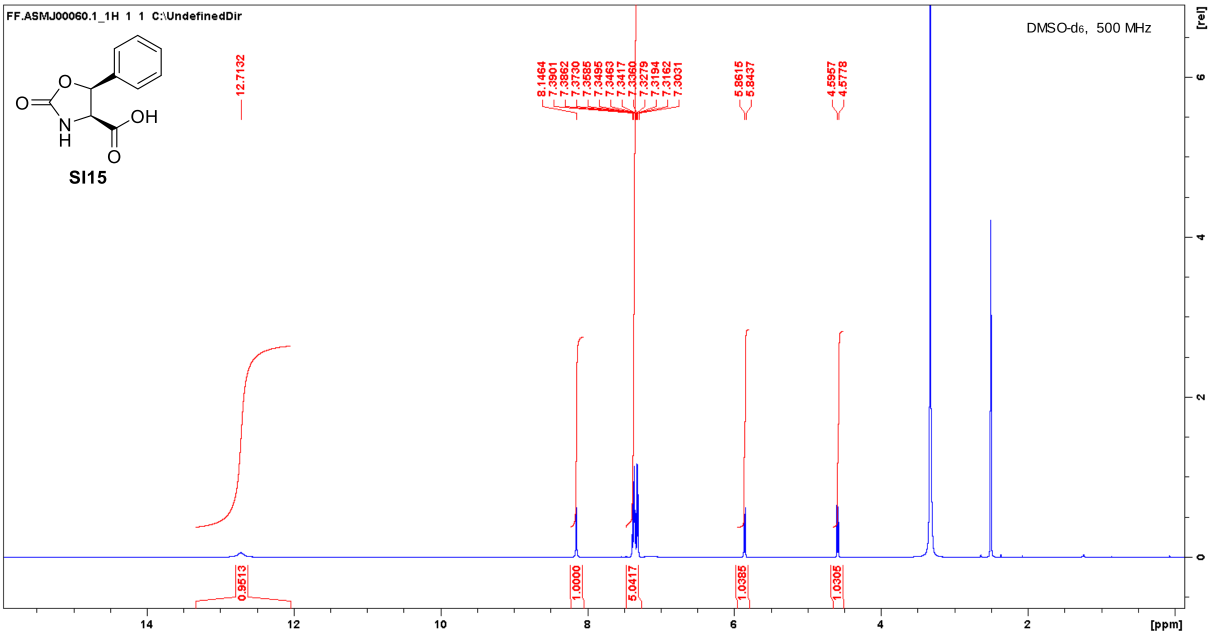
FF.ASMJ00060.1_1H 1 1 C:\UndefinedDir
DMSO-d6, 500 MHz
SI15
OH
O
N
H
12.7132
8.1464
7.3901
7.3862
7.3730
7.3585
7.3495
7.3463
7.3417
7.3360
7.3279
7.3194
7.3162
7.3031
5.8615
5.8437
4.5957
4.5778
0.9513
1.0000
5.0417
1.0385
1.0305
[ppm]
[rel]

605

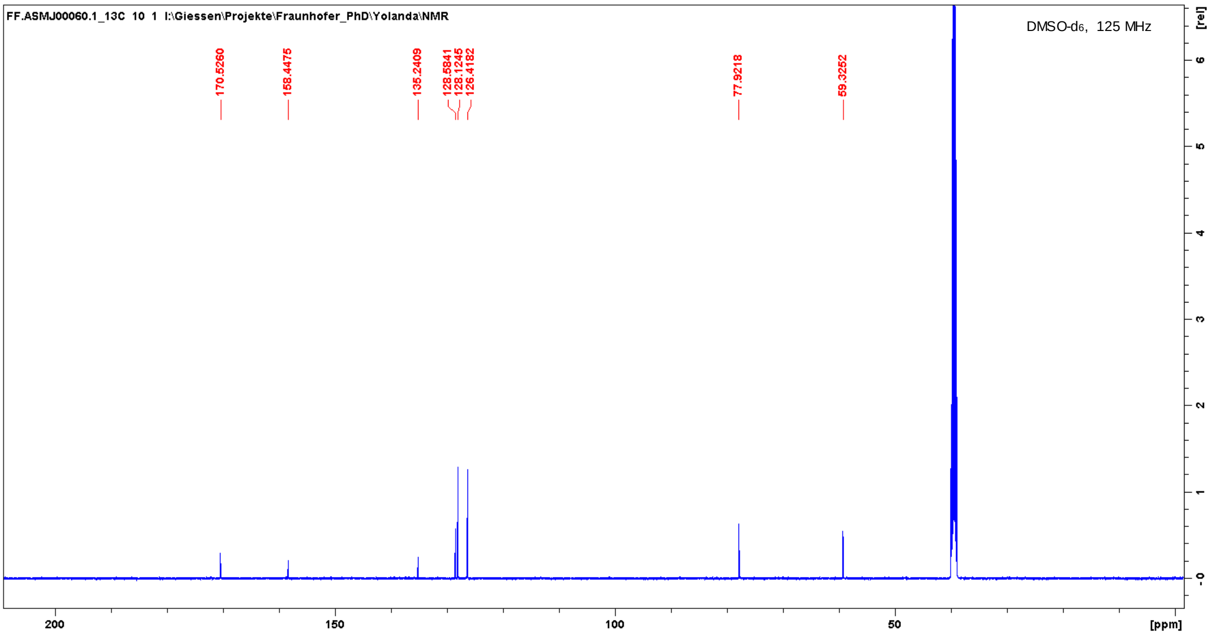
FF.ASMJ00060.1_13C 10 1 I:\Giessen\Projekte\Fraunhofer_PhD\Yolanda\NMR
DMSO-d6, 125 MHz
170.5260
158.4475
135.2409
128.5841
128.1245
126.4182
77.9218
59.3252
[ppm]
[rel]

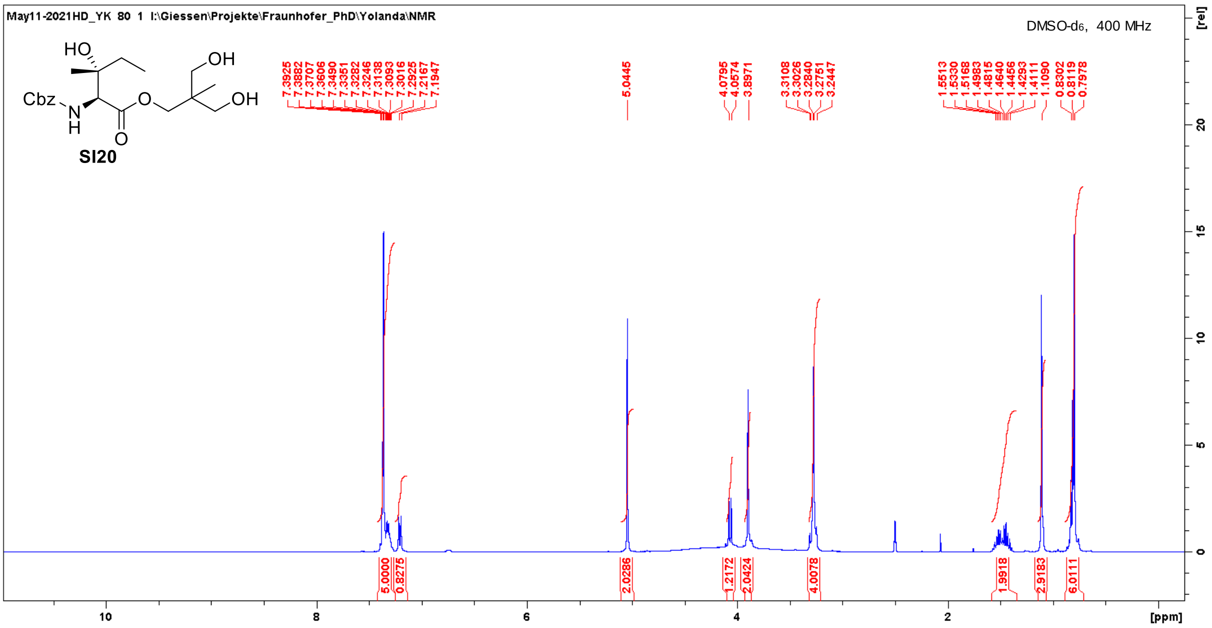
May11-2021HD_YK 80 1 I:\Giessen\Projekte\Fraunhofer_PhD\Yolanda\NMR
DMSO-d6, 400 MHz
HO
OH
Cbz
N
H
O
OH
SI20
7.3925 7.3882 7.3707 7.3606 7.3490 7.3351 7.3282 7.3246 7.3138 7.3093 7.3016 7.2925 7.2167 7.1947
5.0445
4.0795 4.0574 3.8971
3.3108 3.3026 3.2840 3.2751 3.2447
1.5513 1.5330 1.5168 1.4983 1.4815 1.4640 1.4456 1.4293 1.4111 1.1090 0.8302 0.8119 0.7978
5.0000 0.8275 2.0286 1.2172 2.0424 4.0078 1.9918 2.9183 6.0111
[ppm]
[rel]

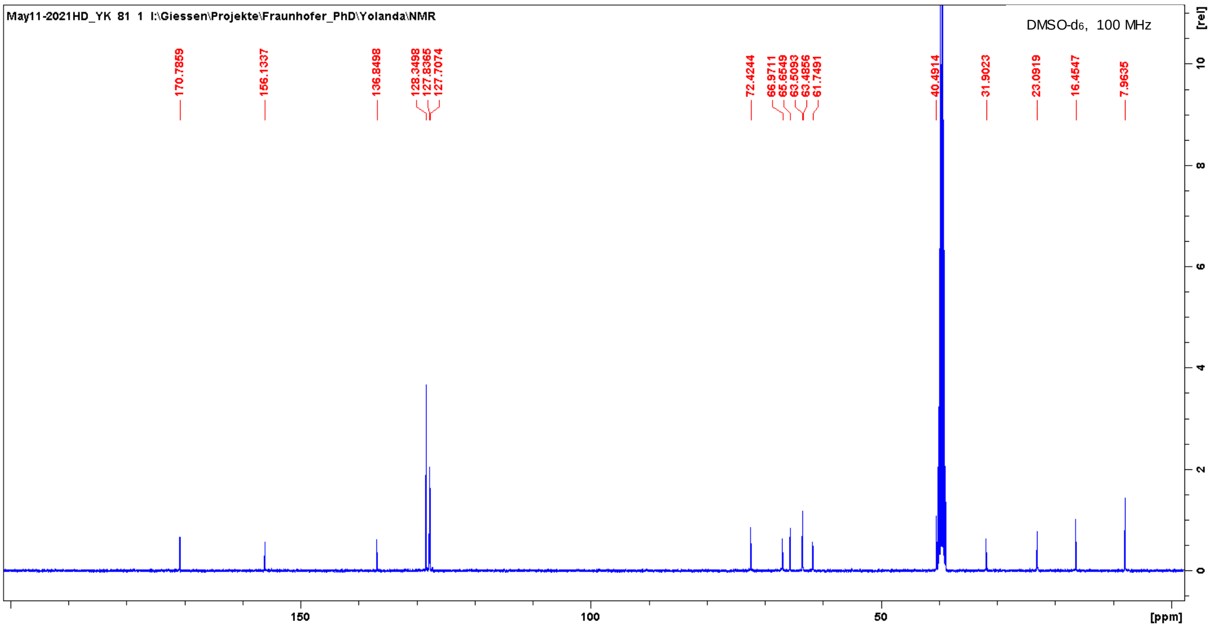
May11-2021HD_YK 81 1 I:\Giessen\Projekte\Fraunhofer_PhD\Yolanda\NMR
DMSO-d6, 100 MHz
170.7859
156.1337
136.8498
128.3488 127.8365 127.7074
72.4244 66.9711 65.6549 63.5093 63.4856 61.7491
40.4914
31.9023
23.0919
16.4547
7.9635
[ppm]
[rel]

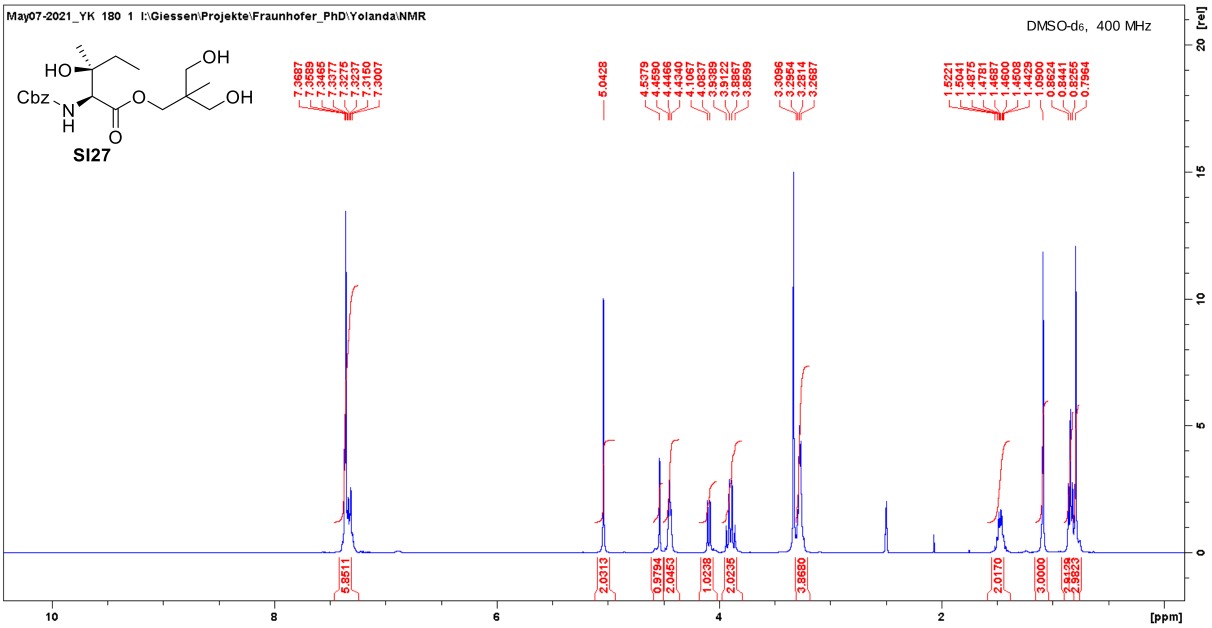
May07-2021_YK 180 1 I:\Giessen\Projekte\Fraunhofer_PhD\Yolanda\NMR
DMSO-d6, 400 MHz
HO
OH
Cbz
N
H
O
OH
SI27
[ppm]
[rel]

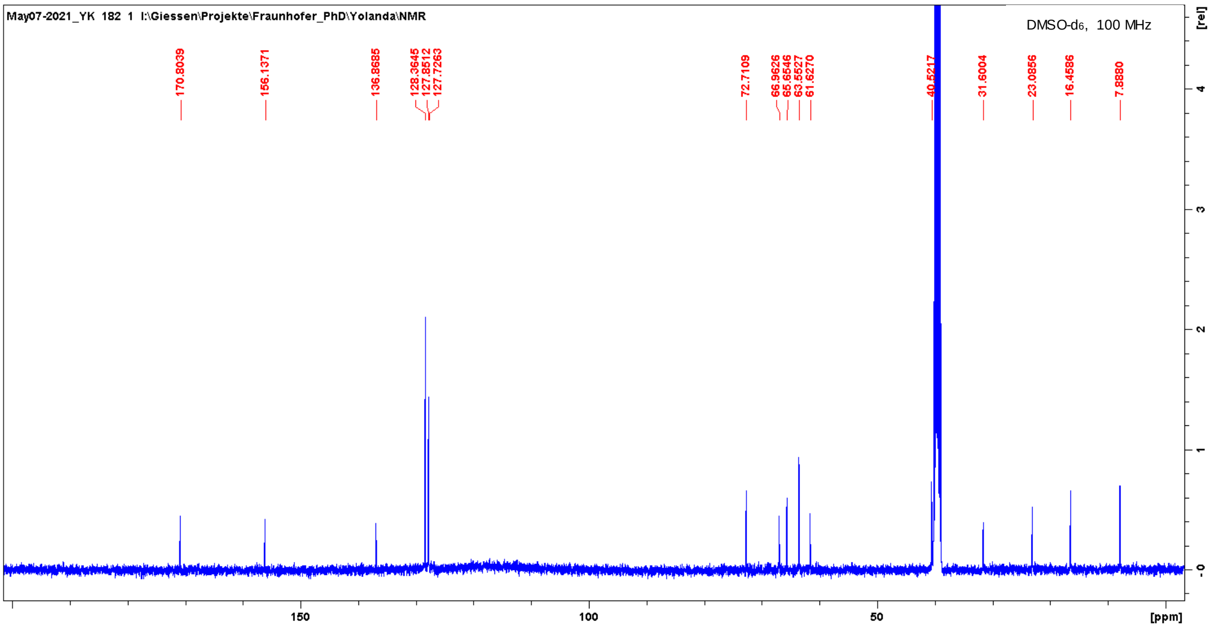
May07-2021_YK 182 1 I:\Giessen\Projekte\Fraunhofer_PhD\Yolanda\NMR
DMSO-d6, 100 MHz
170.8039
156.1371
136.8685
128.3645
127.8512
127.7263
72.7109
66.9626
65.6546
63.5527
61.6270
40.5217
31.6004
23.0856
16.4586
7.8880
[ppm]
[rel]

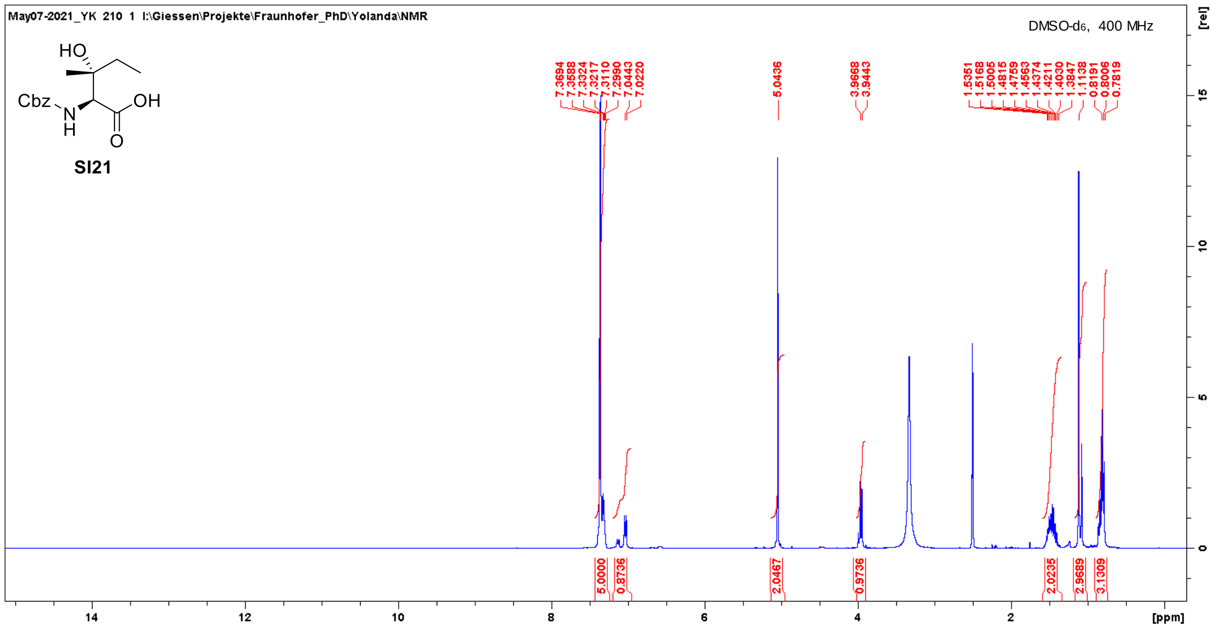
May07-2021_YK 210 1 I:\Giessen\Projekte\Fraunhofer_PhD\Yolanda\NMR
DMSO-d6, 400 MHz
HO
Cbz
N
H
OH
O
SI21
7.3694 7.3588 7.3324 7.3217 7.3110 7.2990 7.0443 7.0220
5.0436
3.9668 3.9443
1.5351 1.5168 1.5005 1.4815 1.4759 1.4563 1.4374 1.4211 1.4030 1.3847 1.1138 0.8191 0.8006 0.7819
5.0000 0.8736 2.0467 0.9736 2.0235 2.9689 3.1309
[ppm]
[rel]

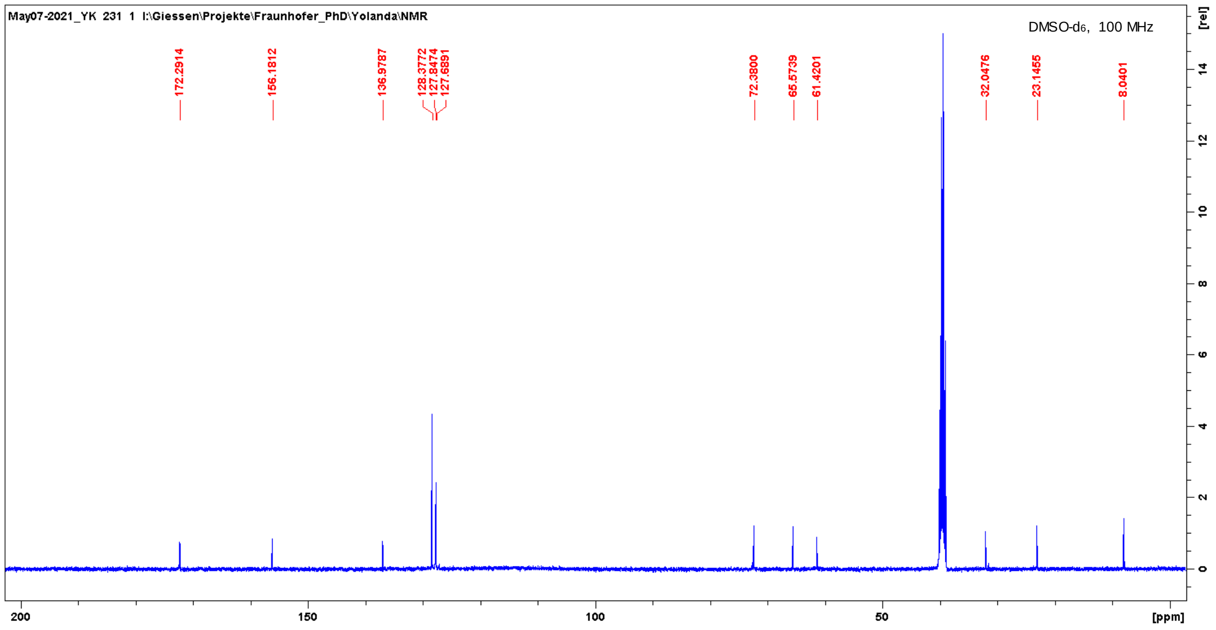
May07-2021_YK 231 1 I:\Giessen\Projekte\Fraunhofer_PhD\Yolanda\NMR
DMSO-d6, 100 MHz
172.2914
156.1812
136.9787
128.3772 127.8474 127.6891
72.3800
65.5739
61.4201
32.0476
23.1455
8.0401
[ppm]
[rel]

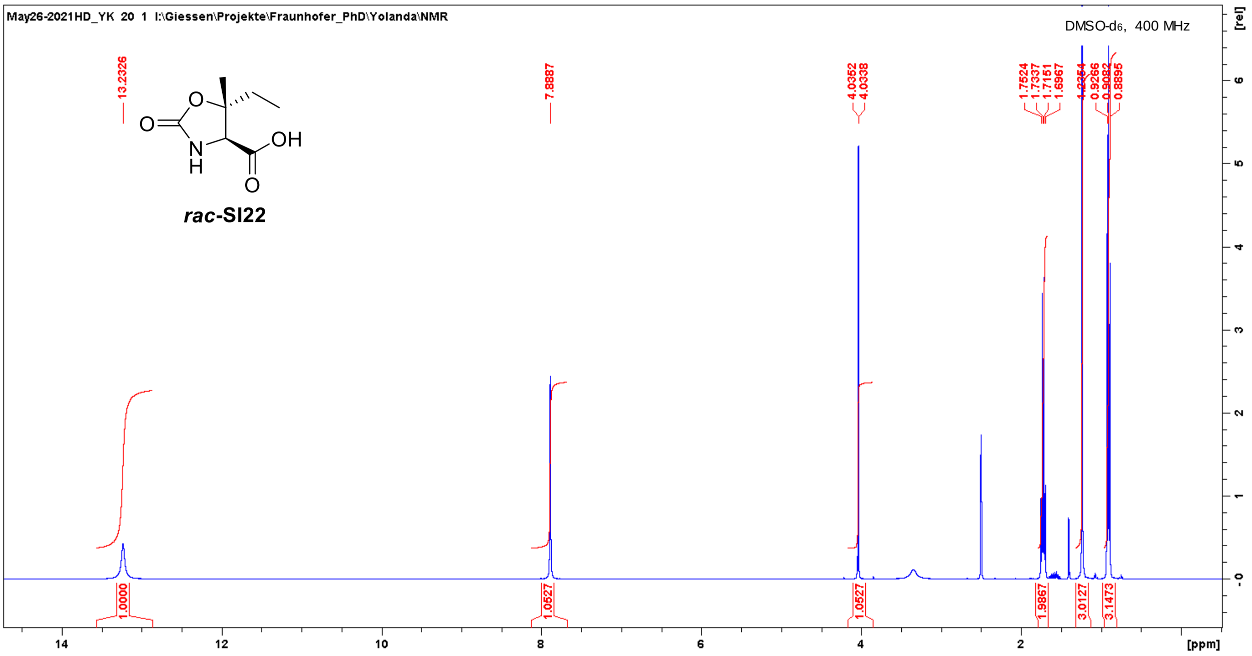
May26-2021HD_YK 20 1 I:\Giessen\Projekte\Fraunhofer_PhD\Yolanda\NMR
DMSO-d6, 400 MHz
rac-SI22
13.2326
7.8887
4.0352
4.0338
1.7524
1.7337
1.7151
1.6967
1.2354
0.9266
0.9082
0.8895
1.0000
1.0527
1.0527
1.9867
3.0127
3.1473
[ppm]
[rel]

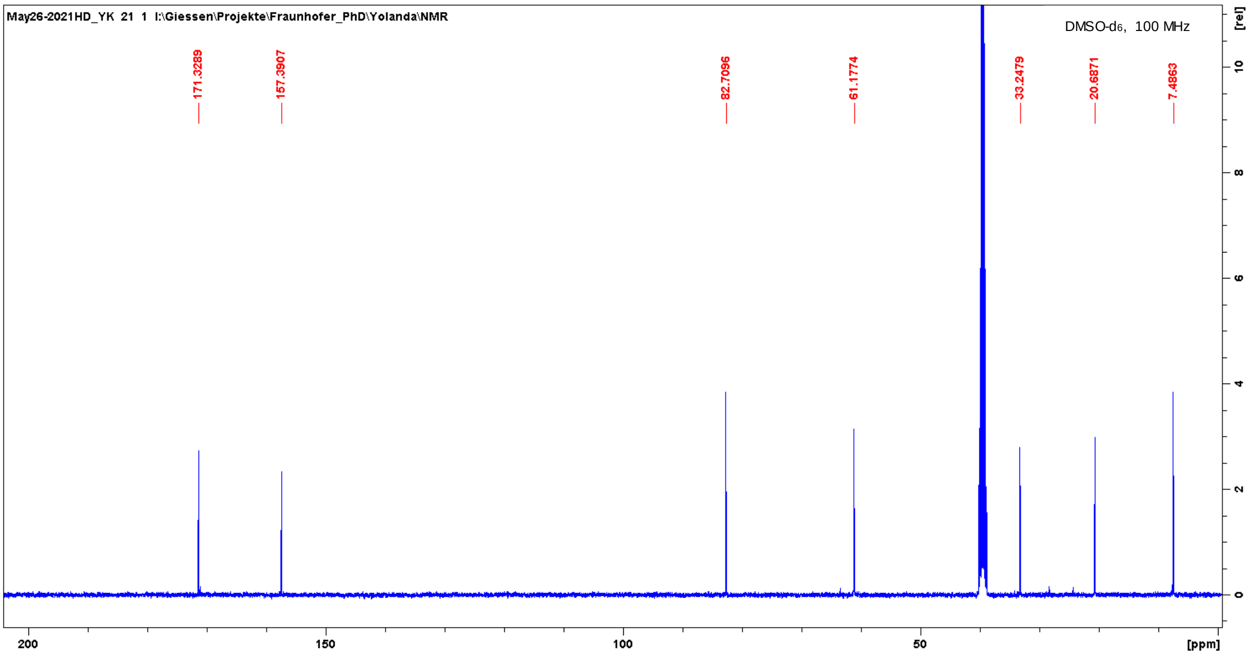
May26-2021HD_YK 21 1 I:\Giessen\Projekte\Fraunhofer_PhD\Yolanda\NMR
DMSO-d6, 100 MHz
171.3289
157.3907
82.7096
61.1774
33.2479
20.6871
7.4863
[ppm]
[rel]

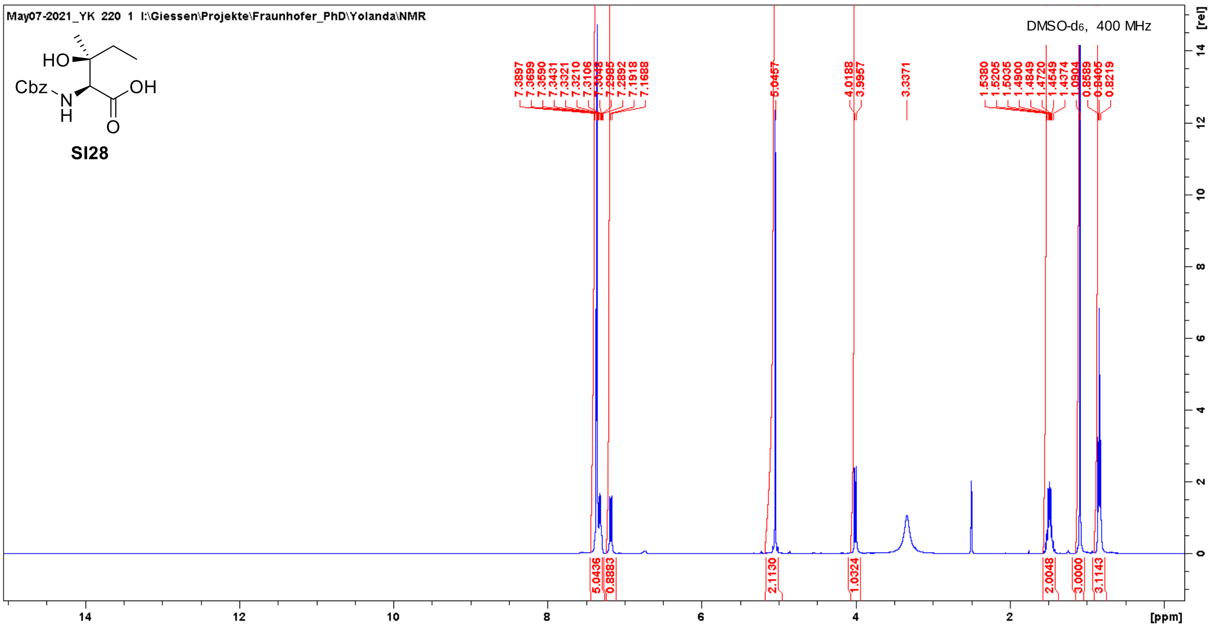
May07-2021_YK 220 1 I:\Giessen\Projekte\Fraunhofer_PhD\Yolanda\NMR
DMSO-d6, 400 MHz
HO
Cbz
N
H
OH
O
SI28
7.3897
7.3699
7.3590
7.3431
7.3321
7.3210
7.3106
7.2985
7.2892
7.1918
7.1688
5.0457
4.0188
3.9957
3.3371
1.5380
1.5205
1.5035
1.4900
1.4849
1.4720
1.4549
1.4374
0.8405
0.8219
5.0436
0.8883
2.1130
1.0324
2.0048
3.0000
3.1143
[ppm]
[rel]

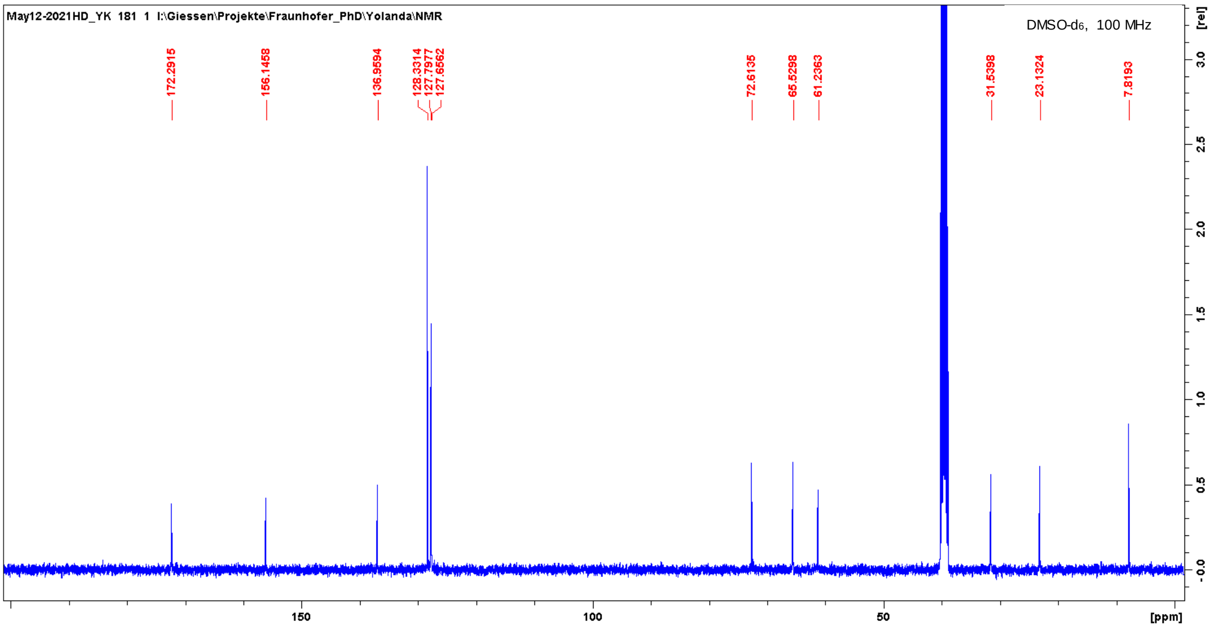
May12-2021HD_YK 181 1 I:\Giessen\Projekte\Fraunhofer_PhD\Yolanda\NMR
DMSO-d6, 100 MHz
172.2915
156.1458
136.9594
128.3314
127.7977
127.6562
72.6135
65.5298
61.2363
31.5398
23.1324
7.8193
[ppm]
[rel]

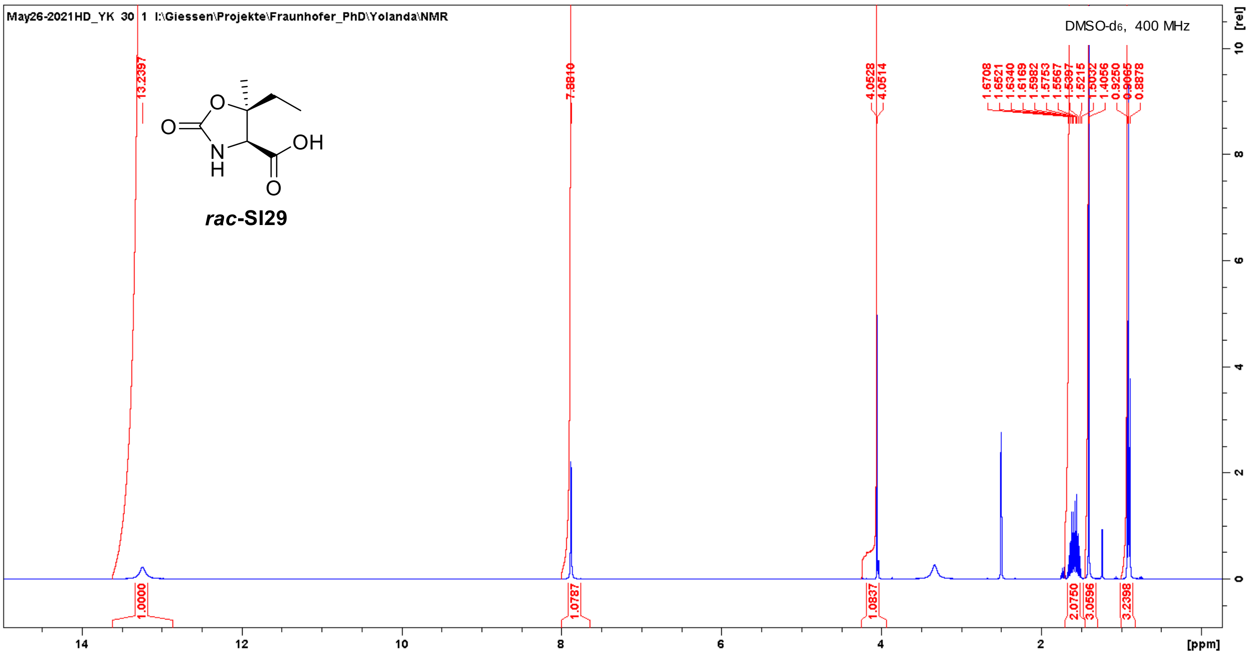
May26-2021HD_YK 30 1 I:\Giessen\Projekte\Fraunhofer_PhD\Yolanda\NMR
DMSO-d6, 400 MHz
rac-SI29
13.2397
7.8810
4.0528
4.0514
1.6708
1.6521
1.6340
1.6169
1.5982
1.5753
1.5567
1.5397
1.5215
1.5032
1.4056
0.9250
0.9065
0.8878
1.0000
1.0787
1.0837
2.0750
3.0596
3.2398
[ppm]
[rel]

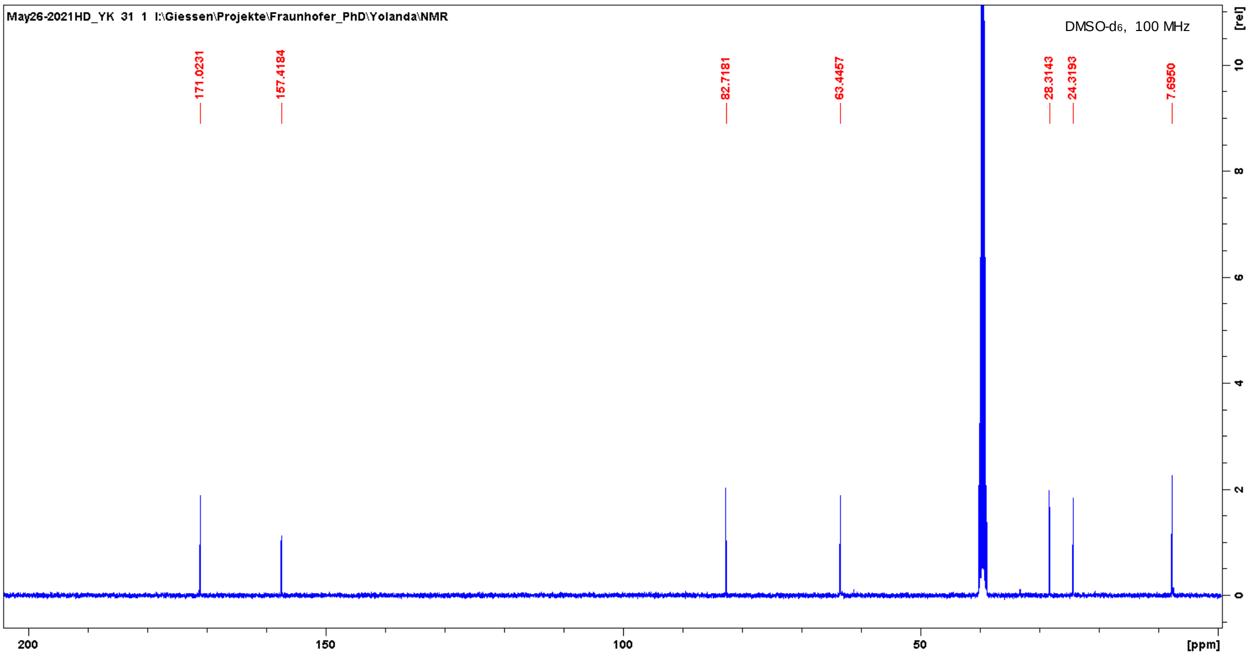
May26-2021HD_YK 31 1 I:\Giessen\Projekte\Fraunhofer_PhD\Yolanda\NMR
DMSO-d6, 100 MHz
171.0231
157.4184
82.7181
63.4457
28.3143
24.3193
7.6950
[ppm]
[rel]

#### 3. Determination of the enantiomeric excesses (*ee*)

##### Chiral Separation

Library, SM, Chiral & Peptide Purification
R&D / IDD In vitro Biology & HT Chemistry
Industriepark Höchst, G 838, Room 007
D-65926 Frankfurt

SANOFI

**Sample Information:**

| | |
|---|---|
| Customer | Dr. Pöverlein |
| Analyst | K. Rahn-Hotze |
| Batch Ref No | FF.ASMJ00012.1 |
| Sample Name | FF.ASMJ00012.1 |
| Racemat | |
| Lab Journal | |
| Comment | |

**Separation Information:**

APC\Poeverlein\Poeverlein_2021

| | |
|---|---|
| HPLC-System | LC_30 |
| Flow rate | 1,0 ml/min |
| Temperature | 30° C |
| HPLC Column | Chiralpak AS-H/122, 250x4,6 mm |
| Eluent | EtOH:MeOH 1:1 |

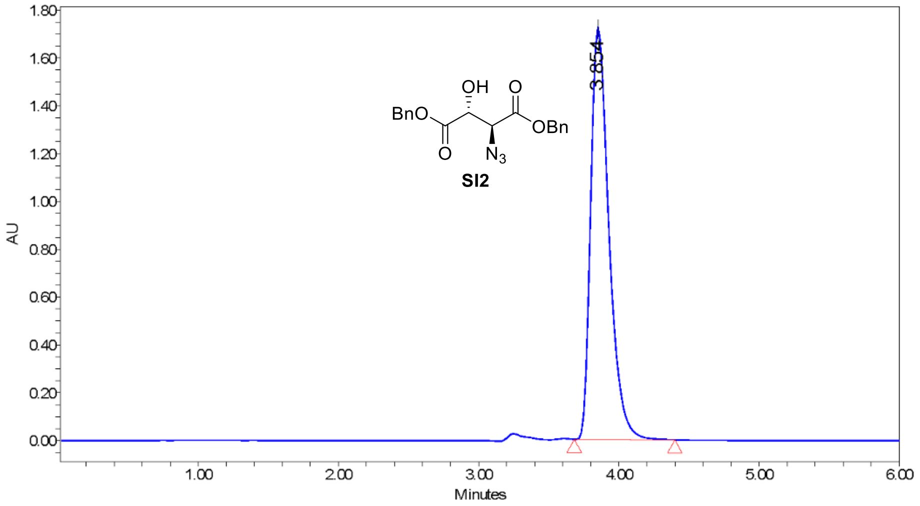

207.0nm; Inj ID 1311; Res ID 1326

Batch_Ref_No FF.ASMJ00012.1 Empower Project: APC\Poeverlein\Poeverlein_2021

**Results**

| | RT | Area | % Area | Height |
|---|---|---|---|---|
| 1 | 3.85 | 15542361 | 100.00 | 1726881 |

Current Date 4/13/2021 Vial 47 1 of 2
Date Acquired 4/13/2021 3:29:30 PM CEST Inj Vol 5.00 uL

#### Sample Information:

Customer Dr. Pöverlein

Batch Ref No FF.ASMJ00012.1

SampleName FF.ASMJ00012.1

Comment

##### Spectrum Index Plot

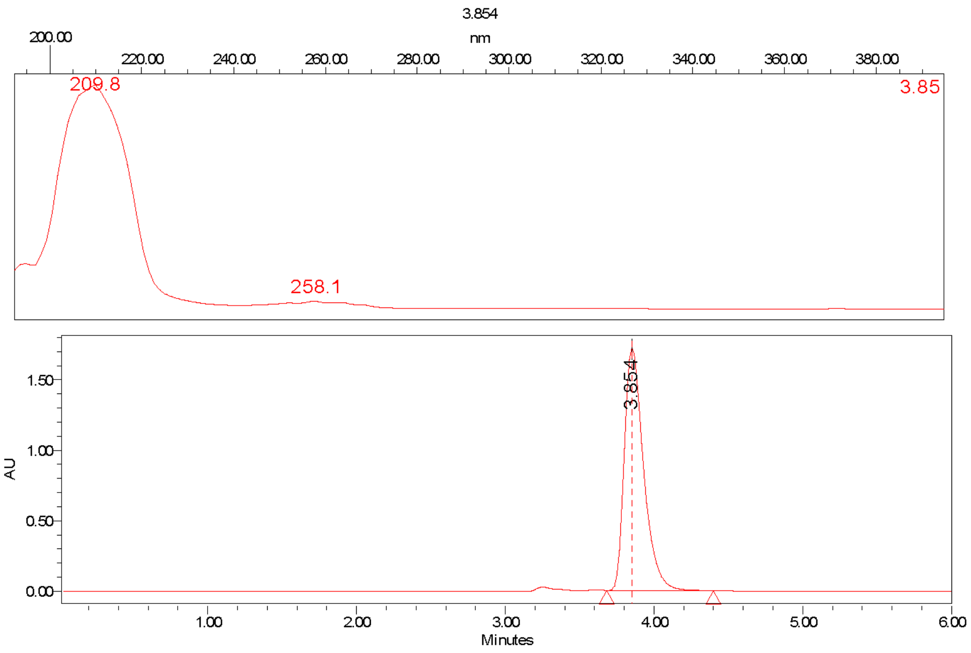

Batch_Ref_No FF.ASMJ00012.1 Project Name: APC\Poeverlein\Poeverlein_2021

Current Date 4/13/2021 Vial 47 2 of 2

Date Acquired 4/13/2021 3:29:30 PM CEST Inj Vol 5.00 uL

626

#### Chiral Separation

Library, SM, Chiral & Peptide Purification
R&D / IDD In vitro Biology & HT Chemistry
Industriepark Höchst, G 838, Room 007
D-65926 Frankfurt

SANOFI

##### Sample Information:

Customer Dr. Pöverlein
Analyst K. Rahn-Hotze
Batch Ref No FF.ASMJ00019.1
Sample Name FF.ASMJ00019.1
Racemat
Lab Journal
Comment

##### Separation Information:

APC\Poeverlein\Poeverlein_2021
HPLC-System LC_30
Flow rate 1,0 ml/min
Temperature 30° C
HPLC Column Chiralpak AS-H/122, 250x4,6 mm
Eluent EtOH:MeOH 1:1

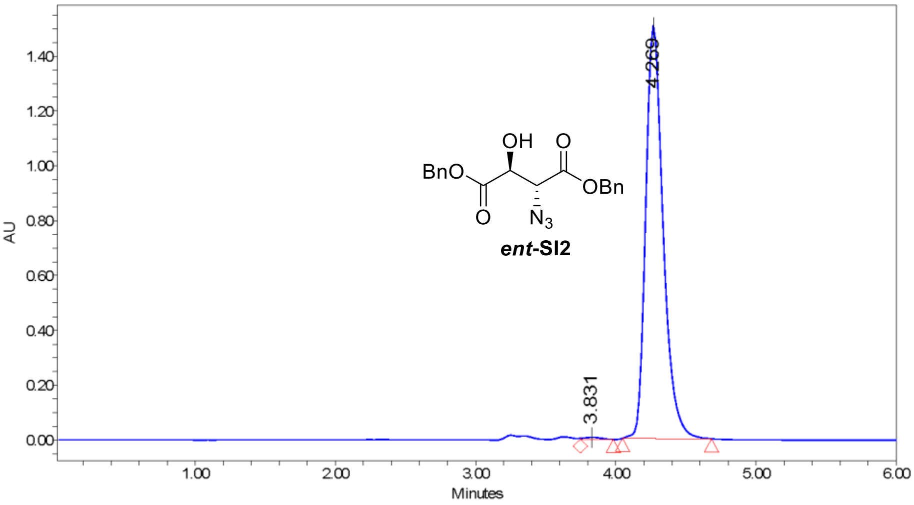

207.0nm; Inj ID 1322; Res ID 1327

Batch_Ref_No FF.ASMJ00019.1 Empower Project: APC\Poeverlein\Poeverlein_2021

##### Results

| | RT | Area | % Area | Height |
|---|---|---|---|---|
| 1 | 3.83 | 65863 | 0.50 | 7974 |
| 2 | 4.27 | 12982699 | 99.50 | 1504163 |

Current Date 4/13/2021 Vial 48 1 of 2
Date Acquired 4/13/2021 3:42:32 PM CEST Inj Vol 2.00 uL

627

#### Sample Information:

Customer Dr. Pöverlein

Batch Ref No FF.ASMJ00019.1

SampleName FF.ASMJ00019.1

Comment

##### Spectrum Index Plot

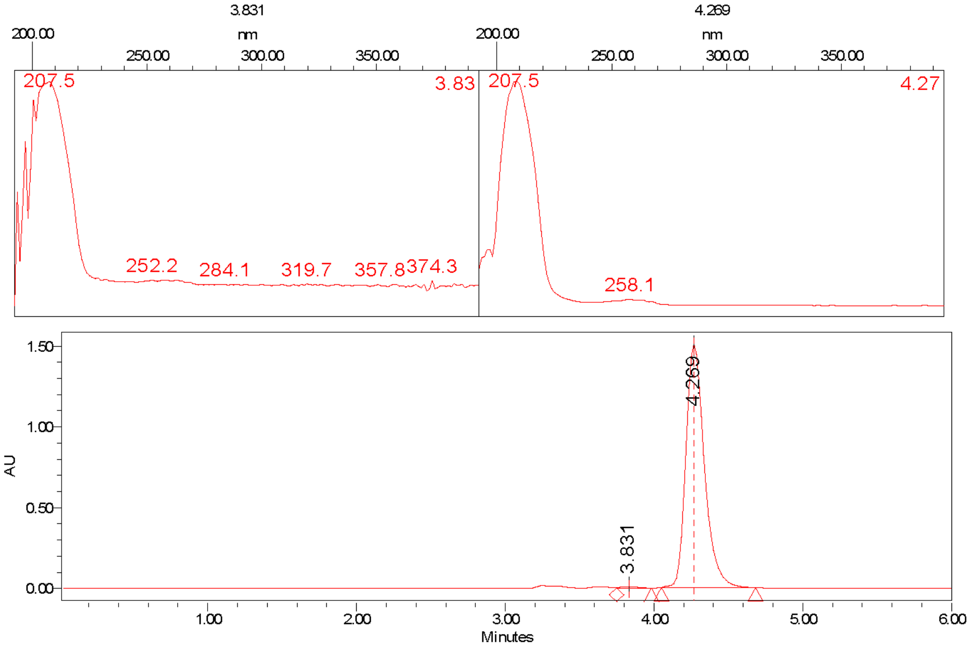

Batch_Ref_No FF.ASMJ00019.1 Project Name: APC\Poeverlein\Poeverlein_2021

Current Date 4/13/2021 Vial 48 2 of 2

Date Acquired 4/13/2021 3:42:32 PM CEST Inj Vol 2.00 uL

628

#### Chiral Separation

Library, SM, Chiral & Peptide Purification
R&D / IDD In vitro Biology & HT Chemistry
Industriepark Höchst, G 838, Room 007
D-65926 Frankfurt

SANOFI

##### Sample Information:

| | |
|---|---|
| Customer | Dr. Pöverlein |
| Analyst | K. Rahn-Hotze |
| Batch Ref No | FF.ASMJ00014.2 |
| Sample Name | FF.ASMJ00014.2 |
| Racemat | |
| Lab Journal | |
| Comment | |

##### Separation Information:

APC\Poeverlein\Poeverlein_2021

| | |
|---|---|
| HPLC-System | LC_30 |
| Flow rate | 1,0 ml/min |
| Temperature | 30° C |
| HPLC Column | Chiralpak AD-H/148, 250x4,6 mm |
| Eluent | EtOH:MeOH 1:1 |

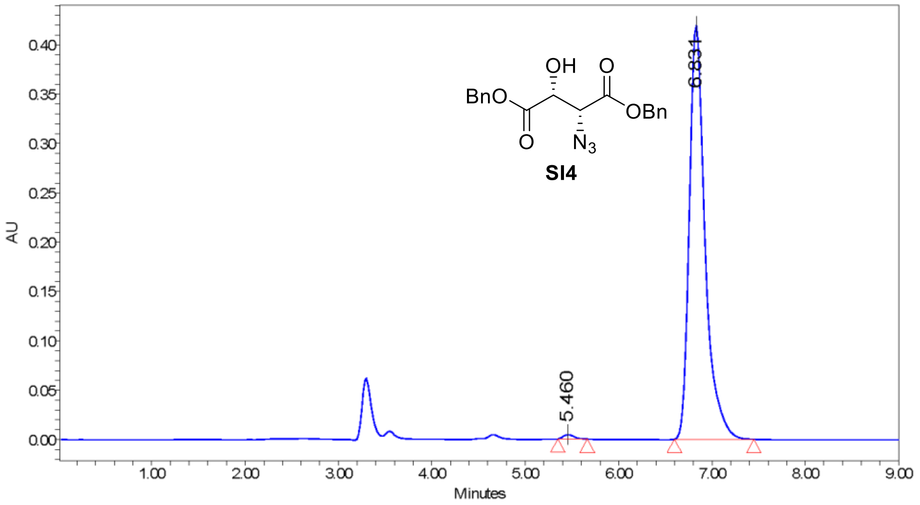

207.0nm; Inj ID 1264; Res ID 1277

Batch_Ref_No FF.ASMJ00014.2 Empower Project: APC\Poeverlein\Poeverlein_2021

##### Results

| | RT | Area | % Area | Height |
|---|---|---|---|---|
| 1 | 5.46 | 34946 | 0.71 | 4388 |
| 2 | 6.83 | 4899448 | 99.29 | 419033 |

Current Date 4/13/2021 Vial 43 1 of 2
Date Acquired 4/13/2021 2:28:50 PM CEST Inj Vol 5.00 uL

629

#### Sample Information:

| | |
|---|---|
| Customer | Dr. Pöverlein |
| Batch Ref No | FF.ASMJ00014.2 |
| SampleName | FF.ASMJ00014.2 |
| Comment | |

#### Spectrum Index Plot

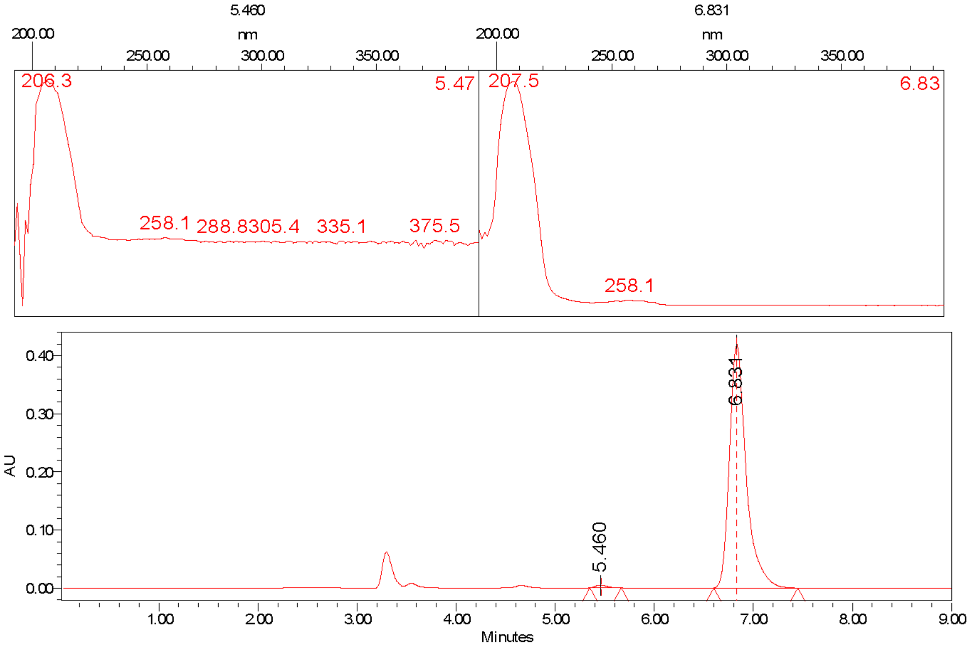

Batch_Ref_No FF.ASMJ00014.2 Project Name: APC\Poeverlein\Poeverlein_2021

### Chiral Separation

Library, SM, Chiral & Peptide Purification
R&D / IDD In vitro Biology & HT Chemistry
Industriepark Höchst, G 838, Room 007
D-65926 Frankfurt

SANOFI

#### Sample Information:

| | |
|---|---|
| Customer | Dr. Pöverlein |
| Analyst | K. Rahn-Hotze |
| Batch Ref No | FF.ASMJ00021.2 |
| Sample Name | FF.ASMJ00021.2 |
| Racemat | |
| Lab Journal | |
| Comment | |

#### Separation Information:

APC\Poeverlein\Poeverlein_2021

| | |
|---|---|
| HPLC-System | LC_30 |
| Flow rate | 1,0 ml/min |
| Temperature | 30° C |
| HPLC Column | Chiralpak AD-H/148, 250x4,6 mm |
| Eluent | EtOH:MeOH 1:1 |

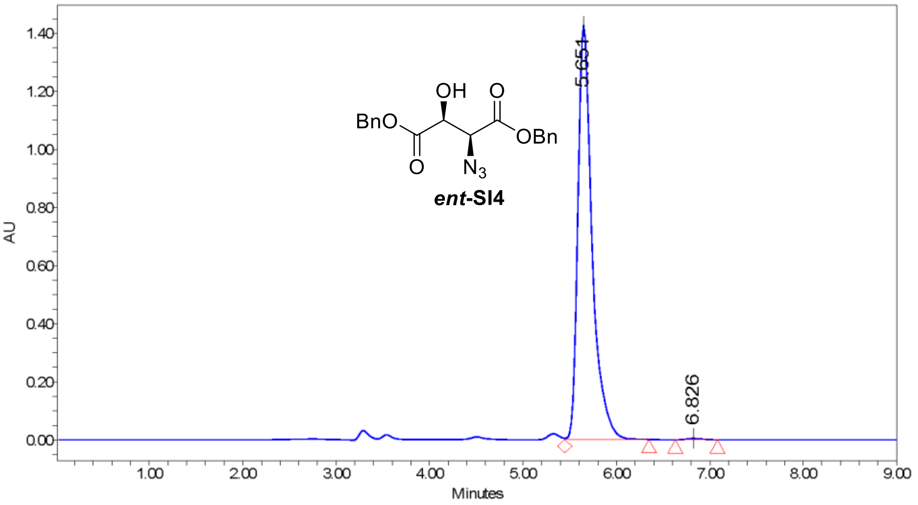

Batch_Ref_No FF.ASMJ00021.2    Empower Project: APC\Poeverlein\Poeverlein_2021

##### Results

| | RT | Area | % Area | Height |
|---|---|---|---|---|
| 1 | 5.65 | 15082936 | 99.67 | 1423205 |
| 2 | 6.83 | 50343 | 0.33 | 4658 |

Current Date 4/13/2021    Vial 44    1 of 2
Date Acquired 4/13/2021 2:38:46 PM CEST    Inj Vol 5.00 uL

631

#### Sample Information:

Customer Dr. Pöverlein

Batch Ref No FF.ASMJ00021.2

SampleName FF.ASMJ00021.2

Comment

**Spectrum Index Plot**

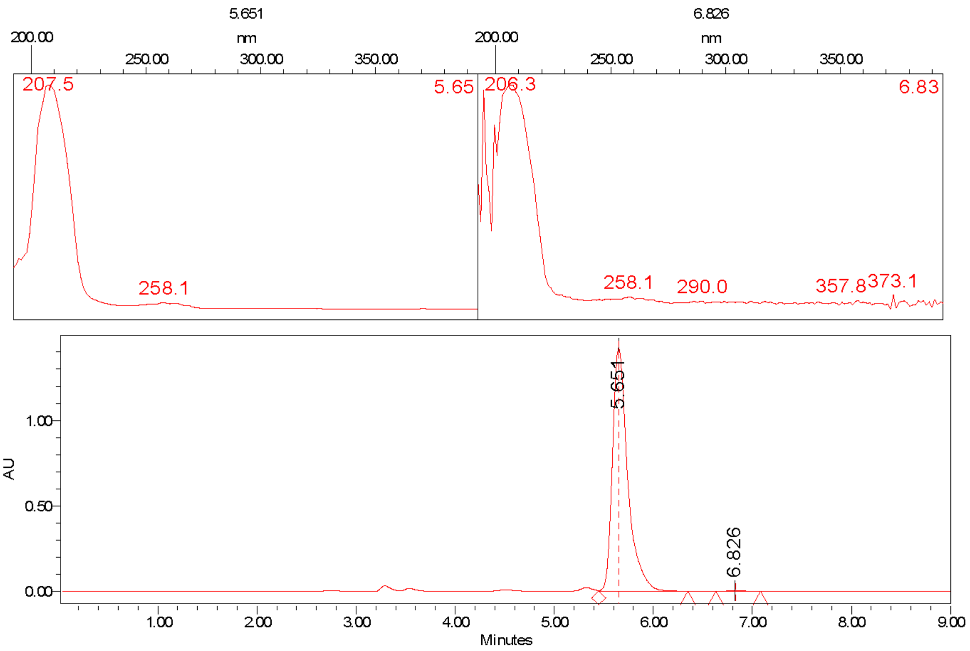

Batch_Ref_No FF.ASMJ00021.2 Project Name: APC\Poeverlein\Poeverlein_2021

| | | | | |
|---|---|---|---|---|
| Current Date | 4/13/2021 | Vial | 44 | 2 of 2 |
| Date Acquired | 4/13/2021 2:38:46 PM CEST | Inj Vol | 5.00 uL | |

#### Chiral Separation

Library, SM, Chiral & Peptide Purification
R&D / IDD In vitro Biology & HT Chemistry
Industriepark Höchst, G 838, Room 007
D-65926 Frankfurt

SANOFI

#### Sample Information:

| | |
|---|---|
| Customer | Dr. Pöverlein |
| Analyst | K. Rahn-Hotze |
| Batch Ref No | FF.ASMJ00075.2 |
| Sample Name | FF.ASMJ00075.2 |
| Racemat | |
| Lab Journal | |
| Comment | |

#### Separation Information:

APC\Poeverlein\Poeverlein_2021

| | |
|---|---|
| HPLC-System | LC_03 |
| Flow rate | 1,0 ml/min |
| Temperature | 30° C |
| HPLC Column | Chiralpak ID/174, 250x4,6 mm |
| Eluent | Hep:EtOH:MeOH 5:1:1 + 0,1% TFA |

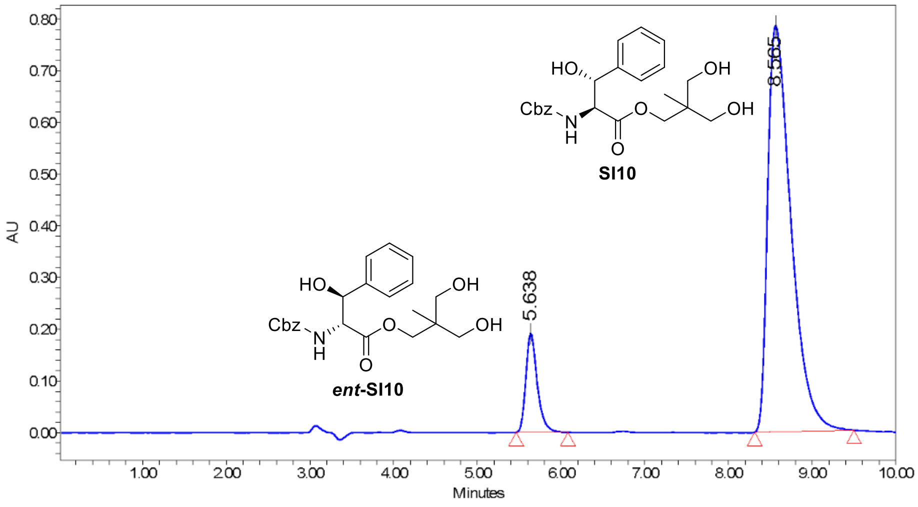

205.0nm; Inj ID 1607; Res ID 1695

Batch_Ref_No FF.ASMJ00075.2 Empower Project: APC\Poeverlein\Poeverlein_2021

##### Results

| | RT | Area | % Area | Height |
|---|---|---|---|---|
| 1 | 5.64 | 1817870 | 10.68 | 191839 |
| 2 | 8.56 | 15197037 | 89.32 | 785502 |

Current Date 5/19/2021 Vial 32
Date Acquired 5/18/2021 3:34:03 PM CEST Inj Vol 2.00 uL

634

#### Sample Information:

Customer Dr. Pöverlein

Batch Ref No FF.ASMJ00075.2

SampleName FF.ASMJ00075.2

Comment

##### Spectrum Index Plot

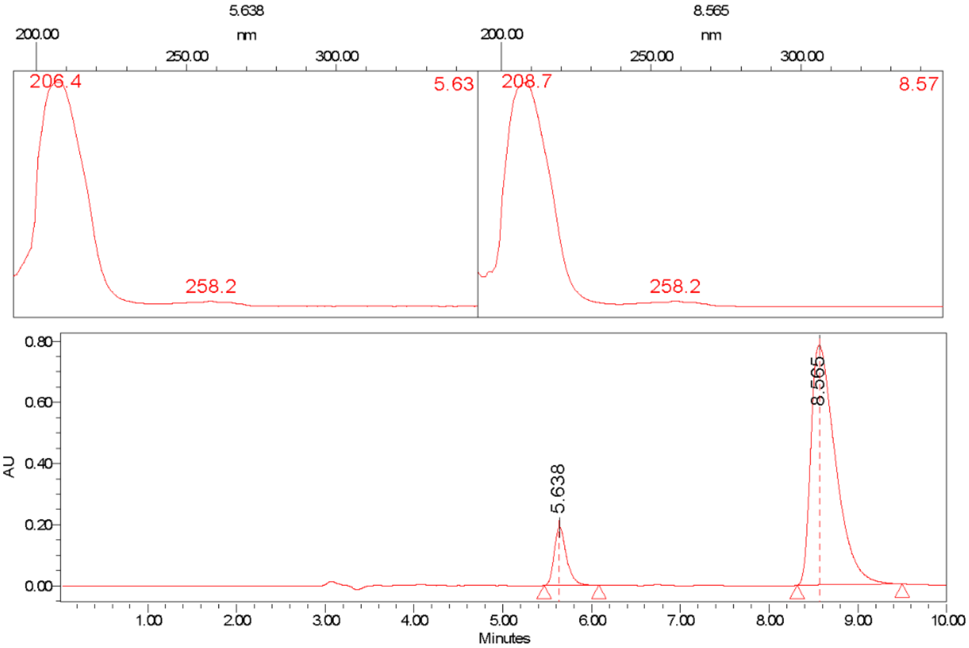

Batch_Ref_No FF.ASMJ00075.2 Project Name: APC\Poeverlein\Poeverlein_2021

#### Chiral Separation

Library, SM, Chiral & Peptide Purification
R&D / IDD In vitro Biology & HT Chemistry
Industriepark Höchst, G 838, Room 007
D-65926 Frankfurt

SANOFI

#### Sample Information:

Customer Dr. Pöverlein
Analyst K. Rahn-Hotze
Batch Ref No FF.ASMJ00072.2
Sample Name FF.ASMJ00072.2
Racemat
Lab Journal
Comment

#### Separation Information:

APC\Poeverlein\Poeverlein_2021
HPLC-System LC_03
Flow rate 1,0 ml/min
Temperature 30° C
HPLC Column Chiralpak ID/174, 250x4,6 mm
Eluent Hep:EtOH:MeOH 5:1:1 + 0,1% TFA

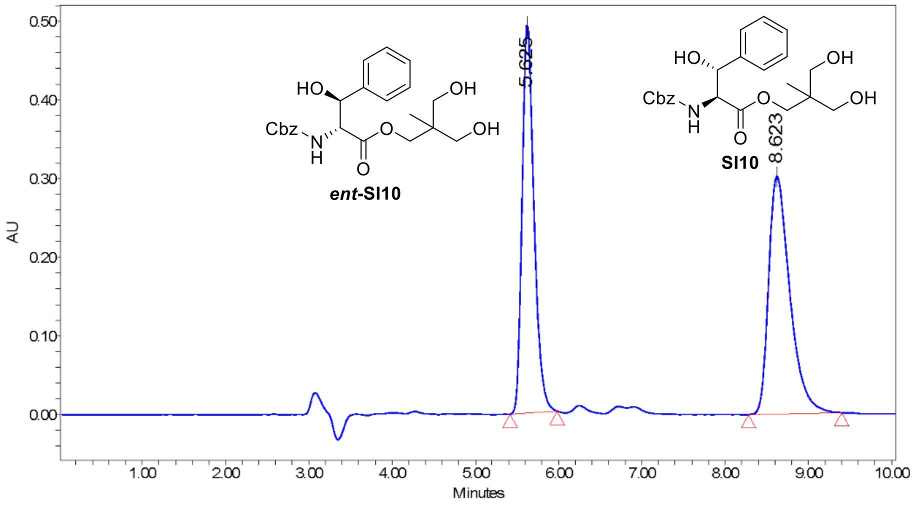

205.0nm; Inj ID 1561; Res ID 1568

Batch_Ref_No FF.ASMJ00072.2 Empower Project: APC\Poeverlein\Poeverlein_2021

##### Results

| | RT | Area | % Area | Height |
|---|---|---|---|---|
| 1 | 5.62 | 4892494 | 47.71 | 496270 |
| 2 | 8.62 | 5362392 | 52.29 | 302904 |

Current Date 5/19/2021 Vial 26 
Date Acquired 5/18/2021 10:18:44 AM CEST Inj Vol 5.00 uL

636

#### Sample Information:

| | |
|---|---|
| Customer | Dr. Pöverlein |
| Batch Ref No | FF.ASMJ00072.2 |
| SampleName | FF.ASMJ00072.2 |
| Comment | |

##### Spectrum Index Plot

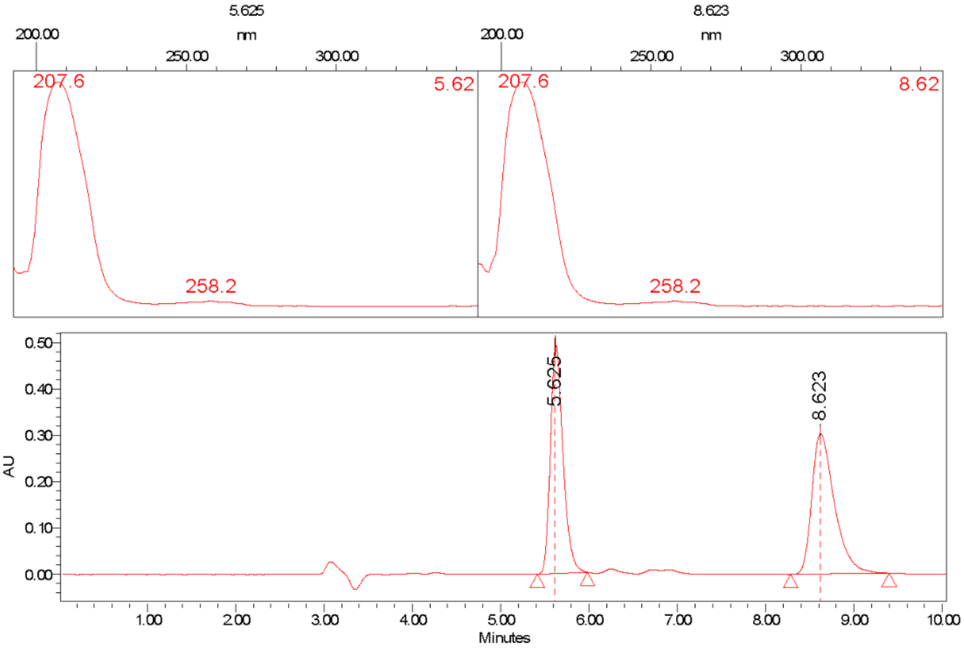

Batch_Ref_No FF.ASMJ00072.2 Project Name: APC\Poeverlein\Poeverlein_2021

Current Date 5/19/2021 Vial 26 2 of 2
Date Acquired 5/18/2021 10:18:44 AM Inj Vol 5.00 uL
CEST

637

#### Chiral Separation

Library, SM, Chiral & Peptide Purification
R&D / IDD In vitro Biology & HT Chemistry
Industriepark Höchst, G 838, Room 007
D-65926 Frankfurt

SANOFI

#### Sample Information:

Customer Dr. Pöverlein
Analyst K. Rahn-Hotze
Batch Ref No FF.ASMJ00075.1
Sample Name FF.ASMJ00075.1
Racemat
Lab Journal
Comment

#### Separation Information:

APC\Poeverlein\Poeverlein_2021
HPLC-System LC_03
Flow rate 1,0 ml/min
Temperature 30° C
HPLC Column Chiralpak IF/181, 250x4,6 mm
Eluent Hep:EtOH:MeOH 5:1:1 + 0,1% TFA

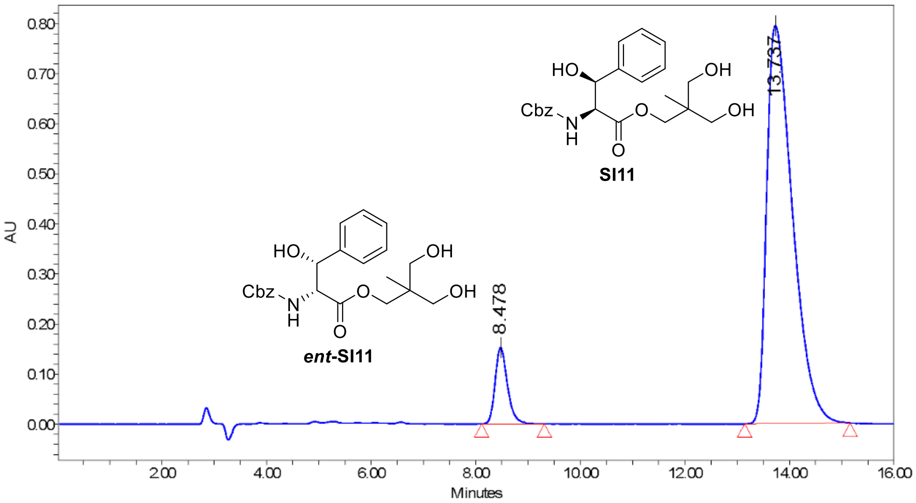

Batch_Ref_No FF.ASMJ00075.1 Empower Project: APC\Poeverlein\Poeverlein_2021

##### Results

| | RT | Area | % Area | Height |
|---|---|---|---|---|
| 1 | 8.48 | 2488866 | 8.38 | 152577 |
| 2 | 13.74 | 27228681 | 91.62 | 794500 |

Current Date 5/19/2021
Date Acquired 5/18/2021 2:30:24 PM CEST
Vial 29
Inj Vol 5.00 uL

638

#### Sample Information:

| | |
|---|---|
| Customer | Dr. Pöverlein |
| Batch Ref No | FF.ASMJ00075.1 |
| SampleName | FF.ASMJ00075.1 |
| Comment | |

##### Spectrum Index Plot

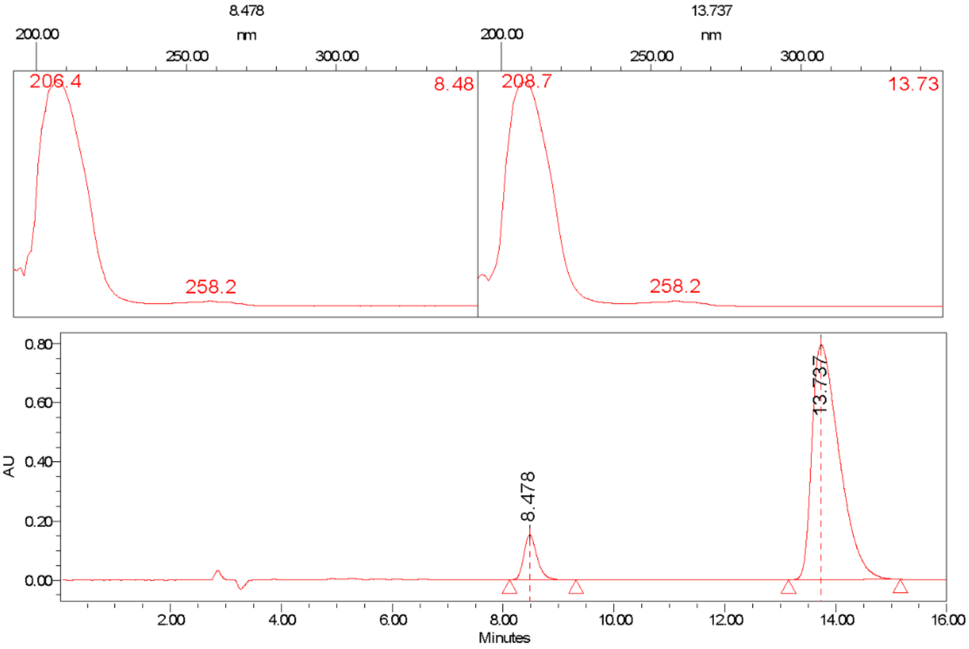

Batch_Ref_No FF.ASMJ00075.1 Project Name: APC\Poeverlein\Poeverlein_2021

#### Chiral Separation

Library, SM, Chiral & Peptide Purification
R&D / IDD In vitro Biology & HT Chemistry
Industriepark Höchst, G 838, Room 007
D-65926 Frankfurt

SANOFI

#### Sample Information:

Customer  Dr. Pöverlein
Analyst  K. Rahn-Hotze
Batch Ref No  FF.ASMJ00072.1
Sample Name  FF.ASMJ00072.1
Racemat
Lab Journal
Comment

#### Separation Information:

APC\Poeverlein\Poeverlein_2021
HPLC-System  LC_03
Flow rate  1,0 ml/min
Temperature  30° C
HPLC Column  Chiralpak IF/181, 250x4,6 mm
Eluent  Hep:EtOH:MeOH 5:1:1 + 0,1% TFA

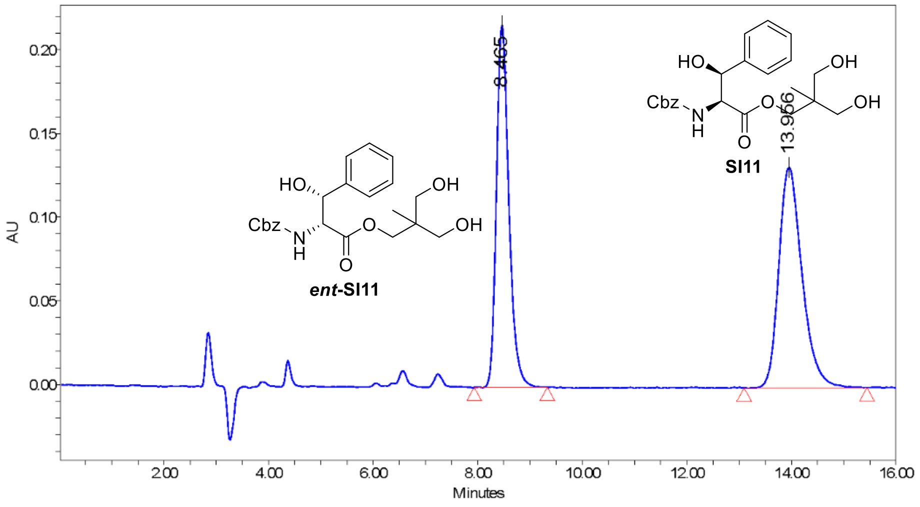

Batch_Ref_No FF.ASMJ00072.1  Empower Project: APC\Poeverlein\Poeverlein_2021

##### Results

| | RT | Area | % Area | Height |
|---|---|---|---|---|
| 1 | 8.46 | 3565304 | 46.77 | 216120 |
| 2 | 13.96 | 4057665 | 53.23 | 131638 |

Current Date  5/19/2021  Vial  25  1 of 2
Date Acquired  5/18/2021 11:04:49 AM CEST  Inj Vol  5.00 uL

640

#### Sample Information:

Customer Dr. Pöverlein
Batch Ref No FF.ASMJ00072.1
SampleName FF.ASMJ00072.1
Comment

##### Spectrum Index Plot

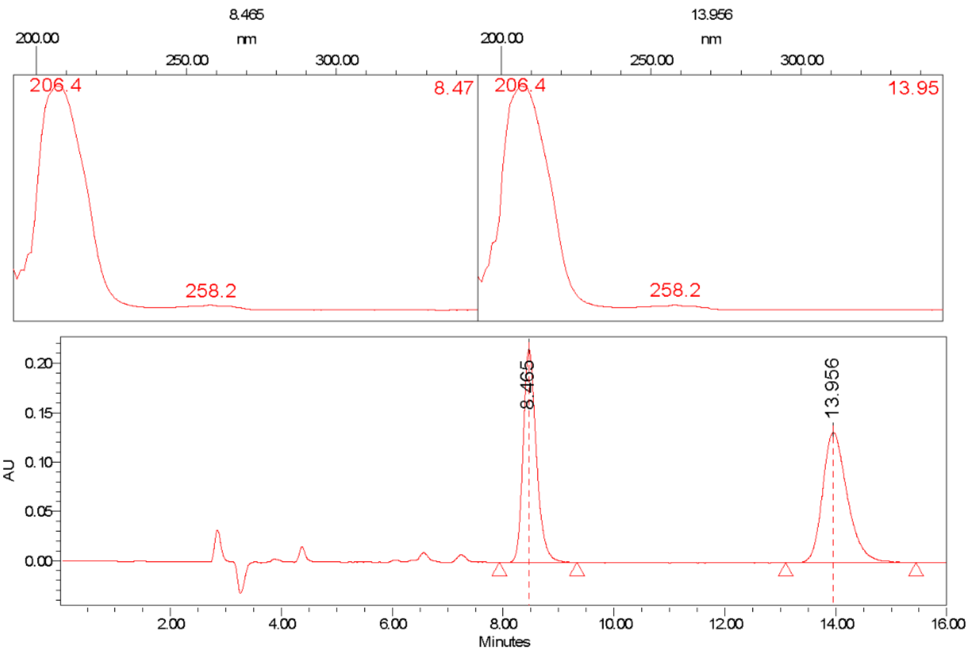

Batch_Ref_No FF.ASMJ00072.1 Project Name: APC\Poeverlein\Poeverlein_2021

Current Date 5/19/2021 Vial 25 2 of 2
Date Acquired 5/18/2021 11:04:49 AM CEST Inj Vol 5.00 uL

641

#### Chiral Separation

Library, SM, Chiral & Peptide Purification
R&D / IDD In vitro Biology & HT Chemistry
Industriepark Höchst, G 838, Room 007
D-65926 Frankfurt

SANOFI

#### Sample Information:

| | |
|---|---|
| Customer | Dr. Pöverlein |
| Analyst | K. Rahn-Hotze |
| Batch Ref No | FF.ASMJ00095.1 |
| Sample Name | FF.ASMJ00095.1 |
| Racemat | |
| Lab Journal | |
| Comment | |

#### Separation Information:

APC\Poeverlein\Poeverlein_2021

| | |
|---|---|
| HPLC-System | LC_03 |
| Flow rate | 1,0 ml/min |
| Temperature | 30° C |
| HPLC Column | Chiralpak IF/181, 250x4,6 mm |
| Eluent | Hep:EtOH:MeOH 5:1:1 + 0,1% TFA |

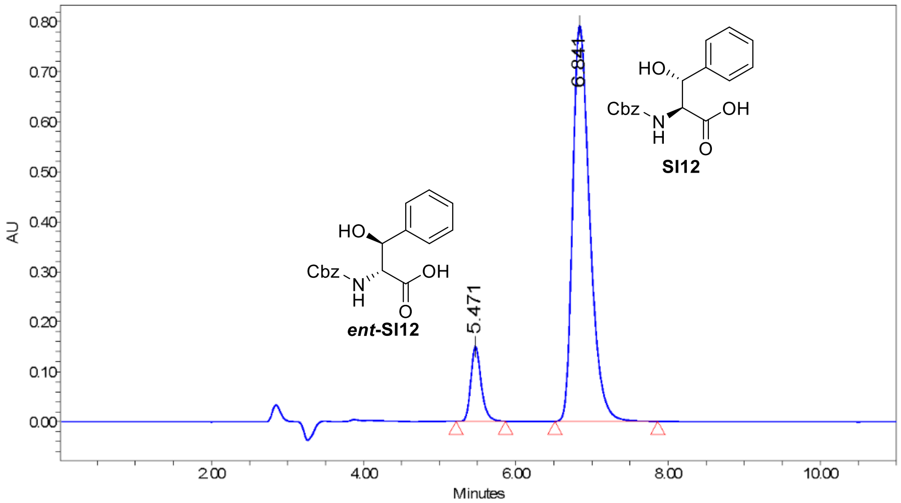

Batch_Ref_No FF.ASMJ00095.1 Empower Project: APC\Poeverlein\Poeverlein_2021

##### Results

| | RT | Area | % Area | Height |
|---|---|---|---|---|
| 1 | 5.47 | 1415425 | 10.28 | 149772 |
| 2 | 6.84 | 12350954 | 89.72 | 793556 |

Current Date 5/19/2021 Vial 31 1 of 2
Date Acquired 5/18/2021 2:57:07 PM CEST Inj Vol 5.00 uL

642

#### Sample Information:

Customer Dr. Pöverlein
Batch Ref No FF.ASMJ00095.1
SampleName FF.ASMJ00095.1
Comment

##### Spectrum Index Plot

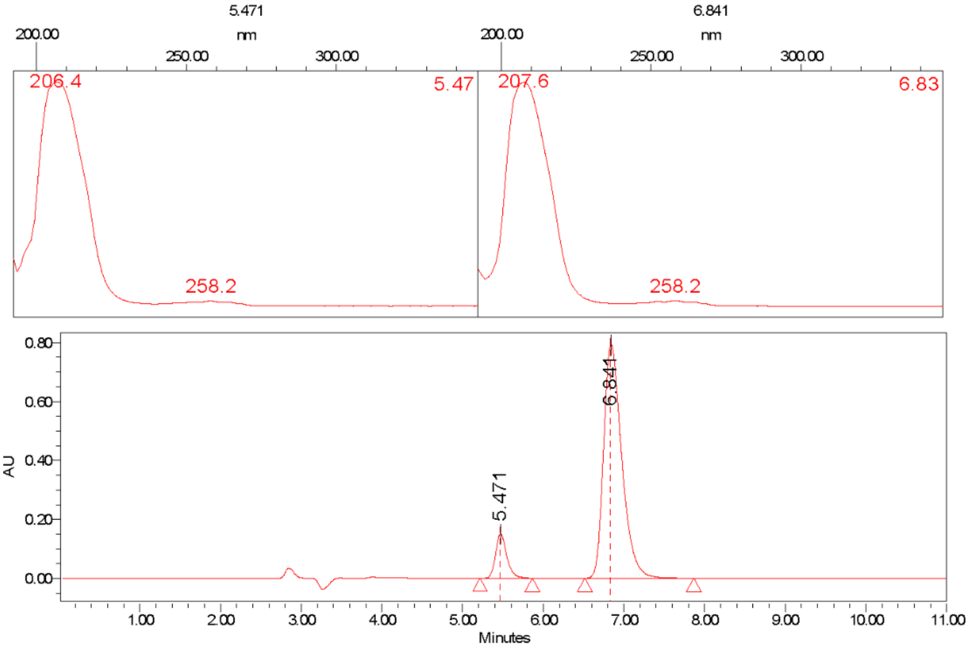

Batch_Ref_No FF.ASMJ00095.1 Project Name: APC\Poeverlein\Poeverlein_2021

#### Chiral Separation

Library, SM, Chiral & Peptide Purification
R&D / IDD In vitro Biology & HT Chemistry
Industriepark Höchst, G 838, Room 007
D-65926 Frankfurt

SANOFI

#### Sample Information:

| | |
|---|---|
| Customer | Dr. Pöverlein |
| Analyst | K. Rahn-Hotze |
| Batch Ref No | FF.ASMJ00091.1 |
| Sample Name | FF.ASMJ00091.1 |
| Racemat | |
| Lab Journal | |
| Comment | |

#### Separation Information:

APC\Poeverlein\Poeverlein_2021

| | |
|---|---|
| HPLC-System | LC_03 |
| Flow rate | 1,0 ml/min |
| Temperature | 30° C |
| HPLC Column | Chiralpak IF/181, 250x4,6 mm |
| Eluent | Hep:EtOH:MeOH 5:1:1 + 0,1% TFA |

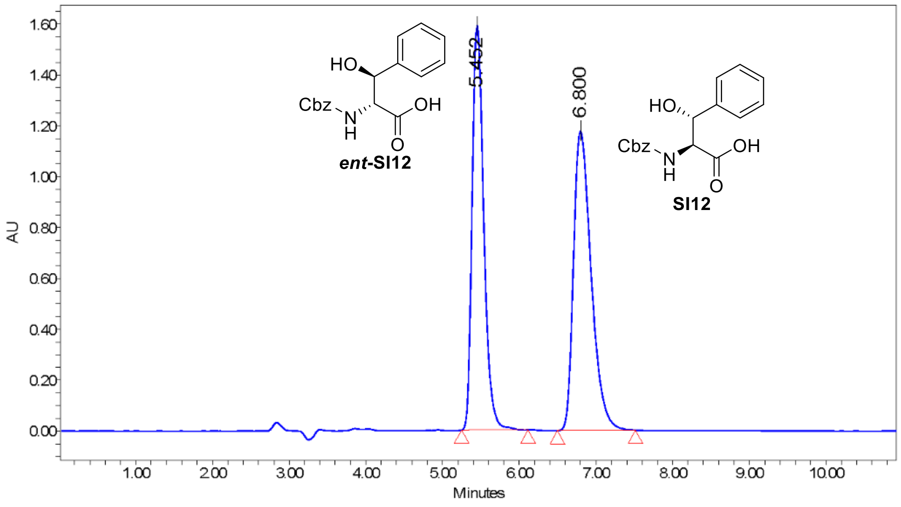

Batch_Ref_No FF.ASMJ00091.1 Empower Project: APC\Poeverlein\Poeverlein_2021

##### Results

| | RT | Area | % Area | Height |
|---|---|---|---|---|
| 1 | 5.45 | 16852027 | 47.19 | 1590853 |
| 2 | 6.80 | 18856420 | 52.81 | 1178025 |

Current Date 5/19/2021
Date Acquired 5/18/2021 9:54:32 AM CEST
Vial 28
Inj Vol 5.00 uL

644

#### Sample Information:

Customer Dr. Pöverlein

Batch Ref No FF.ASMJ00091.1

SampleName FF.ASMJ00091.1

Comment

**Spectrum Index Plot**

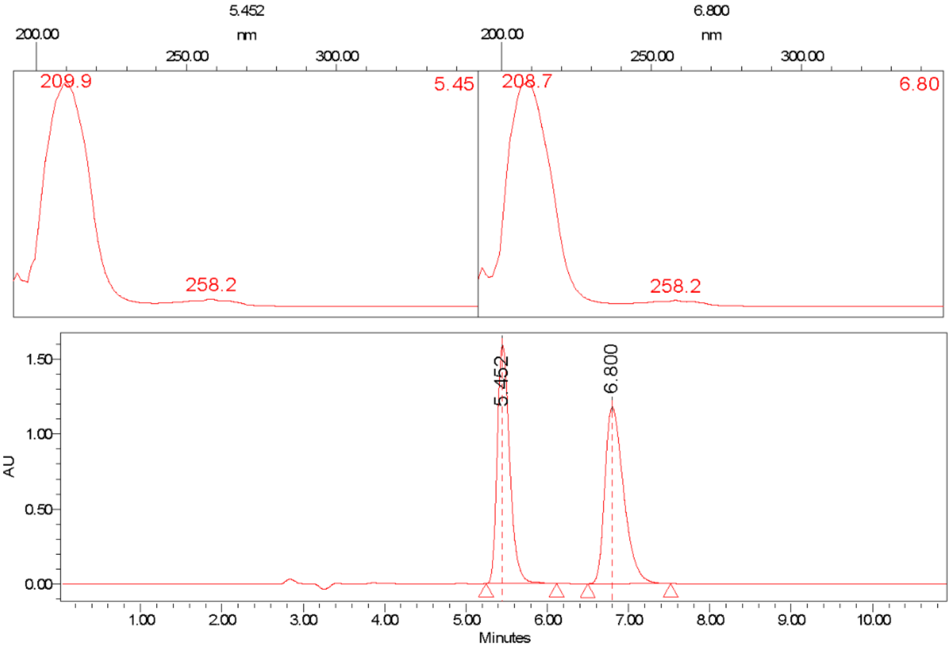

Batch_Ref_No FF.ASMJ00091.1 Project Name: APC\Poeverlein\Poeverlein_2021

Current Date 5/19/2021 Vial 28 2 of 2

Date Acquired 5/18/2021 9:54:32 AM CEST Inj Vol 5.00 uL

645

646

#### Chiral Separation

Library, SM, Chiral & Peptide Purification
R&D / IDD In vitro Biology & HT Chemistry
Industriepark Höchst, G 838, Room 007
D-65926 Frankfurt

SANOFI

#### Sample Information:

Customer Dr. Pöverlein
Analyst K. Rahn-Hotze
Batch Ref No FF.ASMJ00094.1
Sample Name FF.ASMJ00094.1
Racemat
Lab Journal
Comment

#### Separation Information:

APC\Poeverlein\Poeverlein_2021
HPLC-System LC_03
Flow rate 1,0 ml/min
Temperature 30° C
HPLC Column Chiralpak IF/181, 250x4,6 mm
Eluent Hep:EtOH:MeOH 5:1:1 + 0,1% TFA

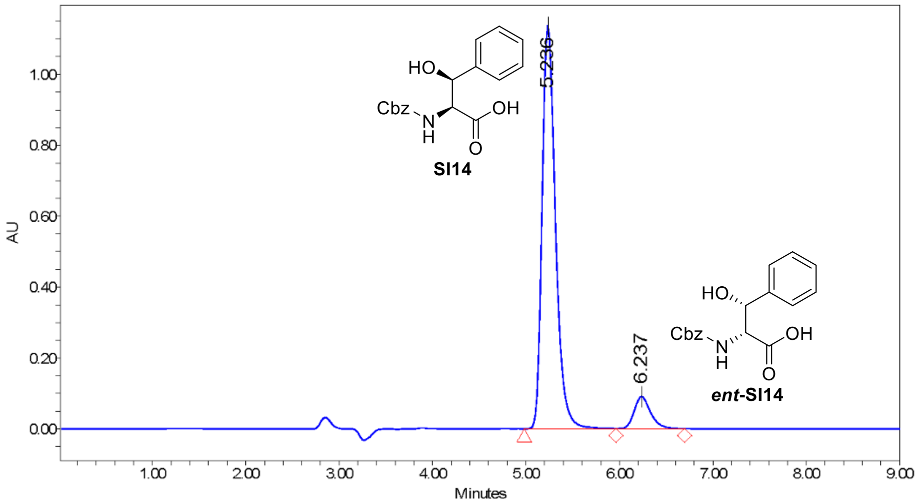

205.0nm; Inj ID 1590; Res ID 1596

Batch_Ref_No FF.ASMJ00094.1 Empower Project: APC\Poeverlein\Poeverlein_2021

##### Results

| | RT | Area | % Area | Height |
|---|---|---|---|---|
| 1 | 5.24 | 11346463 | 90.90 | 1137016 |
| 2 | 6.24 | 1135561 | 9.10 | 91076 |

Current Date 5/19/2021
Date Acquired 5/18/2021 2:47:15 PM CEST
Vial 30
Inj Vol 5.00 uL
1 of 2

647

#### Sample Information:

Customer Dr. Pöverlein

Batch Ref No FF.ASMJ00094.1

SampleName FF.ASMJ00094.1

Comment

#### Spectrum Index Plot

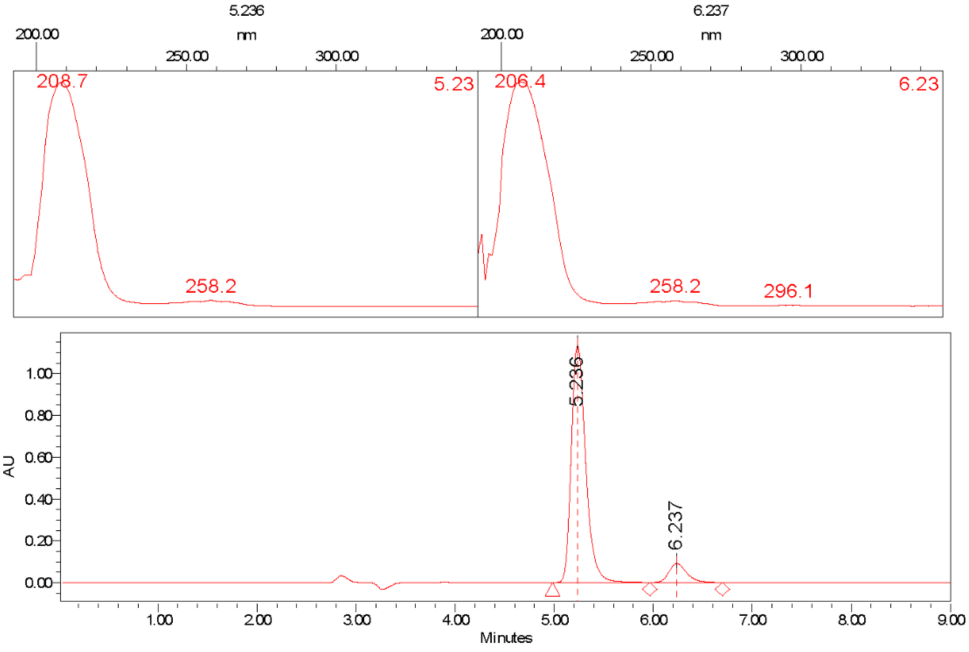

Batch_Ref_No FF.ASMJ00094.1 Project Name: APC\Poeverlein\Poeverlein_2021

Current Date 5/19/2021 Vial 30 2 of 2

Date Acquired 5/18/2021 2:47:15 PM CEST Inj Vol 5.00 uL

648

#### Chiral Separation

Library, SM, Chiral & Peptide Purification
R&D / IDD In vitro Biology & HT Chemistry
Industriepark Höchst, G 838, Room 007
D-65926 Frankfurt

SANOFI

##### Sample Information:

| | |
|---|---|
| Customer | Dr. Pöverlein |
| Analyst | K. Rahn-Hotze |
| Batch Ref No | FF.ASMJ00090.1 |
| Sample Name | FF.ASMJ00090.1 |
| Racemat | |
| Lab Journal | |
| Comment | |

##### Separation Information:

APC\Poeverlein\Poeverlein_2021

| | |
|---|---|
| HPLC-System | LC_03 |
| Flow rate | 1,0 ml/min |
| Temperature | 30° C |
| HPLC Column | Chiralpak IF/181, 250x4,6 mm |
| Eluent | Hep:EtOH:MeOH 5:1:1 + 0,1% TFA |

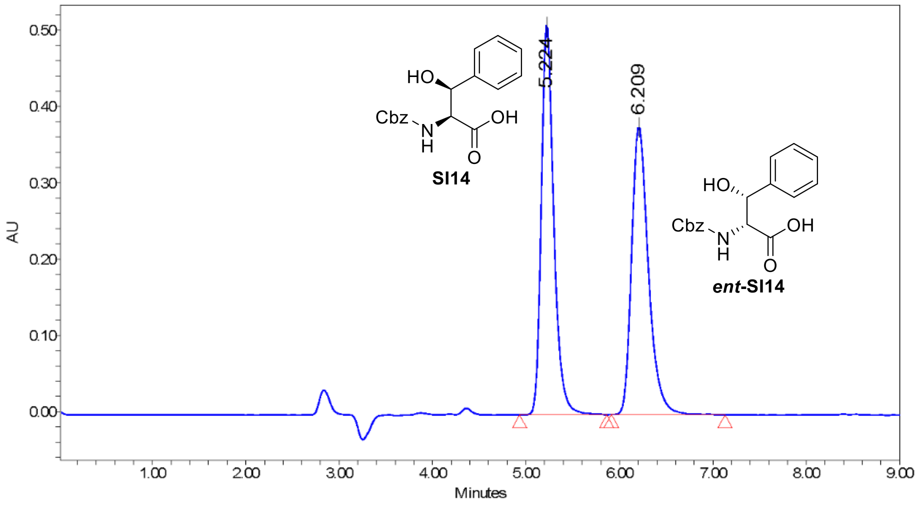

205.0nm; Inj ID 1581; Res ID 1597

Batch_Ref_No FF.ASMJ00090.1 Empower Project: APC\Poeverlein\Poeverlein_2021

##### Results

| | RT | Area | % Area | Height |
|---|---|---|---|---|
| 1 | 5.22 | 4759655 | 50.20 | 513452 |
| 2 | 6.21 | 4722672 | 49.80 | 378452 |

| | | | |
|---|---|---|---|
| Current Date | 5/19/2021 | Vial | 27 |
| Date Acquired | 5/18/2021 11:37:51 AM CEST | Inj Vol | 5.00 uL |

#### Sample Information:

Customer Dr. Pöverlein
Batch Ref No FF.ASMJ00090.1
SampleName FF.ASMJ00090.1
Comment

**Spectrum Index Plot**

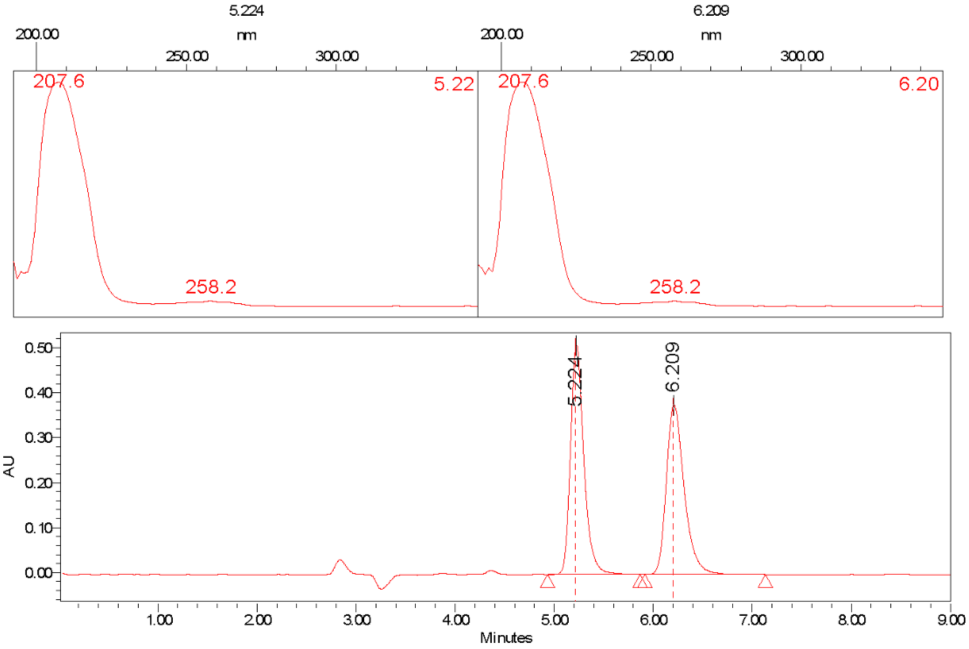

Batch_Ref_No FF.ASMJ00090.1 Project Name: APC\Poeverlein\Poeverlein_2021

Current Date 5/19/2021 Vial 27 2 of 2
Date Acquired 5/18/2021 11:37:51 AM CEST Inj Vol 5.00 uL

650

### Chiral Separation

Library, SM, Chiral & Peptide Purification
R&D / IDD In vitro Biology & HT Chemistry
Industriepark Höchst, G 838, Room 007
D-65926 Frankfurt

SANOFI

#### Sample Information:

| | |
|---|---|
| Customer | Dr. Pöverlein |
| Analyst | K. Rahn-Hotze |
| Batch Ref No | FF.ASMJ00088.1 |
| Sample Name | FF.ASMJ00088.1 |
| Racemat | |
| Lab Journal | |
| Comment | |

#### Separation Information:

APC\Poeverlein\Poeverlein_2021

| | |
|---|---|
| HPLC-System | LC_03 |
| Flow rate | 1,0 ml/min |
| Temperature | 30° C |
| HPLC Column | Chiralpak IF/181, 250x4,6 mm |
| Eluent | Hep:EtOH:MeOH 5:1:1 + 0,1% TFA |

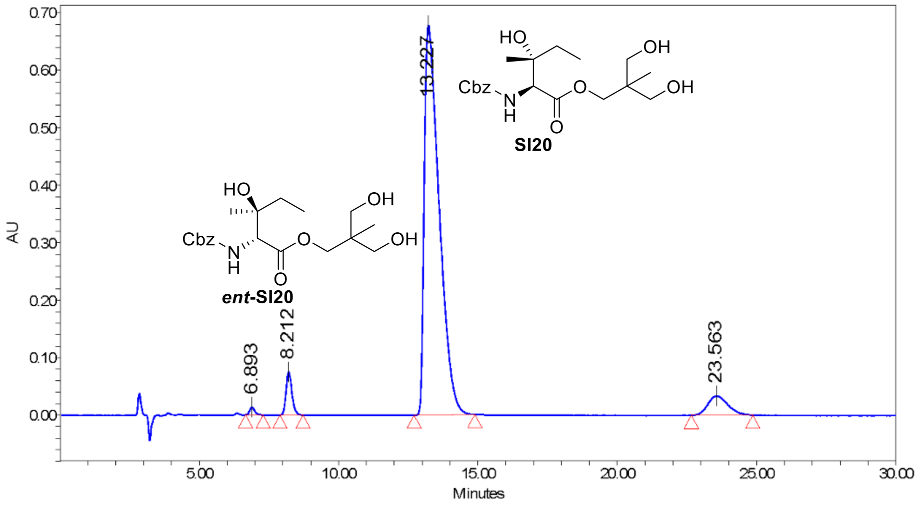

207.0nm; Inj ID 1674; Res ID 1681

Batch_Ref_No FF.ASMJ00088.1 Empower Project: APC\Poeverlein\Poeverlein_2021

##### Results

| | RT | Area | % Area | Height |
|---|---|---|---|---|
| 1 | 6.89 | 168888 | 0.60 | 13440 |
| 2 | 8.21 | 1103333 | 3.93 | 74819 |
| 3 | 13.23 | 25157668 | 89.52 | 677669 |
| 4 | 23.56 | 1671918 | 5.95 | 33362 |

Current Date 5/19/2021
Date Acquired 5/19/2021 9:06:21 AM CEST
Vial 37
Inj Vol 5.00 uL

#### Sample Information:

| | |
|---|---|
| Customer | Dr. Pöverlein |
| Batch Ref No | FF.ASMJ00088.1 |
| SampleName | FF.ASMJ00088.1 |
| Comment | |

#### Spectrum Index Plot

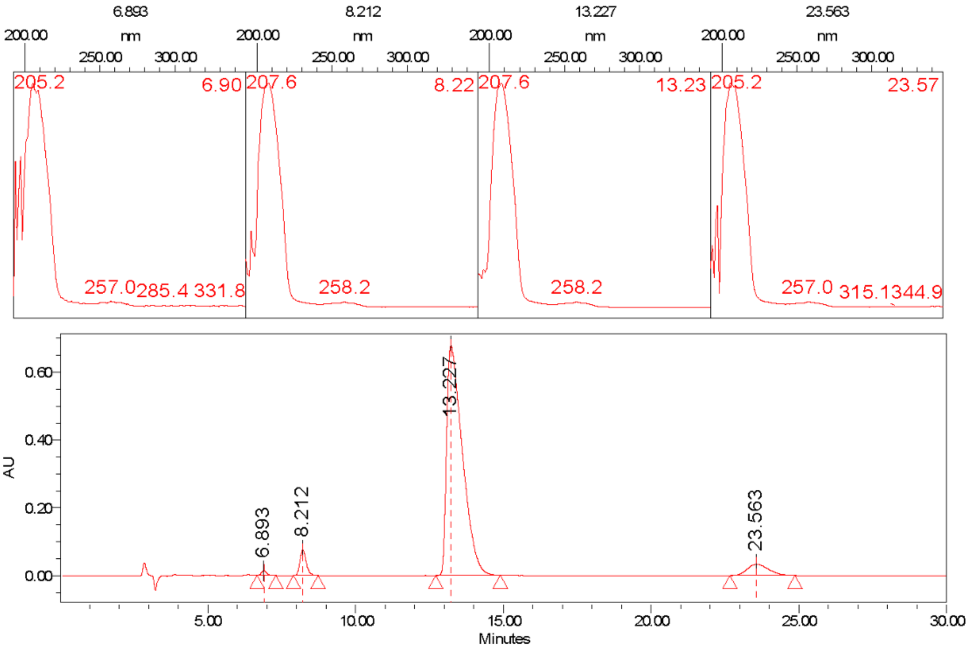

Batch_Ref_No FF.ASMJ00088.1

Project Name: APC\Poeverlein\Poeverlein_2021

| | | | |
|---|---|---|---|
| Current Date | 5/19/2021 | Vial 37 | 2 of 2 |
| Date Acquired | 5/19/2021 9:06:21 AM CEST | Inj Vol 5.00 uL | |

653

### Chiral Separation

Library, SM, Chiral & Peptide Purification
R&D / IDD In vitro Biology & HT Chemistry
Industriepark Höchst, G 838, Room 007
D-65926 Frankfurt

SANOFI

#### Sample Information:

Customer Dr. Pöverlein
Analyst K. Rahn-Hotze
Batch Ref No FF.ASMJ00079.1
Sample Name FF.ASMJ00079.1
Racemat
Lab Journal
Comment

#### Separation Information:

APC\Poeverlein\Poeverlein_2021
HPLC-System LC_03
Flow rate 1,0 ml/min
Temperature 30° C
HPLC Column Chiralpak IF/181, 250x4,6 mm
Eluent Hep:EtOH:MeOH 5:1:1 + 0,1% TFA

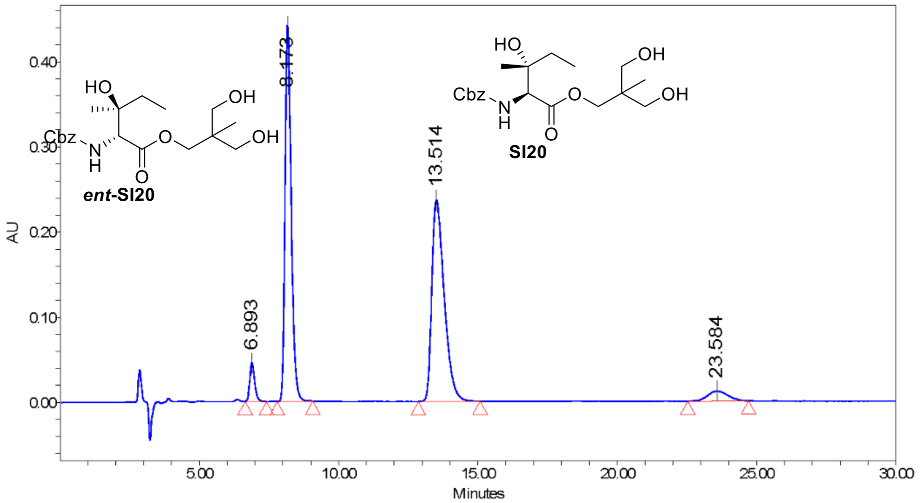

207.0nm; Inj ID 1667; Res ID 1676

Batch_Ref_No FF.ASMJ00079.1 Empower Project: APC\Poeverlein\Poeverlein_2021

##### Results

| | RT | Area | % Area | Height |
|---|---|---|---|---|
| 1 | 6.89 | 557454 | 3.65 | 45324 |
| 2 | 8.17 | 6794692 | 44.55 | 442018 |
| 3 | 13.51 | 7308184 | 47.92 | 236514 |
| 4 | 23.58 | 591832 | 3.88 | 11880 |

Current Date 5/19/2021 Vial 35 1 of 2
Date Acquired 5/19/2021 8:04:37 AM CEST Inj Vol 5.00 uL

654

#### Sample Information:

Customer Dr. Pöverlein
Batch Ref No FF.ASMJ00079.1
SampleName FF.ASMJ00079.1
Comment

##### Spectrum Index Plot

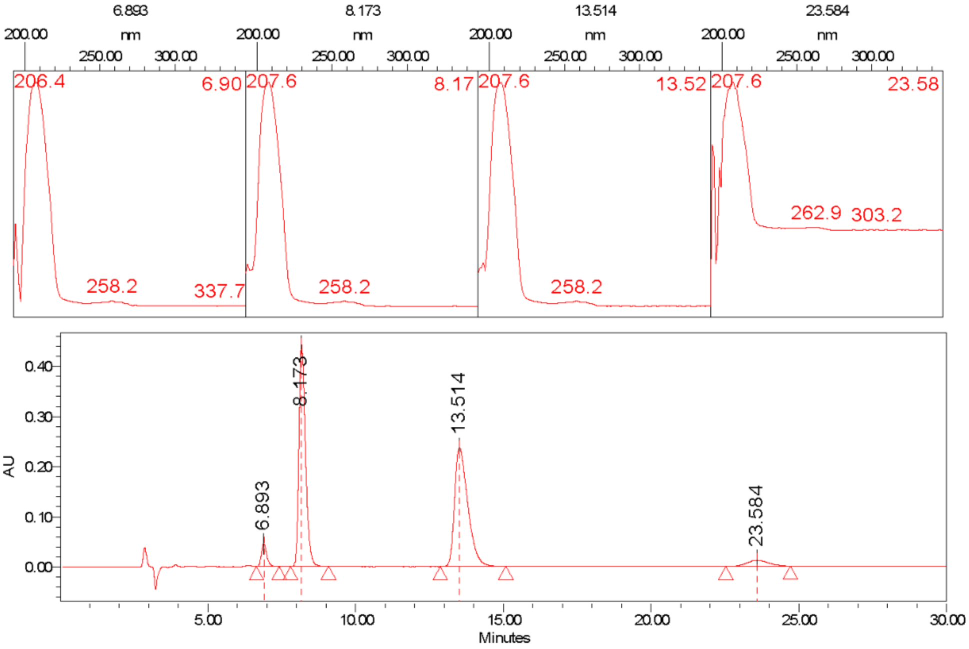

Batch_Ref_No FF.ASMJ00079.1 Project Name: APC\Poeverlein\Poeverlein_2021

#### Chiral Separation

Library, SM, Chiral & Peptide Purification
R&D / IDD In vitro Biology & HT Chemistry
Industriepark Höchst, G 838, Room 007
D-65926 Frankfurt

SANOFI

#### Sample Information:

Customer Dr. Pöverlein
Analyst K. Rahn-Hotze
Batch Ref No FF.ASMJ00089.1
Sample Name FF.ASMJ00089.1
Racemat
Lab Journal
Comment

#### Separation Information:

APC\Poeverlein\Poeverlein_2021
HPLC-System LC_03
Flow rate 1,0 ml/min
Temperature 30° C
HPLC Column Chiralpak IF/181, 250x4,6 mm
Eluent Hep:EtOH:MeOH 5:1:1 + 0,1% TFA

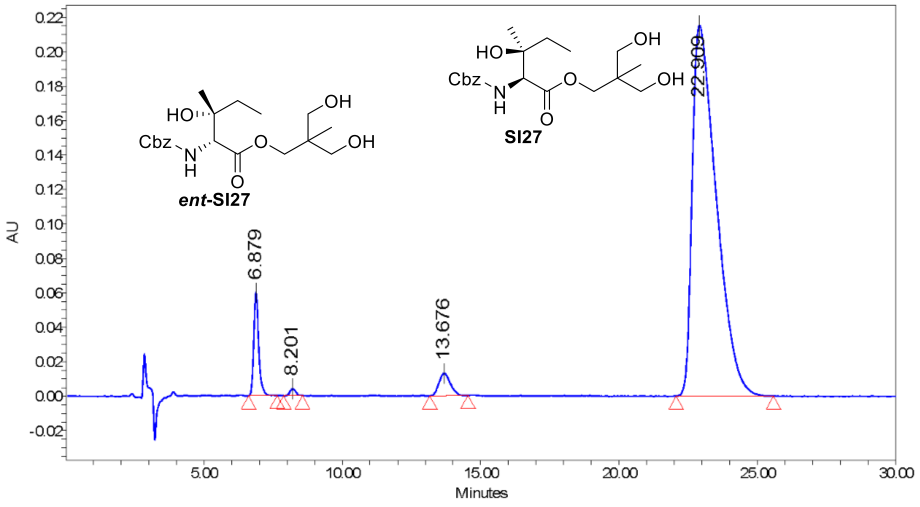

Batch_Ref_No FF.ASMJ00089.1 Empower Project: APC\Poeverlein\Poeverlein_2021

##### Results

| | RT | Area | % Area | Height |
|---|---|---|---|---|
| 1 | 6.88 | 743457 | 5.24 | 60011 |
| 2 | 8.20 | 57360 | 0.40 | 3936 |
| 3 | 13.68 | 370942 | 2.62 | 12923 |
| 4 | 22.91 | 13004059 | 91.73 | 215289 |

Current Date 5/19/2021
Date Acquired 5/19/2021 11:12:48 AM CEST
Vial 38
Inj Vol 3.00 uL
1 of 2

656

#### Sample Information:

Customer Dr. Pöverlein
Batch Ref No FF.ASMJ00089.1
SampleName FF.ASMJ00089.1
Comment

##### Spectrum Index Plot

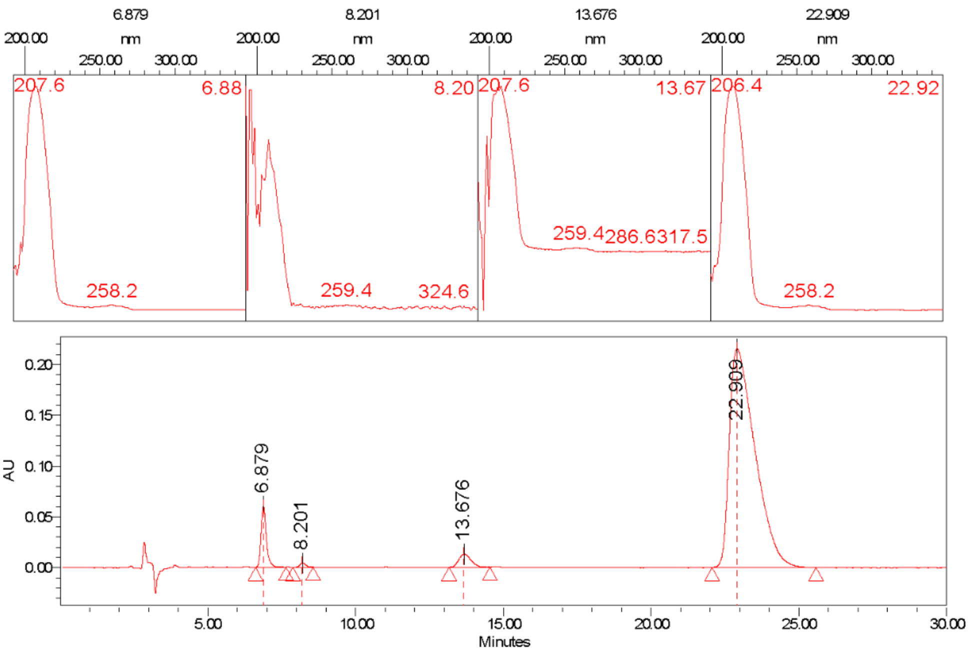

Batch_Ref_No FF.ASMJ00089.1 Project Name: APC\Poeverlein\Poeverlein_2021

#### Chiral Separation

Library, SM, Chiral & Peptide Purification
R&D / IDD In vitro Biology & HT Chemistry
Industriepark Höchst, G 838, Room 007
D-65926 Frankfurt

SANOFI

##### Sample Information:

| | |
|---|---|
| Customer | Dr. Pöverlein |
| Analyst | K. Rahn-Hotze |
| Batch Ref No | FF.ASMJ00080.1 |
| Sample Name | FF.ASMJ00080.1 |
| Racemat | |
| Lab Journal | |
| Comment | |

##### Separation Information:

APC\Poeverlein\Poeverlein_2021

| | |
|---|---|
| HPLC-System | LC_03 |
| Flow rate | 1,0 ml/min |
| Temperature | 30° C |
| HPLC Column | Chiralpak IF/181, 250x4,6 mm |
| Eluent | Hep:EtOH:MeOH 5:1:1 + 0,1% TFA |

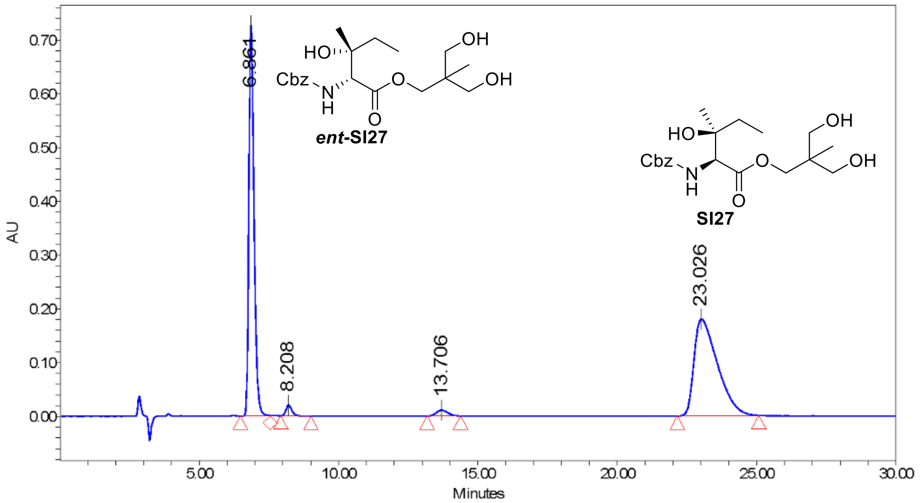

207.0nm; Inj ID 1671; Res ID 1677

Batch_Ref_No FF.ASMJ00080.1 Empower Project: APC\Poeverlein\Poeverlein_2021

##### Results

| | RT | Area | % Area | Height |
|---|---|---|---|---|
| 1 | 6.86 | 9352186 | 45.72 | 727535 |
| 2 | 8.21 | 294051 | 1.44 | 19947 |
| 3 | 13.71 | 306613 | 1.50 | 10859 |
| 4 | 23.03 | 10503751 | 51.35 | 179849 |

Current Date 5/19/2021
Date Acquired 5/19/2021 8:35:28 AM CEST

Vial 36
Inj Vol 5.00 uL

#### Sample Information:

Customer Dr. Pöverlein
Batch Ref No FF.ASMJ00080.1
SampleName FF.ASMJ00080.1
Comment

**Spectrum Index Plot**

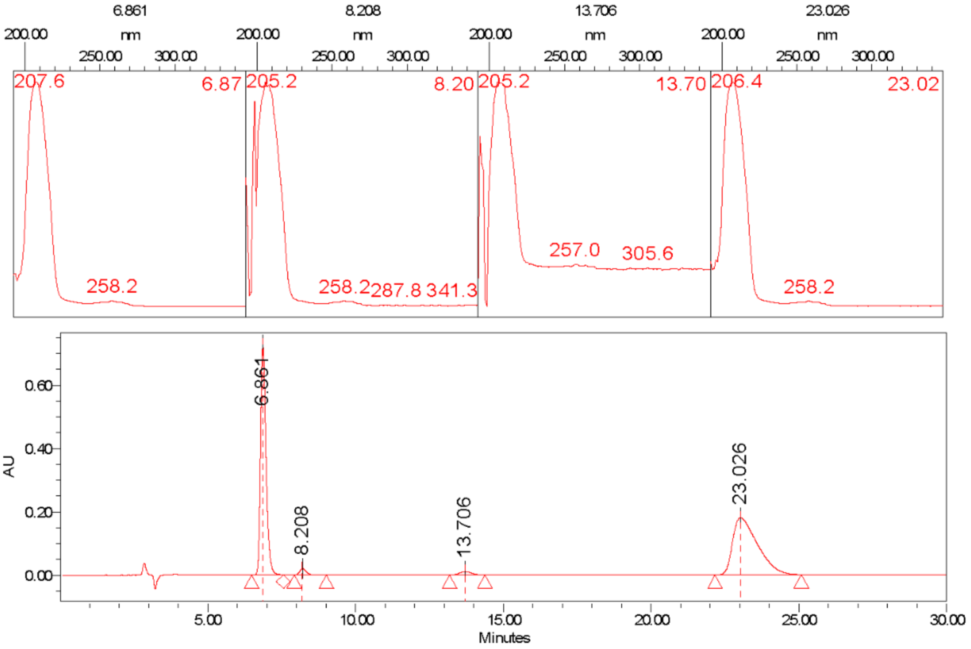

Batch_Ref_No FF.ASMJ00080.1 Project Name: APC\Poeverlein\Poeverlein_2021

Current Date 5/19/2021 Vial 36 2 of 2
Date Acquired 5/19/2021 8:35:28 AM CEST Inj Vol 5.00 uL

#### Chiral Separation

Library, SM, Chiral & Peptide Purification
R&D / IDD In vitro Biology & HT Chemistry
Industriepark Höchst, G 838, Room 007
D-65926 Frankfurt

SANOFI

#### Sample Information:

| | |
|---|---|
| Customer | Dr. Pöverlein |
| Analyst | K. Rahn-Hotze |
| Batch Ref No | FF.ASMJ00096.1 |
| Sample Name | FF.ASMJ00096.1 |
| Racemat | |
| Lab Journal | |
| Comment | |

#### Separation Information:

APC\Poeverlein\Poeverlein_2021

| | |
|---|---|
| HPLC-System | LC_03 |
| Flow rate | 1,0 ml/min |
| Temperature | 30° C |
| HPLC Column | Chiralpak IF/181, 250x4,6 mm |
| Eluent | Hep:EtOH 10:1 + 0,1% TFA |

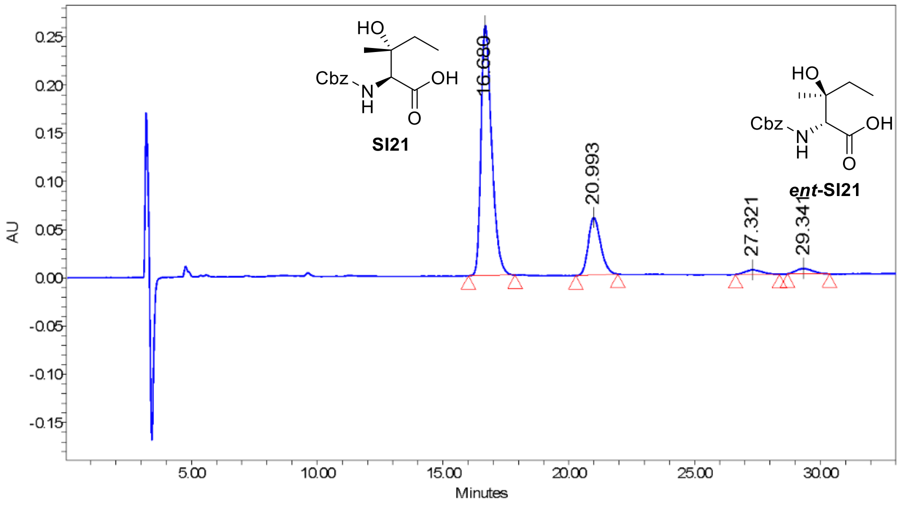

Batch_Ref_No FF.ASMJ00096.1 Empower Project: APC\Poeverlein\Poeverlein_2021

##### Results

| | RT | Area | % Area | Height |
|---|---|---|---|---|
| 1 | 16.68 | 7643383 | 75.21 | 259347 |
| 2 | 20.99 | 2040213 | 20.07 | 59600 |
| 3 | 27.32 | 219056 | 2.16 | 4734 |
| 4 | 29.34 | 260346 | 2.56 | 5668 |

Current Date 6/8/2021 Vial 44 1 of 2
Date Acquired 6/8/2021 10:08:03 AM CEST Inj Vol 5.00 uL

662

663

#### Sample Information:

| | |
|---|---|
| Customer | Dr. Pöverlein |
| Batch Ref No | FF.ASMJ00096.1 |
| SampleName | FF.ASMJ00096.1 |
| Comment | |

##### Spectrum Index Plot

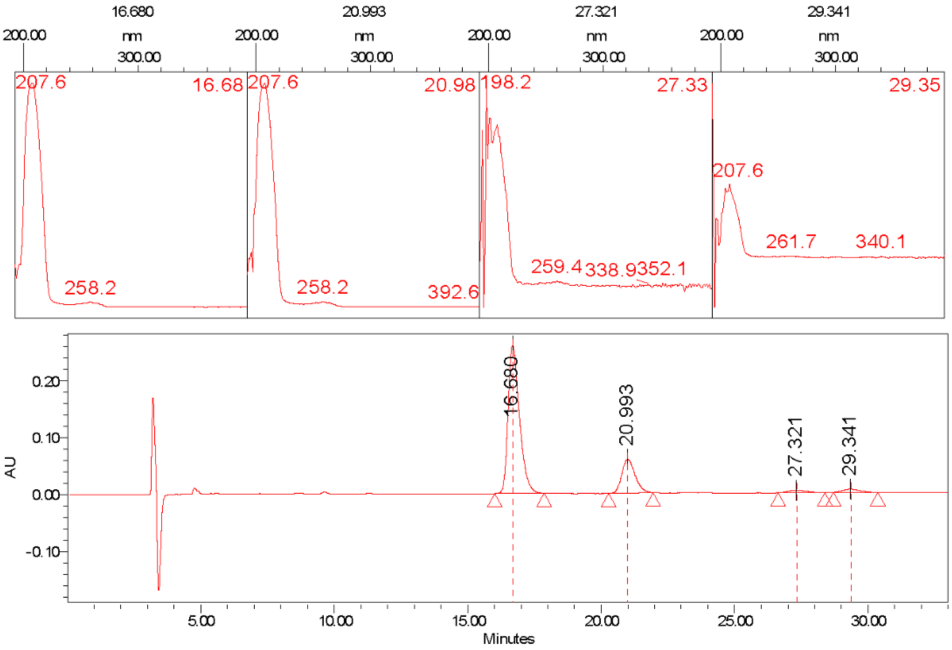

Batch_Ref_No FF.ASMJ00096.1 Project Name: APC\Poeverlein\Poeverlein_2021

| | | | |
|---|---|---|---|
| Current Date | 6/8/2021 | Vial 44 | 2 of 2 |
| Date Acquired | 6/8/2021 10:08:03 AM CEST | Inj Vol 5.00 uL | |

#### Chiral Separation

Library, SM, Chiral & Peptide Purification
R&D / IDD In vitro Biology & HT Chemistry
Industriepark Höchst, G 838, Room 007
D-65926 Frankfurt

SANOFI

##### Sample Information:

Customer Dr. Pöverlein
Analyst K. Rahn-Hotze
Batch Ref No FF.ASMJ00092.1
Sample Name FF.ASMJ00092.1
Racemat
Lab Journal
Comment

##### Separation Information:

APC\Poeverlein\Poeverlein_2021
HPLC-System LC_03
Flow rate 1,0 ml/min
Temperature 30° C
HPLC Column Chiralpak IF/181, 250x4,6 mm
Eluent Hep:EtOH 10:1 + 0,1% TFA

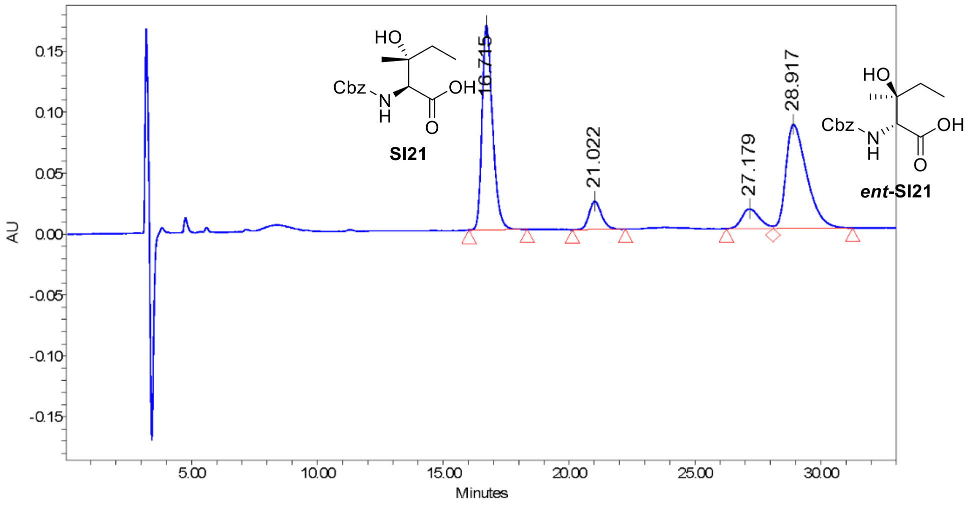

208.0nm; Inj ID 1774; Res ID 1783

Batch_Ref_No FF.ASMJ00092.1 Empower Project: APC\Poeverlein\Poeverlein_2021

##### Results

| | RT | Area | % Area | Height |
|---|---|---|---|---|
| 1 | 16.72 | 4876070 | 42.95 | 167660 |
| 2 | 21.02 | 803425 | 7.08 | 23173 |
| 3 | 27.18 | 787898 | 6.94 | 16162 |
| 4 | 28.92 | 4884366 | 43.03 | 85118 |

Current Date 6/8/2021 Vial 42 1 of 2
Date Acquired 6/8/2021 9:00:19 AM CEST Inj Vol 5.00 uL

#### Sample Information:

| | |
|---|---|
| Customer | Dr. Pöverlein |
| Batch Ref No | FF.ASMJ00092.1 |
| SampleName | FF.ASMJ00092.1 |
| Comment | |

##### Spectrum Index Plot

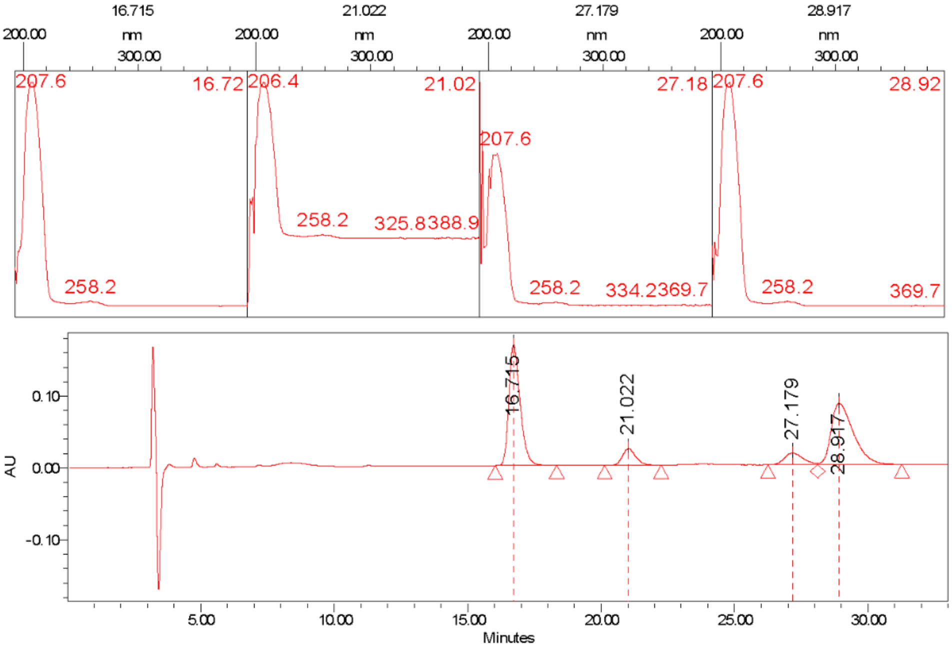

Batch_Ref_No FF.ASMJ00092.1 Project Name: APC\Poeverlein\Poeverlein_2021

Current Date 6/8/2021 Vial 42 2 of 2
Date Acquired 6/8/2021 9:00:19 AM CEST Inj Vol 5.00 uL

### Chiral Separation

Library, SM, Chiral & Peptide Purification
R&D / IDD In vitro Biology & HT Chemistry
Industriepark Höchst, G 838, Room 007
D-65926 Frankfurt

SANOFI

#### Sample Information:

Customer Dr. Pöverlein
Analyst K. Rahn-Hotze
Batch Ref No FF.ASMJ00097.1
Sample Name FF.ASMJ00097.1
Racemat
Lab Journal
Comment

#### Separation Information:

APC\Poeverlein\Poeverlein_2021
HPLC-System LC_03
Flow rate 1,0 ml/min
Temperature 30° C
HPLC Column Chiralpak IF/181, 250x4,6 mm
Eluent Hep:EtOH 10:1 + 0,1% TFA

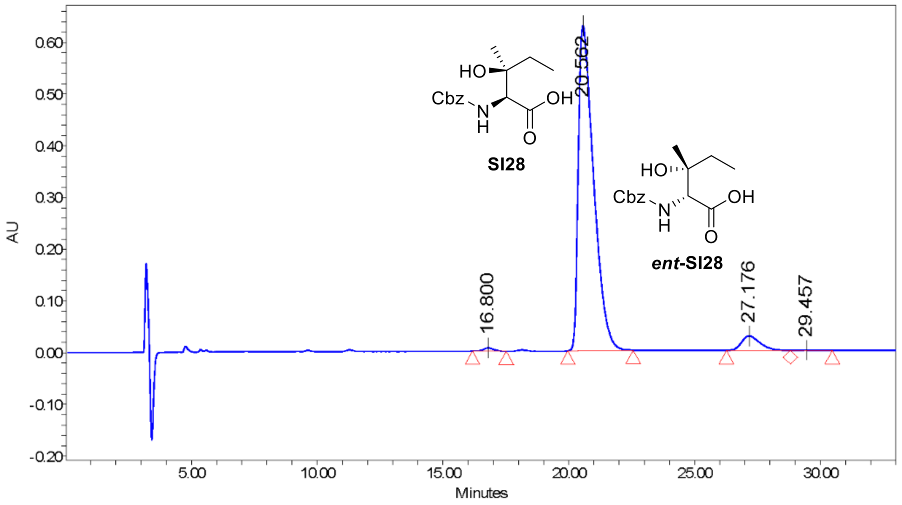

208.0nm; Inj ID 1786; Res ID 1793

Batch_Ref_No FF.ASMJ00097.1 Empower Project: APC\Poeverlein\Poeverlein_2021

##### Results

| | RT | Area | % Area | Height |
|---|---|---|---|---|
| 1 | 16.80 | 165940 | 0.59 | 6167 |
| 2 | 20.56 | 26569318 | 94.16 | 629039 |
| 3 | 27.18 | 1444362 | 5.12 | 28117 |
| 4 | 29.46 | 37232 | 0.13 | 730 |

Current Date 6/8/2021
Date Acquired 6/8/2021 10:41:54 AM CEST
Vial 45
Inj Vol 5.00 uL
1 of 2

#### Sample Information:

Customer Dr. Pöverlein

Batch Ref No FF.ASMJ00097.1

SampleName FF.ASMJ00097.1

Comment

**Spectrum Index Plot**

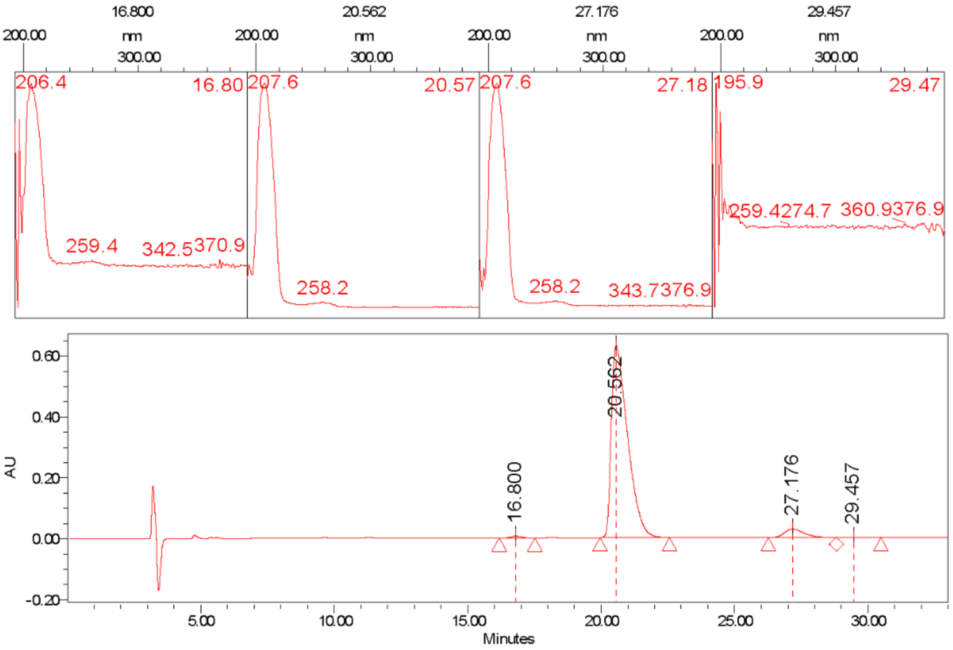

Batch_Ref_No FF.ASMJ00097.1 Project Name: APC\Poeverlein\Poeverlein_2021

| | | |
|---|---|---|
| Current Date 6/8/2021 | Vial 45 | 2 of 2 |
| Date Acquired 6/8/2021 10:41:54 AM CEST | Inj Vol 5.00 uL | |

#### Chiral Separation

Library, SM, Chiral & Peptide Purification
R&D / IDD In vitro Biology & HT Chemistry
Industriepark Höchst, G 838, Room 007
D-65926 Frankfurt

SANOFI

#### Sample Information:

Customer Dr. Pöverlein
Analyst K. Rahn-Hotze
Batch Ref No FF.ASMJ00093.1
Sample Name FF.ASMJ00093.1
Racemat
Lab Journal
Comment

#### Separation Information:

APC\Poeverlein\Poeverlein_2021
HPLC-System LC_03
Flow rate 1,0 ml/min
Temperature 30° C
HPLC Column Chiralpak IF/181, 250x4,6 mm
Eluent Hep:EtOH 10:1 + 0,1% TFA

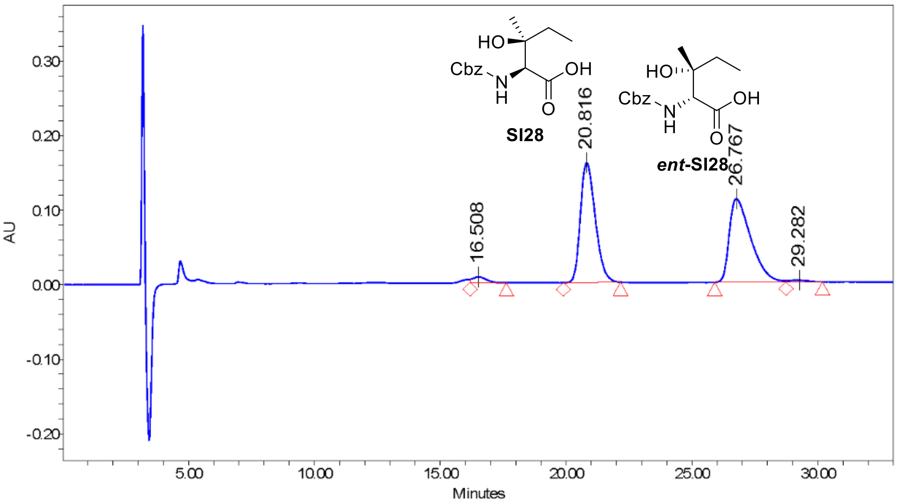

208.0nm; Inj ID 1791; Res ID 1794

Batch_Ref_No FF.ASMJ00093.1 Empower Project: APC\Poeverlein\Poeverlein_2021

##### Results

| | RT | Area | % Area | Height |
|---|---|---|---|---|
| 1 | 16.51 | 288213 | 2.11 | 7812 |
| 2 | 20.82 | 6602833 | 48.38 | 160649 |
| 3 | 26.77 | 6642495 | 48.67 | 111814 |
| 4 | 29.28 | 115445 | 0.85 | 2267 |

Current Date 6/8/2021 Vial 43 1 of 2
Date Acquired 6/8/2021 11:17:59 AM CEST Inj Vol 15.00 uL

674

675

#### Sample Information:

Customer Dr. Pöverlein

Batch Ref No FF.ASMJ00093.1

SampleName FF.ASMJ00093.1

Comment

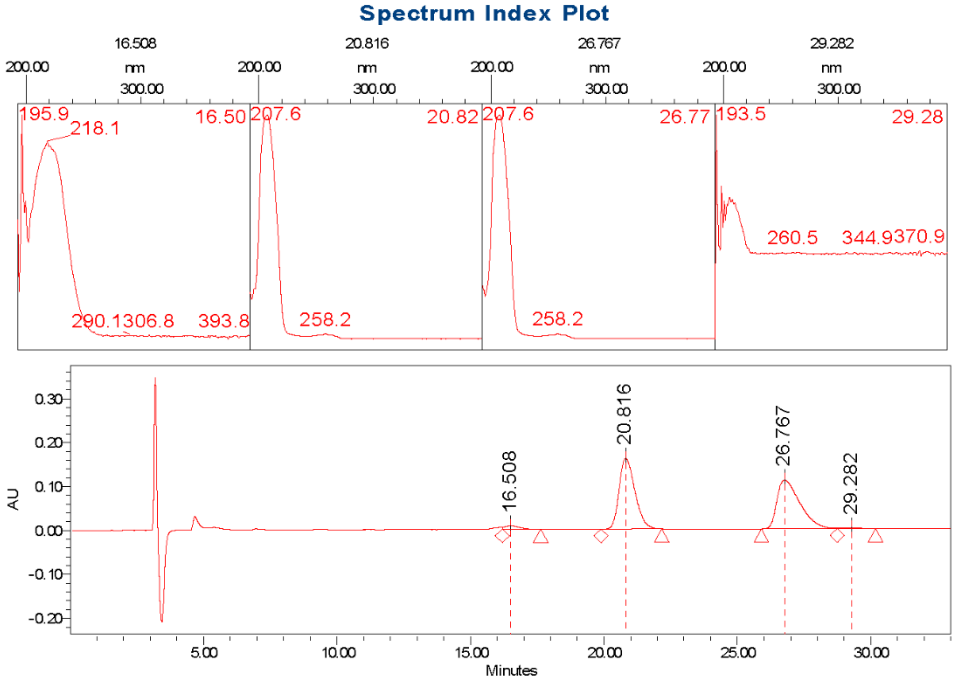

Batch_Ref_No FF.ASMJ00093.1 Project Name: APC\Poeverlein\Poeverlein_2021

Current Date 6/8/2021 Vial 43 2 of 2

Date Acquired 6/8/2021 11:17:59 AM CEST Inj Vol 15.00 uL

676
